# Biological evaluation of novel side chain containing CQTrICh-analogs as antimalarials and their development as *Pf*CDPK1 kinase inhibitors

**DOI:** 10.1101/2022.07.07.498981

**Authors:** Iram Irfan, Amad Uddin, Ravi Jain, Aashima Gupta, Sonal Gupta, John V. Napoleon, Afzal Hussain, Mohamed F. Alajmi, Mukesh C. Joshi, Phool Hasan, Mohammad Abid, Shailja Singh

## Abstract

To combat the emergence of drug resistance against the existing antimalarials, novel side chain containing 7-chloroquinoline-indole-chalcones tethered with a triazole (CQTrICh-analogs **7 (a-s)** and **9)** were designed and synthesized by reacting substituted 1-phenyl-3-(1-(prop-2-yn-1- yl)-1H-indol-3-yl) prop-2-en-1-one and 1-(prop-2-yn-1-yl)-1H-indole-3-carbaldehyde with 4- azido-7-chloroquinoline, respectively via a ‘click’ reaction. The selected CQTrICh-analogs: **7l** and **7r** inhibited chloroquine-sensitive (3D7) and resistant (RKL-9) strains of *Plasmodium falciparum*, with IC_50_ values of **2.4 µM** & **1.8 µM** (**7l**), and **3.5 µM** & **2.7 µM** (**7r**), respectively, and showed insignificant hemolysis and cytotoxicity in mammalian cells. Intra-erythrocytic progression studies revealed that the active hybrids: **7l** and **7r** are effective against the mature stages of the parasite. Given the importance of Calcium-Dependent Protein Kinase 1 (*Pf*CDPK1) in the parasite biology, notably during late schizogony and subsequent invasion of merozoites into host RBCs, we identified this protein as a possible molecular target of these active hybrids. *In silico* interaction analysis indicated that **7l** and **7r** stably interact with the catalytically active ATP-binding pocket of *Pf*CDPK1, by the formation of energetically favorable H-bonds. Furthermore, *in vitro* Microscale Thermophoresis and kinase assays with recombinant *Pf*CDPK1 demonstrated that the active hybrids interact with and inhibit the kinase activity, thus presumably responsible for the parasite growth inhibition. Interestingly, **7l** and **7r** showed no inhibitory effect on the human kinases, indicating that they are selective for the parasite kinase. Conceivably, we report the antiplasmodial potential of novel kinase targeting bio-conjugates, a step towards developing pan-kinase inhibitors, which is a prerequisite for cross-stage anti-malarial protection.

**GRAPHICAL ABSTRACT:** 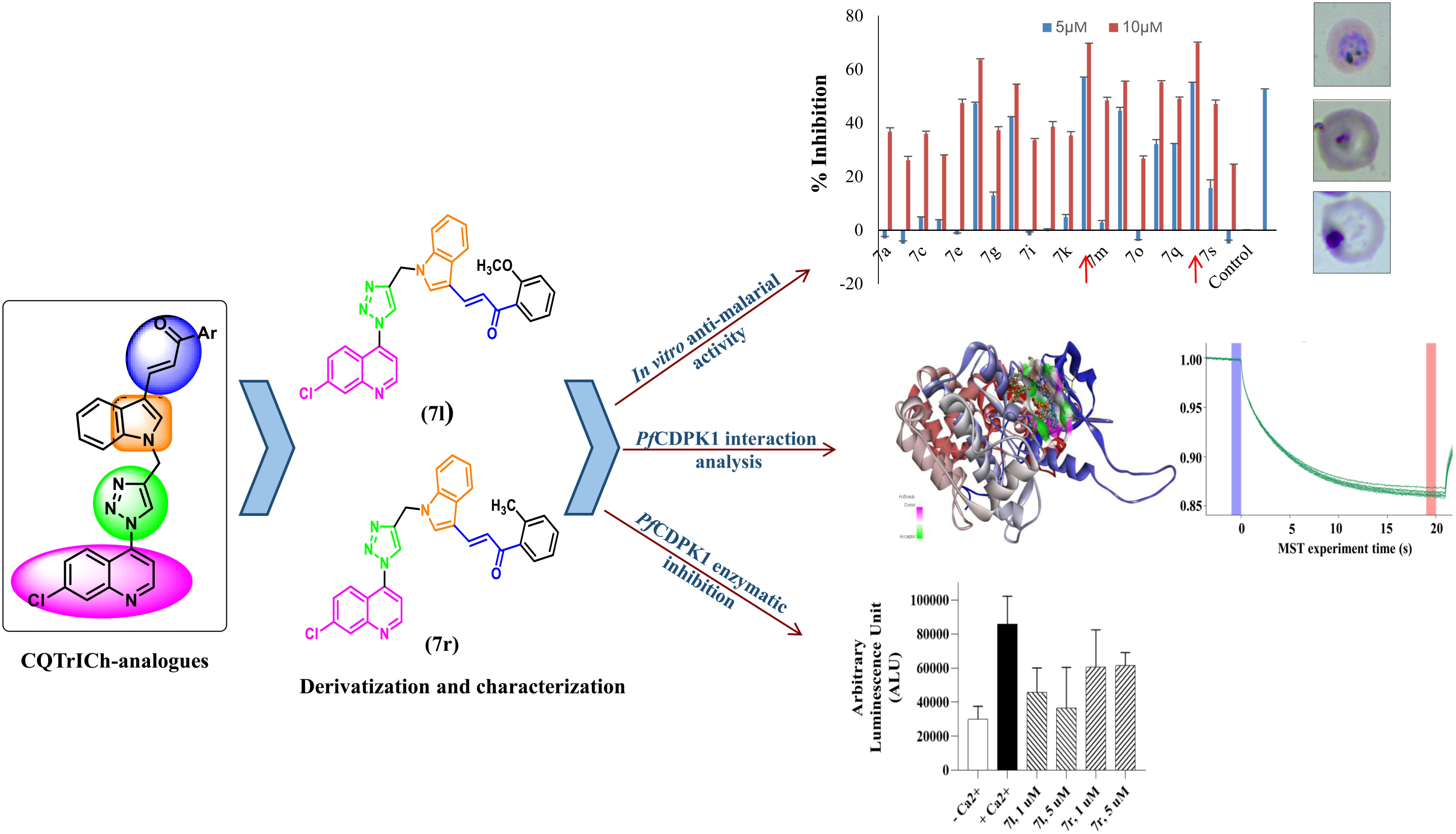

## 1. Introduction

Side chains containing quinoline analogs are an important class of antimalarial drugs in the history of antimalarials. In fact, these analogs compromised by the resistance, remain a choice of interest to make better novel side chains containing antimalarials [1]. Thus, novel anti-malarial resistance combating approaches and low-cost new drugs with favorable safety profiles are urgently needed [2–5]. Recently, the WHO report in the 2021 edition stated 241 million malaria cases whereas an estimated of 627,000 deaths worldwide in 2020 [6]. It is, therefore, crucial to develop effective anti-malarial medications that identify new targets, particularly those with novel mechanisms of action and toxicity profiles to cure the lethal and pandemic malaria parasite. The side chain containing aminoquinolines (*viz.* chloroquine) is well known for its inhibition of crystallization of Fe(III) heam to β-hematin and hemozoin etc. [7,8]. Numerous studies have shown that side-chain alteration and modifications may lead to increased antimalarial activity against various *P. falciparum* strains [9–14].

Scientists are engaged in the search for possible mechanistic approaches to find out how various antimalarials inhibit the *Plasmodium* parasite at different stages of its lifecycle, but the question remains a big task. Thus, several groups are working to design and develop small molecular probes by applying various synthetic and biological approaches to understand their mode of action in the lifecycle of the parasite. *Pf*CDPK1 kinase is one of the biological targets which may act as a vital target for the incoming probes. In the genus *Plasmodium*, Calcium-Dependent Protein Kinases (CDPKs) represent a class of serine/threonine protein kinases that are *activated* by Ca^2+^-ions, belonging to a multigene family comprised of seven members, and each gene is predominantly expressed in a distinct phase of the parasite life cycle. Since they aren’t found in mammals, they could be used as safer pharmacological targets. Previous reports demonstrate the relevance of CDPK1 in *all* asexual intra-erythrocytic stages of the parasite. It can regulate actin-myosin motor during gliding and erythrocyte invasion of the parasite; and, regulate the cAMP-mediated signaling cascade module [15]. With this understanding of the *indispensable* role of CDPK1 in the parasite life cycle, we pursued this as a potential target for the development of novel anti-malarial inhibitors that can block parasite growth by targeting *multiple* stages in malarial infection.

Earlier attempts have been made to identify specific inhibitors against *Pf*CDPK1 protein, including purfalcamine, imidazopyridazines (**A** or 2-amino-3-[(4-methylphenyl) carbonyl]-indolizine-1-carboxamide), imidazopyridine analogs, and a series of inhibitors, identified by an *in vitro* biochemical screening approach against chemical libraries of small molecules against recombinant *Pf*CDPK1 whereas **A** exhibited promising *in vitro* anti-parasite activity, suitable ADME properties and modest *in vivo* efficacy in the mouse model [16–20]. In **Fig. 1**, triazole-linked chalcone **B** [21] and CQ-triazole-linked chalcones **C** [22], and indole-triazole hybrids (**D**) [23], were developed and evaluated for their antiplasmodial activity, whereas indole containing core in indolizines **A** were studied for their *Pf*CDPK1 inhibition. Few other research groups have worked in this field as well [22,24–26]. Based on these active pharmacophore core units *viz.* quinoline, triazole, indole, and chalcones, we have successfully designed and developed a side chain containing novel anti-malarial CQTrICh-analogs (**7a-s)** and **9** and their possible *Pf*CDPK1 inhibitory potential. Here, we reported the synthesis, characterization, ADME properties, and dose-dependent *in vitro* antimalarial activity against the CQ*^S^*-3D7 strain of *P. falciparum*, however, two selective analogs (**7l** and **7r**) were screened for their anti-plasmodial IC_50_ value against CQ*^S^*-3D7 as well as CQ*^R^*-RKL9 strains of *P. falciparum*, *Pf*CDPK-1 inhibition, hematin inhibition, growth progression analysis, cytotoxic properties, and molecular docking studies. Overall, **7l** and **7r** exhibited promising results towards antimalarial activity against both the tested strains and found as *Pf*CDPK1 kinase inhibitors. This study opens the door to making more bioisosteres towards the development of inhibitors of *Pf*CDPK1 kinase.

**Fig. 1.**
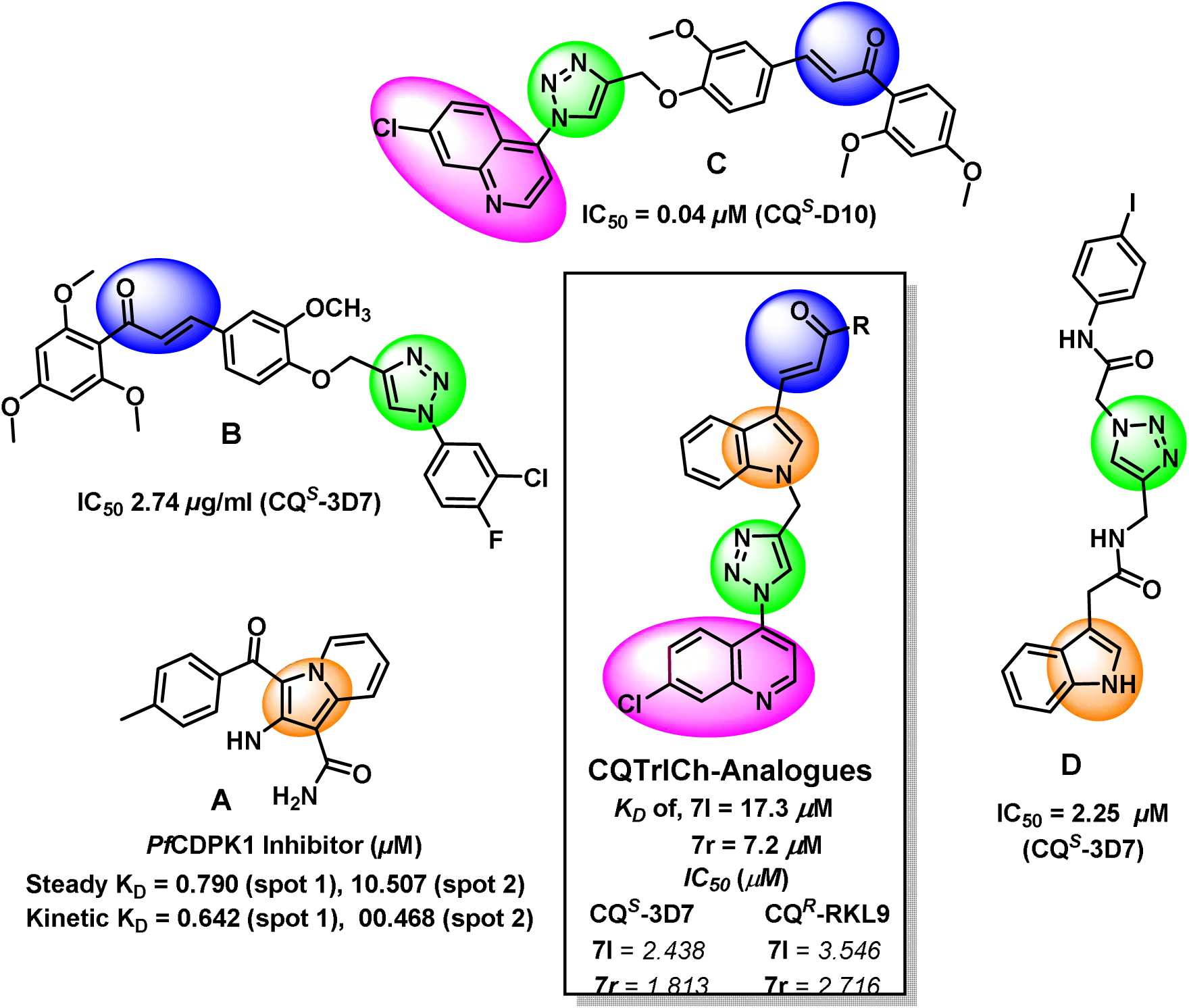
Rationale for the design and development of CQTrICh-analogs

## 2. Chemistry

Synthesis of CQTrICh-analogs; **7a-s** and **9** were accomplished by an efficient synthetic approach illustrated in **Scheme 1**. Indole-3-aldehyde (**1,** 1.0 equiv.) on condensation with desired aryl ketones (**2a-s,** 1.0 equiv.) under reflux condition in anhyd. ethanol for 30 h to give indole-chalcones (**3a-s**) in good yield. The indole-chalcone intermediates (**3a-s**) were further propargylated using propargyl bromide (2.0 equiv.) and K_2_CO_3_ (2.5 equiv.) in DMF at 0°C then the reaction mixture was allowed to stir at room temperature for additional 23 h to produce the corresponding alkynes (**4a-s)** in excellent yield. Simultaneously, azide (**6**) [27,28] was prepared from 4,7-dichloroquinoline (CQ, **5**) by direct azide replacement using NaN_3_ (6.0 equiv.) in anhydrous DMF at 85 °C for 3 h. Then, corresponding alkyne intermediates (**4a-s**) were treated with azide **6** (1 equiv.) at room temperature for 20-24 h under click reaction [29] in presence of THF:H_2_O (1:2) mixture, sodium ascorbate (0.5 equiv.) and CuSO_4_.5H_2_O (0.16 equiv.) as a catalyst to afford CQTrICh analogs (**7a-s**) in moderate to good yield (53-91%). Similarly, we also synthesized compound **9** by treating 1-(prop-2-ynyl)-1H-indole-3-carbaldehyde (**8**) with azide **6** at room temperature for 20 h under Click reaction in presence of THF:H_2_O (1:2) mixture, sodium ascorbate (0.5 equiv.) and CuSO_4_.5H_2_O (0.16 equiv.) as a catalyst.

**Scheme 1.**
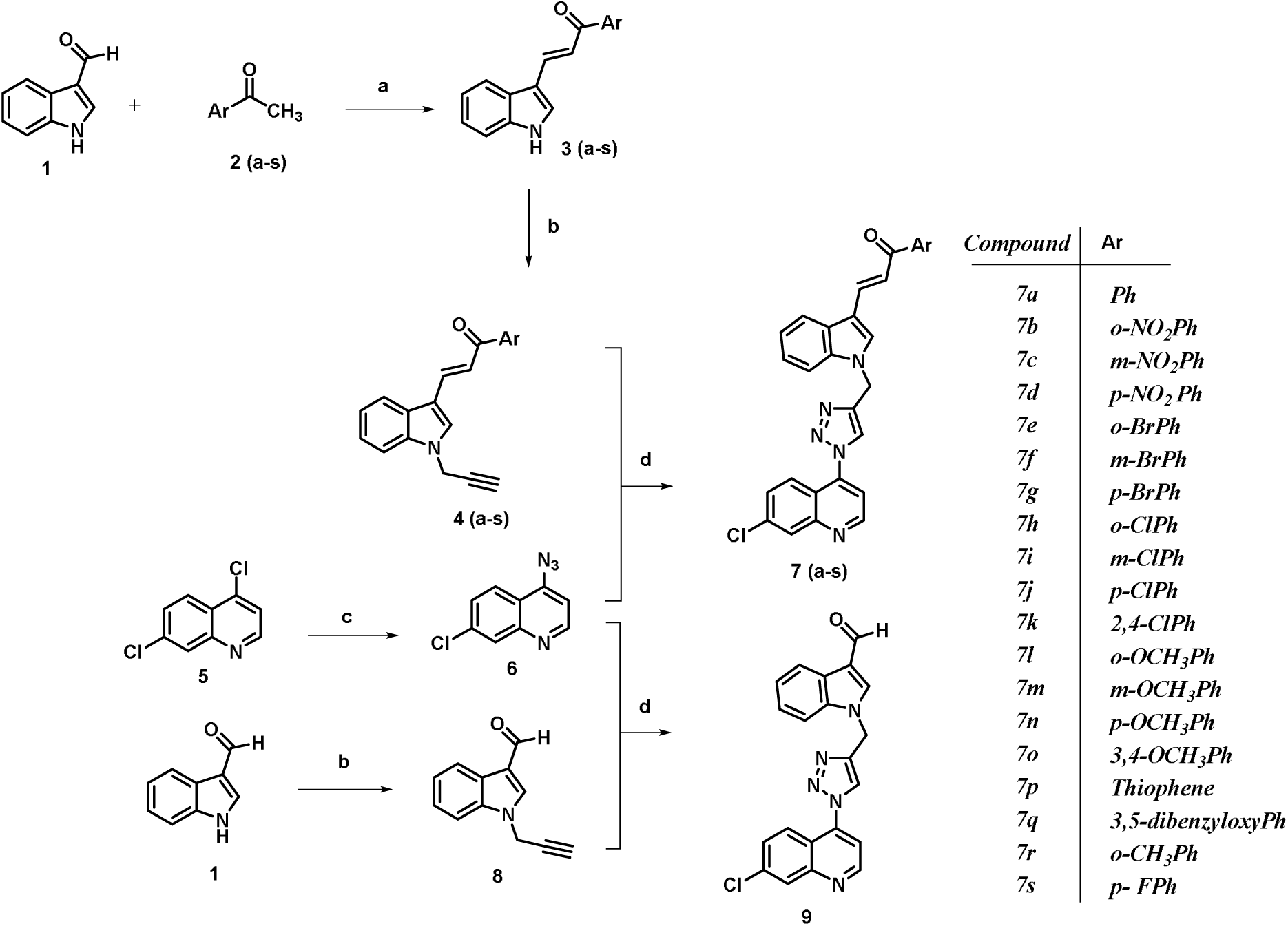
Synthesis of CQTrICh analogs (**7a-s)** and **9**. Reagents and conditions: (a) ethanol, reflux, 30 h; (b) propargyl bromide, K_2_CO_3_, DMF, 0 °C–rt, 23 h; (c) NaN_3_, DMF, 85 °C, 3 h; (d) CuSO_4_.5H_2_O, sodium ascorbate, THF/H_2_O (1:2), rt, 20–24 h.

All the final CQTrICh analogs (**7a-s)** and **9** were purified by column chromatography using 5% methanol in dichloromethane (DCM) as an eluent and obtained an excellent yield followed by characterization and confirmation using spectroscopic techniques such as IR, ^1^H, ^13^C NMR, and mass spectrometry. The purity was calculated using UP-LC and obtained with >95% for most of the analogs whereas few analogs were found with >92% purity.

Among all CQTrICh-analogs (**7a-s)** and **9**, one of the representative compounds (*E*)-1-(2- chlorophenyl)-3-(1-((1-(7-chloroquinolin-4-yl)-1*H*-1,2,3-triazol-4-yl)- methyl)-1*H*-indol-3-yl) prop-2-en-1-one (**7h**) is explained here based on spectroscopic data; however, similar trend obtained for all other reported analogs of the series. IR spectra of **7h** show a characteristic peak of (C-H)_str_ of triazole ring at 3149 cm^-1^, whereas a peak at 1650 cm^-1^ confirms the (C=O)_str_ of the chalcone. ^1^H NMR spectra of **7h** shows a total of 15 peaks in δ scale (ppm). A doublet appeared at 9.12 indicating the presence of N=C-*H* of the quinoline ring whereas a singlet appeared at 8.91 showing the presence of N=C-*H* of the indole ring and other aromatic peaks (protons of benzene and quinoline ring) appear between 8.26-7.04 with their coupling trends. Singlet peak appeared at 8.23 shows the presence of C-*H* of triazole ring whereas doublet at 7.97 and 7.95 assign the =C-*H*_indole side_ with J = 9.03 Hz and =C-*H*_keto side_ with J = 7.98 Hz in *trans*-conformation of *H*- C=C-*H* group, respectively. The singlet at 5.70, assign the C*H*_2_ protons present between the indole and triazole ring. ^13^C NMR spectra (δ, ppm) of **7h** show total of 29 peaks. Some characteristic peaks appeared at 193.38 ppm is due to *C*=O, 152.81 (*C*=N_quinoline_), 149.83 (*C*- N_quart_ of quinoline ring), 143.75 (*trans*-*C_indole side_*=C), 141.32 C_quart_ of benzene, 127.81 (*trans*-C=*C*_ketone side_), 130.41 *C*_quart_ of triazole ring, 117.59 *C*-H of triazole ring, 112.39 *C*_quart_ of indole ring carbon, 41.51 *C*H_2_ carbon. These data are well supported by 2D-COSY, 2D-HMBC, and 2D-HMQC analysis (spectra given in supplementary file). An interesting trend of mass spectrometry also confirms the mass of **7h**. The mass of **7h** was 523.10, whereas in mass spectra we obtained a mass of 496.21 [M-N_2_]^+^. As we know, the triazole ring is not that stable on fragmentation patterns, thus, the elimination of N_2_ (MW = 28) from the triazole ring took place. Therefore, we obtained a specific mass patterns of 496.21 [M-N_2_]^+^, 498.22 [M-N_2_+2]^+^, and 499.24 [M-N_2_+2+1]^+^ (which is also explained in the fragmentation pattern obtained from mass spectrometry, included in the supplementary file as **Fig. S.95**). The purity of all the compounds was checked with the UPLC data. In the case of **7h,** the 96.20 % of purity was recorded by UP- LC.

## 3. Biological activity and physico-chemical testing

### 3.1 *In vitro* growth inhibition assay against 3D7 strain of *P. falciparum*

CQTrICh analogs (**7a-s)** and **9** were tested for their *in vitro* antimalarial activity against the CQ*^S^*-3D7 strain of *P. falciparum* at two different concentrations (5 and 10 *µ*M) and compared to CQ as a standard drug (**Table 1, Fig. S.96**). Initial results were screened with percent inhibition, however, two active analogs **7l** and **7r** screened against the CQ*^R^*-RKL9 strain of *P. falciparum* and also determined their IC_50_ value. At, 10 *µ*M concentration more than 50% percent inhibition was obtained for **7f**, **7h**, **7l**, **7n**, **7p**, and **7r**, whereas other analogs were obtained with >30% inhibition as compared to a standard CQ. However, at 5 *µ*M concentration unexpected trend was obtained and none of the analogs exhibited more than 47% inhibition, except, the **7l** and **7r** which exhibited 56.84% and 55.20% inhibition, respectively as compared to a standard CQ. The unexpected trend arises with the negative inhibitory concentration due to the formation of precipitation during screening. The active CQTrICh-analogs **7l** and **7r** exhibited significant percent growth inhibition (>50% inhibition) as compared to untreated control against the tested strains of *P. falciparum* among the series. For comparison, we also synthesized compound **9** (*without chalcone unit*) to see whether it is more or less active than the CQTrICh- analogs (**7a-s**). Thus, we observed that compound **9** exhibits poor inhibition among the series at the tested concentration against the CQ*^R^*3D7 strain of *P. falciparum* confirming the role of chalcone in antimalarial activity.

**Table 1:**
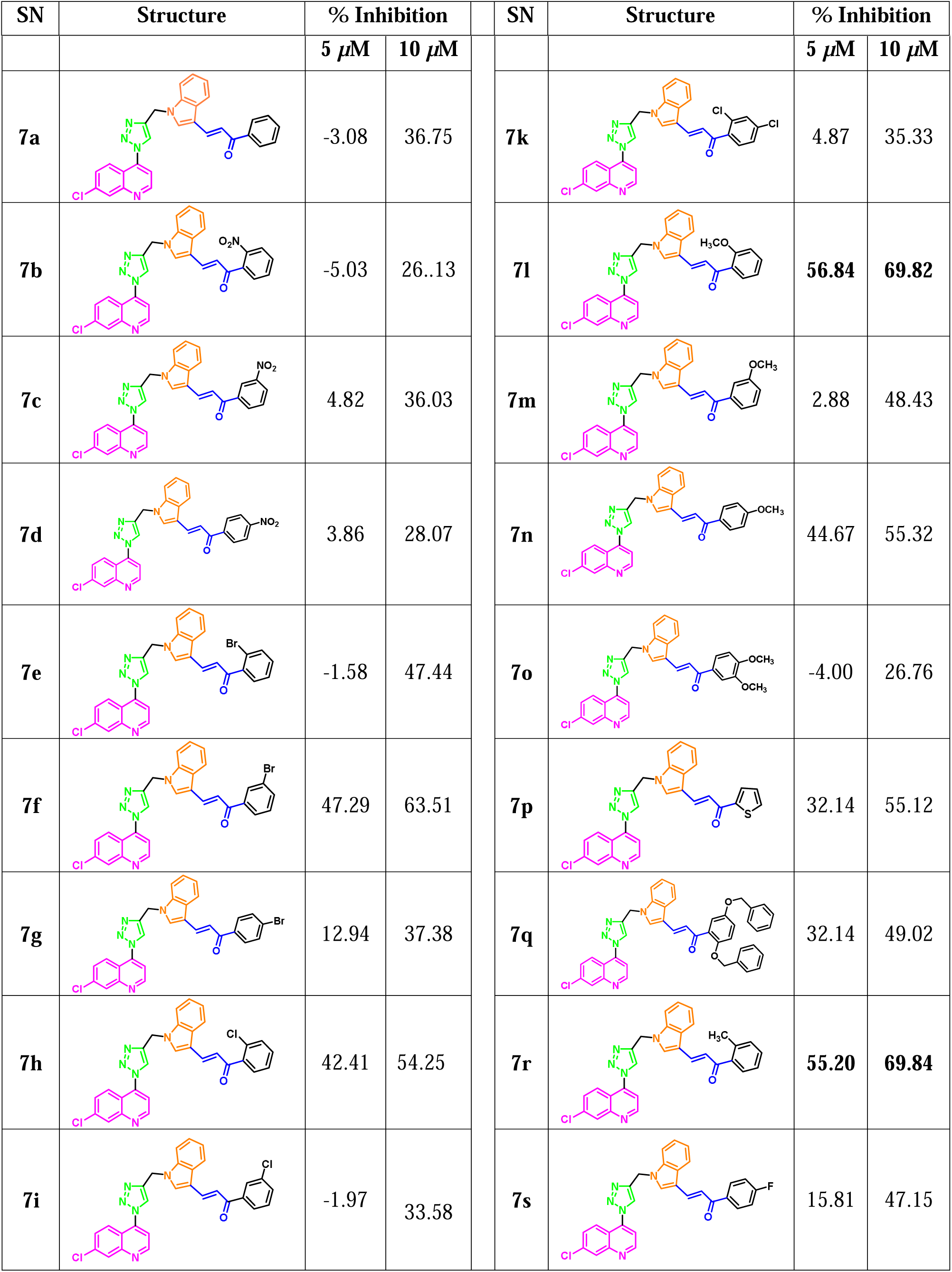
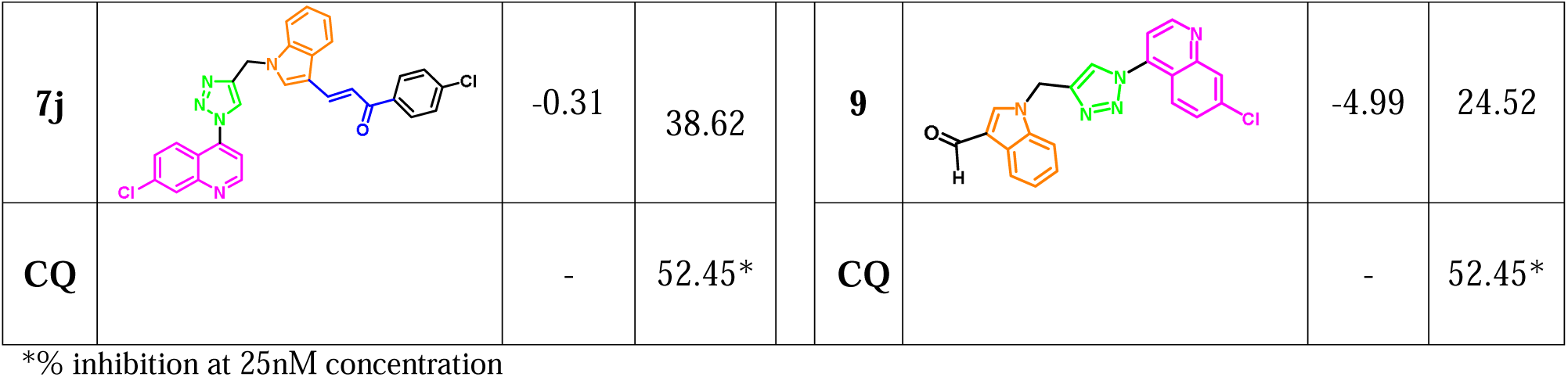
*In vitro* antimalarial inhibition activity of CQTrICh-analogs (**7a-s)** and **9** against the CQ*^S^*-3D7 strain of P. *falciparum*

| SN | Structure | % Inhibition | SN | Structure | % Inhibition |
| --- | --- | --- | --- | --- | --- |

| | | 5 μM | 10 μM | | | 5 μM | 10 μM |
| --- | --- | --- | --- | --- | --- | --- | --- |
| 7a |  | -3.08 | 36.75 | 7k |  | 4.87 | 35.33 |
| 7b |  | -5.03 | 26..13 | 7l |  | **56.84** | **69.82** |
| 7c |  | 4.82 | 36.03 | 7m |  | 2.88 | 48.43 |
| 7d |  | 3.86 | 28.07 | 7n |  | 44.67 | 55.32 |
| 7e |  | -1.58 | 47.44 | 7o |  | -4.00 | 26.76 |
| 7f |  | 47.29 | 63.51 | 7p |  | 32.14 | 55.12 |
| 7g |  | 12.94 | 37.38 | 7q |  | 32.14 | 49.02 |
| 7h |  | 42.41 | 54.25 | 7r |  | **55.20** | **69.84** |
| 7i |  | -1.97 | 33.58 | 7s |  | 15.81 | 47.15 |
| **7j** |  | -0.31 | 38.62 | **9** |  | -4.99 | 24.52 |
| **CQ** |  | - | 52.45* | **CQ** |  | - | 52.45* |

\*% inhibition at 25nM concentration

Based on percent inhibition data, both the active analogs (**7l** and **7r**) were further screened against the CQ*^S^*-3D7 and CQ*^R^*-RKL9 strain of *P. falciparum* to find IC_50_ value. Half maximal inhibition concentration (IC_50_) of the most effective compounds **7l** and **7r** was determined at various concentrations (0.62-20 *µ*M) against both the strains of *P. falciparum*, where one complete intra-erythrocytic cycle of *P. falciparum* was examined after the treatment of DNA-specific dye SYBR green I assay to observe the growth inhibition. Both **7l** and **7r** were found to be effective against the tested strains and showed inhibitory effects in a concentration-dependent manner as compared to untreated control. The IC_50_ of compounds **7l** and **7r** were found to be 2.4 *µ*M and 1.8 *µ*M, respectively against CQ*^S^*-3D7 and strain of *P. falciparum*. Furthermore, **7l** and **7r** were also tested against CQ*^R^*RKL9 of *P. falciparum*, and IC_50_ values were determined with 3.5 *µ*M and 2.7 *µ*M, respectively using Graph Pad Prism 8 software (**Fig. 2**). Thus, our results indicate that analogs **7l** and **7r** are significantly effective against the sensitive as well as the resistant strain of *Plasmodium*.

**Fig. 2:**
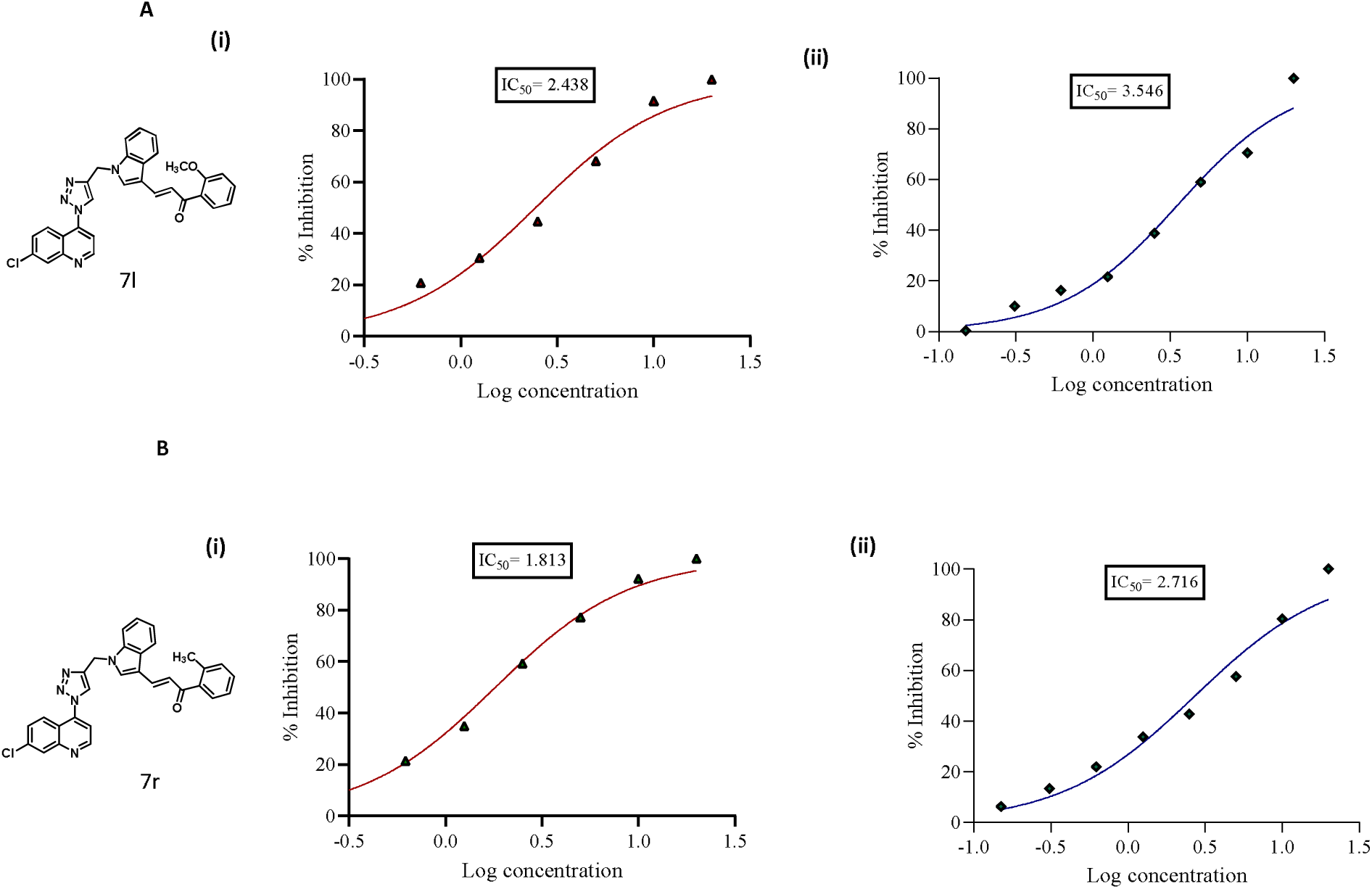
Growth inhibition assay: Evaluation of half-maximal inhibition concentration (IC_50_) of compounds to check their activity against the parasite. Synchronized ring-stage parasites were treated with compounds **7l (A)** and **7r (B)** ranging from 0.31 to 20 *µ*M concentrations. IC_50_ values of each compound in 3D7 **(i)** RKL9 **(ii)** were evaluated by plotting growth inhibition values against log concentration of these compounds. The experiment was done in triplicate and data was expressed as mean values ± SD.

### 3.2 Growth progression analysis

Growth progression evaluation of the potential antimalarial CQTrICh-analogs: **7r** and **7I** was carried out against the CQ*^S^*-3D7 strain of *P. falciparum*. The evaluation was performed for one complete growth cycle of the parasite. The parasite culture of ring-stage was treated with **7r** and **7I** at their IC_50_ concentration and parasites without the treatment of drug were taken as control. Thin Giemsa-stained blood smears were prepared at different time intervals of 20, 40, and 56- Hour Post-Treatment (HPT). Growth progression defect was analyzed by counting ∼2000 cells per Giemsa stained slide by light microscope. Morphological analysis was observed in th presence and absence of compounds. It was observed that after 40 HPT and 56 HPT, **7r** and **7** treated parasites were found to be stuck at the trophozoite stage with altered morphology. In untreated control, the growth of the cells progressed normally. As compared to untreated control, a gradual reduction in the percentage of parasitemia was observed (**Fig. 3**).

**Fig. 3:**
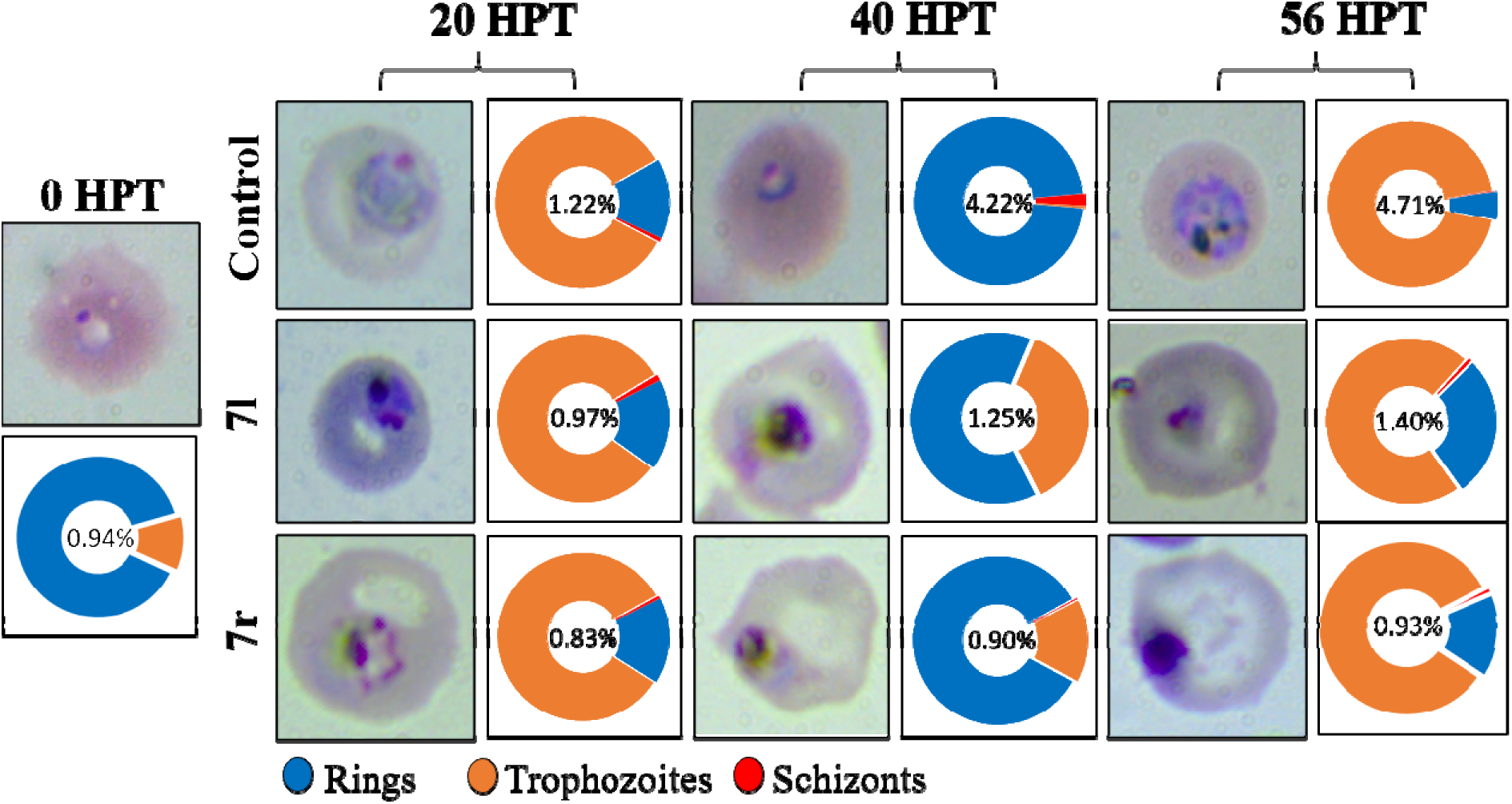
Light microscopic analysis of different blood stages of 3D7 *P. falciparum*, after treatment with the compounds: **7l** and **7r**. Growth progression analysis was evaluated for a complete intra- erythrocytic cycle from the ring-to-ring stage, at different time intervals of 20, 40, and 56 HPT. Synchronized ring-stage parasite culture of 1% parasitemia was treated with compounds **7l** and **7r** at their IC_50_ concentrations. Morphological changes were analyzed by Giemsa staining at intervals of 20, 40, and 56 HPT. Graphs and figures show the growth defect during parasite egress or invasion.

### 3.3 Hemozoin inhibition study

During Haemoglobin (Hb) degradation pathway, ferric heme a by-product of Hb degradation, is toxic to both the malaria parasites and the host cells [30–33]. Consequently, malaria parasite for their protection undergoes a pathway, which converts toxic heme into non-toxic metabolites such as crystallization into hemozoin. Hemozoin is a malaria pigment and insoluble in water [32–33]. Spectrophotometric analysis was performed to examine free monomeric heme obtained from hemozoin. For the quantification of monomeric heme, synchronized ring-stage parasites were treated with **7l** and **7r** and observed up to the schizont stage. The result shows that when parasites treated with different concentrations of **7l** and **7r** exhibit decreased percentage of free monomeric heme in a dose-dependent manner as compared to untreated control. At, 10 *µ*M concentration **7l** showed 44% whereas **7r** showed 48.8% control-free heme (**Fig. 4A**). Absorbance was taken at 405/750 nm corresponds to the free monomeric heme obtained from hemozoin of parasites later evidenced by the reduction in Hb hydrolysis. Crystallization of hemozoin is vital for the persistence of the malaria parasites that creates interest in the development of novel antimalarial drugs [31–34].

**Fig. 4:**
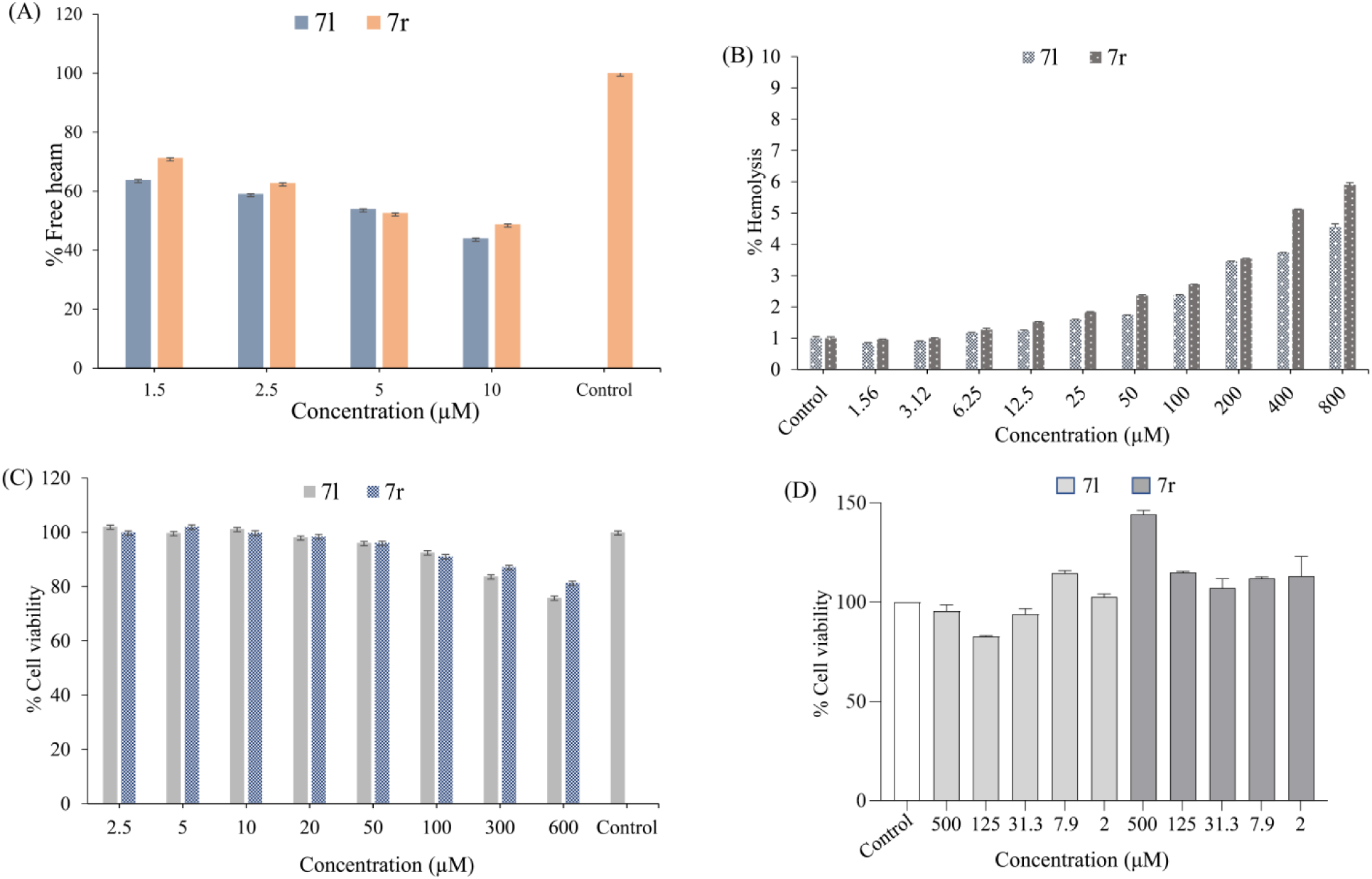
(A) The graph showing the hemozoin inhibition in parasites exposed to compounds (**7l**) and (**7r**). Synchronized ring-stage parasites were treated with compounds (**7l**) and (**7r**) at different concentrations for 30h; control-free heme was calculated by spectrophotometry. **(B)** Effect of triazole hybrids **7l** and **7r** on human RBCs. Hemolytic effect was checked at different concentration (1.5-800 *µ*M) with 10% (v/v) RBCs suspension for 1h. Absorbance was taken at 415 nm indicating no significant lysis occurred even in a higher concentrations of compounds (**7l**) and (**7r**) up to 800 *µ*M. MTT assay was done to check the HEK293 **(C)** and HEPG2 **(D)** cells’ viability after treatment with compounds (**7l**) and (**7r**) at various concentrations.

### 3.4 Hemolysis and cytotoxicity study

The toxicity of compounds: **7l** and **7r** on human red blood cells (hRBCs) was observed to evaluate the hemolytic activity. The hemolytic study was observed at different concentrations ranging from 1.5-800 μM. At a maximum concentration of 800 μM, **7l** and **7r** showed toxicity triggering only 4.5 and 5.9% hemolysis, respectively. However, at 50 μM concentration, the analogue **7l** and **7r** show nontoxic nature, causing only 1.7 and 2.3% cell lysis, respectively. Thus, these analogs have safely exhibited very less toxicity (**Fig. 4B**).

The selected compounds were further evaluated for cytotoxicity by the (3-(4,5-dimethyl- 2-yl)-2,5-diphenyltetrazolium bromide) (MTT) assay on the noncancerous human cell lines: HEK293, which is a specific cell line initially resulting from the cells of the human embryonic kidney grown by a tissue culture; and HEPG2, which is a cell line exhibiting epithelial-like morphology. HEK293 and HEPG2 cells are widely used in cell biological research for decades, because of their consistent growth and inclination for transfection. The selected compounds **7l** and **7r** were screened in the concentration range of 2.5−600 μM (HEK293) and 2−500 μM (HEPG2), and it was found that **7l** and **7r** are non-cytotoxic to the host cells. Further, based on these results, we can speculate that these compounds might be a good anti-malarial entity (**Fig. 4C and 4D**).

### 3.5 Molecular docking study

In the malaria parasite *P. falciparum*, a targeted gene-disruption approach adopted by Kato et al. demonstrated that *Pf*CDPK1 is essential for parasite viability by regulating parasite motility during egress and invasion. He postulated a series of structurally related 2,6,9-trisubstituted purines with an *in vitro* biochemical screening using recombinantly purified *Pf*CDPK1 protein against a library of 20,000 compounds. SAR studies depicted that quinoline core containing 2,6,9-trisubstituted purines derivative exhibited potent cellular (EC_50:_ 0.1 μM) and *Pf*CDPK1- inhibitory (IC_50_: 3.846 μM) activities [16]. In the subsequent year, Lemercier *et al.* studied the structural requirements for the inhibition of *Pf*CDPK1 by developing a primary screening assay, in which a total of 54,000 compounds were tested, yielding two distinct chemical series of the small molecules which inhibited in the nano-molar range, which was further characterized through enzymatic and biophysical analysis. One of the active pharmacophores containing indolizine core (**A**) was shown to inhibit *Pf*CDPK1 with a sub-micromolar potency [17]. Therefore, with this idea, we focused on indole and quinoline core to produce CQTrICh-analogs (**7a-s)** and **9**, whereas quinoline core and chalcones are already known for their potent antimalarial activity [7,8,21,22]. Thus, we have gone through molecular docking studies of **7l** and **7r** to find the possible binding mode with the *Pf*CDPK1 enzyme of *P. falciparum*, indeed it was confirmed by the speculated binding possibilities. Here, we found that the compounds **7l** and **7r** showed good binding affinity against *Pf*CDPK1, and preferentially occupied the active site cavity (**Table 2 upper panel**). Compounds **7l** and **7r** showed -10.7 kcal/mol and -10.2 kcal/mol binding free energy with *Pf*CDPK1, respectively. Both the compounds formed several close interactions such as H-bonds to active site residues and other interactions offered by the protein *Pf*CDPK1 (**Table 2 lower panel**). The molecular docking study provides an accurate and preferred orientation of a compound at the binding site of a receptor. Interaction analysis of all possible docked conformers of compounds was carried out to investigate their binding pattern and possible interactions with the ATP binding pocket of *Pf*CDPK1 (**Fig. 5**). Thus, the analysis of docked conformations indicates that both compounds bind deeper into the binding pocket of *Pf*CDPK1 and perhaps block the accessibility of substrate which may be presumably responsible for growth inhibition by decreasing the accessibility of substrate or binding partner.

**Fig. 5:**
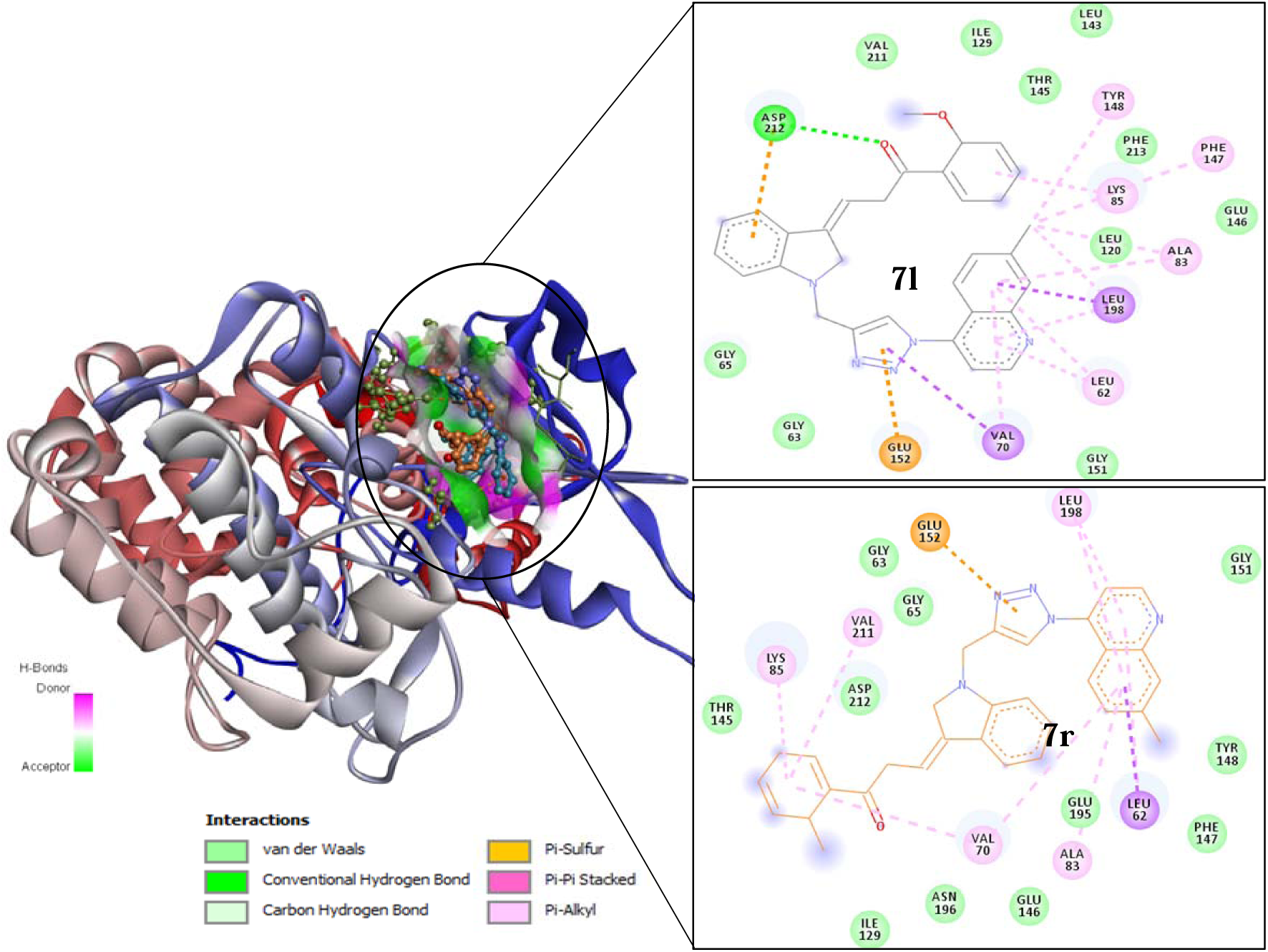
*In silico* studies of lead compounds (**7l**) and (**7r**) with *Pf*CDPK1. Cartoon representation showed the compound (**7l**) and (**7r**) interact within the binding pocket (left). 2D image of compound (**7l**) showing hydrogen bond and hydrophobic interactions (upper right) and **7r** (lower right).

**Table 2:** Molecular interaction of (**7l**) and (**7r**) with *Pf*CDPK1 binding site (upper panel). The binding affinity of compounds with *Pf*CDPK1 generated from molecular docking along with the key interacting residues (lower panel).

| Target protein | SN (Ligand name) | Binding Free Energy (kcal/mol) | pKi | Ligand Efficiency (kcal/mol/non-H atom) | Torsional Energy |
| --- | --- | --- | --- | --- | --- |
| *Pf*CDPK1 | **7l** | -10.7 | 7.85 | 0.2098 | 2.1791 |
|  | **7r** | -10.2 | 7.48 | 0.2082 | 1.8678 |
| Protein-ligand Interactions |  |  |  |  |  |
| SN | Affinity<br>(kcal/mol) | Hydrogen Bonds |  | Other Interacting Residues |  |
|  |  | Amino Acid<br>Residues | Distance (Å) |  |  |
| **7l** | -10.7 | Asp 212 | 2.67 | Val-211, Ile-129, Thr-145, Leu-143, Tyr-148, Phe-213, Phe-147, Lys-85, Leu-120, Phe-147, Glu-146, Ala-83, Leu-198, Leu-62, Gly-151, Val-70, Glu-125, Gly-62, Gly-65, Ala-66, |  |
| **7r** | -10.2 | - | - | Thr-125, Lys-85, Asp-212, Val-211, Gly-85, Gly-63, Glu-152, Leu-198, Gly-151, Tyr-148, Phe-147, Leu-62, Glu-195, Ala-83, Glu-146, Val-17, Asn-196, Ile-129 |  |

### 3.6 Drug-likeliness assessment

Physico-chemical profile (ADME properties) of the CQTrICh-analogs and compound **9** were also determined. We evaluated the druggability of triazoles by calculating its Lipinski’s Ro’5 (Rule of five) descriptors. Out of **20** compounds, only **7a, 7p** and **9** fulfilled all the Ro’5 norms. The physicochemical properties of all hybrids **7a-s** and **9** are given in (**Table S1** in **supplementary data**). All the triazoles under study were found to contain less than 15 rotatable bands. Carcinogenicity of compounds was checked using Carcinopred-EL, we found that all hybrids are non-carcinogenic in nature and not even a single compound passed the PAIN pattern.

### 3.7 Compounds interact with and inhibit CDPK1-activity *in vitro*

MST, a novel method for immobilization-free interaction analysis was used to assess the r*Pf*CDPK1 (**7l** and **7r**) interaction *in vitro*. When fluorescently labeled r*Pf*CDPK1 was titrated against increasing concentrations of **7l** and **7r**, its thermophoretic mobility markedly changed indicating effective interaction with the compounds. Calculation of equilibrium dissociation constants of **7l** and **7r** revealed *K_d_ values* of 17.3 μM and 7.2 μM, respectively (**Fig. 6A(i) and 6A(ii)**). According to Ozbabacan *et al.*, *K_d_* values in this micro-molar range are commonly found for transient interactions [36]. Since our *in silico* interaction analysis depicted the potentiality of the compounds to interact with the ATP binding pocket of CDPK1, we utilized *in vitro* kinase assay to demonstrate the enzymatic activity of the protein in the presence of the compounds: **7l** and **7r**, by measuring the amount of ADP produced in the kinase reactions. Both the analogs exhibited differential inhibition of CDPK1 activity which supports well with our *in silico* docking approach and MST-based interaction analysis. Notably, **7l** and **7r** were significantly inhibited phosphorylation activity of CDPK1, accounting for 28.1% and 54.5% (at 1 μM) and 11.6% and 56.2 % inhibition (at 5 μM), respectively (**Fig. 6B(i)**). We further analyzed the effect of these compounds on mammalian kinome by taking HepG2 cytosolic fraction as kinome source, interestingly both compounds **7l** and **7r** showed no inhibition of the host kinase activity (**Fig. 6B(ii)**). Overall, these results suggest that analogs **7l** and **7r** might be an eye-opener and active candidate to inhibit the CDPK1 enzymatic activity, without affecting the host kinome.

**Fig. 6:**
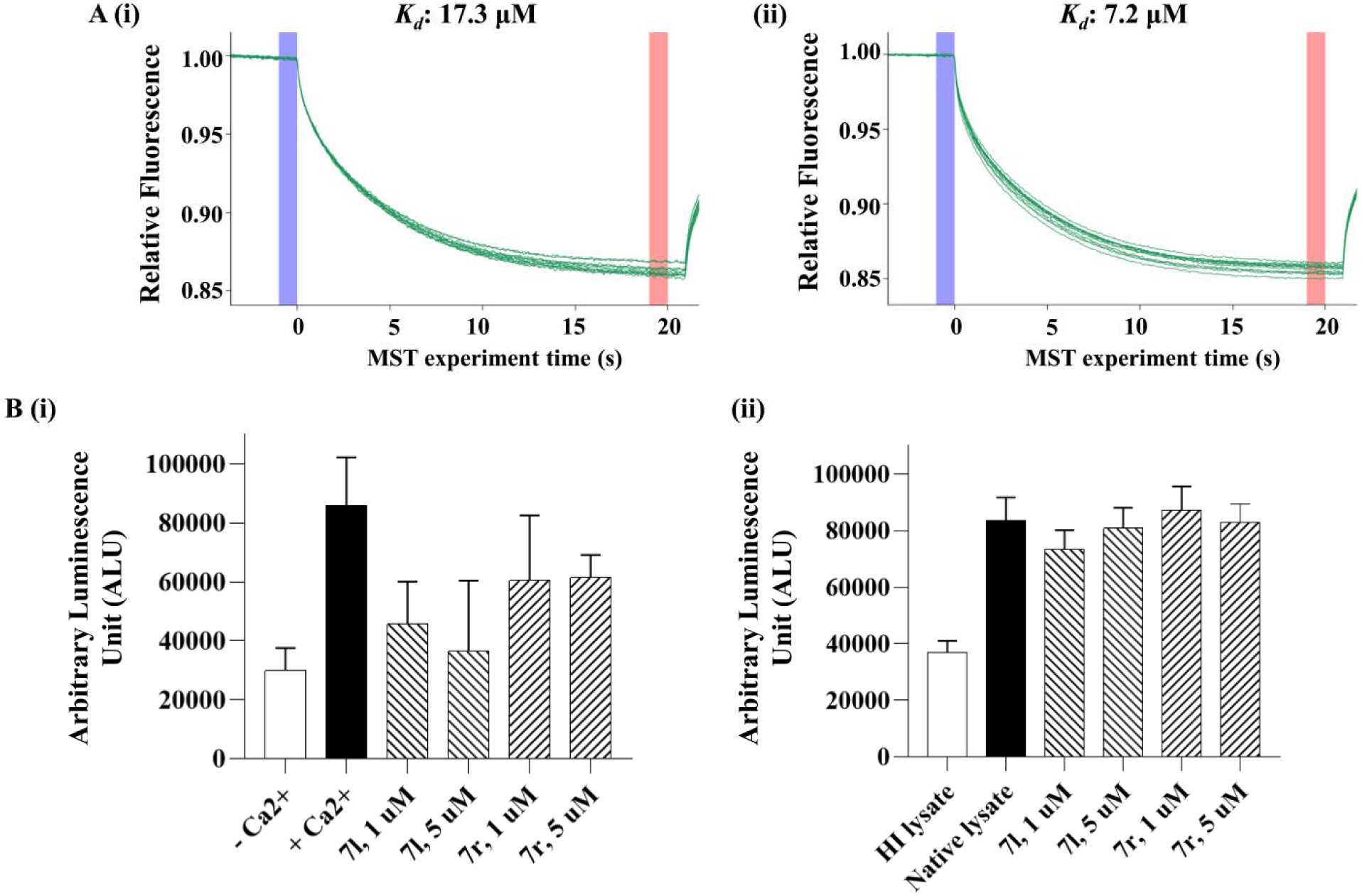
7l and 7r interact with and inhibit CDPK1-activity *in vitro*. MST was used to assess the r*Pf*CDPK/(**7l** or **7r**) interaction *in vitro* **(A)**. Thermophoretic mobility of fluorescently labeled r*Pf*CDPK1 altered dramatically, showing that **7l** and **7r** had an effective interaction, with *K_d_* values of 17.3 μM and 7.2 μM, respectively. Further, *in vitro* kinase assay in the presence of **7l** and **7r** depicted that both compounds showed differential inhibition of CDPK1 activity which supports well with our *in silico* docking approach and MST-based interaction analysis **(B(i))**. **7l** and **7r** inhibited phosphorylation activity of CDPK1, accounting for 28.1% and 54.5% (at 1 μM) and 11.6% and 56.2 % inhibition (at 5 μM), respectively. **7l** and **7r** were assessed for their effect on host kinase activity inhibition and showed no inhibitory effect on the human kinome **(B(ii))**.

## 4. Discussion

4-Aminoquinoline antimalarials contain side chains with a basic amino group thought to play a key role in pH trapping where the basic amino group in the side chain is one of the major factors to enhance its antimalarial activity. [26,37]. Indole rings (which also contains basic amino group) containing indolizines are well explored to inhibit *Pf*CDPK1 with a sub-micromolar potency [7]. Other hand chalcone-containing side chains in 4-Aminoquinoline are also known for their antimalarial activity. [7,8] Thus, keeping these ideas of antimalarial activity and indole ring involvement to inhibit the *Pf*CDPK1 enzyme (which is essential for the parasite viability by regulating the parasite motility during egress and invasion), we have designed and developed CQTrICh-analogs (**7a-s** and **9**). All the CQTrICh-analogs were determined for their Log P-value and ranging between 3.52 and 4.50, whereas compounds **7q** and **9** were reported with the value of 5.80 and 3.01, respectively.

The most active CQTrICh-analogs (**7l** and **7r**) against *in vitro* CQ*^S^*-3D7 strain, were selected and further screened for their *in vitro* antimalarial IC_50_ value against CQ*^S^*-3D7 and CQ*^R^*- RKL-9 strains of *P. falciparum*. Selected CQTrICh-analogs (**7l** and **7r**) exhibited an IC_50_ value of 2.4, 3.5 *µ*M, and 1.8, 2.7 *µ*M, respectively, against CQ*^S^*-3D7 and CQ*^R^*-RKL-9 strain of *P. falciparum*, whereas Resistance Index (RI) value was obtained 1.45 (**7l**) and 1.49 (**7r**) (**Table 3**). The RI value was found to increase with decreasing log P in both the tested compounds. Therefore, almost a linear dependence of RI on log P suggests that the lack of possible cross-resistance in the tested analogs may be related to its physiochemical properties rather than CQ chemosensitizing itself.

**Table 3:** *In vitro* IC_50_ (*µ*M) of selected potential CQTrICh-analogs (**7l** and **7r**) with their RI and Log P-value and *Pf*CDPK1 inhibition activity

| SN | Structure | IC<sub>50</sub> (μM) | | RI<sup>a</sup> | Log P | Cytotoxicity (HEK293)<sup>b</sup> | PI of CDPK1<sup>c</sup> | |
| --- | --- | --- | --- | --- | --- | --- | --- | --- |
| | | CQ<sup>S</sup> 3D7 | CQ<sup>R</sup> RKL9 | | | | 1 μM (con.) | 5 μM (con.) |
| **7l** |  | 2.438 | 3.546 | 1.45 | 4.29 | >90% | 28.1% | 11.6% |
| **7r** |  | 1.813 | 2.716 | 1.49 | 4.15 | >90% | 54.5% | 56.2% |

<sup>a</sup>Resistance Index (RI) = IC<sub>50</sub> ( μM) CQ<sup>R</sup>-RKL-9 / CQ<sup>S</sup>-3D7
<sup>b</sup>At 100 μM concentration more than 90% cell are viable, so can be speculated they are non-toxic to tested non-cancerous human cell lines.
<sup>c</sup>**7l** and **7r** were significantly inhibited phosphorylation (PI) activity of CDPK1, accounting for 28.1% and 54.5% (at 1 μM) and 11.6% and 56.2 % inhibition (at 5 μM), respectively.

Growth progression analysis was also performed for the active CQTrICh-analogs (**7l** and **7r**) against the CQ*^S^*-3D7 strain of *P. falciparum*, where parasite culture of the ring stage was treated with both CQTrICh-analogs **7l** and **7r** at their respective IC_50_ concentration, while parasites without the treatment of drug were taken as control by thin Giemsa-stained blood smears with 20, 40 and 56 hrs post-treatment (HPT) intervals. Morphological analysis was observed in the presence and absence of the analogs tested and it was observed that after 40 HPT and 56 HPT, **7l** and **7r** treated parasites were stuck at a trophozoite stage with altered morphology whereas in untreated control the growth of the cell progressed normally. Overall, in treated cells, a gradual reduction in parasitemia percentage was observed as compared to untreated control (**Fig. 4**). Spectrophotometric analysis of CQTrICh-analogs (**7l** and **7r**) was also performed to examine free monomeric heme obtained from hemozoin. Thus, **7l** and **7r** were treated with ring-stage parasites and observed up to the schizont stage at various concentrations and compared with the untreated control in a dose-dependent manner. At 10 μM concentration **7l** and **7r** exhibited 44% and 48.8% control-free heme, respectively. These CQTrICh-analogs (**7l** and **7r**) were also studied for their toxicity behavior on human Red Blood Cells (hRBCs) at various concentrations between 1.56 to 800 μM. The toxicity was triggering only 4.5% and 5.9% hemolysis at a maximum concentration (800 μM), however; non-toxic nature was observed with 1.7% and 2.3% cell lysis, respectively for **7l** and **7r**. This indicate that the two CQTrICh-analogs are safer to be carried out further for pharmacological investigation. This study was also supported by hemozoin inhibition. In a further study CQTrICh-analogs (**7l** and **7r**) were also evaluated for cytotoxicity at various concentration ranges on noncancerous human cell lines: HEK293 and HEPG2. Both the tested compounds significantly exhibited with >90% cells are viable at even 100 μM concentration. Thus, we can speculate that these tested analogs are nontoxic and may be potential antimalarial scaffolds.

Overall, if we look at the analogs **7h**, **7l,** and **7r**, interestingly an increased inhibition activity (concentration based) was observed when, -Cl (**7h**), -OMe (**7l**), and -Me (**7r**) were located at *ortho*-position in benzene attached to the chalcone unit which might create special geometry and helping in the development of interaction with *Pf*CDPK1 kinase enzyme. These groups might be playing a major role in increasing the antimalarial inhibition activity and later these were proven by hemozoin inhibition, hemolysis, and docking study followed by interaction with *Pf*CDPK1. These speculations were also well supported by the *in vitro Pf*CDPK1 inhibition activity, where the equilibrium dissociation constant revealed *K_d_*values of 17.3 *µ*M and 7.2 *µ*M for **7l** and **7r**, respectively. According to the *in silico* interaction study, we found that these CQTrICh-analogs (**7l** and **7r**) possibly interact with the ATP binding pocket of CDPK1. Molecular interaction of **7l** and **7r** exhibit -10.7 and -10.2 kcal/mol of binding free energy with p*K_i_* values of 7.85 and 7.48. Simultaneously, 0.2098 and 0.2082 kcal/mol/non-h atom ligand efficiency was observed along with torsional energy of 2.1791 and 1.8678, respectively. Analogue **7l** also exhibited possible protein-ligand interaction with an affinity of -10.7 kcal/mol and formed H-bond with Asp 212 residue of the protein testes, whereas **7r** exhibited possible protein-ligand interaction with an affinity of -10.2 kcal/mol, along with other interacting residues (**Table 2**).

*In silico* and *in vitro* binding experiments with the recombinantly purified *Pf*CDPK1 depicted that the lead compounds: **7l** and **7r** interact with the protein, with K_d_ values of 17.3 and 7.2 μM, respectively, and these values are comparable to those of ATP which usually reaches high concentrations within cells, in the range of 1 to 10 mmol/L, indicating that the compounds are unlikely to have a significant effect. However, earlier attempts have been made to utilize the scaffolds: indole, triazole, and quinoline, to synthesize novel *specific* **inhibitors** with their inhibitory activities at the μM range, against numerous kinases. In this direction, Xu *et al.* synthesized a novel [1,2,4]**triazol**o[4,3*-b*][1,2,4,5]tetrazine derivative which was found to be a potent anti-proliferative agent and inhibited c-Met kinase activity (IC_50_: **11.77** μ**M**) [38]. New 1,2,4-**triazole** scaffolds were synthesized by El-Sherief *et al.* and most of the tested compounds exhibited noteworthy anti-proliferative effects against a panel of cancer cell lines [39]. The mechanistic study showed that the lead compounds strongly inhibited EGFR and BRAF^V600E^ kinases in the **low** μ**M** range. In the subsequent year, Goma’a *et al.* designed and synthesized several 1,2,4-**triazole** derivatives with ethyl 2-((5-amino-1H-1,2,4-**triazol**-3-yl)thio)acetate as the starting material [40]. The lead compound could reduce the viral plaques by 50% at a dose of **80** μ**M** against Herpes Simplex Virus-1 (HSV-1), grown on Vero African green monkey kidney cells. Docking studies revealed that the lead compound interacted with the active site of HSV-1 thymidine kinase. In the malaria parasite *P. falciparum*, a targeted gene-disruption approach adopted by Nobutaka Kato et al. (2008) demonstrated that *pfcdpk1* is essential for the parasite viability by regulating parasite motility during egress and invasion, and identified a series of structurally related 2,6,9-trisubstituted purines by an *in vitro* biochemical screening using recombinantly purified *Pf*CDPK1 protein against a library of 20,000 compounds [16]. SAR studies of the 2,6,9-trisubstituted purines yielded compound **14** harboring **quinoline** moiety, which showed potent *Pf*CDPK1-inhibitory (IC_50_: **3.8** μ**M**) activities. More recently, our research group adopted a structure-based virtual chemical library screening approach in combination with extensive biochemical and biophysical characterization-based tools to identify novel lead candidates: Compound 1_(ST092793)_ and Compound 2_(S344699)_, capable of inhibiting CDPK1 activity *in vitro*. Dose-response curves for CDPK1-inhibitory activity of the compounds depicted their IC_50_ values to be **33.8** and **42.6** μ**M**, respectively; and, the compounds showed intra-erythrocytic growth inhibition of the parasite, depicting their IC_50_ value of **9.5** and **17.8** μ**M**, respectively [41].

Based on *in silico* output, we utilized *in vitro* kinase assay to demonstrate the enzymatic activity of the protein in presence of tested CQTrICh-analogs (**7l** and **7r**) by measuring the ADP produced during kinase reaction. They both (**7l** and **7r**) significantly inhibited phosphorylation activity of CDPK1, accounting for 28.2% and 54.5% (at 1 *µ*M), and, 11.6% and 56.2% (at 5 *µ*M) inhibition, respectively (**Table 3**). Analogue **9** (without chalcone) was not a good inhibitor at concentration-based analysis. Thus, the overall findings and observations synchronize well with each other.

## 5. Conclusion

A successful synthesis of a series of CQTrICh-analogs (**7a-s)** and **9** using click chemistry has produced **7l** and **7r** likely to be potential antimalarial analogs. Infect, these analogs have proved to be a potential and valuable probes of antimalarial activity, resistance, and nontoxic in nature. The 4-aminoquinoline bearing with triazole, *indole,* and chalcone unit was also found to be reasonably eye-opener probes due to its significant contribution in inhibition of phosphorylation activity of CDPK1 protein. These probes or analogs might open a new way of thinking in the history of antimalarials. All the synthesized analogs were screened for their *in vitro* percent inhibition antimalarial activity. However, the most active analogs among them were evaluated further for their *in vitro* IC_50_ value against CQ*^S^*-3D7 and CQ*^R^*-RKL-9 strains of *P. falciparum* where **7l** (*o*-methoxy) and **7r** (*o*-methyl) were found to be effective against the tested strains with an IC_50_ value against CQ*^S^*-3D7 and CQ*^R^*-RKL-9 strain of *P. falciparum* with 2.4, 3.5 *µ*M and 1.8, 2.7 *µ*M, respectively. The RI value was obtained at 1.45 (**7l**) and 1.49 (**7r**). Hemozoin inhibition at 10 *µ*M concentration exhibited 44% and 48% control free heme for **7l** and **7r**, respectively whereas at 50 *µ*M concentration both compounds were found to be non-toxic in nature. Both the selected analogs were also found non-cytotoxic to HEK293 and HEPG2 cells. Docking study also revealed that **7r** and **7l** exhibit good binding affinity against *Pf*CDPK1 protein and occupied the active site cavity where the ATP were bonded. *In vitro* activity also supports *in silico* docking approach to identify the lead probe. Analogs, **7l** (28% at 1 *µ*M, 11.6% at 5 *µ*M) exhibit better inhibition as compared to **7r** (54.5% at 1 µM and 56.2 % at 5 µM) against *Pf*CDPK1 protein. Overall, results suggested that these selected CQTrICh-analogs (**7r** and **7l**) shown to be a promising probe with antimalarial activity and futuristic landmark as the CDPK1 inhibitors.

This is the first attempt to design CDPK inhibitors using the hybridization approach, but it still needs further improvements. The antimalarial properties of indole-based chalcones (side chain part of the hybrid) and several chalcones based on different aryl/heteroaryl moieties are very well documented in the literature. The idea to link these chalcones with CQ via a triazole linker was to improve their efficacy and selectivity against the malaria parasite. Overall, the strategy resulted in the formation of new scaffolds, but none of them demonstrated activity in the nanomolar range, which was our aim in designing such molecules. Fortunately, we found two analogs that were equipotent against the CQ^S^ and CQ^R^ isolates, which is a reasonable start for a new series. We are now excited to explore these two analogs using computational as well as SAR approaches to improve their efficacy by optimizing the CQ and side-chain parts. This would necessitate additional planning to design, synthesize, characterize, and conduct their biological studies, including CDPK inhibition studies.

## 6. Material and Methods

### 6.1. Chemistry

#### 6.1.1. Experimental Protocols

All the chemicals and solvents (analytical grade) were purchased from Sigma-Aldrich, USA. Thin-layer chromatographic analysis was carried out on precoated Merck silica gel 60 F_254_ TLC aluminium sheets and spots were visualized under UV light at 254 nm. IR spectra were recorded on Agilent Cary 630 FT-IR spectrometer and only major peaks are reported in cm**^-1^**. **^1^**H and ^13^C NMR spectra were obtained in DMSO-*d_6_* as a solvent with tetramethylsilane (TMS) as an internal standard on Bruker Spectrospin DPX-700 spectrometer at 700 MHz and 75 MHz, respectively. Splitting patterns are designated as follows: s (singlet), d (doublet), t (triplet), m (multiplet) or brs (broad). **^1^**H NMR chemical shifts (δ) values are reported in parts per million (ppm) relative to residual solvent (DMSO-*d_6_*, δ 2.54) and ^13^C NMR chemical shifts (δ) are reported in ppm relative to (DMSO-*d_6_*, δ 39.5) and coupling constants (*J*) are expressed in Hertz (Hz). Mass spectra were recorded on an Agilent Quadrupole-6150 LC/MS spectrometer. Melting points were measured on a digital Buchi melting point apparatus (M-560) and are uncorrected. Purities were determined by Agilent RRLC MS 6320 ion trap spectrometer using an X-bridge C18 1.7 µm column (50 mm × 2.1 mm). Mobile phase channel A consisted of 5 mM ammonium acetate in water. Mobile phase B consisted of acetonitrile with a flow rate = 0.8 mL/min; detection was done by UV@214, and all final compounds were confirmed to have ≥95% purity. Purification of the compounds was carried out by silica gel column chromatography (230-400 mesh size) with the indicated eluent.

#### 6.1.2 Synthesis of chalcone (3a-s)

In a dried two-neck round bottom flask purged with argon, Indole 3-aldehyde **1** (1.0 mmol) was dissolved in anhydrous ethanol (10 ml). To this solution, substituted ketone **2a-s** (1.0 mmol) and piperidine (3.0 mmol) were added with constant stirring. The reaction mixture was then allowed to reflux for 24 hr. A yellow precipitate formed during the course of the reaction was filtered and washed with chilled ethanol, dried, and recrystallized from ethanol to afford pure product [42]. All the chalcones were reported earlier [43–45].

#### 6.1.3 Synthesis of alkyne (4a-s), (8)

To a solution of indole-chalcone **3a-s** (1.0 g, 3.42 mmol) in anhydrous DMF (10 ml), potassium carbonate (1.2 g, 8.55 mmol) was added. The reaction mixture was allowed to stir for 15 min at room temperature. To this solution, propargyl bromide (0.76 mL, 6.84 mmol) was added dropwise at room temperature. The reaction mixture was stirred overnight under argon and then quenched by the addition of water. The aqueous phase was extracted with ethyl acetate (2×30 mL). The combined organic layers were washed with brine, dried over anhydrous sodium sulfate, and evaporated under vacuum. The crude was purified by column chromatography using 30% ethyl acetate in hexane to give the indole-chalcone alkyne good to excellent yields [46].

##### 6.1.3.1 (E)-1-phenyl-3-(1-(prop-2-yn-1-yl)-1H-indol-3-yl)prop-2-en-1-one (4a)

Yellow solid, yield: 40.4 %, R_f_ = 0.75 (Ethylacetate : Hexane = 50 : 50), mp: 226.5 °C, IR (neat): ν (cm^-1^) 3222 (≡C-H)_str_, 1641 (C=C)_str._ chalcone, ^1^H NMR (700 MHz, DMSO-*d_6_*) (δ-ppm): 8.20 (s, 1H, Ar-H), 8.14 (d, *J* = 7.2 Hz, 3H, CH, Ar-*H*), 8.03 (d, *J* = 15.5 Hz, 1H, Ar-*H*), 7.70 (d, *J* = 15.5 Hz, 1H, C*H*), 7.67 – 7.65 (m, 2H, Ar-*H*), 7.58 (t, *J* = 7.6 Hz, 2H, Ar-H), 7.36 (t, *J* = 7.5 Hz, 1H, Ar-*H*), 7.32 (d, *J* = 7.8 Hz, 1H, Ar-*H*), 5.20 (d, *J* = 2.5 Hz, 2H, C*H_2_*), 3.53 (t, *J* = 2.5 Hz, ≡C-*H*). ^13^C NMR (176 MHz, DMSO-*d_6_*) (δ-ppm): 188.81, 138.32, 138.02, 136.91, 134.92, 132.52, 128.75, 128.71, 128.18, 125.81, 123.04, 121.76, 120.70, 116.27, 112.58, 111.11, 78.39, 76.40, 35.67. ESI-MS (*m/z*) calcd. for C_20_H_15_N0: 285.34; Found: 286.16 [M+H]^+^.

##### 6.1.3.2 (E)-1-(2-nitrophenyl)-3-(1-(prop-2-yn-1-yl)-1H-indol-3-yl)prop-2-en-1-one (4b)

Yellow solid, yield: 42.1 %, R_f_ = 0.73 (Ethylacetate : Hexane = 50 : 50) mp: 150.1 °C, IR (neat): ν (cm^-1^) 3252 (≡C-H)_str_, 1620 (C=C)_str._ chalcone, ^1^H NMR (700 MHz, DMSO-*d_6_*) (δ, ppm): 8.18 (dd, *J* = 8.2, 0.9 Hz, 1H, C*H*), 8.11 (s, 1H, Ar-*H)*, 7.99 (d, *J* = 8.0 Hz, 1H, C-*H*), 7.90 (td, *J* = 7.5, 1.1 Hz, 1H, Ar-*H*), 7.80 (td, *J* = 8.2, 1.4 Hz, 1H, Ar-*H*), 7.75 (dd, *J* = 7.5, 1.3 Hz, 1H, Ar-*H*), 7.66 (t, *J* = 11.8 Hz, 2H, Ar-*H*), 7.36 – 7.33 (m, 1H, Ar-*H)*, 7.29 – 7.26 (m, 1H, Ar-*H*), 7.09 (d, *J* = 16.1 Hz, 1H, Ar-*H*), 5.17 (d, *J* = 2.5 Hz, 2H, C*H_2_*), 3.52 (t, *J* = 2.5 Hz, 1H, ≡C*H*). ^13^C NMR (176 MHz, DMSO-*d_6_*) (δ-ppm): 191.51, 147.06, 140.50, 136.97, 135.91, 135.66, 134.07, 131.00, 129.10, 125.45, 124.44, 123.18, 121.95, 120.57, 119.98, 111.90, 111.21, 78.19, 76.50, 35.70. ESI-MS (*m/z*) calcd. for C_20_H_14_N_2_0_3_: 330.34; Found: 331.04 [M+H]^+^.

##### 6.1.3.3 (E)-1-(3-nitrophenyl)-3-(1-(prop-2-yn-1-yl)-1H-indol-3-yl)prop-2-en-1-one (4c)

Yellow solid, yield: 40.1 %, R_f_ = 0.75 (Ethylacetate : Hexane = 50 : 50) mp: 200.1 °C, IR (neat): ν (cm^-1^) 3235 (≡C-H)_str_, 1641 (C=C)_str._ chalcone ^1^H NMR (700 MHz, DMSO-*d_6_*) (δ, ppm): 8.78 – 8.77 (m, 1H), 8.63 – 8.61 (m, 1H, Ar-H), 8.49 (ddd, *J* = 8.1, 2.3, 0.9 Hz, 1H, Ar-*H*), 8.28 (s, 1H, Ar-*H*), 8.18 (d, *J* = 7.8 Hz, 1H, Ar-*H*), 8.13 (d, *J* = 15.4 Hz, 1H, Ar-*H*), 7.89 (t, *J* = 7.9 Hz, 1H, Ar-*H*), 7.74 (d, *J* = 15.4 Hz, 1H, Ar-*H*), 7.67 (d, *J* = 8.1 Hz, 1H, Ar-*H*), 7.39 – 7.36 (m, 1H, Ar- *H*), 7.35 – 7.32 (m, 1H, Ar-*H*), 5.22 (d, *J* = 2.5 Hz, 2H, C*H_2_*), 3.55 (t, *J* = 2.5 Hz, ≡C-*H*). ^13^C NMR (176 MHz, DMSO-*d_6_*) (δ-ppm): 187.47, 148.66, 140.09, 137.43, 136.11, 134.96, 131.00, 127.18, 126.30, 123.67, 122.40, 121.24, 115.95, 113.12, 111.69, 87.73, 78.77, 77.01, 36.24., ESI-MS (*m/z*) calcd. for C_20_H_14_N_2_0_3_: 330.34;Found: 331.08 [M+H]^+^.

##### 6.1.3.4. (E)-1-(4-nitrophenyl)-3-(1-(prop-2-yn-1-yl)-1H-indol-3-yl)prop-2-en-1-one (4d)

Yellow solid, yield: 89.1 %, R_f_ = 0.75 (Ethylacetate : Hexane = 50 : 50), mp: 189.8 °C, IR (neat): ν (cm^-1^) 3267 (≡C-H)_str_, 1654 (C=C)_str._ chalcone, ^1^H NMR (700 MHz, DMSO-*d_6_*) (δ, ppm): 8.37 (dd, *J* = 22.9, 8.9 Hz, 2H, Ar-*H*), 8.26 (s, 1H, Ar-*H*), 8.18 (d, *J* = 7.8 Hz, 1H, C*H*), 8.10 (d, *J* = 15.4 Hz, 1H, Ar-*H*), 7.75 (d, *J* = 15.2 Hz, 3H, Ar-*H*), 7.68 (t, *J* = 11.3 Hz, 2H, C*H*), 7.35 (t, *J* = 7.9 Hz, 2H, Ar-*H*), 5.22 (d, *J* = 2.5 Hz, 1H, Ar-*H*), 3.55 (t, *J* = 2.5 Hz, ≡C-*H*). ^13^C NMR (176 MHz, DMSO-*d_6_*) (δ-ppm): 187.77, 149.47, 143.38, 139.75, 137.00, 135.90, 129.56, 125.73, 123.81, 123.23, 121.97, 120.84, 115.89, 112.65, 111.23, 78.26, 76.54, 35.76. ESI-MS (*m/z*) calcd. for C_20_H_14_N_2_O_3_: 330.3; Found: 331.17 [M+H]^+^.

##### 6.1.3.5. (E)-1-(2-bromophenyl)-3-(1-(prop-2-yn-1-yl)-1H-indol-3-yl)prop-2-en-1-one (4e)

Yellow solid, yield: 76.9 %, R_f_ = 0.78 (Ethylacetate : Hexane = 50 : 50),mp: 90.1 °C, IR (neat): ν (cm^-1^) 3278 (≡C-H)_str_, 1663 (C=C)_str._ chalcone, ^1^H NMR (700 MHz, DMSO-*d_6_*) (δ, ppm): 8.12 (s, 1H, Ar-*H*), 7.96 (d, *J* = 7.9 Hz, 1H, C*H*), 7.75 (dd, *J* = 8.0, 0.7 Hz, 1H, Ar-*H*), 7.65 (d, *J* = 8.2 Hz, 1H, C*H*), 7.59 (d, *J* = 16.0 Hz, 1H, Ar-*H*), 7.52 (dd, J = 4.7, 1.5 Hz, 1H, Ar-*H*), 7.46 (dd, *J* = 2.1, 1.0 Hz, 1H, Ar-*H*), 7.35 (t, *J* = 7.2 Hz, 1H, Ar-*H*), 7.28 (t, J = 7.1 Hz, 1H, Ar-*H*), 7.04 (d, *J* = 16.0 Hz, 1H, Ar-*H*), 5.16 (d, *J* = 2.4 Hz, 2H, C*H_2_*), 3.51 (t, *J* = 2.5 Hz, ≡C-*H*). ^13^C NMR (176 MHz, DMSO-*d_6_*) (δ-ppm): 193.78, 141.50, 140.78, 137.00, 135.62, 133.02, 131.29, 129.00, 127.79, 125.49, 123.19, 122.02, 120.79, 120.44, 118.63, 111.99, 111.26, 78.21, 76.47, 35.71. ESI-MS (*m/z*) calcd. for C_20_H_14_BrNO: 364.2; Found: 366.10 [M+2H]^+^.

##### 6.1.3.6. (E)-1-(3-bromophenyl)-3-(1-(prop-2-yn-1-yl)-1H-indol-3-yl)prop-2-en-1-one (4f)

Yellow solid, yield: 45 %, R_f_ = 0.78 (Ethylacetate : Hexane = 50 : 50), mp: 148 °C, IR (neat): ν (cm^-1^) 3235 (≡C-H)_str_, 1641 (C=C)_str._ chalcone, ^1^H NMR (700 MHz, DMSO-*d_6_*) (δ, ppm): 8.24 (dd, *J* = 4.5, 2.7 Hz, 2H, Ar-H), 8.16 (t, *J* = 7.0 Hz, 2H, Ar-*H*), 8.06 (d, *J* = 15.3 Hz, 1H, C*H*), 7.86 – 7.85 (m, 1H, Ar-*H*), 7.66 (dd, *J* = 7.9, 3.4 Hz, 2H, Ar-*H*), 7.56 – 7.53 (m, 1H, Ar-*H*), 7.37 – 7.35 (m, 1H, Ar-*H*), 5.31 (d, *J* = 2.5 Hz, 1H, Ar-*H*), 5.21 (d, *J* = 2.5 Hz, 2H, C*H_2_*), 3.54 (t, *J* = 2.5 Hz, 1H, ≡C-*H*). ^13^C NMR (176 MHz, DMSO-*d_6_*) (δ-ppm): 187.45, 140.43, 138.79, 136.89, 135.64, 135.16, 135.11, 134.89, 134.75, 131.02, 130.95, 130.58, 130.46, 127.29, 126.91, 125.86, 123.10, 122.23, 121.82, 121.42, 120.72, 118.28, 115.74, 114.95, 112.61, 111.11, 110.97, 78.33, 76.50, 76.46, 35.71. ESI-MS (*m/z*) calcd. for C_20_H_14_BrNO: 364.2; Found: 366.09 [M+H]^+^.

##### 6.1.3.7. (E)-1-(4-bromophenyl)-3-(1-(prop-2-yn-1-yl)-1H-indol-3-yl)prop-2-en-1-one (4g)

Yellow solid, yield: 44.4 %, R_f_ = 0.75 (Ethylacetate : Hexane = 50 : 50), mp: 193.5 °C, IR (neat): ν (cm^-1^) 3227 (≡C-H)_str_, 1642 (C=C)_str._ chalcone, ^1^H NMR (700 MHz, DMSO-*d_6_*) (δ, ppm): 8.22 (s, 1H, Ar-*H*), 8.15 (d, *J* = 7.9 Hz, 1H, Ar-*H*), 8.10 – 8.08 (m, 2H, Ar-*H*), 8.05 (d, *J* = 15.4 Hz, 1H, Ar-*H*), 8.02 – 8.00 (m, 1H, Ar-*H*), 7.79 – 7.76 (m, 3H, Ar-*H*), 7.65 (t, *J* = 7.5 Hz, 2H, Ar-*H*), 5.21 (d, *J* = 2.5 Hz, 2H, C*H_2_*), 3.53 (t, *J* = 2.5 Hz, 1H, ≡C-*H*). ^13^C NMR (176 MHz, DMSO-*d_6_*) (δ-ppm): 188.37, 188.30, 139.05, 137.78, 137.40, 135.97, 135.78, 135.69, 135.13, 132.25, 132.20, 130.75, 130.45, 126.98, 126.25, 123.57, 122.28, 121.84, 121.23, 118.75, 116.26, 115.53, 113.07, 111.61, 78.81, 76.92, 36.17. ESI-MS (*m/z*) calcd. for C_20_H_14_BrN0: 364.24; Found: 364.08 [M+H]^+^.

##### 6.1.3.8. (E)-1-(2-chlorophenyl)-3-(1-(prop-2-yn-1-yl)-1H-indol-3-yl)prop-2-en-1-one (4h)

Yellow solid, yield: 84.2 %, R_f_ = 0.75 (Ethylacetate : Hexane = 50 : 50), mp: 95.1 °C, IR (neat): ν (cm^-1^) 3261 (≡C-H)_str_, 1646 (C=C)_str._ chalcone, ^1^H NMR (700 MHz, DMSO-*d_6_*) (δ, ppm): 8.12 (s, 1H, Ar-*H*), 7.96 (d, *J* = 7.9 Hz, 1H, C*H*), 7.65 (d, J = 9.5 Hz, 2H, C*H*), 7.59 (d, *J* = 9.0 Hz, 1H, Ar-*H*), 7.56 (d, *J* = 7.8 Hz, 2H, Ar-*H*), 7.49 (t, *J* = 6.8 Hz, 1H, Ar-*H*), 7.35 (t, *J* = 7.2 Hz, 1H, Ar-*H*), 7.28 (t, *J* = 7.5 Hz, 1H, Ar-*H*), 7.07 (d, *J* = 16.0 Hz, 1H, Ar-*H*), 5.17 (d, *J* = 2.4 Hz, 2H, C*H_2_*), 3.51 (t, *J* = 2.5 Hz, 1H, ≡C-*H*). ^13^C NMR (176 MHz, DMSO-*d_6_*) (δ-ppm): 192.81, 140.55, 139.45, 137.00, 135.62, 131.27, 129.95, 129.80, 129.12, 127.34, 125.50, 123.18, 122.00, 120.97, 120.43, 111.97, 111.26, 78.21, 76.47, 35.71. ESI-MS (*m/z*) calcd. for C_20_H_14_ClN0: 319.8; Found: 320.13 [M+H]^+^.

##### 6.1.3.9. (E)-1-(3-chlorophenyl)-3-(1-(prop-2-yn-1-yl)-1H-indol-3-yl)prop-2-en-1-one (4i)

Yellow solid, yield: 76.51 %, R_f_ = 0.75 (Ethylacetate : Hexane = 50 : 50), mp: 144.6 °C, IR (neat): ν (cm^-1^) 3291 (≡C-H)_str_, 1652 (C=C)_str._ chalcone, ^1^H NMR (700 MHz, DMSO-*d_6_*) (δ, ppm): 8.25 (s, 1H, Ar-*H*), 8.16 (d, *J* = 7.8 Hz, 1H, C*H*), 8.13 – 8.12 (m, 2H, Ar-*H*), 8.06 – 8.05 (m, 1H, Ar-*H*), 7.95 (d, *J* = 7.9 Hz, 1H, CH), 7.66 (dd, *J* = 8.5, 3.3 Hz, 2H, Ar-*H*), 7.33 – 7.32 (m, 1H, Ar-*H*), 7.04 (d, *J* = 12.3 Hz, 1H, Ar-*H*), 5.21 (d, *J* = 2.5 Hz, 2H, CH_2_), 3.54 (t, *J* = 2.5 Hz, 1H, ≡C-*H*). ^13^C NMR (176 MHz, DMSO-*d_6_*) (δ-ppm): ^13^C NMR (176 MHz, DMSO-*d_6_*) (δ- ppm): 187.51, 141.10, 140.24, 138.81, 136.90, 135.64, 135.52, 135.20, 134.77, 133.71, 132.22, 132.01, 130.77, 130.70, 129.03, 127.72, 127.56, 126.91, 126.57, 125.85, 123.11, 122.84, 121.82, 121.42, 120.75, 118.28, 115.77, 114.98, 112.61, 111.12, 110.97, 78.34, 76.50, 76.4690 35.8504 35.7217. ESI-MS (*m/z*) calcd. for C_20_H_14_ClNO: 319.18; Found: 320.11 [M+H]^+^.

##### 6.1.3.10. (E)-1-(4-chlorophenyl)-3-(1-(prop-2-yn-1-yl)-1H-indol-3-yl)prop-2-en-1-one (4j)

Yellow solid, yield: 57.8 %, R_f_ = 0.75 (Ethylacetate : Hexane = 50 : 50), mp: 192.1 °C, IR (neat): ν (cm^-1^) 3267 (≡C-H)_str_, 1652 (C=C)_str._ chalcone, ^1^H NMR (700 MHz, DMSO-*d_6_*) (δ, ppm): 8.22 (s, 1H, Ar-*H*), 8.18 – 8.16 (m, 2H, Ar-*H*), 8.10 – 8.08 (m, 1H, Ar-*H*), 8.05 (d, *J* = 15.4 Hz, 1H, C*H*), 7.69 (s, 1H, Ar-*H*), 7.66 (d, *J* = 8.6 Hz, 2H, C*H*), 7.64 (d, *J* = 1.8 Hz, 1H, Ar-*H*), 7.63 (d, *J* = 2.0 Hz, 1H, Ar-*H*), 7.36 (t, *J* = 7.1 Hz, 1H, Ar-*H*), 5.21 (d, *J* = 2.5 Hz, 2H, C*H_2_*), 3.53 (t, *J* = 2.5 Hz, 1H, ≡C-*H*). ^13^C NMR (176 MHz, DMSO-*d_6_*) (δ-ppm): 187.63, 138.55, 137.38, 136.98, 136.93, 135.28, 135.21, 134.64, 130.14, 129.84, 128.83, 128.78, 125.78, 123.10, 121.80, 121.36, 120.76, 118.28, 115.81, 115.12, 112.60, 111.13, 110.94, 78.35, 76.45, 35.70. ESI-MS (*m/z*) calcd. for C_20_H_14_ClN0: 319.8; Found: 320.11 [M+H]^+^.

##### 6.1.3.11. (E)-1-(2,4-dichlorophenyl)-3-(1-(prop-2-yn-1-yl)-1H-indol-3-yl)prop-2-en-1-one (4k)

Yellow solid, yield: 56.3 %, R_f_ = 0.75 (Ethylacetate : Hexane = 50 : 50), mp: 147.9 °C, IR (neat): ν (cm^-1^) 3283 (≡C-H)_str_, 1648 (C=C)_str._ chalcone, ^1^H NMR (700 MHz, DMSO-*d_6_*) (δ, ppm): 8.14 (s, 1H, Ar-*H*), 7.98 (d, *J* = 8.0 Hz, 1H, C*H*), 7.78 (d, *J* = 1.6 Hz, 1H, C*H*), 7.67 – 7.64 (m, 2H, Ar-*H*), 7.59 – 7.58 (m, 2H, Ar-*H*), 7.35 (t, *J* = 8.1 Hz, 1H, Ar-*H*), 7.28 (t, *J* = 7.5 Hz, 1H, Ar-*H*), 7.05 (d, *J* = 16.0 Hz, 1H, Ar-*H*), 5.17 (d, *J* = 2.5 Hz, 2H,C*H_2_*), 3.52 (t, *J* = 2.5 Hz, 1H, ≡C-*H*). ^13^C NMR (176 MHz, DMSO-*d_6_*) (δ-ppm): 192.42, 141.71, 138.75, 137.5043 136.37, 135.42, 131.56, 131.01, 129.98, 128.06, 125.95, 123.70, 122.52, 121.1580 121.02, 112.52, 111.74, 78.65, 77.02, 36.21. ESI-MS (*m/z*) calcd. for C_20_H_13_Cl_2_NO: 354.2; Found: 356.21 [M+H]^+^.

##### 6.1.3.12. (E)-1-(2-methoxyphenyl)-3-(1-(prop-2-yn-1-yl)-1H-indol-3-yl)prop-2-en-1-one (4l)

Cream solid, yield: 48 %, R_f_ = 0.78 (Ethylacetate : Hexane = 50 : 50), mp: 109 °C, IR (neat): ν (cm^-1^) 3296 (≡C-H)_str_, 1655 (C=C)_str._ chalcone, ^1^H NMR (700 MHz, DMSO-*d_6_*) (δ, ppm): 8.37 (s, 1H, Ar-H), 8.13 (d, *J* = 7.8 Hz, 1H, C*H*), 7.67 (d, J = 8.2 Hz, 1H, C*H*), 7.38 – 7.36 (m, 1H, Ar-*H*), 7.32 – 7.29 (m, 3H, Ar-*H*), 5.26 (d, *J* = 2.5 Hz, 2H, CH_2_), 3.57 (t, J = 2.5 Hz, 1H, ≡C-*H*), 3.33 (s, 6H, C*H_3_*). ^13^C NMR (176 MHz, DMSO-*d_6_*) (δ-ppm): 184.92, 140.15, 136.58, 124.68, 123.73, 122.79, 121.12, 117.55, 111.22, 78.01, 76.83, 36.02. ESI-MS (*m/z*) calcd. For C_21_H_17_NO_2_: 315.4; Found: 314.97 [M-H]^+^.

##### 6.1.3.13. (E)-1-(3-methoxyphenyl)-3-(1-(prop-2-yn-1-yl)-1H-indol-3-yl)prop-2-en-1-one (4m)

Yellow solid, yield: 61.05 %, R_f_ = 0.75 (Ethylacetate : Hexane = 50 : 50), mp: 118.7 °C, IR (neat): ν (cm^-1^) 3259 (≡C-H)_str_, 1646 (C=C)_str._ chalcone, ^1^H NMR (700 MHz, DMSO-*d_6_*) (δ, ppm): 8.37 (s, 1H, Ar-*H),* 8.21 (s, 1H, Ar-*H*), 8.13 (d, *J* = 7.9 Hz, 1H, C*H*), 8.03 (d, *J* = 15.5 Hz, 1H, Ar-*H*), 7.75 (d, *J* = 7.7 Hz, 1H, C*H*), 7.66 (d, J = 5.6 Hz, 2H, Ar-H), 7.36 (d, *J* = 8.3 Hz, 1H, Ar-*H*), 7.23 (dd, *J* = 7.9, 2.3 Hz, 1H, Ar-H), 5.26 (d, *J* = 2.4 Hz, 2*H*), 5.20 (d, *J* = 2.4 Hz, 2*H*), 3.86 (d, *J* = 6.6 Hz, 3H, C*H_3_*), 3.57(t, *J* = 2.1 Hz, 1H, ≡C-*H*). ^13^C NMR (176 MHz, DMSO-*d_6_*) (δ-ppm): 188.57, 184.92, 159.49, 140.14, 139.84, 138.01, 136.88, 136.57, 134.82, 129.85, 124.68, 123.73, 123.02, 122.79, 121.76, 121.12, 120.69, 120.62, 118.45, 117.55, 116.39, 112.72, 112.55, 111.226, 111.10, 76.83, 76.40, 55.31, 36.01, 35.67. ESI-MS (*m/z*) calcd. for C_21_H_17_NO_2_: 315.4; Found: 316.16[M+H]^+^.

##### 6.1.3.14. (E)-1-(4-methoxyphenyl)-3-(1-(prop-2-yn-1-yl)-1H-indol-3-yl)prop-2-en-1-one (4n)

Yellow solid, yield: 36.6 %, R_f_ = 0.80 (Ethylacetate : Hexane = 50 : 50), mp: 179.3 °C, IR (neat): ν (cm^-1^) 3235 (≡C-H)_str_, 1642 (C=C)_str._ chalcone. ^1^H NMR (700 MHz, DMSO-*d_6_*) (δ, ppm): 8.17 (s, 1H, Ar-*H*), 8.15 (d, *J* = 8.9 Hz, 2H, Ar-*H*), 8.13 (d, J = 7.8 Hz, 1H, C*H*), 7.99 (d, *J* = 15.4 Hz, 1H, C*H*), 7.70 (d, *J* = 15.5 Hz, 1H, Ar-H), 7.65 (d, *J* = 8.1 Hz, 1H, Ar-H), 7.35 (t, *J* = 7.5 Hz, 1H, Ar-*H*), 7.31 (t, *J* = 7.4 Hz, 1H, Ar-*H*), 7.10 (t, J = 5.8 Hz, 2H, Ar-*H*), 5.20 (d, *J* = 2.4 Hz, 2H, C*H_2_*), 3.88 (s, 3H, C*H_3_*), 3.52 (t, *J* = 2.5 Hz, 1H, ≡C-*H*). ^13^C NMR (176 MHz, DMSO-*d_6_*) (δ- ppm): 187.14, 162.75, 137.04, 136.85, 134.42, 131.07, 130.48, 130.20, 125.84, 122.94, 121.62, 120.64, 116.26, 113.94, 113.91, 112.59, 111.05, 78.44, 76.34, 55.49, 35.62. ESI-MS (*m/z*) calcd. for C_21_H_17_N0_2_: 315.37;Found: 316.15 [M+H]^+^.

##### 6.1.3.15. (E)-1-(3,4-dimethoxyphenyl)-3-(1-(prop-2-yn-1-yl)-1H-indol-3-yl)prop-2-en-1-one (4o)

Yellow solid, yield: 35.4 %, R_f_ = 0.80 (Ethylacetate : Hexane = 50 : 50), mp: 173.8 °C, IR (neat): ν (cm^-1^) 3259 (≡C-H)_str_, 1642 (C=C)_str._ chalcone. ^1^H NMR (700 MHz, DMSO-*d_6_*) (δ, ppm): 8.18 (s, 1H, Ar-*H*), 8.12 (d, *J* = 7.8 Hz, 1H, C*H*), 7.99 (d, *J* = 15.4 Hz, 1H, C*H*), 7.88 (dd, *J* = 8.4, 2.0 Hz, 1H, Ar-*H*), 7.71 (d, *J* = 15.5 Hz, 1H, Ar-*H*), 7.65 (d, *J* = 8.1 Hz, 1H, Ar-*H*), 7.60 (d, *J* = 1.9 Hz, 1H, Ar-*H*), 7.35 (t, *J* = 7.5 Hz, 1H, Ar-*H*), 7.31 (t, *J* = 7.0 Hz, 1H, Ar-*H*), 7.12 (d, *J* = 8.4 Hz, 1H, Ar-*H*), 5.20 (d, *J* = 2.4 Hz, 2H, C*H_2_*), 3.88 (d, *J* = 6.1 Hz, 6H, C*H_3_*), 3.52 (t, *J* = 2.5 Hz, 1H, ≡C-*H*). ^13^C NMR (176 MHz, DMSO-*d_6_*) (δ-ppm): 187.20, 152.71, 148.74, 136.88, 136.84, 134.26, 131.18, 125.90, 122.94, 122.74, 121.63, 120.57, 116.33, 112.59, 111.05, 110.85, 110.59, 78.46, 76.34, 55.73, 55.54, 35.63. ESI-MS (*m/z*) calcd. for C_22_H_19_N0_3_: 345.39; Found: 346.14 [M+H]^+^.

##### 6.1.3.16. (E)-3-(1-(prop-2-yn-1-yl)-1H-indol-3-yl)-1-(thiophen-2-yl)prop-2-en-1-one (4p)

Yellow solid, yield: 42.9 %, R_f_ = 0.75 (Ethylacetate : Hexane = 50 : 50), mp: 162.4 °C, IR (neat): ν (cm^-1^) 3270 (≡C-H)_str_, 1642 (C=C)_str._ chalcone, ^1^H NMR (700 MHz, DMSO-*d_6_*) (δ, ppm): 8.28 (dd, *J* = 3.8, 1.0 Hz, 1H, Ar-*H*), 8.19 – 8.17 (m, 2H, Ar-H,C*H*), 8.01 (dd, *J* = 4.1, 1.8 Hz, 1H, Ar- *H*), 7.99 – 7.98 (m, 1H,C*H*), 7.67 – 7.63 (m, 2H, Ar-*H*), 7.61 (s, 1H, Ar-*H*), 7.37 – 7.35 (m, 1H, Ar-*H*), 7.32 (dd, *J* = 5.9, 2.2 Hz, 1H, Ar-*H*), 7.28 – 7.27 (m, 1H, Ar-*H*), 5.20 (s, 1H, Ar-*H)*, 3.53 (d, *J* = 2.5 Hz, 1H, ≡C-*H*). ^13^C NMR (176 MHz, DMSO-*d_6_*) (δ-ppm): 181.60, 181.41, 147.07, 146.10, 137.12, 136.89, 135.44, 134.95, 134.81, 134.46, 134.35, 134.18, 132.35, 131.53, 128.84, 128.75, 125.76, 123.04, 122.72, 121.73, 121.30, 120.79, 118.22, 116.02, 115.08, 112.39, 111.07, 110.89, 78.37, 76.40, 35.66. ESI-MS (*m/z*) calcd. for C_18_H_13_NOS: 291.4; Found: 292.11 [M+H]^+^.

##### 6.1.3.17. (E)-1-(3,5-bis(benzyloxy)phenyl)-3-(1-(prop-2-yn-1-yl)-1H-indol-3-yl)prop-2-en-1- one (4q)

Yellow solid, yield: 55.3 %, R_f_ = 0.75 (Ethylacetate : Hexane = 50 : 50), mp: 147.0 °C, IR (neat): ν (cm^-1^) 3278 (≡C-H)_str_, 1654 (C=C)_str._ chalcone. ^1^H NMR (700 MHz, DMSO-*d_6_*) (δ, ppm): 8.37 (s, 1H, Ar-*H*), 8.22 (s, 3H, Ar-*H*), 8.07 (d, *J* = 7.7 Hz, 2H, Ar-H, C*H*), 8.01 (d, *J* = 15.4 Hz, 2H, Ar-*H*), 7.65 (d, *J* = 8.1 Hz, 3H, Ar-H, C*H*), 7.58 (d, *J* = 15.5 Hz, 3H, Ar-*H*), 7.18 (s, 1H, Ar-*H*), 6.97 (d, *J* = 2.2 Hz, 3H, Ar-*H*), 5.30 (s, 1H, Ar-*H*), 5.26 (d, *J* = 2.5 Hz, 1H, Ar-*H*), 5.17 (s, 3H,CH_2_, Ar-*H*), 5.11 (s, 1H, Ar-H), 3.53 (s, 3H, Ar-H, ≡C-*H*). ^13^C NMR (176 MHz, DMSO-*d_6_*) (δ-ppm): 188.2847 184.92, 159.64, 140.52, 138.06, 136.81, 136.70, 134.66, 128.47, 128.43, 127.92, 127.79, 127.77, 127.75, 125.92, 123.73, 123.01, 122.79, 121.75, 121.12, 120.52, 116.28, 112.54, 111.08, 107.09, 107.02, 105.97, 76.41, 69.57, 69.54, 35.69. ESI-MS (*m/z*) calcd. for C_34_H_27_NO_3_: 497.6; Found: 498.21 [M+H]^+^.

##### 6.1.3.18. (E)-3-(1-(prop-2-yn-1-yl)-1H-indol-3-yl)-1-(o-tolyl)prop-2-en-1-one (4r)

Yellow solid, yield: 34.4 %, R_f_ = 0.75 (Ethylacetate : Hexane = 50 : 50), mp: 221.5 °C, IR (neat): ν (cm^-1^) 3221 (≡C-H)_str_, 1643 (C=C)_str._ chalcone. ^1^H NMR (700 MHz, DMSO-*d_6_*) (δ-ppm): 8.10 (s, 1H, Ar-*H*), 8.12 (d, *J* = 7.1 Hz, 2H, CH, Ar-*H*), 8.03 (d, *J* = 15.5 Hz, 2H,Ar-*H*), 7.73 (d, *J* = 15.1 Hz, 1H, C*H*), 7.61 – 7.69 (m, 1H, Ar-*H*), 7.46 (t, *J* = 7.4 Hz, 2H, Ar-*H*), 7.33 (t, *J* = 7.2 Hz, 1H, Ar- *H*), 7.31 (d, *J* = 7.7 Hz, 1H, Ar-*H*), 5.22(d, *J* = 2.4 Hz, 2H, C*H_2_*), 3.53 (t, *J* = 2.5 Hz, 1H, ≡C-*H*), 2.12(s, 3H, C*H_3_*). ^13^C NMR (176 MHz, DMSO-*d_6_*) (δ-ppm): 188.51, 138.33, 138.21, 135.53, 134.12, 133.51, 129.45, 128.41, 128.12, 125.31, 124.44, 121.75, 120.52, 116.25, 112.57, 112.01, 78.32, 76.40, 35.77. ESI-MS (*m/z*) calcd. for C_21_H_17_NO: 299.37; Found: 300.52 [M+H]^+^.

##### 6.1.3.19. (E)-1-(4-fluorophenyl)-3-(1-(prop-2-yn-1-yl)-1H-indol-3-yl)prop-2-en-1-one (4s)

Yellow solid, yield: 66.7 %, R_f_ = 0.75 (Ethylacetate : Hexane = 50 : 50), mp: 168.6 °C, IR (neat): ν (cm^-1^) 3265 (≡C-H)_str_, 1652 (C=C)_str._ chalcone. ^1^H NMR (700 MHz, DMSO-*d_6_*) (δ, ppm): 8.24 (dd, *J* = 8.8, 5.6 Hz, 2H, Ar-*H*), 8.20 (s, 1H, Ar-*H)*, 8.16 (d, *J* = 7.9 Hz, 1H, C*H*), 8.03 (d, *J* = 15.4 Hz, 1H, C*H*), 7.69 (d, *J* = 15.4 Hz, 1H, Ar-*H*), 7.66 (d, *J* = 8.2 Hz, 1H, Ar-*H*), 7.40 (t, *J* = 8.8 Hz, 2H, Ar-*H*), 7.37 – 7.35 (m, 1H, Ar-*H*), 7.31 (t, *J* = 7.1 Hz, 1H, Ar-*H*), 5.21 (d, *J* = 2.5 Hz, 2H, C*H_2_*), 3.53 (t, *J* = 2.5 Hz, 1H, ≡C-*H*). ^13^C NMR (176 MHz, DMSO-*d_6_*) (δ-ppm): 187.79, 165.88, 164.46, 138.67, 137.38, 135.49, 135.41, 135.36, 131.61, 131.56, 131.25, 126.27, 123.53, 122.22, 121.23, 116.40, 116.18, 116.06, 113.05, 111.58, 78.84, 76.89, 36.15. ESI-MS (*m/z*) calcd. for C_20_H_14_FN0: 303.3; Found: 304.23 [M+H]^+^.

##### 6.1.3.20. 1-(prop-2-yn-1-yl)-1H-indole-3-carbaldehyde (8)

Cream solid, yield: 90.4 %, R_f_ = 0.75 (Ethylacetate : Hexane = 50 : 50), mp: 108.9 °C, IR (neat): ν (cm^-1^) 3298 (≡C-H)_str_, 1641 (C=C)_str._ chalcone, ^1^H NMR (700 MHz, DMSO-*d_6_*) (δ, ppm): 9.95 (s, 1H, CHO), 8.37 (s, 1H, Ar- *H*), 8.14 (d, *J* = 7.8 Hz, 1H, Ar-*H*), 7.67 (d, *J* = 8.2 Hz, 1H, Ar-*H*), 7.38 – 7.36 (m, 1H, Ar-*H*), 7.32 – 7.29 (m, 1H, Ar-*H*), 5.26 (d, *J* = 2.4 Hz, 2H, C*H_2_*), 3.57 (t, *J* = 2.5 Hz, 1H, ≡C-*H*). ^13^C NMR (176 MHz, DMSO-*d_6_*) (δ-ppm): 184.92, 140.15, 136.57, 124.68, 123.73, 122.79, 121.12, 117.55, 111.22, 78.00, 76.83, 36.01. ESI-MS (*m/z*) calcd. for C_12_H_9_NO: 183.21; Found: 184.34 [M+H]^+^.

#### 6.1.4. Synthesis of azide (6)

In a dried round bottom flask, to a solution of 4,7-dichloroquinoline (**5**) (1.0 mmol) in anhyd. DMF (5 ml), sodium azide (6.0 mmol) was added and the reaction mixture was allowed to stir at 85°C for 3 h. After completion of the reaction, the mixture was poured into water, extracted with chloroform (2×30 mL), dried over sodium sulphate and concentrated in vacuum to give azide which was used further without purification [47].

#### 6.1.5. General procedure for the synthesis of triazole hybrids (7a-s) and 9

In a dried round bottom flask, Indole-chalcone alkynes **4a-s/**indole alkyne **8** (0.20 g, 0.61 mmol) and azide **6** (0.093 g, 0.61 mmol) were dissolved in a mixture of solvent system, THF/H_2_O (1:2, 9 ml). After that, sodium ascorbate (0.063 g, 0.32 mmol) was added followed by the addition of CuSO_4_.5H_2_O (0.026 g, 0.11 mmol). The reaction mixture was stirred till the complete disappearance of starting materials. The solid obtained was filtered and washed with water. The crude product was purified by silica gel chromatography eluting with a solution of methanol (5%) in dichloromethane to yield pure compounds [29].

##### 6.1.5.1. (E)-3-(1-((1-(7-chloroquinolin-4-yl)-1H-1,2,3-triazol-4-yl)methyl)-1H-indol-3-yl)-1- phenylprop-2-en-1-one (7a)

Yellow solid, yield: 62 %, R_f_ = 0.69 (Methanol:DCM = 5 : 95), mp: 225.1-225.3 °C, IR (neat): ν (cm^-1^) 3152 (C-H)_str._ triazole ring, 1648 (C=C)_str._ chalcone, ^1^H NMR (700 MHz, DMSO-*d_6_*) (δ, ppm): 9.12 (t, *J* = 4.83 Hz, 1H, CQ-*H*), 8.94 (s, 1H, NC-*H*_indole_), 8.32 (s, 1H, Ar-H), 8.28 (d, *J* = 2.03 Hz,1H, C-*H*_triazole_), 8.13-8.11 (m, 3H, Ar-*H*), 8.05 (m, 1H, Ar-*H*), 7.98 (d, *J* = 9.17 Hz, 1H, Ar-*H*), 7.83-7.81 (m, 3H, Ar-*H*), 7.78 (d, *J* = 1.89 Hz, 1H, Ar- *H*), 7.68 (d, *J* = 15.5 Hz, 1H, Ar-*H*), 7.64 (s, 1H, Ar-*H*), 7.55-7.54 (m, 3H, Ar-*H*), 5.74 (s, 2H, C*H_2_*). ^13^C NMR (176 MHz, DMSO-*d_6_*) (δ, ppm): 189.26, 152.83, 149.85, 143.92, 140.75, 138.83, 138.65, 137.71, 136.05, 135.84, 132.96, 129.47, 129.22, 129.18, 128.63, 128.38, 126.26, 126.33, 125.88, 123.49, 122.13, 121.14, 120.73, 117.67, 117.63, 116.51, 112.99, 111.77. ESI-MS (*m/z*) calcd. for C_29_H_20_ClN_5_O: 489.96; Found: 462.24 [M^_^N_2_+1]^+^, 464.27 [M-N_2_+2+1]^+^, 465.30[M-N_2_+2+2]^+^. Purity: UP-LC 91.04%.

##### 6.1.5.2 (E)-3-(1-((1-(7-chloroquinolin-4-yl)-1H-1,2,3-triazol-4-yl)methyl)-1H-indol-3-yl)-1-(2- nitrophenyl)prop-2-en-1-one (7b)

Yellow solid, yield: 73 %, R_f_ = 0.85 (Methanol : DCM = 5 : 95), mp: 210.4-210.7 °C, IR (neat): ν (cm^-1^) 3136 (C-H)_str._ triazole ring, 1611 (C=C)_str._ chalcone, ^1^H NMR (700 MHz, DMSO-*d_6_*) (δ, ppm): 9.11 (d, *J* =4.69 Hz, 1H, CQ-*H*), 8.91 (s, 1H, NC- *H*_indole_), 8.26 (d, *J* = 1.75 Hz, 1H, Ar-*H*), 8.22 (s, 1H, C-*H*_triazole_), 8.15 (d, *J* = 8.19 Hz, 1H, Ar- H), 7.96 (t, *J* = 7.56 Hz, 2H, Ar-*H*), 7.87 (d, *J* = 14.98 Hz, 1H, Ar-*H*), 7.81 (t, *J* = 4.48 Hz, 2H, Ar-*H*), 7.76 (d, J = 9.66 Hz, 1H, Ar-*H*), 7.74 (d, J = 10.15 Hz, 2H, Ar-*H*), 7.66 (s, 1H, Ar-*H*), 7.30 (t, J = 7.56 Hz, 1H, Ar-*H*), 7.25 (t, *J* = 7.63 Hz, 1H, Ar-*H*), 7.07 (d, J = 16.1 Hz, 1H, Ar-*H*), 5.70 (s, 2H, C*H_2_*). ^13^C NMR (176 MHz, DMSO-*d_6_*) (δ, ppm): 192.03, 152.81, 149.84, 147.53, 143.74, 141.19, 140.72, 137.79, 136.77, 136.44, 135.83, 134.58, 131.46, 129.57, 129.44, 128.61, 126.68, 125.99, 125.87, 124.95, 123.65, 122.36, 120.99, 120.69, 120.36, 117.59, 112.34, 111.89, 41.51. ESI-MS (*m/z*) calcd. for C_29_H_19_ClN_6_O_3_: 534.96, Found: 507.26 [M-N_2_+1]^+^, 509.25 [M- N_2_+2+1]^+^, 465.30 [M-N_2_+2+2]^+^. Purity: UP-LC 98.88 %.

##### 6.1.5.3 (E)-3-(1-((1-(7-chloroquinolin-4-yl)-1H-1,2,3-triazol-4-yl)methyl)-1H-indol-3-yl)-1-(3- nitrophenyl)prop-2-en-1-one (7c)

Yellow solid, yield: 71%, R_f_ = 0.82 (Methanol : DCM = 5 : 95), mp: 236.1-236.9 °C, IR (neat): ν (cm^-1^) 3157 (C-H)_str._ triazole ring, 1657 (C=C)_str._ chalcone. ^1^H NMR (700 MHz, DMSO-*d_6_*) (δ, ppm): 9.12 (s, 1H, CQ-*H*), 8.94 (s, 1H, NC-*H*_indole_), 8.75 (s, 1H, Ar-*H*), 8.59 (d, *J* = 6.86 Hz,1H, C-*H*_triazole_), 8.45 (d, J = 7.63 Hz, 1H, Ar-*H*), 8.40 (s, 1H, Ar-*H*), 8.26 (s, 1H, Ar-*H*), 8.16 – 8.12 (m, 3H, Ar-*H*), 7.97 (d, *J* = 8.75Hz, 1H, Ar-*H*), 7.86 – 7.82 (m, 3H, Ar-*H*), 7.76 – 7.70 (m, 2H, Ar-*H*), 7.35 – 7.30 (m, 2H, Ar-*H*), 5.74 (s, 2H, C*H_2_*). ^13^C NMR (176 MHz, DMSO-*d_6_*) (δ, ppm): 187.39, 152.83, 149.84, 148.64, 143.80, 140.74, 140.23, 140.09, 137.76, 136.81, 135.84, 134.92, 130.99, 129.46, 128.62, 127.16, 126.69, 126.36, 125.87, 123.65, 122.96, 122.31, 121.19, 120.72, 117.63, 115.67, 113.06, 111.87, 41.61. ESI-MS (*m/z*) calcd. for C_29_H_19_ClN_6_O_3_: 534.96, Found: 507.26 [M-N_2_+1]^+^, 509.25 [M-N_2_+2+1]^+^, 510.26 [M- N_2_+2+2]^+^. Purity: UP-LC 94.99 %.

##### 6.1.5.4. (E)-3-(1-((1-(7-chloroquinolin-4-yl)-1H-1,2,3-triazol-4-yl)methyl)-1H-indol-3-yl)-1-(4- nitrophenyl)prop-2-en-1-one (7d)

Orange solid, yield: 67 %, R_f_ = 0.79 (Methanol : dichloromethane = 5 : 95), mp: 247.5-247.9 °C, IR (neat): ν (cm^-1^) 3142 (C-H)_str._ triazole ring, 1641 (C=C)_str._ chalcone. ^1^H NMR (700 MHz, DMSO-*d_6_*) (δ, ppm): 9.12 (d, 1H, *J* = 4.62 Hz, CQ- *H*), 8.94 (s, 1H, NC-*H*_indole_), 8.36 (t, *J* = 4.55 Hz, 3H, C-*H*_triazole_, Ar-*H*), 8.32 (d, *J* = 8.75 Hz, 2H, C*H*), 8.26 (s, 1H, Ar-*H*), 8.15 (d, *J* = 7.91 Hz, 1H, Ar-*H*), 8.10 (d, *J* = 15.14 Hz, 1H, Ar-*H*), 7.97 (d, *J* = 9.03 Hz, 1H, Ar-*H*), 7.83 (t, *J* = 7.07 Hz, 2H, Ar-*H*), 7.77-7.75 (m, 1H, Ar-*H*), 7.66 (d, *J* = 15.4 Hz, 1H, Ar-*H*), 7.35 (t, *J* = 7.35 Hz, 1H, Ar-*H*), 7.30 (t, *J* = 7.56 Hz, 1H, Ar-*H*), 5.75 (s, 2H, C*H_2_*). ^13^C NMR (176 MHz, DMSO-*d_6_*) (δ, ppm): 188.1743, 152.81, 149.92, 149.84, 143.8, 143.78, 140.73, 140.37, 137.81, 137.07, 135.84, 129.99, 129.45, 128.62, 126.69, 126.26, 125.85, 124.27, 123.68, 122.35, 121.27, 120.70, 117.61, 116.11, 113.08, 111.89, 41.59. ESI-MS (*m/z*) calcd. for C_29_H_19_ClN_6_O_3_: 534.95; Found: 535.07 [M+H]^+^, 536.36 [M+H+1]^+^. Purity: UP- LC 92.70 %.

##### 6.1.5.5. (E)-1-(2-bromophenyl)-3-(1-((1-(7-chloroquinolin-4-yl)-1H-1,2,3-triazol-4-yl)methyl)- 1H-indol-3-yl)prop-2-en-1-one (7e)

Orange solid, yield: 54 %, R_f_ = 0.67 (Methanol : DCM = 5 : 95), mp: 190.3-193.0 °C, IR (neat): ν (cm^-1^) 3151 (C-H)_str._ triazole ring, 1562 (C=C)_str._ chalcone. ^1^H NMR (700 MHz, DMSO-d_6_) (δ, ppm): 9.12 (d, *J* = 4.62 Hz, 1H, CQ-*H*), 8.91 (s, 1H, NC-*H*_indole_), 8.26 (s, 1H, Ar-*H*), 8.22 (s,1H, C-*H*_triazole_), 7.97 (d, 1H_trans indole side_, *J* = 9.1 Hz), 7.95 (m, 1H_trans keto side_ *J* = 7.98 Hz), 7.81 (t, *J* = 4.62 Hz, 2H, Ar-*H*), 7.75 (d, *J* = 9.03, 1H, Ar- *H*), 7.73 (d, *J* = 7.93 Hz, 1H, Ar-*H*), 7.58 (d, *J* = 16.03 Hz, 1H, Ar-*H*), 7.51 – 7.49 (m, 2H, Ar-*H*), 7.44 – 7.42 (m, 1H, Ar-H), 7.36 (t, *J* = 7.56 Hz, 1H, Ar-*H*), 7.25 (t, *J* = 7.63 Hz, 1H, Ar-*H*), 7.03 (s, 1H, Ar-*H*), 5.69 (s, 2H, C*H_2_*). ^13^C NMR (176 MHz, DMSO-*d_6_*) (δ, ppm): 194.34, 152.81, 149.83, 143.73, 142.00, 141.54, 140.72, 137.82, 136.73, 135.83, 133.47, 131.71, 129.43, 129.41, 128.61, 128.25, 126.69, 125.99, 125.87, 123.64, 122.40, 121.14, 120.86, 120.69, 119.09, 117.58, 112.39, 111.91, 41.51. ESI-MS (*m/z*) calcd. for C_29_H_19_BrClN_5_O: 568.86, Found: 542.11 [M- N_2_+2]^+^, 543.09 [M-N_2_+2+1]^+^, 545.15 [M-N_2_+2+2+1]^+^. Purity: UP-LC 91.90 %.

##### 6.1.5.6. (E)-1-(3-bromophenyl)-3-(1-((1-(7-chloroquinolin-4-yl)-1H-1,2,3-triazol-4-yl)methyl)- 1H-indol-3-yl)prop-2-en-1-one (7f)

Yellow solid, yield: 69 %, R_f_ = 0.68 (Methanol : DCM = 5: 95), mp: 221.2-221.9 °C, IR (neat): ν (cm^-1^) 3144 (C-H)_str._ triazole ring, 1655 (C=C)_str._ chalcone. ^1^H NMR (700 MHz, DMSO-d_6_) (δ, ppm): 8.91 (s, 1H, NC-*H*_indole_), 8.37 (s, 1H, Ar-H), 8.27 (d, 1H, J = 1.96 Hz, C-*H*_triazole_), 8.22 (s, 1H, Ar-*H*), 8.14 (d, *J* = 7.84 Hz, Ar-*H*), 8.08 (s, 1H, Ar-*H*), 7.97 (d, 1H, *J* = 9.03 Hz, Ar-*H*), 7.83 – 7.81 (m, 3H, Ar-*H*), 7.77 – 7.75 (m, 1H, Ar-*H*), 7.65 (d, *J* = 15.4 Hz, 1H, Ar-*H*), 7.52 (t, *J* = 7.84 Hz, 1H, Ar-*H*), 7.34 (t, *J* = 7.42 Hz, 1H, Ar-*H*), 7.29 (t, *J* = 7.77 Hz, 1H, Ar-*H*), 5.74 (s, 2H, C*H_2_*). ^13^C NMR (176 MHz, DMSO-*d_6_*) (δ, ppm): 187.88, 152.81, 149.84, 143.85, 140.94, 140.74, 139.43, 137.69, 136.31, 135.83, 135.56, 131.42, 131.04, 129.45, 128.62, 127.73, 126.66, 126.40, 125.87, 123.55, 122.70, 122.20, 121.15, 120.71, 117.61, 115.98, 113.03, 111.79, 41.59. ESI-MS (*m/z*) calcd. for C_29_H_19_BrClN_5_O: 568.86, Found: 542.10 [M-N_2_+2]^+^, 543.16 [M-N_2_+2+1]^+^, 545.05 [M-N_2_+2+2+1]^+^. Purity: UP-LC 92.37 %.

##### 6.1.5.7. (E)-1-(4-bromophenyl)-3-(1-((1-(7-chloroquinolin-4-yl)-1H-1,2,3-triazol-4-yl)methyl)- 1H-indol-3-yl)prop-2-en-1-one (7g)

Yellow solid, yield: 62 %, R_f_ = 0.73 (Methanol : DCM = 5: 95), mp: 228.6-229.1 °C, IR (neat): ν (cm^-1^) 3136 (C-H)_str._ triazole ring, 1652 (C=C)_str._ chalcone. ^1^H NMR (700 MHz, DMSO-d_6_) (δ, ppm): 9.12 (d, 1H, *J* = 4.76 Hz, CQ-*H*), 8.93 (s, 1H, NC-*H*_indole_), 8.33 (s, 1H, Ar-*H*), 8.27 (d, *J* = 1.96 Hz, 1H, C-*H*_triazole_), 8.13 (d, 1H, *J* = 7.91 Hz, Ar-*H*), 8.06 (t, *J* = 8.19 Hz, 3H, Ar-*H*), 7.99 – 7.96 (m, 2H, Ar-*H*), 7.82 (t, *J* = 4.48 Hz, 2H, Ar-*H*), 7.76 – 7.73 (m, 3H, Ar-*H*), 7.66 (s, 1H, Ar-*H*), 7.34 (t, *J* = 7.49 Hz, 1H, Ar-*H*), 7.29 – 7.25 (m, 1H, Ar-*H*), 5.73 (s, 2H, C*H_2_*). ^13^C NMR (176 MHz, DMSO-*d_6_*) (δ, ppm): 188.25, 152.81, 149.84, 143.86, 140.73, 139.21, 137.81, 137.72, 136.37, 135.83, 132.24, 130.42, 129.45, 128.62, 126.96, 126.65, 126.31, 125.86, 125.78, 123.55, 122.18, 121.18, 120.71, 117.63, 117.60, 116.02, 113.02, 111.79, 41.55. ESI-MS (*m/z*) calcd. for C_29_H_19_BrClN_5_O: 568.86, Found: 542.16 [M-N_2_+2]^+^, 543.16 [M-N_2_+2+1]^+^, 545.05 [M-N_2_+2+2+1]^+^. Purity: UP-LC 91.31 %.

##### 6.1.5.8. (E)-1-(2-chlorophenyl)-3-(1-((1-(7-chloroquinolin-4-yl)-1H-1,2,3-triazol-4-yl)methyl)- 1H-indol-3-yl)prop-2-en-1-one (7h)

Yellow solid, yield: 80 %, R_f_ = 0.84 (Methanol : DCM = 5 : 95), mp: 207.2-207.6 °C, IR (neat): ν (cm^-1^) 3149 (C-H)_str._ triazole ring, 1650 (C=C)_str._ chalcone. ^1^H NMR (700 MHz, DMSO-*d_6_*) (δ, ppm): 9.12 (d, 1H, J = 4.96 Hz, CQ-*H*), 8.91 (s, 1H, NC-*H*_indole_), 8.26 (d, *J* = 2.1 Hz, 1H, Ar-*H*), 8.23 (s, 1H, C-*H*_triazole_), 7.97(d, 1H_trans indole side_, *J* = 9.03 Hz), 7.95 (m, 1H_trans keto side_ *J* = 7.98 Hz), 7.80 (d, 2H, *J* = 3.71 Hz, Ar-*H*), 7.76 – 7.74 (m, 1H, Ar-*H*), 7.61 (d, *J* = 16.03 Hz, 1H, Ar-*H*), 7.56 (d, *J* = 1.26 Hz, 1H, Ar-*H*), 7.53 – 7.51 (m, 2H, Ar-*H*), 7.48 – 7.46 (m, 2H, Ar-*H*), 7.33 (t, *J* = 7.56 Hz, 1H, Ar-*H*), 7.25 (t, *J* = 7.42 Hz, 1H, Ar-*H*), 7.04 (s, 1H, Ar-*H*), 5.70 (s, 2H, C*H_2_*). ^13^C NMR (176 MHz, DMSO-*d_6_*) (δ, ppm): 193.38, 152.81, 149.83, 143.75, 141.32, 140.72, 139.96, 137.81, 136.74, 135.83, 131.69, 130.41, 130.25, 129.54, 129.44, 128.61, 127.81, 126.68, 126.01, 125.87, 123.64, 122.39, 121.33, 120.86, 120.69, 117.59, 112.39, 111.91, 41.51. ESI-MS (*m/z*) calcd. for C_29_H_19_Cl_2_N_5_O: 524.41, Found: 496.21 [M-N_2_]^+^, 498.22 [M-N_2_+2]^+^, 499.24 [M-N_2_+2+1]^+^. Purity: UP-LC 96.20 %.

##### 6.1.5.9. (E)-1-(3-chlorophenyl)-3-(1-((1-(7-chloroquinolin-4-yl)-1H-1,2,3-triazol-4-yl)methyl)- 1H-indol-3-yl)prop-2-en-1-one (7i)

Yellow solid, yield: 91.4 %, R_f_ = 0.82 (Methanol : DCM = 5 : 95), mp: 216.6-218.1 °C, IR (neat): ν (cm^-1^) 3105 (C-H)_str._ triazole ring, 1652 (C=C)_str._ chalcone. ^1^H NMR (700 MHz, DMSO-*d_6_*) (δ, ppm): 8.37 (s, 1H, Ar-*H*), 8.27 (d, 1H, *J* = 1.75 Hz, C-*H*_triazole_), 8.15 (d, *J* = 7.84 Hz, 1H, Ar-*H*), 8.10-8.08 (m, 2H, CH, Ar-*H*), 8.01 (s, 1H, Ar-*H*), 7.97 (t, *J* = 9.1 Hz, 1H, Ar-*H*), 7.83-7.81 (m, 3H, Ar-*H*), 7.77-7.75 (m, 1H, Ar-*H*), 7.70 (d, *J* = 6.16 Hz, 1H, Ar-*H*), 7.67 (s, 1H, Ar-*H*), 7.59-7.58 (m, 2H, Ar-*H*), 7.33 (d, *J* = 7.77 Hz, 1H, Ar- *H*), 7.30 (d, *J* = 7.77 Hz, 1H, Ar-*H*), 5.74 (s, 2H, C*H_2_*). ^13^C NMR (176 MHz, DMSO-*d_6_*) (δ, ppm): 187.94, 152.82, 149.84, 143.85, 140.74, 139.45, 137.69, 136.35, 135.84, 134.18, 132.67, 131.23, 131.17, 129.46, 128.62, 128.18, 127.36, 126.66, 126.39, 125.87, 123.56, 122.21, 121.18, 120.72, 117.62, 116.00, 113.03, 111.78, 41.58. ESI-MS (*m/z*) calcd. for C_29_H_19_Cl_2_N_5_O: 524.41, Found: 496.21 [M-N_2_]^+^, 498.17[M-N_2_+2]^+^, 499.24 [M-N_2_+2+1]^+^. Purity: UP-LC 95.68 %.

##### 6.1.5.10. (E)-1-(4-chlorophenyl)-3-(1-((1-(7-chloroquinolin-4-yl)-1H-1,2,3-triazol-4-yl)methyl) *-1H-indol-3-yl)prop-2-en-1-one* (7j)

Yellow solid, yield: 52 %, Rf = 0.76 (Methanol : DCM = 5 : 95), mp: 223.2-223.8 °C, IR (neat): ν (cm^-1^) 3153 (C-H)_str._ triazole ring, 1648 (C=C)_str._ chalcone. ^1^H NMR (700 MHz, DMSO-*d_6_*) (δ, ppm): 9.12 (d, *J* = 4.9Hz, 1H, CQ-*H*), 8.93 (s, 1H, NC-*H*_indole_), 8.33 (s, 1H, Ar-*H*), 8.27 (d, *J* = 1.82 Hz, 1H, (C-*H*_triazole_), 8.14 (t, *J* = 8.47 Hz, 3H, Ar-*H*), 8.06 (t, *J* = 3.5 Hz, 1H, Ar-*H*), 7.97 (d, *J* = 9.1 Hz, 1H, Ar-*H*), 7.83 – 7.81 (m, 2H, Ar- *H*), 7.77 – 7.75 (m, 1H, Ar-*H*), 7.66 (s, 1H, Ar-*H*), 7.61 (d, *J* = 8.33 Hz, 2H, Ar-*H*), 7.35 – 7.33 (m, 1H, Ar-*H*), 7.30 – 7.28 (m, 1H, Ar-*H*), 5.73 (s, 2H, C*H_2_*). ^13^C NMR (176 MHz, DMSO-*d_6_*) (δ, ppm): 188.07, 152.83, 149.84, 143.86, 140.74, 139.18, 137.83, 137.73, 137.48, 136.37, 135.87, 135.84, 130.59, 130.28, 129.46, 129.29, 129.25, 128.62, 126.66, 126.31, 125.87, 123.57, 122.18, 121.19, 120.72, 117.65, 117.63, 116.04, 113.02, 111.79, 41.55. ESI-MS (*m/z*) calcd. For C_29_H_19_Cl_2_N_5_O: 524.41, Found: 496.21 [M-N_2_+1]^+^, 498.22 [M-N_2_+2+1]^+^, 499.21 [M-N_2_+2+2]^+^. Purity: UP-LC 96.16 %.

##### 6.1.5.11. (E)-3-(1-((1-(7-chloroquinolin-4-yl)-1H-1,2,3-triazol-4-yl)methyl)-1H-indol-3-yl)-1- (2,4-dichlorophenyl)prop-2-en-1-one (7k)

Yellow solid, yield: 64 %, R_f_ = 0.77 (Methanol : DCM = 5 : 95), mp: 181.1-182.4 °C, IR (neat): ν (cm^-1^) 3153 (C-H)_str._ triazole ring, 1652 1648 (C=C)_str._ chalcone. ^1^H NMR (700 MHz, DMSO-*d_6_*) (δ, ppm): 9.12 (d, 1H, *J* = 7.69 Hz, CQ-*H*), 8.91 (s, 1H, NC-*H*_indole_), 8.25 (t, 2H, J = 1.82 Hz, Ar-H, (C-*H*_triazole_), 7.96 (t, 2H, *J* = 6.44 Hz, Ar- *H*), 7.81 (d, 2H, *J* = 5.04 Hz, Ar-*H*), 7.75 (s, 2H, Ar-*H*), 7.63 (d, *J* = 15.96 Hz, 1H, Ar-*H*), 7.56 (s, 2H, Ar-H), 7.33 (t, *J* = 7.56 Hz, 1H, Ar-*H*), 7.25 (t, *J* = 7.56 Hz, 1H, Ar-*H*), 7.03 (d, *J* = 16.03 Hz, 1H, Ar-*H*), 5.70 (s, 2H, C*H_2_*). ^13^C NMR (176 MHz, DMSO-*d_6_*) (δ, ppm): 192.47, 152.81, 149.84, 143.70, 141.98, 140.72, 138.77, 137.84, 137.01, 135.83, 135.38, 131.54, 130.95, 129.96, 129.44, 128.62, 128.05, 126.69, 125.99, 125.86, 123.68, 122.44, 121.02, 120.96, 120.69, 117.59, 112.46, 111.93, 41.52. ESI-MS (*m/z*) calcd. for C_29_H_18_Cl_3_N_5_O: 558.85, Found: 532.19 [M-N_2_+2]^+^, 534.17 [M-N_2_+2+2]^+^, 535.18 [M-N_2_+2+2+1]^+^. Purity: UP-LC 95.71 %.

##### 6.1.5.12. (E)-3-(1-((1-(7-chloroquinolin-4-yl)-1H-1,2,3-triazol-4-yl)methyl)-1H-indol-3-yl)-1- (2-methoxyphenyl)prop-2-en-1-one (7l)

Cream solid, yield: 53 %, R_f_ = 0.80 (Methanol : DCM = 5 : 95), mp: 176.4-176.8 °C, IR (neat): ν (cm^-1^) 3112 (C-H)_str._ triazole ring, 1663 (C=C)_str._ chalcone.^1^H NMR (700 MHz, DMSO-d_6_) (δ, ppm): 9.96 (s, 1H, CQ-*H*), 9.93 (s, 1H, NC-*H*_indole_), 9.12 (d, *J* = 4.69 Hz, 1H, Ar-*H*), 8.94 (s, 1H, (C-*H*_triazole_), 8.50 (s, 1H, Ar-*H*), 8.36 (s, 1H, Ar-*H*), 8.27 (d, *J* = 1.82 Hz, 1H, Ar-*H*), 8.13 – 8.11 (m, 2H, Ar- *H*), 7.96 (d, *J* = 9.03 Hz, 1H, Ar-*H*), 7.82 (t, *J* = 4.55 Hz, 2H, Ar-*H*), 7.77 – 7.75 (m, 1H, Ar-*H*), 7.65 (d, *J* = 8.26 Hz, 1H, Ar-*H*), 7.36 – 7.33 (m, 2H, Ar-*H*), 7.03 – 7.27 (m, 2H, Ar-*H*), 5.27 (s, 2H, C*H_2_*), 5.24 (s, 2H, OC*H_3_*). ^13^C NMR (176 MHz, DMSO-*d_6_*) (δ, ppm): 185.39, 185.31, 152.82, 149.89, 143.63, 141.29, 140.72, 140.62, 137.40, 137.06, 135.84, 129.46, 128.63, 126.76, 125.83, 125.23, 125.17, 124.21, 124.19, 123.27, 123.16, 121.61, 121.56, 120.71, 118.03, 117.61, 111.88, 111.69, 78.48, 77.31, 41.78. ESI-MS (*m/z*) calcd. for C_30_H_22_ClN_5_O_2_: 519.99; Found: 518.95 [M-H]^-^. Purity: UP-LC 96.20 %.

##### 6.1.5.13. (E)-3-(1-((1-(7-chloroquinolin-4-yl)-1H-1,2,3-triazol-4-yl)methyl)-1H-indol-3-yl)-1- (3-methoxyphenyl)prop-2-en-1-one (7m)

Yellow solid, yield: 76 %, R_f_ = 0.79 (Methanol : DCM = 5 : 95), mp: 198.4 – 198.8 °C, IR (neat): ν (cm^-1^) 3151 (C-H)_str._ triazole ring, 1642 (C=C)_str._ chalcone. ^1^H NMR (700 MHz, DMSO-*d_6_*) (δ, ppm): 9.13 (d, 1H, *J* = 4.9 Hz, CQ-*H*), 8.93 (s, 1H, NC-*H*_indole_), 8.34 (s, 1H, Ar-*H*), 8.27 (s, 1H, (C-*H*_triazole_), 8.10 (d, *J* = 7.84 Hz, 1H, C*H*), 8.05 (s, 1H, Ar-H), 7.98 (d, *J* = 9.03 Hz, 1H, Ar-*H*), 7.83 – 7.81 (m, 3H, Ar-*H*), 7.77 (d, *J* = 9.03 Hz, 1H, Ar-*H*), 7.73 (d, *J* = 7.91 Hz, 1H, Ar-*H*), 7.65 (t, *J* = 10.92 Hz, 1H, Ar-*H*), 7.54 (t, *J* = 5.25 Hz, 2H, Ar-*H*), 7.49 – 7.46 (m, 1H, Ar-*H*), 7.33 (t, 1H, *J* = 7.28 Hz, Ar-*H*), 7.30 (t, *J* = 8.47 Hz, 1H, Ar-*H*), 7.21 – 7.20 (m, 1H, Ar-*H*), 5.73 (s, 2H, C*H_2_*). ^13^C NMR (176 MHz, DMSO- *d_6_*) (δ, ppm): 189.02, 159.97, 152.89, 149.84, 143.91, 140.75, 140.39, 138.65, 137.67, 135.94, 135.84, 130.35, 130.33, 129.46, 128.62, 126.65, 126.39, 125.88, 123.48, 122.15, 121.14, 121.05, 120.79, 120.72, 118.89, 117.65, 116.63, 113.21, 112.97, 111.77, 55.78, 41.54. ESI-MS (*m/z*) calcd. for C_30_H_22_ClN_5_O_2_: 519.99, Found: 492.23 [M-N_2_+1]^+^, 494.24 [M-N_2_+2+1]^+^, 495.19 [M- N_2_+2+2]^+^. Purity: UP-LC 92.14 %.

##### 6.1.5.14. (E)-3-(1-((1-(7-chloroquinolin-4-yl)-1H-1,2,3-triazol-4-yl)methyl)-1H-indol-3-yl)-1- (4-methoxyphenyl)prop-2-en-1-one (7n)

Yellow solid, yield: 63%, R_f_ = 0.90 (Methanol : DCM = 5 : 95), mp: 204.2-204.7 °C, IR (neat): ν (cm^-1^) 3155 (C-H)_str._ triazole ring, 1646 (C=C)_str._ chalcone. ^1^H NMR (700 MHz, DMSO-d_6_) (δ, ppm): 9.12 (d, *J* = 4.69Hz, CQ-*H*), 8.93 (s, 1H, NC-*H*_indole_), 8.29 (s, 1H, Ar-*H*), 8.27 (d, *J* = 2.1Hz, 1H, (C-*H*_triazole_), 8.14 – 8.11 (m, 3H, CH, Ar- *H*), 8.01 – 7.97 (m, 2H, CH, Ar-*H*), 7.83 – 7.80 (m, 2H, Ar-*H*), 7.77 – 7.75 (m, 1H, Ar-*H*), 7.70 (s, 1H, Ar-*H*), 7.33 (t, *J* = 7.42 Hz, 1H, Ar-*H*), 7.28 (t, *J* = 7.63 Hz, 1H, Ar-*H*), 7.08 (d, *J* = 8.82 Hz, 2H, Ar-*H*), 5.73 (s, 2H, C*H_2_*), 3.86 (s, 3H, OC*H_3_*). ^13^C NMR (176 MHz, DMSO-*d_6_*) (δ, ppm): 184.22, 160.56, 152.29, 148.57, 143.11, 140.35, 140.47, 138.63, 137.27, 136.23, 135.35, 130.67, 130.23, 128.74, 128.55, 126.35, 126.19, 125.73, 123.18, 122.13, 121.13, 121.65, 120.19, 118.32, 118.19, 117.64, 116.61, 113.71, 112.47, 110.57, 55.38, 41.64. ESI-MS (m/z) calcd. For C_30_H_22_ClN_5_O_2_: 519.99, Found: 492.27 [M-N_2_+1]^+^, 494.24 [M-N_2_+2+1]^+^, Purity: UP-LC 94.99 %.

##### 6.1.5.15. (E)-3-(1-((1-(7-chloroquinolin-4-yl)-1H-1,2,3-triazol-4-yl)methyl)-1H-indol-3-yl)-1- (3,4-dimethoxyphenyl)prop-2-en-1-one (7o)

Yellow solid, yield: 60 %, R_f_ = 0.73 (Methanol : DCM = 5 : 95), mp: 169.8-171.5 °C, IR (neat): ν (cm^-1^) 3106 (C-H)_str._ triazole ring, 1644 (C=C)_str._ chalcone. ^1^H NMR (700 MHz, DMSO-*d_6_*) (δ, ppm): 9.12 (d, *J* = 4.62Hz, 1H, CQ-*H*), 8.93 (s, 1H, NC-*H*_indole_), 8.30 (s, 1H, Ar-*H*), 8.27 (s, 1H, (C-*H*_triazole_),), 8.10 (d, *J* = 7.84 Hz, 1H, C*H*), 7.99 (d, *J* = 9.73 Hz, 2H, Ar-*H*), 7.86 (d, *J* = 8.47 Hz, 1H, CH), 7.82 (t, *J* = 5.81 Hz, 2H, Ar-*H*), 7.76 (d, *J* = 9.03 Hz, 1H, Ar-*H*), 7.69 (d, *J* = 15.47 Hz, 1H, Ar-*H*), 7.59 (s, 1H, Ar-*H*), 7.33 (t, *J* = 7.35 Hz, 1H, Ar-*H*), 7.28 (t, *J* = 7.49 Hz, 1H, Ar-*H*), 7.09 (d, *J* = 8.4 Hz, 1H, Ar-*H*), 5.73 (s, 2H, C*H_2_*), 3.86 (s, 3H, OC*H_3_*), 3.86 (s, 3H, OC*H_3_*). ^13^C NMR (176 MHz, DMSO-*d_6_*) (δ, ppm): 187.63, 153.17, 152.82, 149.84, 149.22, 143.96, 140.74, 137.62, 137.52, 135.83, 135.37, 131.68, 129.44, 128.62, 126.62, 126.43, 125.86, 123.38, 123.17, 122.01, 120.99, 120.71, 117.59, 116.55, 113.00, 111.71, 111.31, 111.06, 65.39, 56.19, 41.51. ESI-MS (*m/z*) calcd. For C_31_H_24_ClN_5_O_3_: 550.02, Found: 522.28 [M-N_2_]^+^, 524 [M-N_2_+2]^+^. Purity: UP-LC 95.56 %.

##### 6.1.5.16. (E)-3-(1-((1-(7-chloroquinolin-4-yl)-1H-1,2,3-triazol-4-yl)methyl)-1H-indol-3-yl)-1- (thiophen-2-yl)prop-2-en-1-one (7p)

Yellow solid, yield: 52 %, R_f_ = 0.74 (Methanol : DCM = 5 : 95), mp: 218.6-219.1 °C, IR (neat): ν (cm^-1^) 3147 (C-H)_str._ triazole ring, 1631 (C=C)_str._ chalcone. ^1^H NMR (700 MHz, DMSO-*d_6_*) (δ, ppm): 9.12 (d, *J* = 4.69 Hz, 1H, CQ-*H*), 8.93 (s, 1H, NC-*H*_indole_), 8.31 (s, 1H, Ar-*H*), 8.26 – 8.25 (m, 2H, Ar-H, (C-*H*_triazole_), 8.16 (d, *J* = 7.84 Hz, 1H, Ar-*H*), 8.01 (d, *J* = 15.33 Hz, 1H, Ar-*H*), 7.98 (t, *J* = 6.09 Hz, 2H, Ar-*H*), 7.82 (t, *J* = 4.13 Hz, 2H, Ar-*H*), 7.76 – 7.75 (m, 1H, Ar-*H*), 7.61 (s, 1H, Ar-*H*), 7.34 (t, *J* = 7.35 Hz, 1H, Ar-*H*), 7.30 – 7.28 (m, 2H, Ar-*H*), 5.73 (s, 2H, C*H_2_*). ^13^C NMR (176 MHz, DMSO-*d_6_*) (δ, ppm): 181.86, 152.81, 149.84, 146.63, 143.90, 140.73, 137.76, 137.69, 136.12, 135.83, 134.77, 132.74, 129.45, 129.27, 128.62, 126.64, 126.30, 125.87, 123.50, 122.18, 121.23, 120.71, 117.60, 116.26, 112.82, 111.75,41.54. ESI-MS (*m/z*) calcd. for C_27_H_18_ClN_5_OS: 495.98; Found: 496.13 [M+1]^+^, 498.07 [M+2+1]^+^, 499.38 [M+2+2]^+^. Purity: UP-LC 95.58 %.

##### 6.1.5.17. (E)-1-(3,5-bis(benzyloxy)phenyl)-3-(1-((1-(7-chloroquinolin-4-yl)-1H-1,2,3-triazol-4- yl)methyl)-1H-indol-3-yl)prop-2-en-1-one (7q)

Off white solid, yield: 71%, R_f_ = 0.81 (Methanol : dichloromethane = 5 : 95), mp: 209.8-210.1 °C, IR (neat): ν (cm^-1^) 3105 (C-H)_str._ triazole ring, 1651 (C=C)_str._ chalcone. ^1^H NMR (700 MHz, DMSO-*d_6_*) (δ, ppm): 9.11 (d, 1H, *J* = 4.62 Hz, CQ-*H*), 8.93 (s, 1H, NC-*H*_indole_), 8.35 (s, 1H, Ar-*H*), 8.27 (s, 1H, (C-*H*_triazole_), 8.04 (d, *J* = 8.89 Hz, 2H, Ar-*H*), 7.97 (d, *J* = 9.1 Hz, 1H, Ar-*H*), 7.81 (t, *J* = 3.78 Hz, 2H, Ar-*H*), 7.76 – 7.74 (m, 1H, Ar-H), 7.57 (d, *J* = 15.47 Hz, 1H, Ar-*H*), 7.48 (d, *J* = 7.77 Hz, 4H, Ar-*H*), 7.40 (t, *J* = 7.56 Hz, 4H, Ar-*H*), 7.29 (s, 3H, Ar-*H*), 7.34 (t, *J* = 7.28 Hz, 3H, Ar-*H*), 6.95 (s, 1H, Ar-*H*), 5.73 (s, 2H, C*H_2_*), 5.20 (s, 4H, C*H_2_*). ^13^C NMR (176 MHz, DMSO-*d_6_*) (δ, ppm): 188.70, 160.12, 152.80, 149.84, 143.89, 141.03, 140.73, 138.69, 137.62, 137.31, 135.83, 135.77, 129.44, 128.95, 128.62, 128.39, 128.26, 126.64, 126.46, 125.87, 123.46, 122.13, 120.95, 120.71, 117.60, 116.51, 112.97, 111.75, 107.49, 106.40, 70.05, 41.56. ESI-MS (*m/z*) calcd. for C_43_H_32_ClN_5_O_3_:702.20; Found:702.32 [M]^+^, 703.32 [M+1]^+^, 705.36 [M+2+1]^+^. Purity: UP-LC 94.99 %.

##### 6.1.5.18. (E)-3-(1-((1-(7-chloroquinolin-4-yl)-1H-1,2,3-triazol-4-yl)methyl)-1H-indol-3-yl)-1-o- tolylprop-2-en-1-one (7r)

Yellow solid, yield: 40 %, R_f_ = 0.71 (Methanol : DCM = 5 : 95), mp: 178.4-179.2 °C, IR (neat): ν (cm^-1^) 3144 (C-H)_str._ triazole ring, 1646 (C=C)_str._ chalcone.^1^H NMR (700 MHz, DMSO-*d_6_*) (δ, ppm): 9.11 (d,1H, *J* = 4.62 Hz, CQ-*H*), 8.91 (s, 1H, NC-*H*_indole_), 8.26 (s, 1H, Ar-*H*), 8.21 (s, 1H, (C-*H*_triazole_), 7.95 (t, J = 8.47 Hz, 2H, Ar-H), 7.80 (t, *J* = 3.78 Hz, 2H, Ar-*H*), 7.75 (d, *J* = 7.28 Hz, 1H, Ar-*H*), 7.67 (d, *J* = 15.89 Hz, 1H, Ar-*H*), 7.52 (d, J = 7.63 Hz, 1H, Ar-*H*), 7.40 (t, *J* =7.42 Hz, 1H, Ar-*H*), 7.31 (d, *J* = 14.21 Hz, 3H, Ar-*H*), 7.25 (t, *J* = 7.7 Hz, 1H, Ar-*H*), 7.13 (d, *J* = 15.96 Hz, 1H, Ar-*H*), 5.70 (s, 2H, C*H_2_*). ^13^C NMR (176 MHz, DMSO-*d_6_*) (δ, ppm): 195.82, 152.81, 149.84, 143.85, 140.72, 140.29, 139.96, 137.74, 136.14, 136.0, 135.83, 131.42, 130.37, 129.43, 128.61, 128.12, 126.64, 126.13, 125.86, 123.51, 122.21, 121.57, 120.85, 120.69, 117.58, 112.46, 111.82, 41.47. ESI-MS (*m/z*) calcd. for C_30_H_22_ClN_5_O: 503.99, Found: 504.14 [M+1]^+^, 505.16 [M+2]^+^. Purity: UP-LC 95.56 %.

##### 6.1.5.19. (E)-3-(1-((1-(7-chloroquinolin-4-yl)-1H-1,2,3-triazol-4-yl)methyl)-1H-indol-3-yl)-1- (4-fluorophenyl)prop-2-en-1-one (7s)

Yellow solid, yield: 80 %, R_f_ = 0.69 (Methanol : DCM = 5 : 95), mp: 234.5-234.8 °C, IR (neat): ν (cm^-1^) 3151 (C-H)_str._ triazole ring, 1648 (C=C)_str._ chalcone. ^1^H NMR (700 MHz, DMSO-*d_6_*) (δ, ppm): 9.12 (t, 1H, *J* = 4.97 Hz, CQ-*H*), 8.93 (s, 1H, NC-*H*_indole_), 8.32 (s, 1H, Ar-*H*), 8.28 (d, *J* = 1.96 Hz, 1H, (C-*H*_triazole_), 8.23 -8.21 (m, 2H, Ar- *H*), 8.14 (t, *J* = 8.12 Hz, 2H, Ar-*H*), 8.04 (d, *J* = 15.47 Hz, 1H, Ar-*H*), 7.98 -7.95 (m, 1H, Ar- *H*), 7.82 (t, *J* = 4.83 Hz, 3H, Ar-*H*), 7.77 – 7.76 (m, 1H, Ar-*H*), 7.69 (s, 1H, Ar-*H*), 7.39 – 7.34 (m, 3H, Ar-*H*), 7.30 -7.23 (m, 1H, Ar-*H*), 5.73 (s, 2H, C*H_2_*). ^13^C NMR (176 MHz, DMSO-*d_6_*) (δ, ppm): 187.76, 165.87, 152.83, 149.84, 144.03, 143.89, 138.84, 137.71, 136.18, 135.81, 131.59, 129.47, 128.62, 126.31, 125.79, 123.51, 122.13, 121.19, 120.72, 117.63, 116.06, 112.99, 111.67, 55.39, 41.53. ESI-MS (*m/z*) calcd. for C_29_H_19_ClFN_5_O: 507.95, Found: 780.23 [M-N_2_+1]^+^, 482.22 [M-N_2_+2+1]^+^,483.22 [M-N_2_+2+2]^+^, Purity: UP-LC 95.04 %.

##### 6.1.5.20. 1-((1-(7-chloroquinolin-4-yl)-1H-1,2,3-triazol-4-yl)methyl)-1H-indole-3- carbaldehyde (9)

Cream solid, yield: 59 %, R_f_ = 0.71 (Methanol : dichloromethane = 5 : 95), mp: 209.1–209.9 °C, IR (neat): ν (cm^-1^) 3112 (C-H)_str._ triazole ring. ^1^H NMR (700 MHz, DMSO- *d_6_*) (δ, ppm): 9.96 (s, 1H, C-*H*_CHO_), 9.12 (d, 1H, *J* = 4.69 Hz, CQ-*H*), 8.94 (s, 1H, Ar-*H*), 8.49 (s, 1H, NC-*H*_indole_), 8.26 (d, 1H, *J* = 2.03 Hz, (C-*H*_triazole_), 8.12 (d, *J* = 7.84 Hz, 1H, Ar-*H*), 7.96 (d, *J* = 9.1 Hz, 1H, Ar-*H*), 7.82 (t, *J* = 4.48 Hz, 2H, Ar-*H*), 7.76-7.75 (m, 1H, Ar-*H*), 7.33 (d, *J* = 7.35 Hz, 1H, Ar-*H*), 7.28 (d, *J* = 7.7 Hz, 1H, Ar-*H*), 5.79 (s, 2H, C*H_2_*). ^13^C NMR (176 MHz, DMSO-*d_6_*) (δ, ppm): 185.30, 152.82, 149.84, 143.63, 137.39, 135.84, 129.46, 128.62, 126.76, 125.83, 125.23, 124.18, 123.16, 121.55, 120.69, 118.03, 117.61, 111.88, 41.78. ESI-MS (*m/z*) calcd. for C_21_H_14_ClN_5_O: 387.83, Found: 360.07 [M-N_2_+1]^+^, 362.06 [M-N_2_+2+1]^+^. Purity: UP- LC 95.68 %.

### 6.2. Biology

#### 6.2. Antimalarial Activity

*P. falciparum* 3D7 strain was used in this study, cultured in human O^+^ RBCs supplemented with complete RPMI 1640 (Invitrogen, USA), 0.5% AlbuMax I (Invitrogen, USA), 0.2% NaHCO_3_ (Sigma, USA), 25 mg ml^-1^ gentamicin (Invitrogen, USA), 0.1 mM hypoxanthine (Invitrogen, USA) at 37 °C prepared according to method described previously [48]. Parasite culture was maintained in a mixed gas environment (5% O_2_, 5% CO_2_, and 90% N_2_). Parasites were routinely monitored by Giemsa staining of thin blood smears. Parasites were synchronized at the ring stage by 5% D-sorbitol.

#### 6.2.1. Growth inhibition assay

DNA-specific dye SYBR green I (Invitrogen, Carlsbad, USA) was used to perform a growth inhibition assay to calculate the inhibitory effect of CQTrICh-analogs against parasite *P. falciparum* 3D7 [49,50]. Out of 20 compounds, **7l** and **7r** were selected for further study. Initially, parasites (3D7 and RKL-9 strains of *P. falciparum*) at the trophozoite stage with 1% parasitemia and 2% hematocrit incubated with different concentrations (0.62-20µM) of **7l** and **7r** for one complete intra-erythrocytic cycle; parasites without treatment were used as a negative control. Giemsa staining was done to observe the parasite morphologically. SYBR green dye assay was used to find out the percent growth inhibition. The fluorescence was taken with the 485 nm excitation and 530 nm emission. The experiment was done in triplicate. Data were expressed as mean ± SD. Growth inhibition (% Inhibition) was calculated as follows: % Inhibition = [1- % Parasitemia (Treatment) / % Parasitemia (Control)] × 100.

#### 6.2.2. Blood stage specific effect

Blood stage-specific effect of **7l** and **7r** on CQ*^S^*-3D7 *P. falciparum* was performed, as described previously [46]. Briefly, synchronized ring stage (10-12 hrs) parasites of 0.8% parasitemia and 2% hematocrit were treated with **7l** and **7r** at IC_50_ concentration in a complete RPMI-1640 medium and seeded in 96-well flat-bottom microtiter plates respectively. Parasite culture was incubated and monitored up to 56 hrs post-treatment at 37 °C. Morphological analysis was performed by counting Giemsa-stained slides in duplicate with the help of a light microscope (Olympus).

#### 6.2.3. Hemozoin Assay

Synchronized ring-stage parasites (6% parasitemia) were treated with different concentrations of compounds **7l** and **7r** (1.5, 2.5, 5, and 50 *µ*M). Samples were seeded into 96-well flat bottom plates in triplicate. After 30 hrs, samples were treated with 2.5% SDS in 0.1 M sodium bicarbonate pH 8.8; incubated at room temperature for 30 min. Centrifugation was done and resuspended in 5% SDS and 50 mM NaOH. The suspension was incubated for 30 min. Monomeric heme was quantified at 405/750 nm as described previously [51].

#### 6.2.4. Hemolytic activity

The hemolytic assay was performed as described previously [52]. In brief, RBCs suspension with 10% (v/v) washed with 1 ×PBS (pH 7.4) and re-suspended in 1 × PBS. The suspension was incubated with **7l** and **7r** compounds at different concentrations for 1 h at 37 °C. Samples were centrifuged and supernatant was taken for absorbance at 415 nm. Triton X-100 with 1% (v/v) was used as a positive control. The experiment was done in triplicate.

#### 6.2.5. Cytotoxicity assay

Cytotoxicity of compounds toward HEK293 and HEPG2 cells was measured by the colorimetric MTT assay as previously described [53,54]. A total of 2×10^4^ (HEK293) and 10^4^ (HEPG2) cells supplemented with DMEM and 10% FBS were seeded in 96-well microtitration plate. The plate was incubated overnight at 37 °C in a humidified, 5% CO_2_ incubator. Compounds **7l** and **7r** were added at conc. of 2.5-600 μM (HEK293) & 2-500 μM (HEPG2), and untreated cells were used as control. The plate was incubated for 24 h at 37 °C and 10 μL of MTT (5 mg/mL in 1X-PBS) was added to each well. After incubation formazan crystals were dissolved with DMSO. Absorbance was taken at 570 nm using a microplate reader (Thermo scientific). Each assay was carried out in triplicate.

#### 6.2.6. Molecular docking-studies

To ascertain the compound’s affinity for CDPK1, molecular docking was used. Docking was performed through InstaDock software employing the AutoDock vina-based software [46]. With a blind search space of 62, 58, and 60, centred at -1.465, -20.489, and -19.967 for the X, Y, and Z axes, respectively, the docking was performed using the default docking parameters generated by InstaDock. Using the “ligand splitter” programme of the InstaDock software, the docking output was generated to all feasible conformations of the ligand. The generated output was examined based on the docking score, which established the ligand’s affinity for binding to CDPK1. Docking poses were visualized by PyMOL and Discovery Studio Visualizer. ADME properties were calculated from SwissADME [55], and its carcinogenicity is predicted by using CarcinoPred-EL [56].

#### 6.2.7. Microscale Thermophoresis (MST) for interaction analysis

Binding affinities of the compounds: **7l** & **7r** with the recombinant 6xHis-CDPK1 protein (r*Pf*CDPK1) were evaluated by MST analyses, using Monolith NT.115 instrument (NanoTemper Technologies, Munich, Germany). MST relies on binding-induced changes in thermophoretic mobility, which depends on several molecular properties including particle charge, size, conformation, hydration state, and solvation entropy. Thus, under constant buffer conditions, thermophoresis of unbound proteins typically differs from the thermophoresis of proteins bound to their interaction partners. The thermophoretic movement of a fluorescently labeled protein is measured by monitoring the fluorescence distribution. The protocol for r*Pf*CDPK1/(**7l** or **7r**) interaction analysis was followed as described previously [57,58]. Briefly, 10 μM r*Pf*CDPK1 was prepared in HEPES-NaCl Buffer, pH 7.5 supplemented with 0.05% Tween-20 to prevent sample aggregation, followed by labeling with 30 μM Cysteine reactive dye (Monolith Protein Labelling Kit Red-Maleimide 2^nd^ Generation, NanoTemper), and incubated in dark at Room Temperature (RT) for 30 minutes. A column (provided by the supplier) was prepared by washing and equilibrating with the buffer. The labeled r*Pf*CDPK1 along with the buffer was added to the column, followed by collecting elution fractions. The elution fraction with fluorescence counts less than 1,000 was taken further for interaction analysis. Increasing concentrations of **7l** and **7r** (4.8 nM to 80 μM and 2.4 nM to 40 μM, respectively) diluted in the buffer supplemented with 0.05% tween-20 were titrated against the constant concentration of the labeled r*Pf*CDPK1. Samples were pre-mixed and incubated for 10 minutes at RT in the dark, followed by loading of the samples into the standard treated capillaries (K002 Monolith NT.115). All experiments were carried out at RT, at 20% LED power and 40% MST power. Data evaluation was performed with the Monolith software (Nano Temper, Munich, Germany)

#### 6.2.8. In vitro kinase assay with rPfCDPK1 and HEPG2 lysate

Functional activity of r*Pf*CDPK1 was monitored by utilizing ADP-Glo™ Kinase Assay (Promega Corporation), which is an ATP regeneration-based luciferase reaction system resulting from nascent ADP phosphorylation. The luminescence signal, thus generated, is proportional to the amount of ADP released in a given kinase reaction. Phosphorylation experiments were performed with 100 ng of 6xHis-CDPK1, per reaction, in assay buffer (100 mM Tris-Cl, pH 7.4; 2.5 mM DTT; 50 mM MgCl_2_ and 2.5 mM MnCl_2_), by following previously described protocols [41,59]. The enzymatic reaction was carried out in the absence and presence of calcium ions. 10 μg of dephosphorylated casein from bovine milk, per reaction, was used as an exogenous substrate for the enzyme. 2.5 mM EGTA was added for conditions requiring the absence of Ca^2+^ ions. Kinase reactions were initiated by adding 1 μM ATP and allowed to take place at 30°C for 1 h. To test for any inhibitory effect of the compounds, **7l** and **7r**, CDPK1 was allowed to pre-incubate in the reaction buffer with 1 and 5 μM of each compound at RT for 20 minutes, followed by initiating the reactions by adding ATP. Upon completion of the reactions, ADP- Glo™ Kinase Assay was performed. Luminescence signal, thus generated, was measured with Lumat³ LB 9508 Ultra-Sensitive Tube Luminometer (Berthold Technologies, U.S.A.).

A similar assay was performed to assess the functional activity of the mammalian kinome, as described previously [60]. Briefly, HepG2 cells (10^7^) were re-suspended in 1mL of lysis buffer (20mM Tris, pH 7.7, 0.5% (v/v) Nonidet P-40, 200mM NaCl, 50mM NaF, 0.2mM sodium orthovanadate, 1× PIC and 0.1% (v/v) 2-mercaptoethanol) for 1 h at 4°C, centrifuged at 13,000 rpm for 10 min and supernatant was collected. Kinase activity was assessed in the presence of **7l** and **7r,** at 1 and 5 μM concentrations.

#### 6.2.9 Statistical analysis

The data were calculated as one-way analysis of variance (ANOVA) to observe the mean values obtained for control and after treatment with analogs. Dunnett’s test was used to compare the treatment and control and statistical significance was set at P ≤ 0.01.

## ANCILLARY INFORMATION

Contains spectral data of compounds (**S.3-S.94**), Possible fragmentation pattern in the mass spectrometry (**S.95**), growth Inhibition assay of **7a-s** and **9** compounds (**S.96**), and physicochemical properties of **7a-s** and **9** compounds (**Table S1**).

## AUTHOR INFORMATION

### Corresponding authors

Shailja Singh, Special Center for Molecular Medicine, Jawaharlal Nehru University, Delhi: 110067, India; email address

Mohammad Abid, Medicinal Chemistry Laboratory, Department of Biosciences, Jamia Millia Islamia, New Delhi, 110025, India; email address

## AUTHOR CONTRIBUTIONS

M.A. and S.S. designed the project and optimized the experiments, interpreted the data, and wrote the manuscript; I.I., P.H., and M.C.J. performed the synthesis and characterized the compounds; A.H. and M.F.A. did NMR and interpretation of spectral data. A.U. and S.G. performed all the biological experiments and in silico study; R.J. and A.G. did biochemical and biophysical assays with purified protein, and J.V.N. helped in drafting the manuscript. M.A. and S.S. provided funding acquisition and aided administrative processing. All the authors approved and reviewed the final version of the manuscript.

## ACKNOWLEDGMENTS

MA and SS gratefully acknowledge the financial support in the form of a Core Research Grant from Science & Engineering Research Board (SERB), Govt. of India (Project No. CRG/2018/003967). SS is a recipient of the National Bioscientist award. SS is acknowledging the Drug and Pharmaceuticals Research Programme (DPRP) (Project No. P/569/2016-1/TDT) and DST-SERB (Project No. CRG/2019/002231). Iram Irfan acknowledges DST for postdoctoral fellowship under the women scientist scheme (WOS-A) (File no: SR/WOS-A/CS-116/2018). AU is a recipient of a Senior Research Fellowship from ICMR (Fellowship/48/2019-ECD-II). RJ acknowledges University Grant Commission (UGC) for doctoral research fellowship (Sr. no: 2061430573, Ref. no.: 22/06/2014(i)EU-V). We acknowledge the National Institute of Malaria Research for providing the RKL-9 chloroquine-resistant line of *P. falciparum*. The funders had no role in study design, data collection, and analysis, decision to publish, or preparation of the manuscript.

## NOTES

The authors declare no competing financial interest.

## REFERENCES

[1] H. Hussain, S. Specht, S.R. Sarite, A. Hoerauf, K. Krohn, New quinoline-5,8-dione and hydroxynaphthoquinone derivatives inhibit a chloroquine resistant Plasmodium falciparum strain, Eur. J. Med. Chem. 54 (2012). https://doi.org/10.1016/j.ejmech.2012.06.046.

[2] S. Kumar, M. Guha, V. Choubey, P. Maity, U. Bandyopadhyay, Antimalarial drugs inhibiting hemozoin (β-hematin) formation: A mechanistic update, Life Sci. 80 (2007) 813–828. https://doi.org/10.1016/j.lfs.2006.11.008.

[3] K.K. Roy, Targeting the active sites of malarial proteases for antimalarial drug discovery: approaches, progress and challenges, Int. J. Antimicrob. Agents. 50 (2017) 287–302. https://doi.org/10.1016/j.ijantimicag.2017.04.006.

[4] B. Li, T.J. Webster, Bacteria antibiotic resistance: New challenges and opportunities for implant-associated orthopedic infections, J. Orthop. Res. 36 (2018) 22–32. https://doi.org/10.1002/jor.23656.

[5] A. Uddin, M. Chawla, I. Irfan, S. Mahajan, S. Singh, M. Abid, Medicinal chemistry updates on quinoline- And endoperoxide-based hybrids with potent antimalarial activity, RSC Med. Chem. 12 (2021). https://doi.org/10.1039/d0md00244e.

[6] WHO World Malaria Report, World Malaria Report, 2021. https://www.who.int/teams/global-malaria-programme/reports/world-malaria-report-2021.

[7] T. Hänscheid, T.J. Egan, M.P. Grobusch, Haemozoin: from melatonin pigment to drug target, diagnostic tool, and immune modulator, Lancet Infect. Dis. 7 (2007). https://doi.org/10.1016/S1473-3099(07)70238-4.

[8] T.J. Egan, Recent advances in understanding the mechanism of hemozoin (malaria pigment) formation, J. Inorg. Biochem. 102 (2008). https://doi.org/10.1016/j.jinorgbio.2007.12.004.

[9] V.K. Zishiri, M.C. Joshi, R. Hunter, K. Chibale, P.J. Smith, R.L. Summers, R.E. Martin, T.J. Egan, Quinoline antimalarials containing a dibemethin group are active against chloroquinone-resistant plasmodium falciparum and inhibit chloroquine transport via the P. falciparum chloroquine-resistance transporter (PfCRT), J. Med. Chem. 54 (2011). https://doi.org/10.1021/jm2009698.

[10] S. Andrews, S.J. Burgess, D. Skaalrud, J.X. Kelly, D.H. Peyton, Reversal agent and linker variants of reversed chloroquines: Activities against Plasmodium falciparum, J. Med. Chem. 53 (2010). https://doi.org/10.1021/jm900972u.

[11] S.J. Burgess, A. Selzer, J.X. Kelly, M.J. Smilkstein, M.K. Riscoe, D.H. Peyton, A chloroquine-like molecule designed to reverse resistance in Plasmodium falciparum, J. Med. Chem. 49 (2006). https://doi.org/10.1021/jm060399n.

[12] S.J. Burgess, J.X. Kelly, S. Shomloo, S. Wittlin, R. Brun, K. Liebmann, D.H. Peyton, Synthesis, Structure-activity relationship, and mode-of-action studies of antimalarial reversed chloroquine compounds, J. Med. Chem. 53 (2010). https://doi.org/10.1021/jm1006484.

[13] M.C. Joshi, T.J. Egan, Quinoline Containing Side-chain Antimalarial Analogs: Recent Advances and Therapeutic Application, Curr. Top. Med. Chem. 20 (2020). https://doi.org/10.2174/1568026620666200127141550.

[14] M.C. Joshi, J. Okombo, S. Nsumiwa, J. Ndove, D. Taylor, L. Wiesner, R. Hunter, K. Chibale, T.J. Egan, 4-Aminoquinoline Antimalarials Containing a Benzylmethylpyridylmethylamine Group Are Active against Drug Resistant Plasmodium falciparum and Exhibit Oral Activity in Mice, J. Med. Chem. 60 (2017). https://doi.org/10.1021/acs.jmedchem.7b01537.

[15] S. Kumar, M. Kumar, R. Ekka, J.D. Dvorin, A.S. Paul, A.K. Madugundu, T. Gilberger, H. Gowda, M.T. Duraisingh, T.S. Keshava Prasad, P. Sharma, PfCDPK1 mediated signaling in erythrocytic stages of Plasmodium falciparum, Nat. Commun. 8 (2017) 63. https://doi.org/10.1038/s41467-017-00053-1.

[16] N. Kato, T. Sakata, G. Breton, K.G. Le Roch, A. Nagle, C. Andersen, B. Bursulaya, K. Henson, J. Johnson, K.A. Kumar, F. Marr, D. Mason, C. McNamara, D. Plouffe, V. Ramachandran, M. Spooner, T. Tuntland, Y. Zhou, E.C. Peters, A. Chatterjee, P.G. Schultz, G.E. Ward, N. Gray, J. Harper, E.A. Winzeler, Gene expression signatures and small-molecule compounds link a protein kinase to Plasmodium falciparum motility, Nat. Chem. Biol. 4 (2008) 347–356. https://doi.org/10.1038/nchembio.87.

[17] G. Lemercier, A. Fernandez-Montalvan, J.P. Shaw, D. Kugelstadt, J. Bomke, M. Domostoj, M.K. Schwarz, A. Scheer, B. Kappes, D. Leroy, Identification and characterization of novel small molecules as potent inhibitors of the plasmodial calcium-dependent protein kinase 1, Biochemistry. 48 (2009) 6379–6389. https://doi.org/10.1021/bi9005122.

[18] K.H. Ansell, H.M. Jones, D. Whalley, A. Hearn, D.L. Taylor, E.C. Patin, T.M. Chapman, S.A. Osborne, C. Wallace, K. Birchall, J. Large, N. Bouloc, E. Smiljanic-Hurley, B. Clough, R.W. Moon, J.L. Green, A.A. Holder, Biochemical and Antiparasitic Properties of Inhibitors of the Plasmodium falciparum Calcium-Dependent Protein Kinase PfCDPK1, Antimicrob. Agents Chemother. 58 (2014) 6032–43. https://doi.org/10.1128/aac.02959-14.

[19] T.M. Chapman, S.A. Osborne, N. Bouloc, J.M. Large, C. Wallace, K. Birchall, K.H. Ansell, H.M. Jones, D. Taylor, B. Clough, J.L. Green, A.A. Holder, Substituted imidazopyridazines are potent and selective inhibitors of Plasmodium falciparum calcium-dependent protein kinase 1 (PfCDPK1), Bioorganic Med. Chem. Lett. 23 (2013) 3064–3069. https://doi.org/10.1016/j.bmcl.2013.03.017.

[20] T.M. Chapman, S.A. Osborne, C. Wallace, K. Birchall, N. Bouloc, H.M. Jones, K.H. Ansell, D.L. Taylor, B. Clough, J.L. Green, A.A. Holder, Optimization of an imidazopyridazine series of inhibitors of plasmodium falciparum calcium-dependent protein kinase 1 (Pf CDPK1), J. Med. Chem. 57 (2014) 3570–3587. https://doi.org/10.1021/jm500342d.

[21] R. Kant, D. Kumar, D. Agarwal, R.D. Gupta, R. Tilak, S.K. Awasthi, A. Agarwal, Synthesis of newer 1,2,3-triazole linked chalcone and flavone hybrid compounds and evaluation of their antimicrobial and cytotoxic activities, Elsevier Ltd, 2016. https://doi.org/10.1016/j.ejmech.2016.02.041.

[22] E.M. Guantai, K. Ncokazi, T.J. Egan, J. Gut, P.J. Rosenthal, P.J. Smith, K. Chibale, Design, synthesis and in vitro antimalarial evaluation of triazole-linked chalcone and dienone hybrid compounds, Bioorganic Med. Chem. 18 (2010). https://doi.org/10.1016/j.bmc.2010.10.009.

[23] N. Devender, S. Gunjan, R. Tripathi, R.P. Tripathi, Synthesis and antiplasmodial activity of novel indoleamide derivatives bearing sulfonamide and triazole pharmacophores, Eur. J. Med. Chem. 131 (2017) 171–184. https://doi.org/10.1016/j.ejmech.2017.03.010.

[24] F. Shah, P. Mukherjee, J. Gut, J. Legac, P.J. Rosenthal, B.L. Tekwani, M.A. Avery, Identification of novel malarial cysteine protease inhibitors using structure-based virtual screening of a focused cysteine protease inhibitor library, J. Chem. Inf. Model. 51 (2011) 852–864. https://doi.org/10.1021/ci200029y.

[25] S. Oramas-Royo, P. López-Rojas, Á. Amesty, D. Gutiérrez, N. Flores, P. Martín-Rodríguez, L. Fernández-Pérez, A. Estévez-Braun, Synthesis and antiplasmodial activity of 1,2,3-triazole-naphthoquinone conjugates, Molecules. 24 (2019) 1–23. https://doi.org/10.3390/molecules24213917.

[26] M.C. Joshi, K.J. Wicht, D. Taylor, R. Hunter, P.J. Smith, T.J. Egan, In vitro antimalarial activity, β-haematin inhibition and structure-activity relationships in a series of quinoline triazoles, Eur. J. Med. Chem. 69 (2013). https://doi.org/10.1016/j.ejmech.2013.08.046.

[27] M.V.N. de Souza, K.C. Pais, C.R. Kaiser, M.A. Peralta, M. de L. Ferreira, M.C.S. Lourenço, Synthesis and in vitro antitubercular activity of a series of quinoline derivatives, Bioorganic Med. Chem. 17 (2009). https://doi.org/10.1016/j.bmc.2009.01.013.

[28] B. Aneja, A. Queen, P. Khan, F. Shamsi, A. Hussain, P. Hasan, M.M.A. Rizvi, C.G. Daniliuc, M.F. Alajmi, M. Mohsin, M.I. Hassan, M. Abid, Design, synthesis & biological evaluation of ferulic acid-based small molecule inhibitors against tumor-associated carbonic anhydrase IX, Bioorganic Med. Chem. 28 (2020) 115424. https://doi.org/10.1016/j.bmc.2020.115424.

[29] B. Aneja, M. Azam, S. Alam, A. Perwez, R. Maguire, U. Yadava, K. Kavanagh, C.G. Daniliuc, M.M.A. Rizvi, Q.M.R. Haq, M. Abid, Natural Product-Based 1,2,3- Triazole/Sulfonate Analogues as Potential Chemotherapeutic Agents for Bacterial Infections, ACS Omega. 3 (2018) 6912–6930. https://doi.org/10.1021/acsomega.8b00582.

[30] J.A. Rowe, A. Claessens, R.A. Corrigan, M. Arman, Adhesion of Plasmodium falciparum-infected erythrocytes to human cells: Molecular mechanisms and therapeutic implications, Expert Rev. Mol. Med. 11 (2009) 1–29. https://doi.org/10.1017/S1462399409001082.

[31] N.T. Huy, Y. Shima, A. Maeda, T.T. Men, K. Hirayama, A. Hirase, A. Miyazawa, K. Kamei, Phospholipid Membrane-Mediated Hemozoin Formation: The Effects of Physical Properties and Evidence of Membrane Surrounding Hemozoin, PLoS One. 8 (2013). https://doi.org/10.1371/journal.pone.0070025.

[32] A.P. Gorka, J.N. Alumasa, K.S. Sherlach, L.M. Jacobs, K.B. Nickley, J.P. Brower, A.C. De Dios, P.D. Roepe, Cytostatic versus cytocidal activities of chloroquine analogues and inhibition of hemozoin crystal growth, Antimicrob. Agents Chemother. 57 (2013) 356–364. https://doi.org/10.1128/AAC.01709-12.

[33] N.T. Huy, K. Kamei, T. Yamamoto, Y. Kondo, K. Kanaori, R. Takano, K. Tajima, S. Hara, Clotrimazole binds to heme and enhances heme-dependent hemolysis: Proposed antimalarial mechanism of clotrimazole, J. Biol. Chem. 277 (2002) 4152–4158. https://doi.org/10.1074/jbc.M107285200.

[34] S.E. Francis, D.J. Sullivan, D.E. Goldberg, Hemoglobin metabolism in the malaria parasite Plasmodium falciparium, Annu. Rev. Microbiol. 51 (1997) 97–123. https://doi.org/10.1146/annurev.micro.51.1.97.

[35] B. Tekwani, L. Walker, Targeting the Hemozoin Synthesis Pathway for New Antimalarial Drug Discovery: Technologies for In Vitro &#946;-Hematin Formation Assay, Comb. Chem. High Throughput Screen. 8 (2005) 63–79. https://doi.org/10.2174/1386207053328101.

[36] S.E.A. Ozbabacan, H.B. Engin, A. Gursoy, O. Keskin, Transient proteinprotein interactions, Protein Eng. Des. Sel. 24 (2011). https://doi.org/10.1093/protein/gzr025.

[37] C.H. Kaschula, T.J. Egan, R. Hunter, N. Basilico, S. Parapini, D. Taramelli, E. Pasini, D. Monti, Structure - Activity relationships in 4-aminoquinoline antiplasmodials. The role of the group at the 7-position, J. Med. Chem. 45 (2002). https://doi.org/10.1021/jm020858u.

[38] F. Xu, Z.Z. Yang, J.R. Jiang, W.G. Pan, X. Le Yang, J.Y. Wu, Y. Zhu, J. Wang, Q.Y. Shou, H.G. Wu, Synthesis, antitumor evaluation and molecular docking studies of [1,2,4]triazolo[4,3-b][1,2,4,5]tetrazine derivatives, Bioorganic Med. Chem. Lett. 26 (2016). https://doi.org/10.1016/j.bmcl.2016.05.007.

[39] H.A.M. El-Sherief, B.G.M. Youssif, S.N.A. Bukhari, M. Abdel-Aziz, H.M. Abdel-Rahman, Novel 1,2,4-triazole derivatives as potential anticancer agents: Design, synthesis, molecular docking and mechanistic studies, Bioorg. Chem. 76 (2018). https://doi.org/10.1016/j.bioorg.2017.12.013.

[40] H.A.M. Goma’a, M.A. Ghaly, L.A. Abou-zeid, F.A. Badria, I.A. Shehata, M.M. El-Kerdawy, Synthesis, Biological Evaluation and In Silico Studies of 1,2,4-Triazole and 1,3,4-Thiadiazole Derivatives as Antiherpetic Agents, ChemistrySelect. 4 (2019). https://doi.org/10.1002/slct.201900814.

[41] R. Jain, S. Gupta, M. Munde, S. Pati, S. Singh, Development of novel anti-malarial from structurally diverse library of molecules, targeting plant-like CDPK1, a multistage growth regulator of P. falciparum, Biochem. J. 447 (2020). https://doi.org/10.1042/BCJ20200045.

[42] B. Aneja, R. Arif, A. Perwez, J. V. Napoleon, P. Hasan, M.M.A. Rizvi, A. Azam, Rahisuddin, M. Abid, N-Substituted 1,2,3-Triazolyl-Appended Indole-Chalcone Hybrids as Potential DNA Intercalators Endowed with Antioxidant and Anticancer Properties, ChemistrySelect. 3 (2018) 2638–2645. https://doi.org/10.1002/slct.201702913.

[43] K. Xue, G. Sun, Y. Zhang, X. Chen, Y. Zhou, J. Hou, H. Long, Z. Zhang, M. Lei, W. Wu, A new method for the synthesis of chalcone derivatives promoted by PPh3/I2under non-alkaline conditions, Synth. Commun. 51 (2021). https://doi.org/10.1080/00397911.2020.1847295.

[44] V. Rajkumar, S.A. Babu, R. Padmavathi, Regio- and diastereoselective construction of a new set of functionalized pyrrolidine, spiropyrrolidine and spiropyrrolizidine scaffolds appended with aryl- and heteroaryl moieties via the azomethine ylide cycloadditions, Tetrahedron. 72 (2016). https://doi.org/10.1016/j.tet.2016.07.053.

[45] S.A. Gamage, J.A. Spicer, G.J. Atwell, G.J. Finlay, B.C. Baguley, W.A. Denny, Structure-activity relationships for substituted bis(acridine-4- carboxamides): A new class of anticancer agents, J. Med. Chem. 42 (1999). https://doi.org/10.1021/jm980687m.

[46] A. Uddin, V. Singh, I. Irfan, T. Mohammad, R.S. Hada, M.I. Hassan, M. Abid, S. Singh, Identification and structure–activity relationship (SAR) studies of carvacrol derivatives as potential anti-malarial against Plasmodium falciparum falcipain-2 protease, Bioorg. Chem. 103 (2020). https://doi.org/10.1016/j.bioorg.2020.104142.

[47] M. Irfan, B. Aneja, U. Yadava, S.I. Khan, N. Manzoor, C.G. Daniliuc, M. Abid, Synthesis, QSAR and anticandidal evaluation of 1,2,3-triazoles derived from naturally bioactive scaffolds, Eur. J. Med. Chem. 93 (2015) 246–254. https://doi.org/10.1016/j.ejmech.2015.02.007.

[48] H.B. Reilly, H. Wang, J.A. Steuter, A.M. Marx, M.T. Ferdig, Quantitative dissection of clone-specific growth rates in cultured malaria parasites, Int. J. Parasitol. 37 (2007). https://doi.org/10.1016/j.ijpara.2007.05.003.

[49] M.T. Makler, J.M. Ries, J.A. Williams, J.E. Bancroft, R.C. Piper, B.L. Gibbins, D.J. Hinrichs, Parasite lactate dehydrogenase as an assay for Plasmodium falciparum drug sensitivity, Am. J. Trop. Med. Hyg. 48 (1993) 739–741. https://doi.org/10.4269/ajtmh.1993.48.739.

[50] V. Singh, R.S. Hada, A. Uddin, B. Aneja, M. Abid, K.C. Pandey, S. Singh, Inhibition of Hemoglobin Degrading Protease Falcipain-2 as a Mechanism for Anti-Malarial Activity of Triazole-Amino Acid Hybrids, Curr. Top. Med. Chem. 20 (2020) 377–389. https://doi.org/10.2174/1568026620666200130162347.

[51] T.T. Men, N.T. Huy, D.T.X. Trang, M.N. Shuaibu, K. Hirayama, K. Kamei, A simple and inexpensive haemozoin-based colorimetric method to evaluate anti-malarial drug activity, Malar. J. 11 (2012). https://doi.org/10.1186/1475-2875-11-272.

[52] B.C. Evans, C.E. Nelson, S.S. Yu, K.R. Beavers, A.J. Kim, H. Li, H.M. Nelson, T.D. Giorgio, C.L. Duvall, Ex vivo red blood cell hemolysis assay for the evaluation of pH-responsive endosomolytic agents for cytosolic delivery of biomacromolecular drugs., J. Vis. Exp. (2013). https://doi.org/10.3791/50166.

[53] H.J. Woerdenbag, T.A. Moskal, N. Pras, T.M. Malingré, F.S. El-Feraly, H.H. Kampinga, A.W.T. Konings, Cytotoxicity of artemisinin-related endoperoxides to Ehrlich ascites tumor cells, J. Nat. Prod. 56 (1993) 849–856. https://doi.org/10.1021/np50096a007.

[54] F.A. Wani, Amaduddin, B. Aneja, G. Sheehan, K. Kavanagh, R. Ahmad, M. Abid, R. Patel, Synthesis of Novel Benzimidazolium Gemini Surfactants and Evaluation of Their Anti-Candida Activity, ACS Omega. 4 (2019). https://doi.org/10.1021/acsomega.9b01056.

[55] A. Daina, O. Michielin, V. Zoete, SwissADME: A free web tool to evaluate pharmacokinetics, drug-likeness and medicinal chemistry friendliness of small molecules, Sci. Rep. 7 (2017). https://doi.org/10.1038/srep42717.

[56] L. Zhang, H. Ai, W. Chen, Z. Yin, H. Hu, J. Zhu, J. Zhao, Q. Zhao, H. Liu, CarcinoPred-EL: Novel models for predicting the carcinogenicity of chemicals using molecular fingerprints and ensemble learning methods, Sci. Rep. 7 (2017). https://doi.org/10.1038/s41598-017-02365-0.

[57] S. Duhr, D. Braun, Why molecules move along a temperature gradient, Proc. Natl. Acad. Sci. U. S. A. 103 (2006). https://doi.org/10.1073/pnas.0603873103.

[58] C.J. Wienken, P. Baaske, U. Rothbauer, D. Braun, S. Duhr, Protein-binding assays in biological liquids using microscale thermophoresis, Nat. Commun. 1 (2010). https://doi.org/10.1038/ncomms1093.

[59] R. Jain, P. Dey, S. Gupta, S. Pati, A. Bhattacherjee, M. Munde, S. Singh, Molecular dynamics simulations and biochemical characterization of Pf14-3-3 and PfCDPK1 interaction towards its role in growth of human malaria parasite, Biochem. J. 477 (2020). https://doi.org/10.1042/BCJ20200145.

[60] D. Ramu, R. Jain, R.R. Kumar, V. Sharma, S. Garg, R. Ayana, T. Luthra, P. Yadav, S. Sen, S. Singh, Design and synthesis of imidazolidinone derivatives as potent anti- leishmanial agents by bioisosterism, Arch. Pharm. (Weinheim). 352 (2019). https://doi.org/10.1002/ardp.201800290.

