## Supplementary information for "Biological evaluation of novel side chain containing CQTrICh-analogs as antimalarials and their development as *Pf*CDPK1 kinase inhibitors"

**Table of Contents**

1. Spectral data of compounds	S.1-S.94

2. Possible fragmentation pattern in the mass spectrometry	S.95

3. Growth Inhibition assay	S.96

4. Physicochemical properties	Table S1

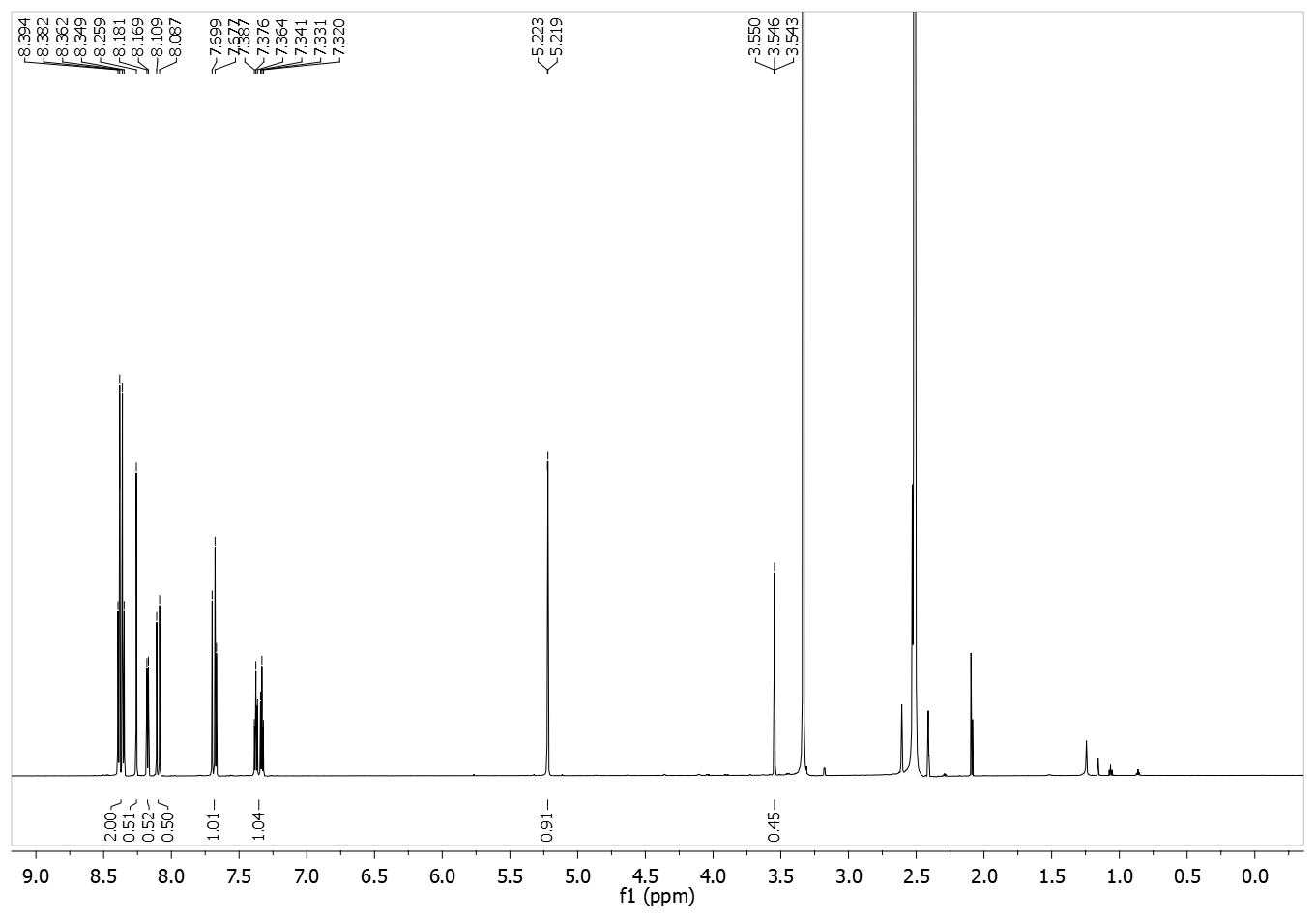
**1.** (*E*)-1-(4-nitrophenyl)-3-(1-(prop-2-yn-1-yl)-1H-indol-3-yl)prop-2-en-1-one **(4d):**

**(4d):**

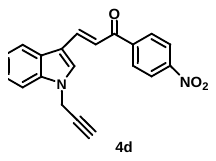

**(E)-3-(1-((1-(7-chloroquinolin-4-yl)-1H-1,2,3-triazol-4-yl)methyl)-1H-indol-3-yl)-1-phenylprop-2-en-1-one (7a)**:

**Figure S.1:** ^1^H NMR spectrum of **4d.**

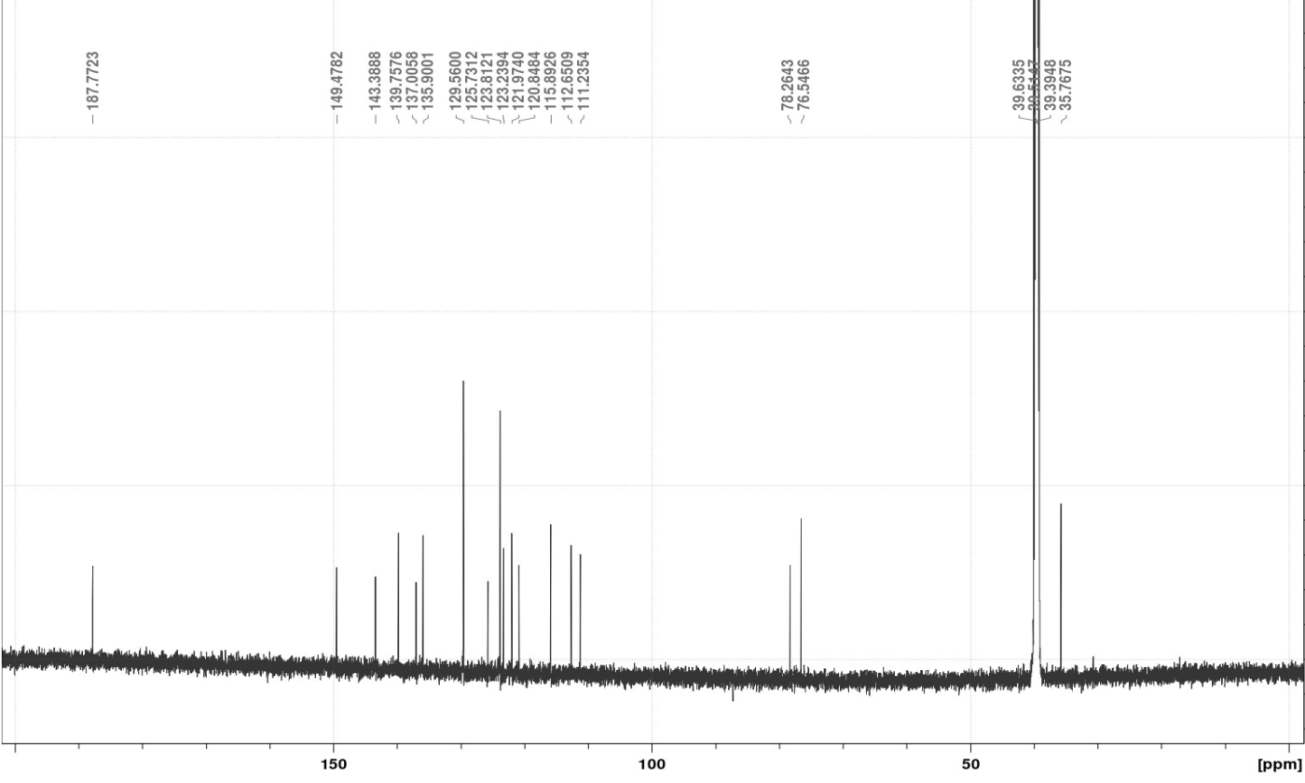

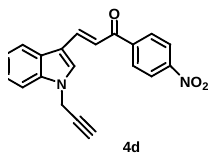

**Figure S.2:** ^13^C NMR spectrum of **4d.**

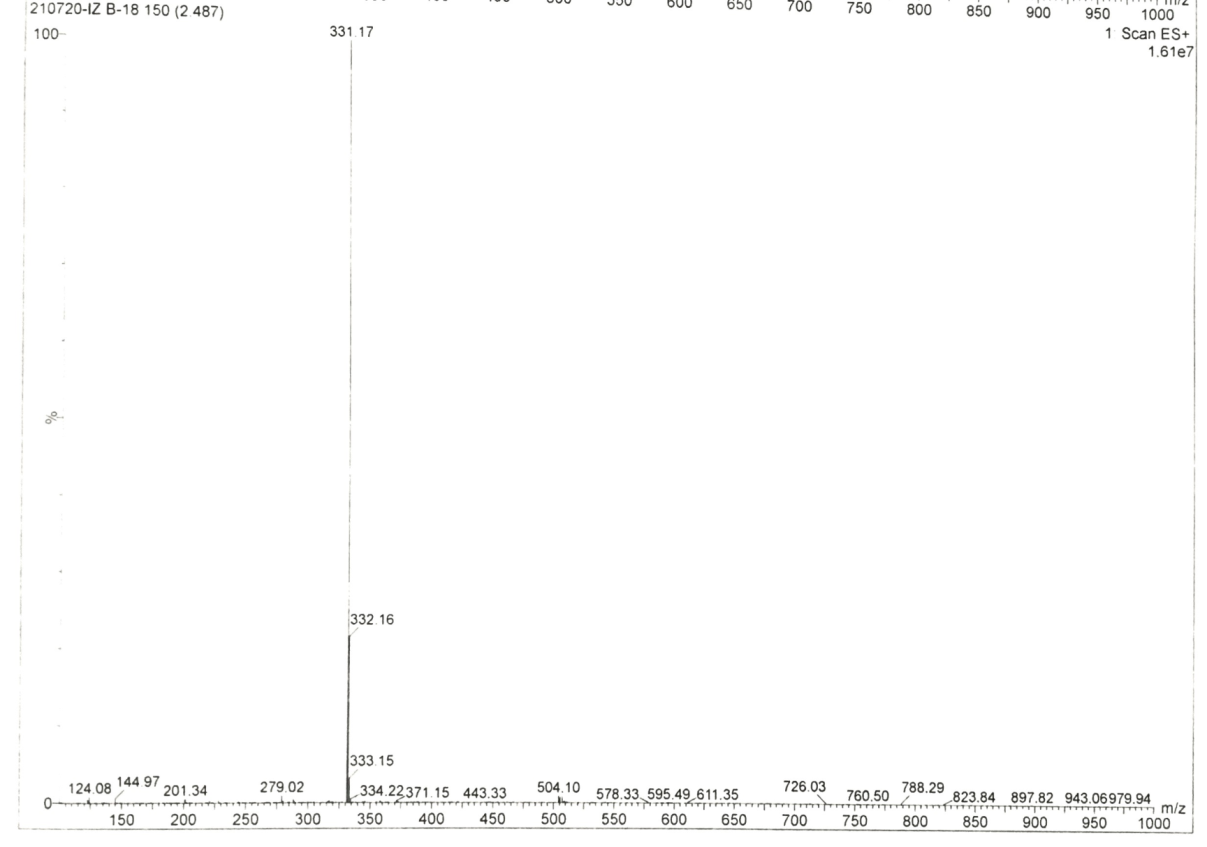

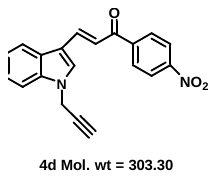

**Figure S.3:** Mass spectrum of **4d.**

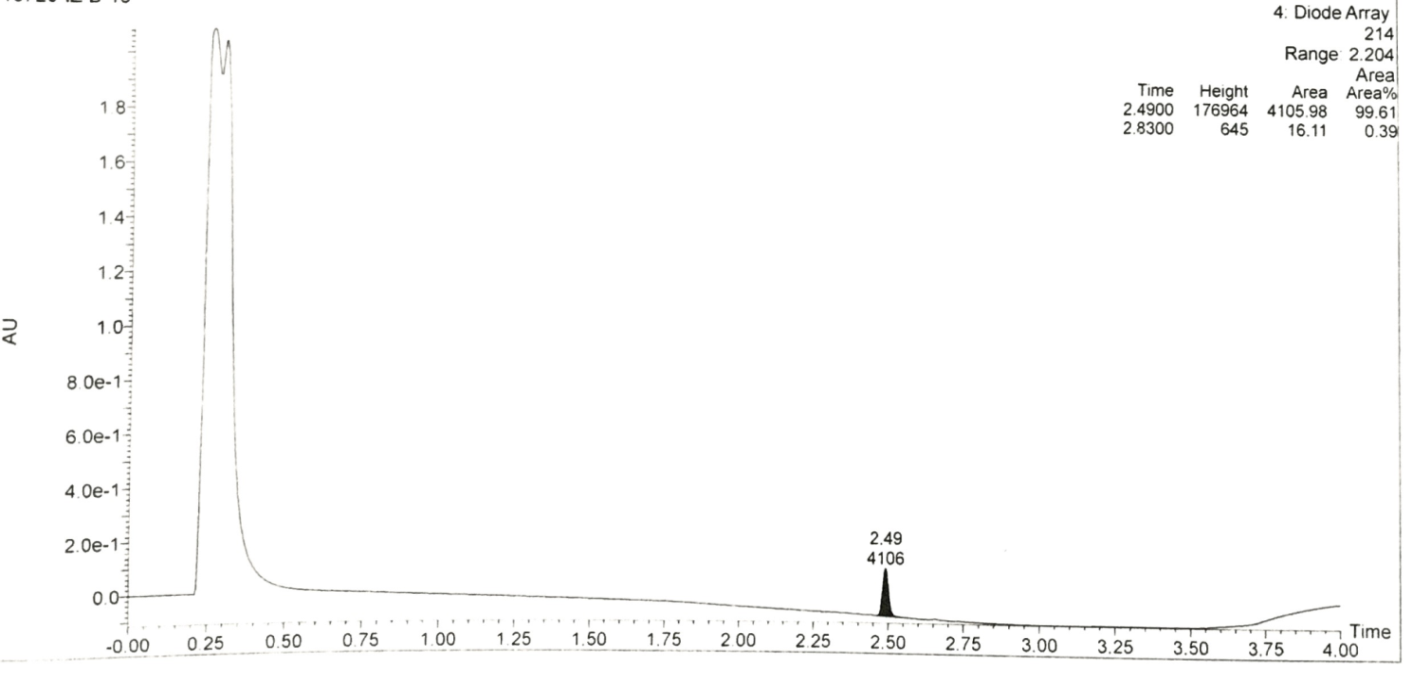

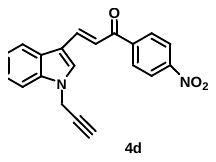

**Figure S.4:** Purity analysis of **4d.**

**2.** (*E*)-1-(2-chlorophenyl)-3-(1-(prop-2-yn-1-yl)-1H-indol-3-yl)prop-2-en-1-one **(4h):**

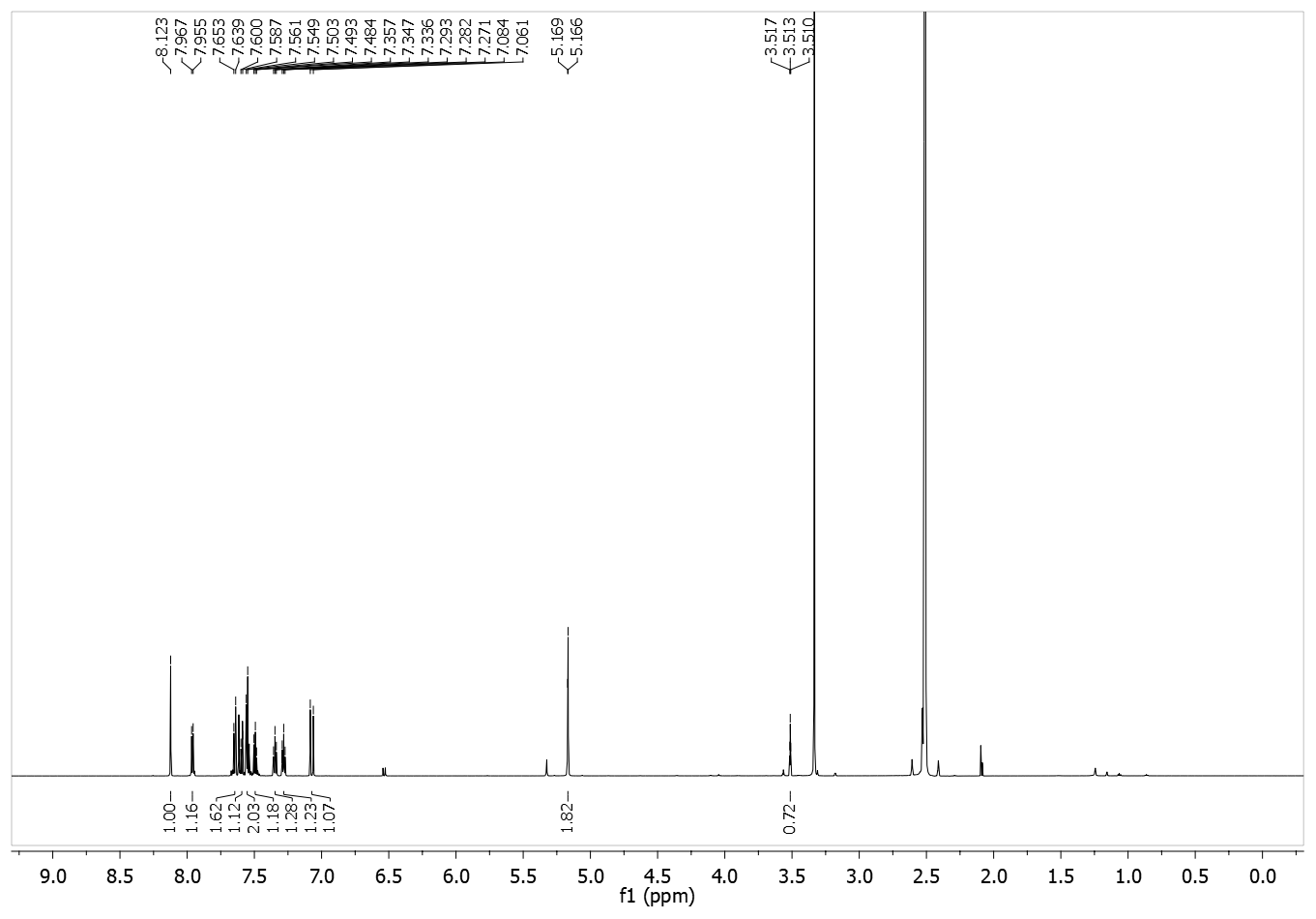

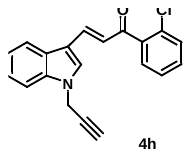

**Figure S.5:** ^1^H NMR spectrum of **4h.**

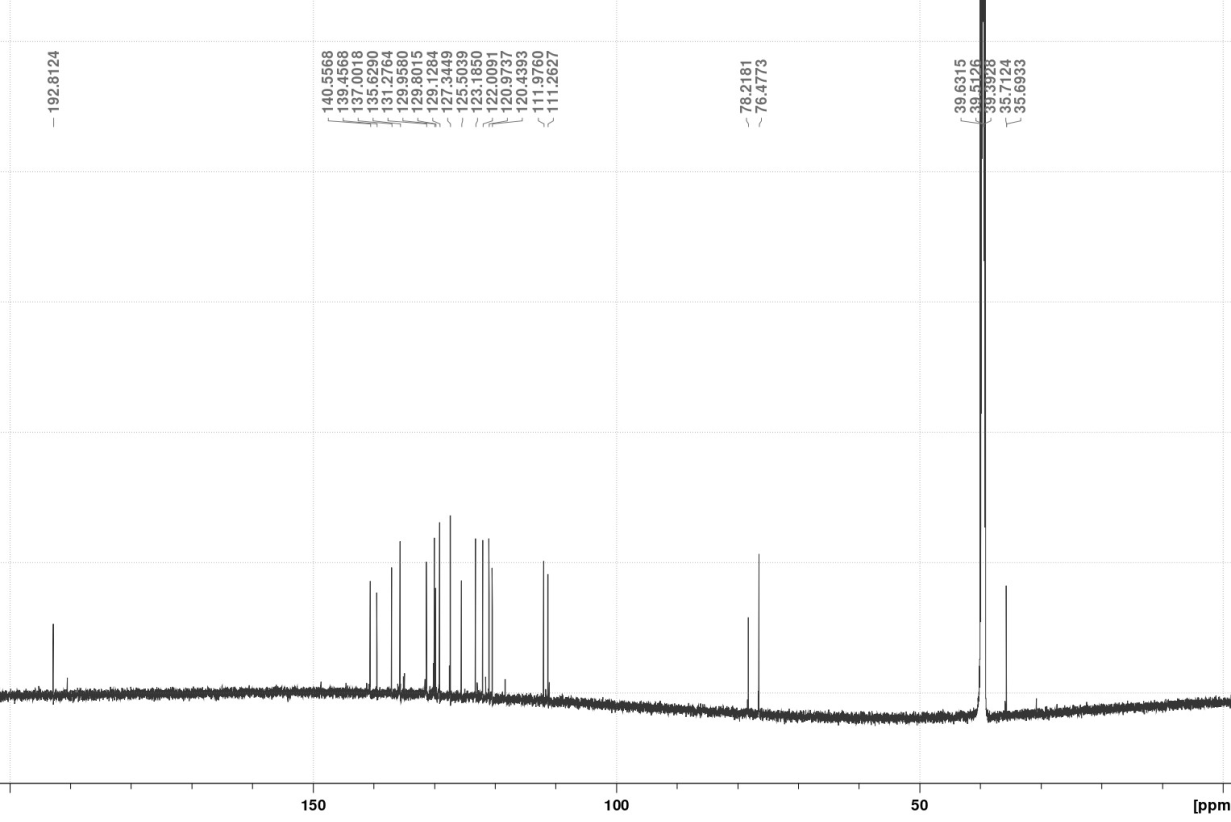

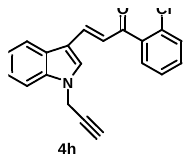

**Figure S.6:** ^13^C NMR spectrum of **4h.**

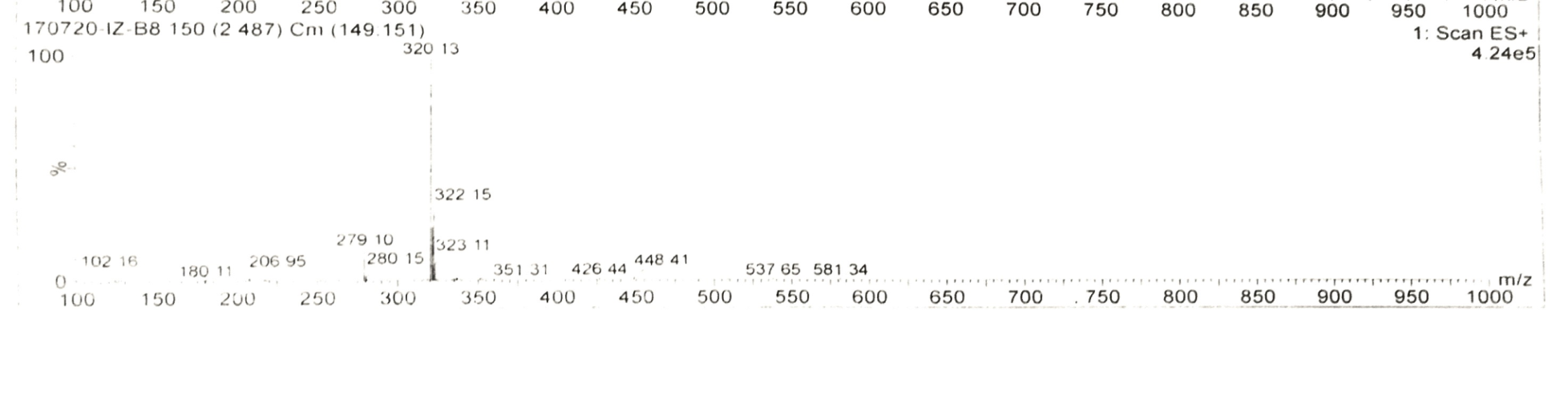

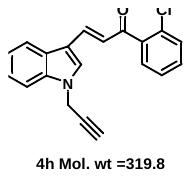

**Figure S.7:** Mass spectrum of **4h.**

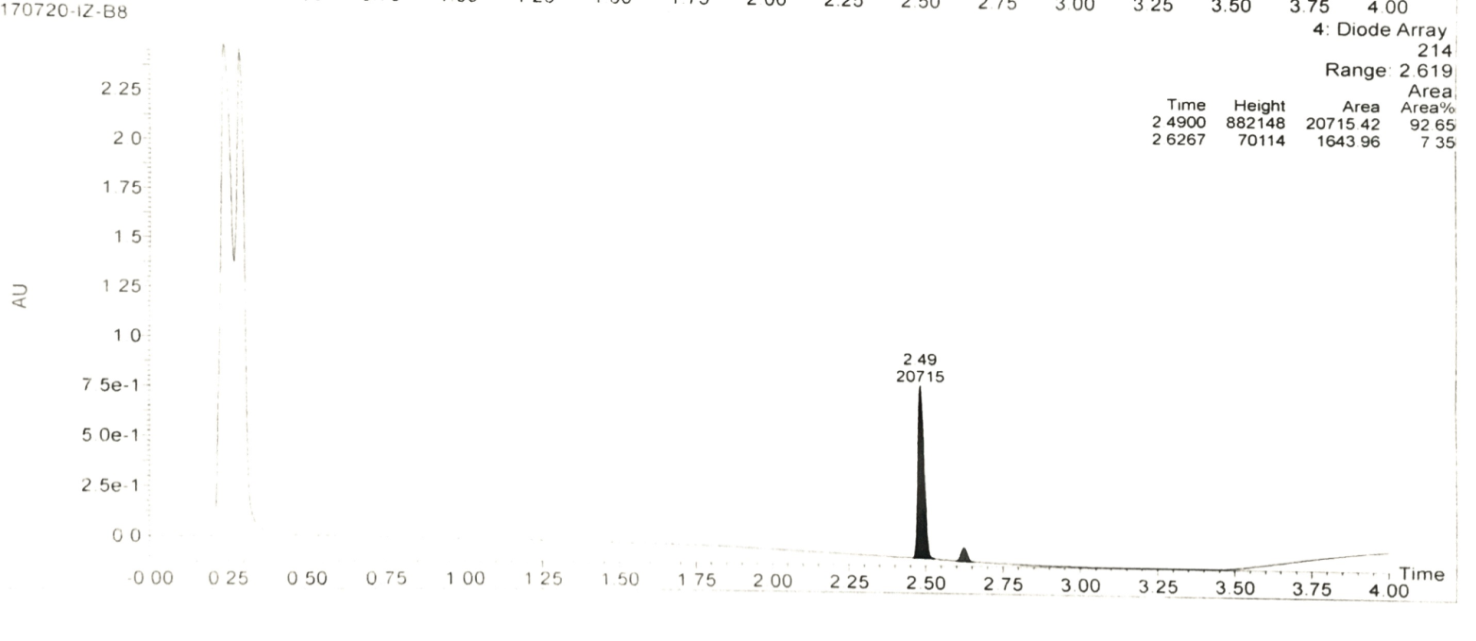

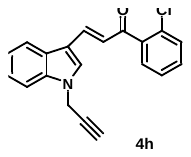

**Figure S.8:** Mass spectrum of **4h.**

**3.** (*E*)-1-(3-chlorophenyl)-3-(1-(prop-2-yn-1-yl)-1H-indol-3-yl)prop-2-en-1-one **(4i):**

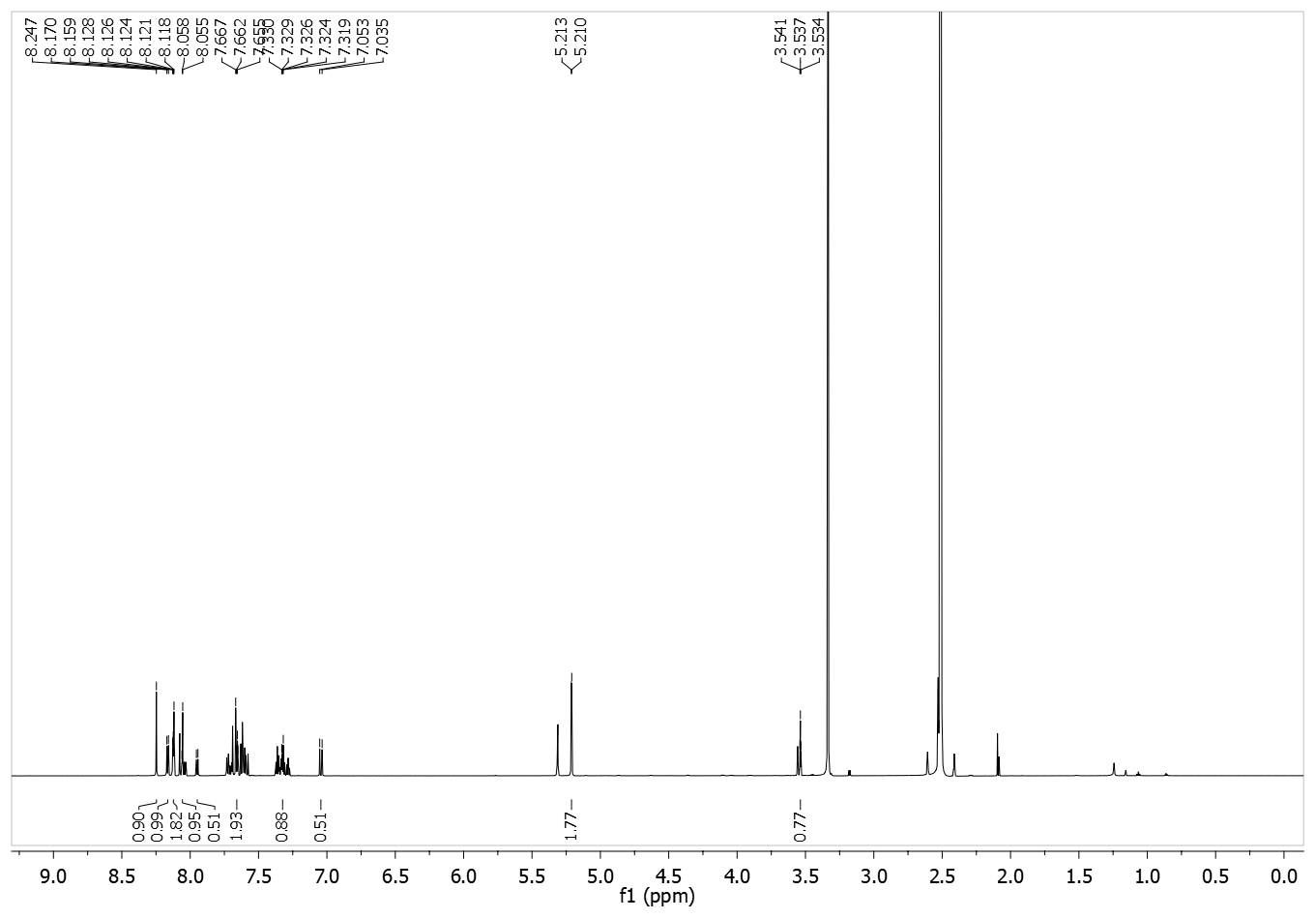

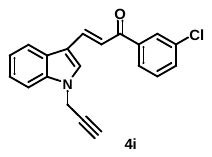

**Figure S.9:** ^1^H NMR spectrum of **4i.**

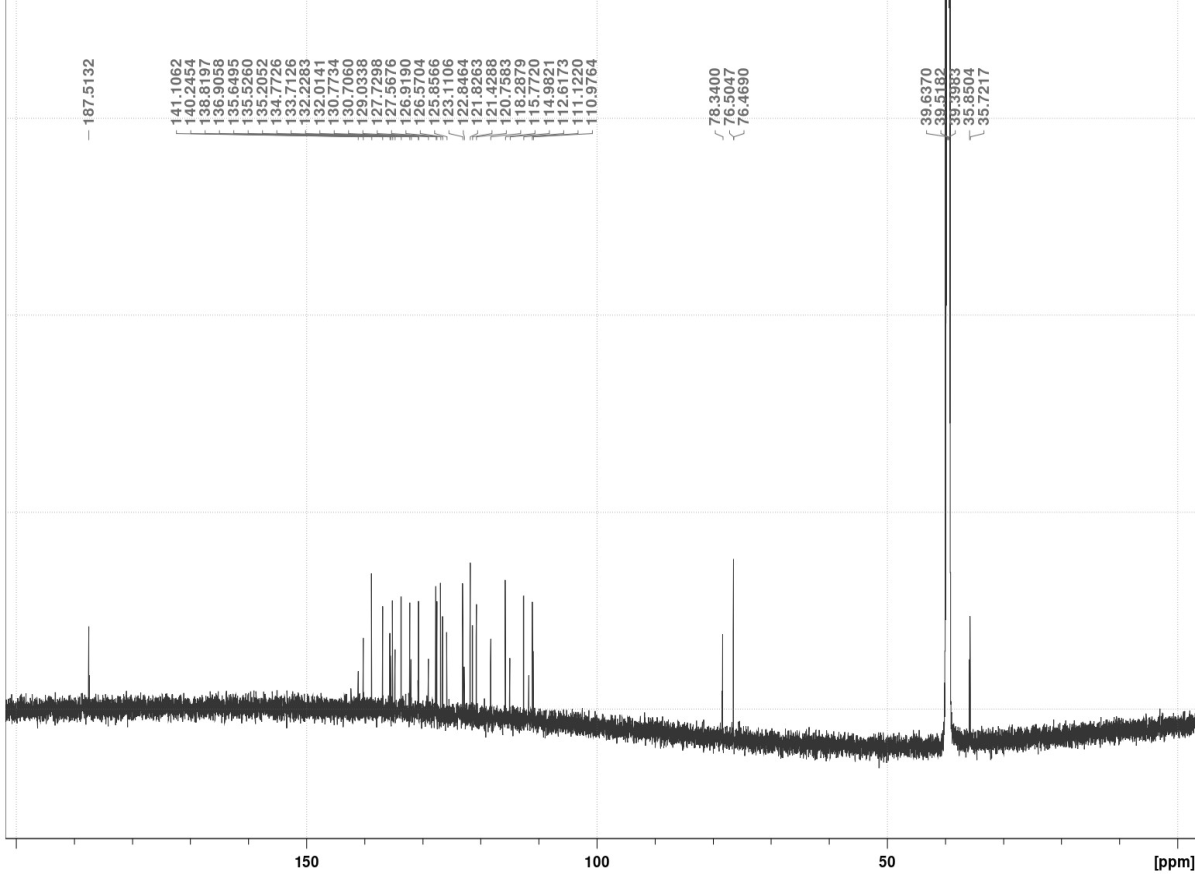

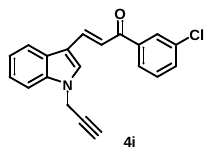

**Figure S.10:** ^13^C NMR spectrum of **4i.**

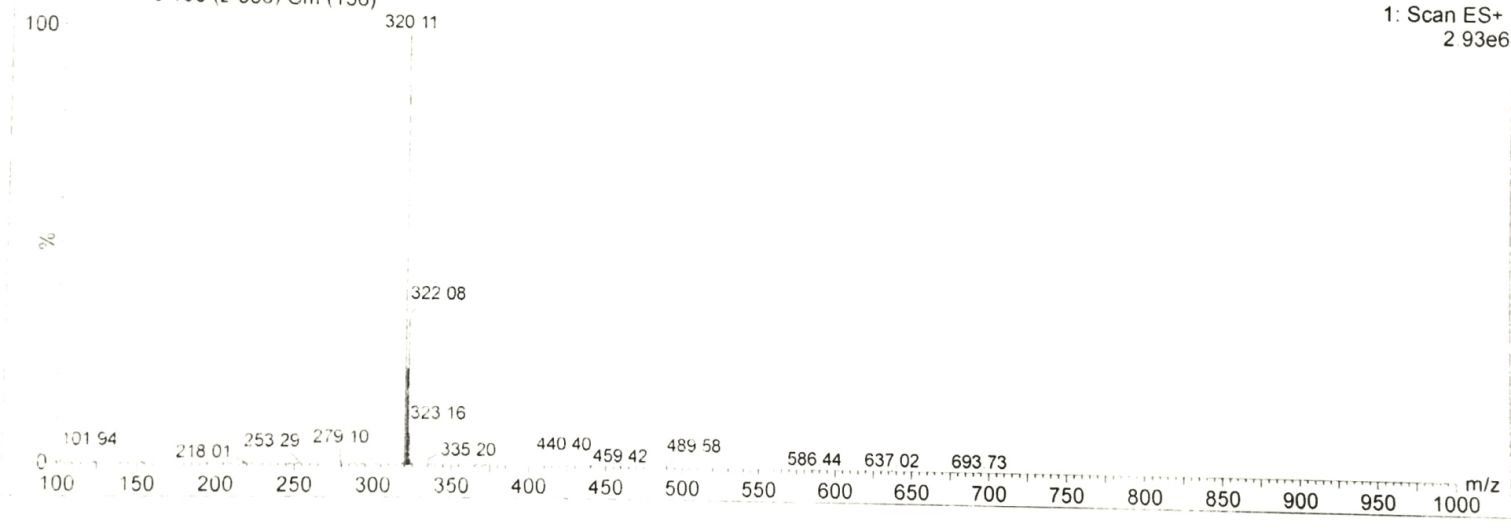

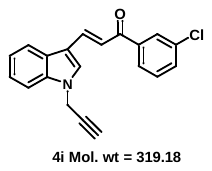

**Figure S.11:** Mass spectrum of **4i.**

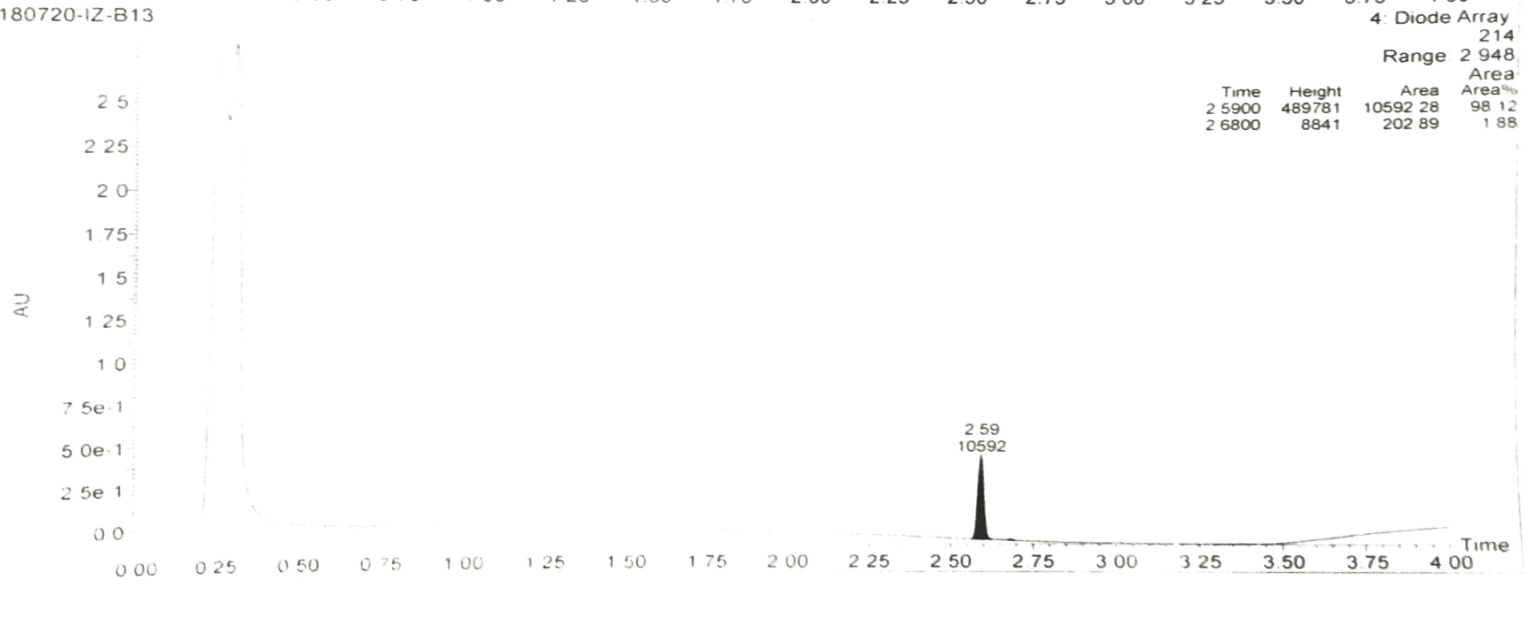

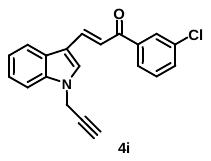

**Figure S.12:** Purity of **4i.**

**4.** (*E*)-1-(2,4-dichlorophenyl)-3-(1-(prop-2-yn-1-yl)-1H-indol-3-yl)prop-2-en-1-one	**(4k):**

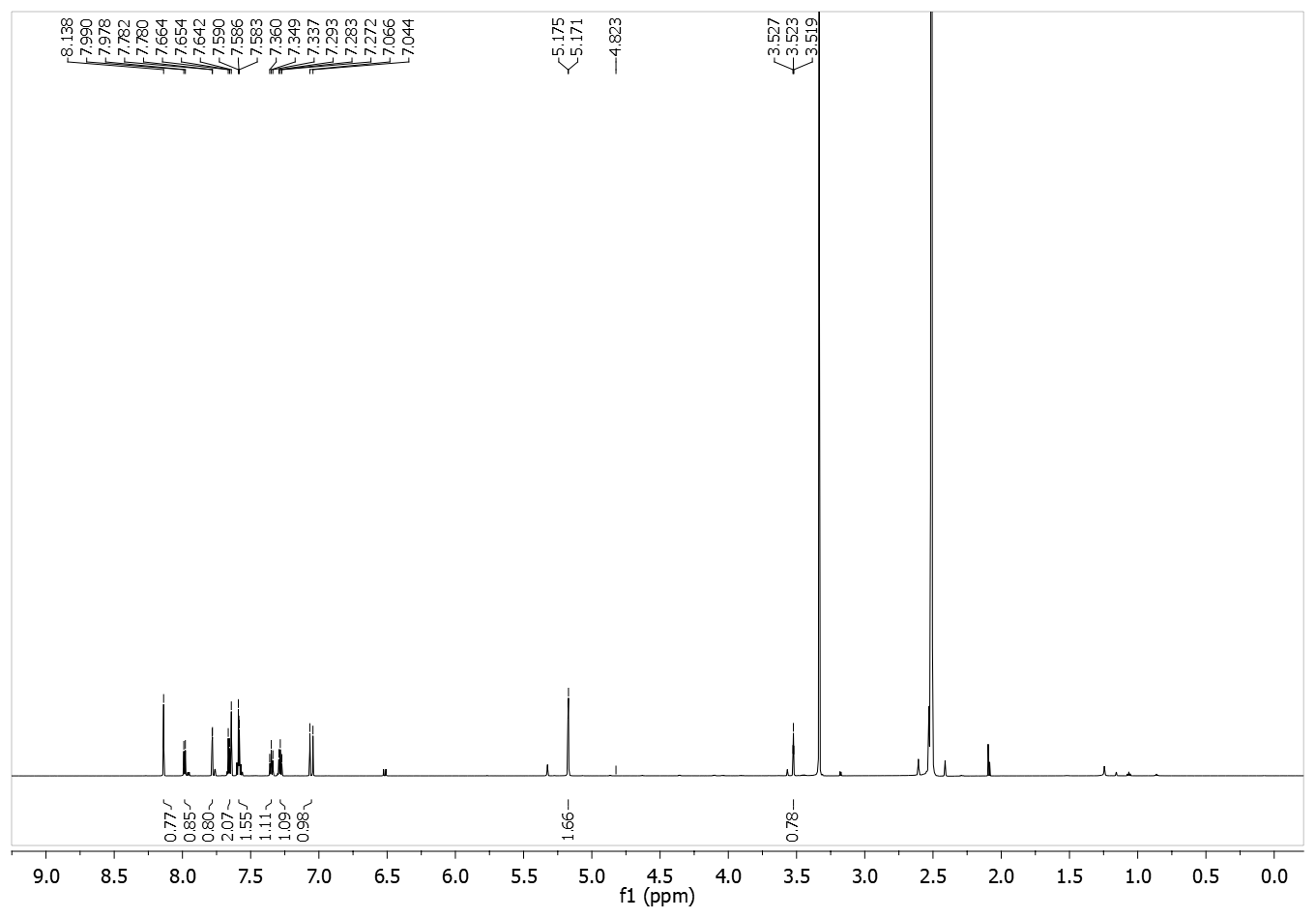

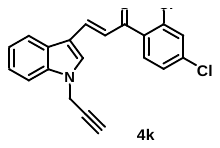

**Figure S.13:** ^1^H NMR spectrum of **4k.**

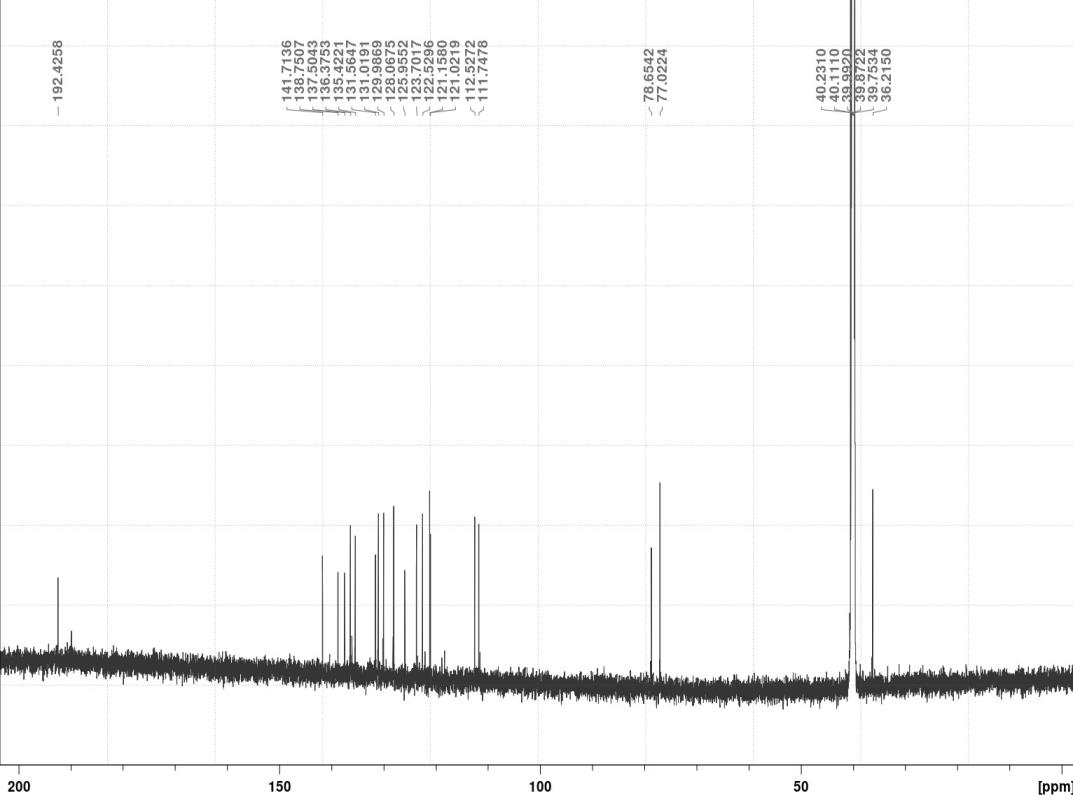

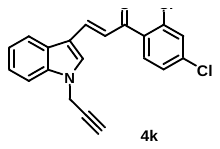

**Figure S.14:** ^1^H NMR spectrum of **4k.**

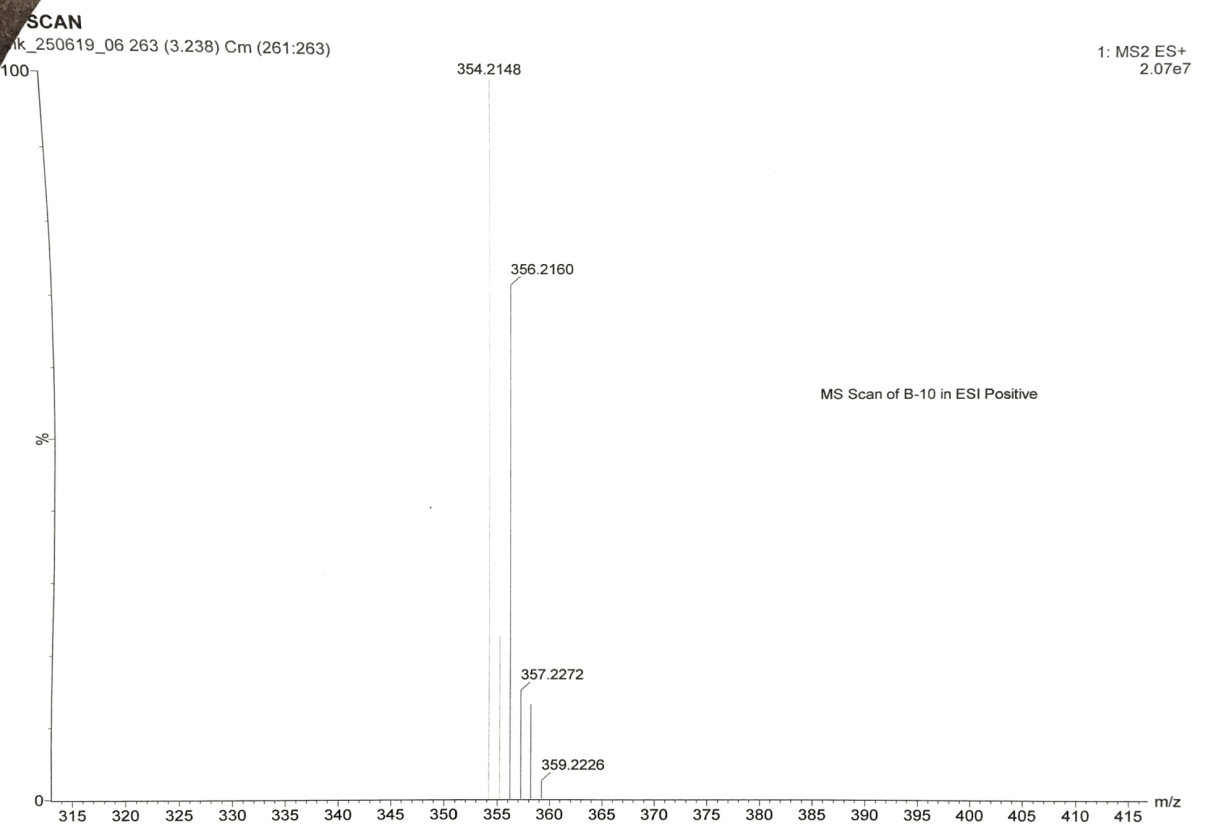

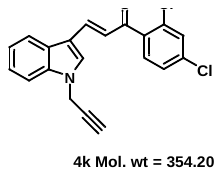

**Figure S.15:** Mass spectrum of **4k.**

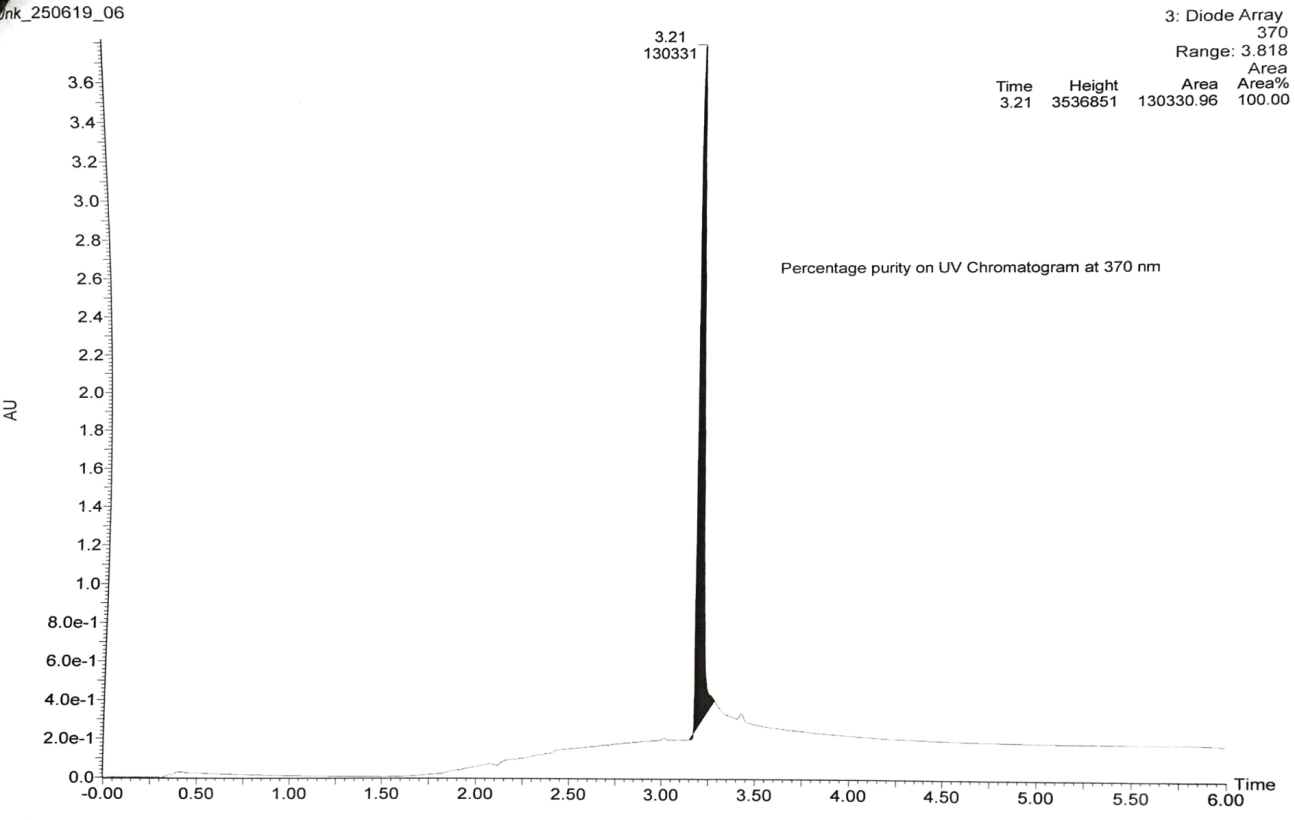

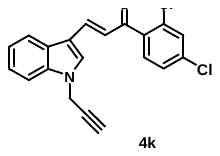

**Figure S.16:** Mass spectrum of **4k.**

**5.** (*E*)-1-(4-methoxyphenyl)-3-(1-(prop-2-yn-1-yl)-1H-indol-3-yl)prop-2-en-1-one **(4n):**

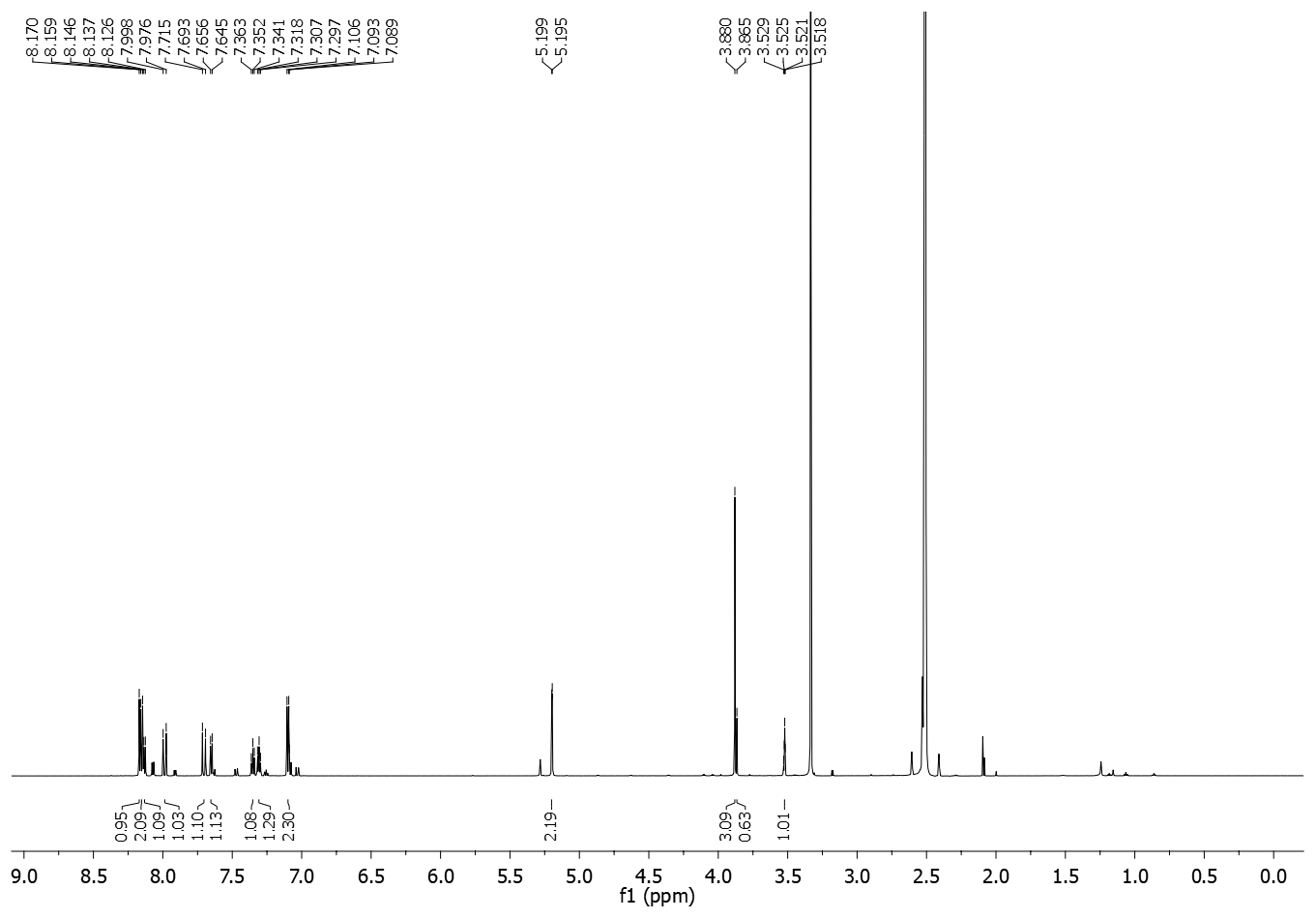

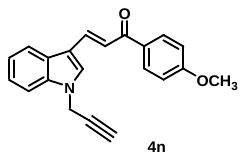

**Figure S.17:** ^1^H NMR spectrum of **4n.**

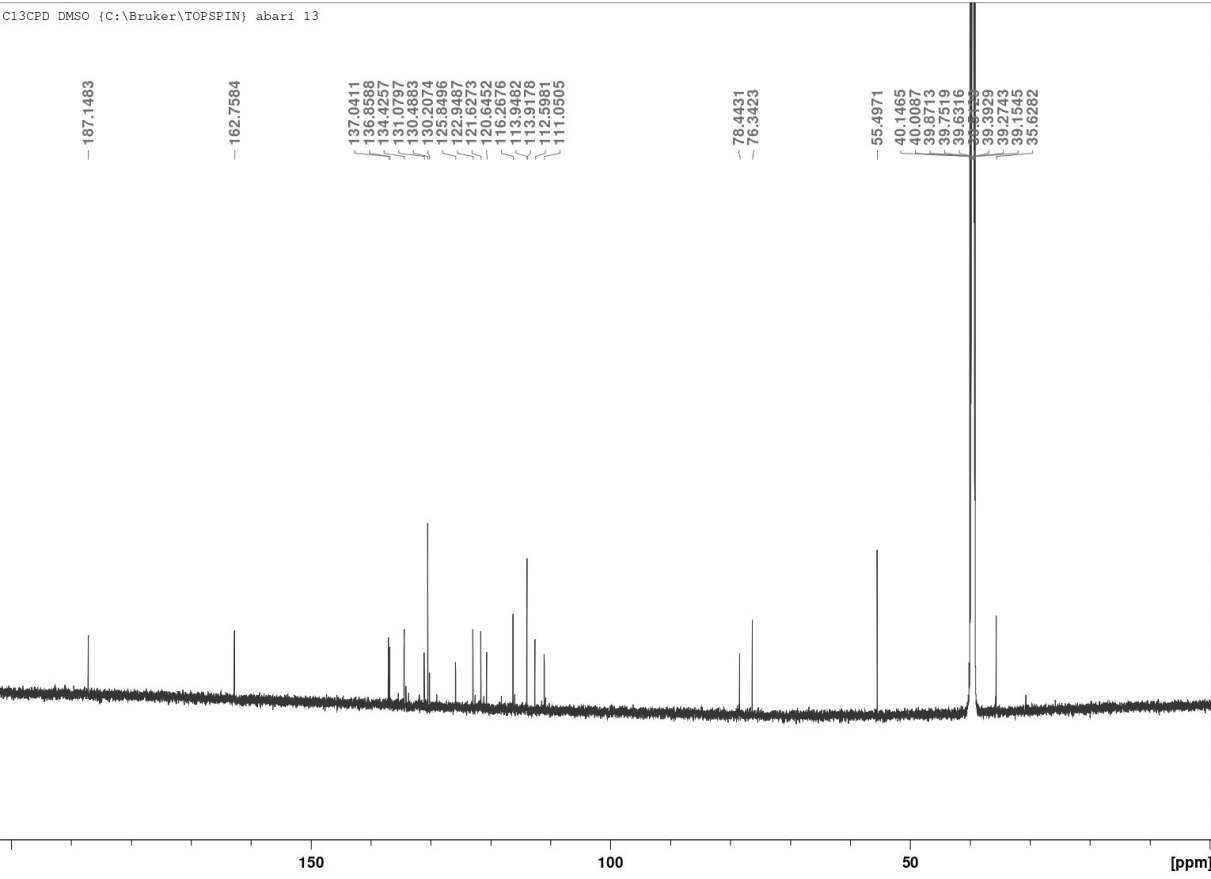

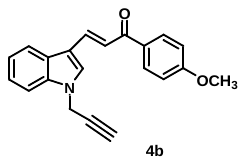

**Figure S.18:** ^1^H NMR spectrum of **4n.**

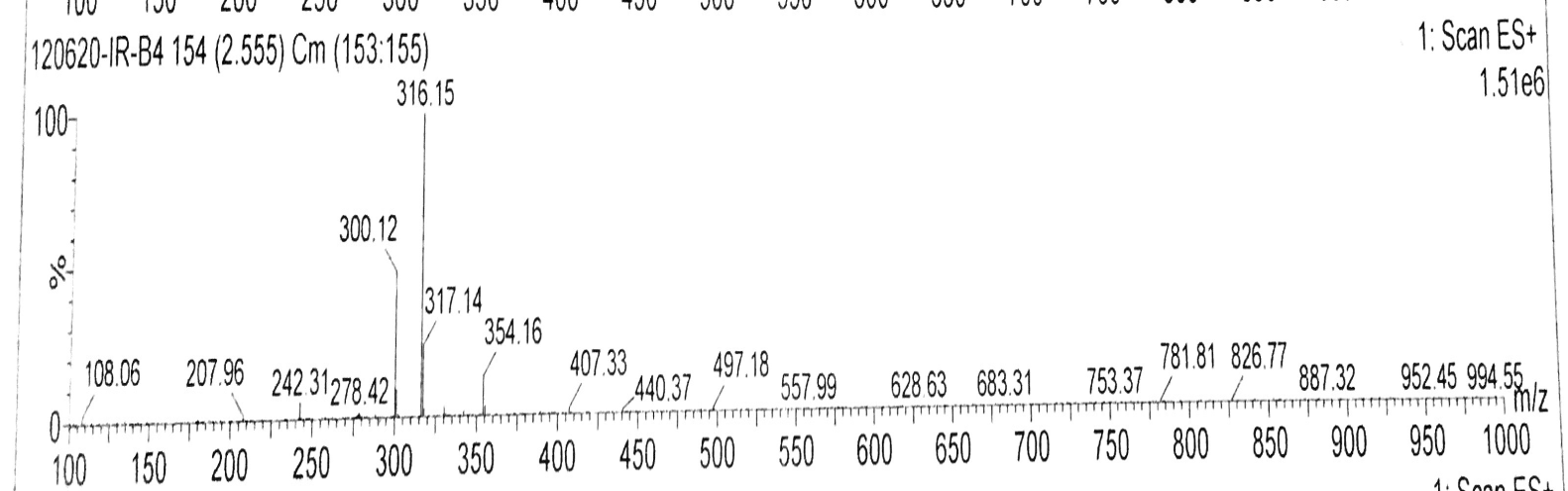

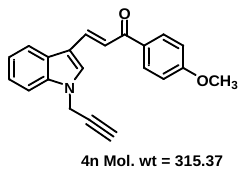

**Figure S.19:** Mass spectrum of **4n.**

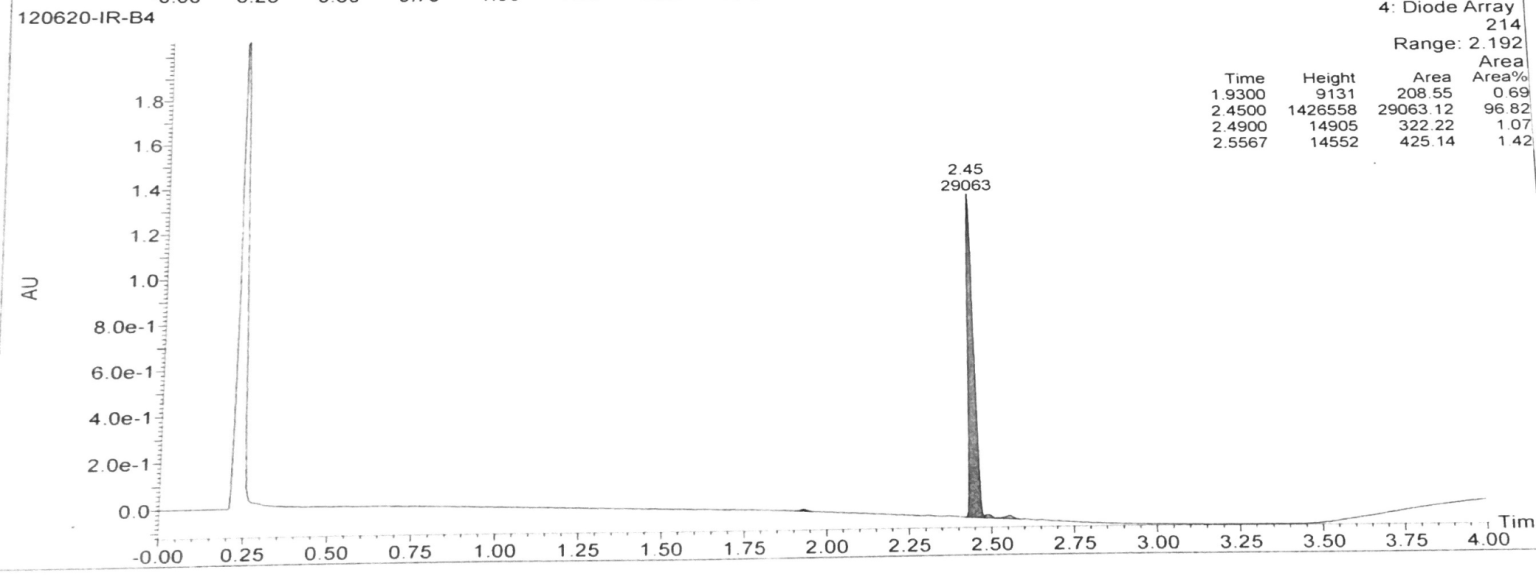

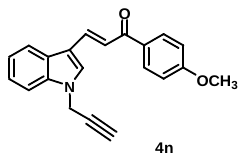

**Figure S.20:** Purity of **4n.**

**6.** (*E*)-1-(3,4-dimethoxyphenyl)-3-(1-(prop-2-yn-1-yl)-1H-indol-3-yl)prop-2-en-1-one **(4o):**

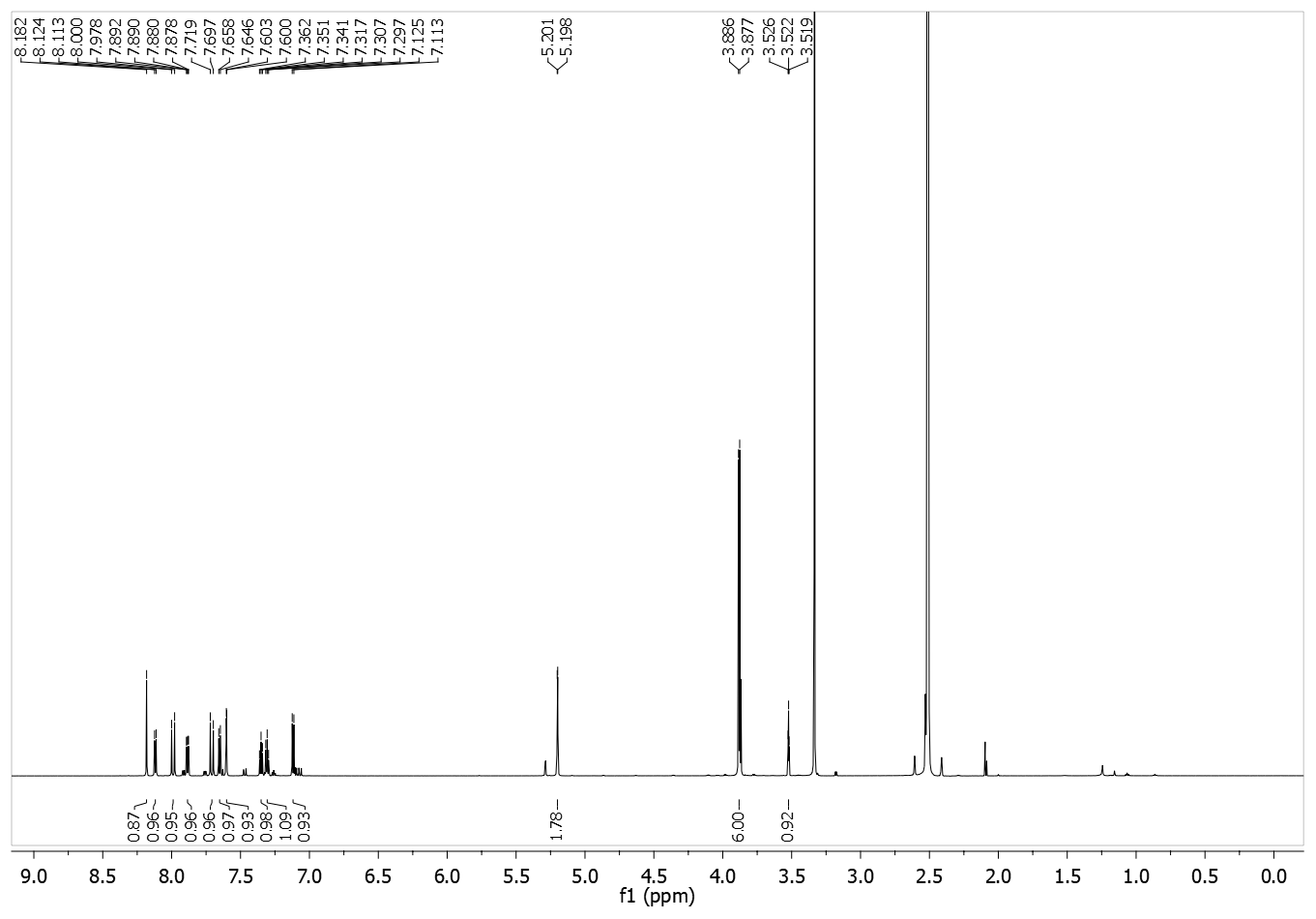

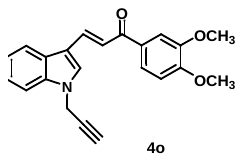

**Figure S.21:** ^1^H NMR spectrum of **4o.**

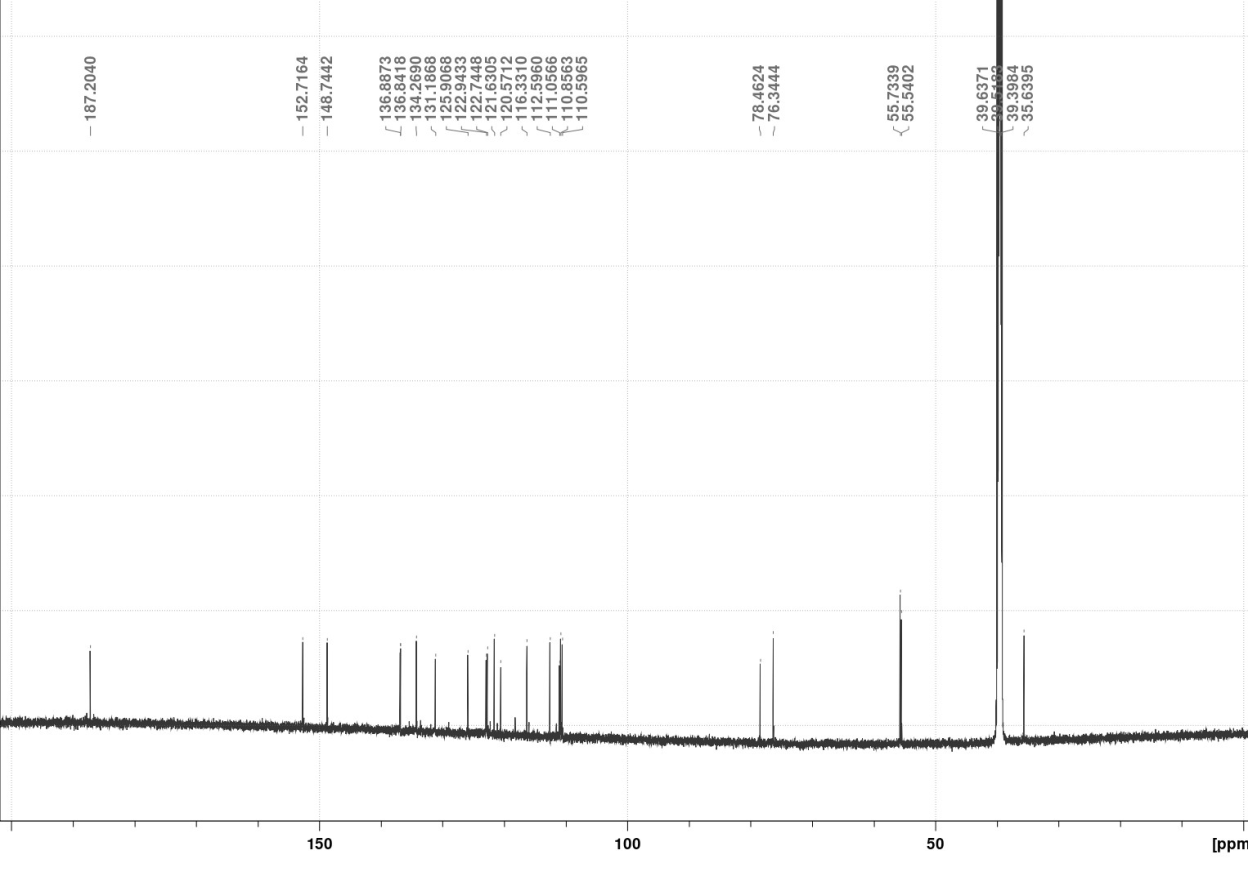

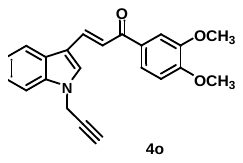

**Figure S.22:** ^12^C NMR spectrum of **4o.**

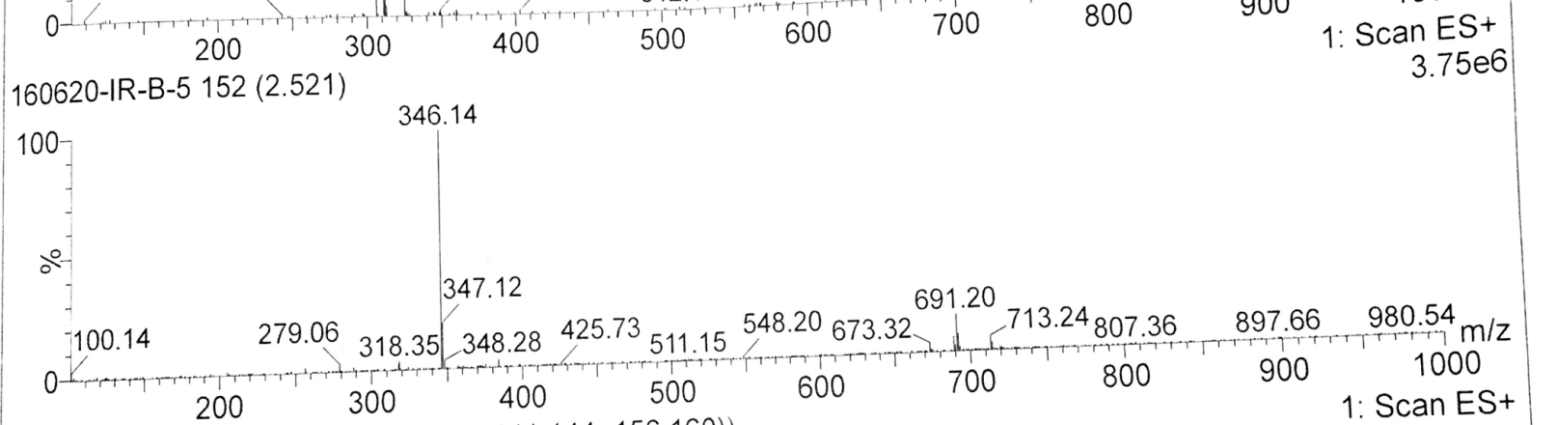

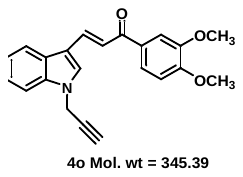

**Figure S.23:** Mass spectrum of **4o.**

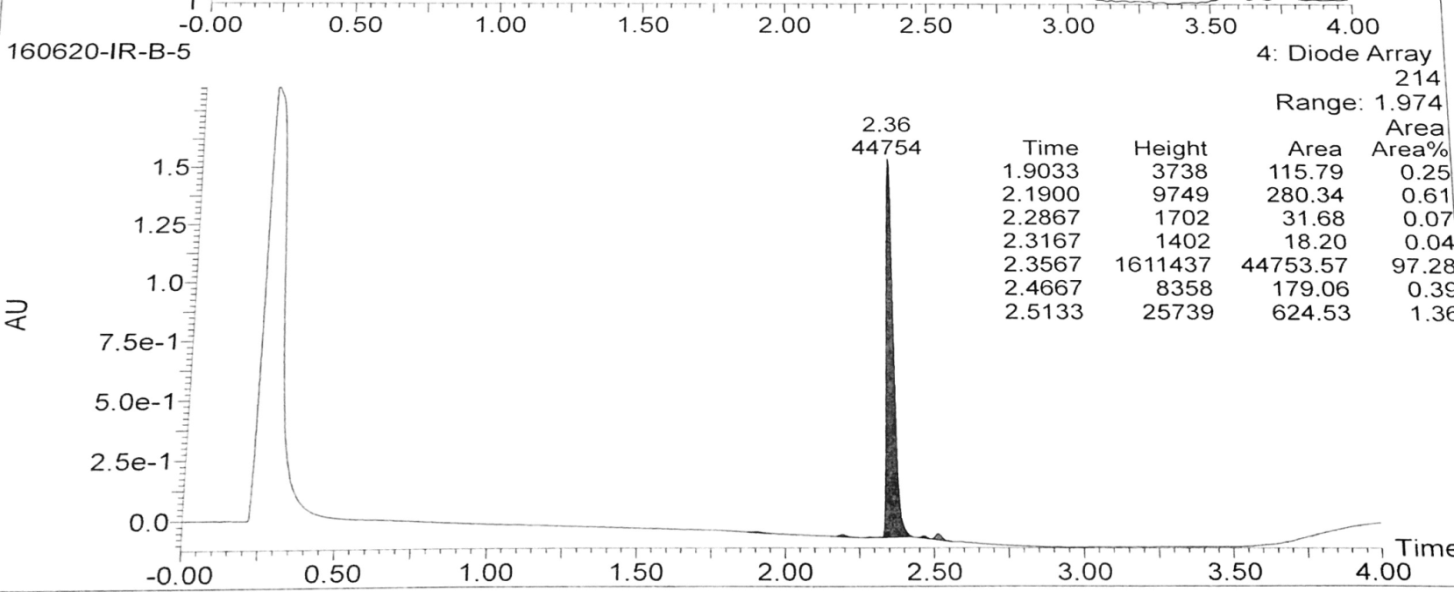

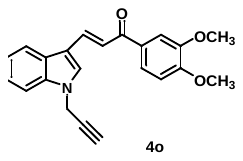

**Figure S.24:** Purity of **4o.**

**7.** (*E*)-1-(3,5-bis(benzyloxy)phenyl)-3-(1-(prop-2-yn-1-yl)-1H-indol-3-yl)prop-2-en-1-one **(4q):**

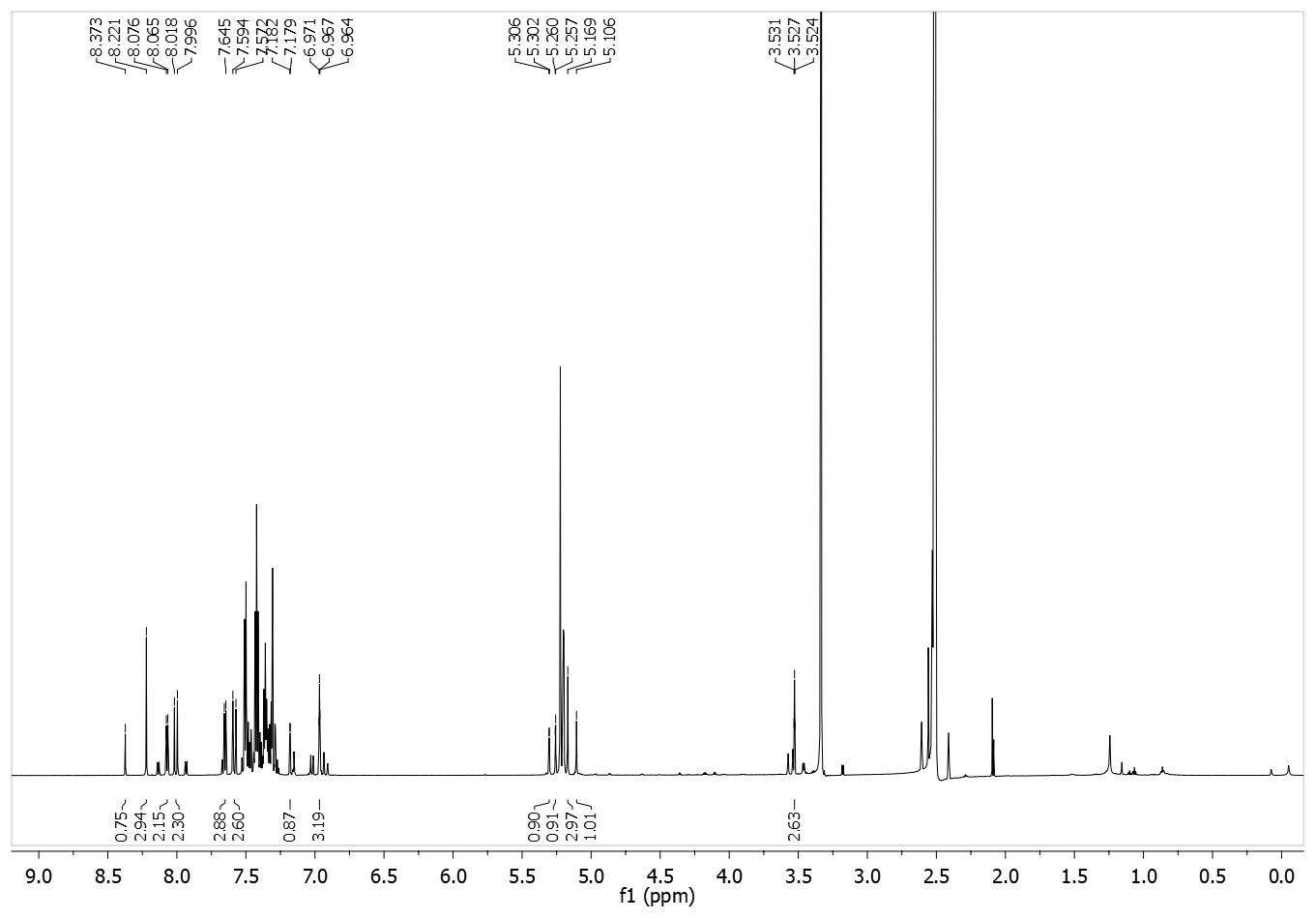

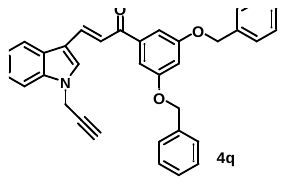

**Figure S.25:** ^1^H NMR spectrum of **4q.**

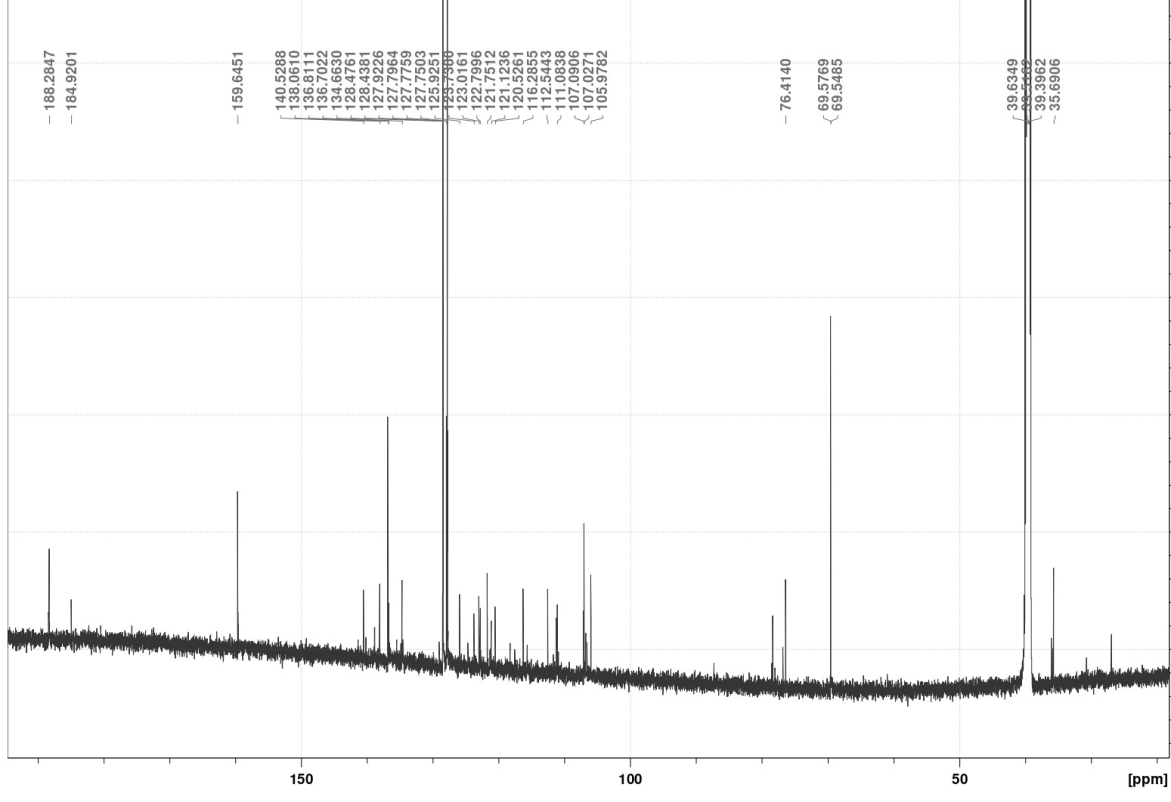

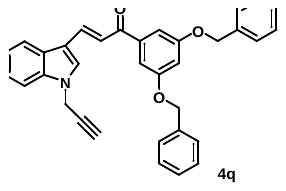

**Figure S.26:** ^13^C NMR spectrum of **4q.**

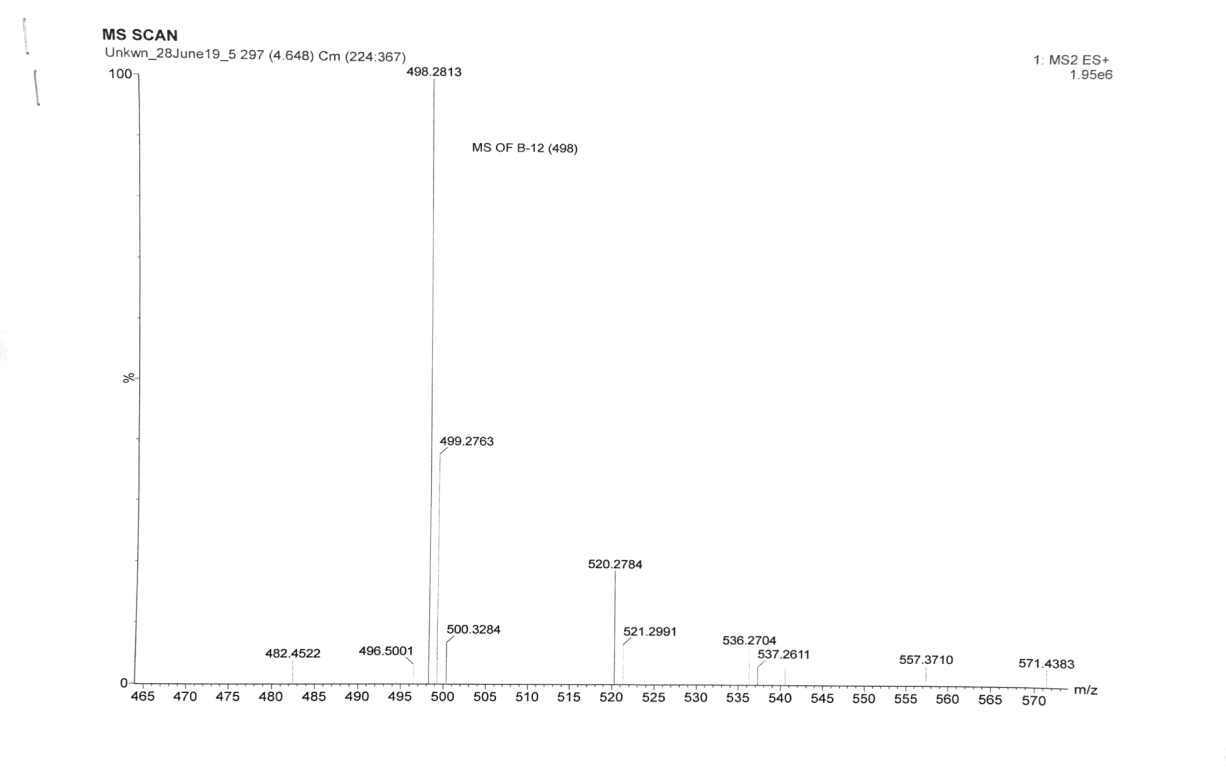

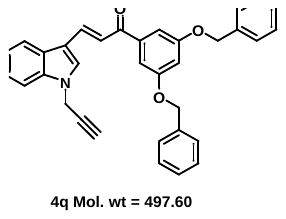

**Figure S.27:** Mass spectrum of **4q.**

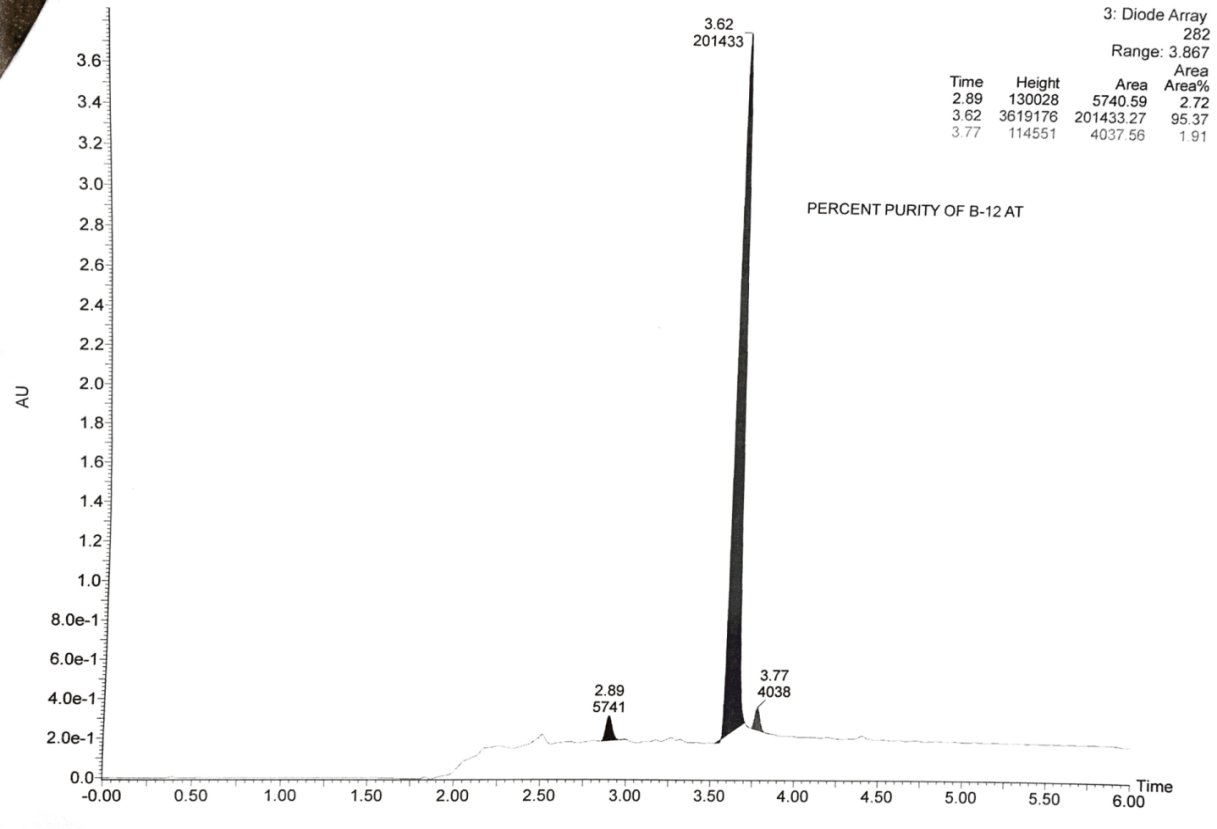

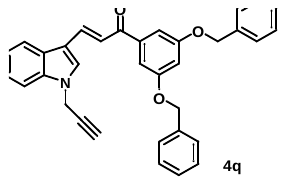

**Figure S.28:** Purity of **4q.**

**8.** 1-(prop-2-yn-1-yl)-1H-indole-3-carbaldehyde **(8):**

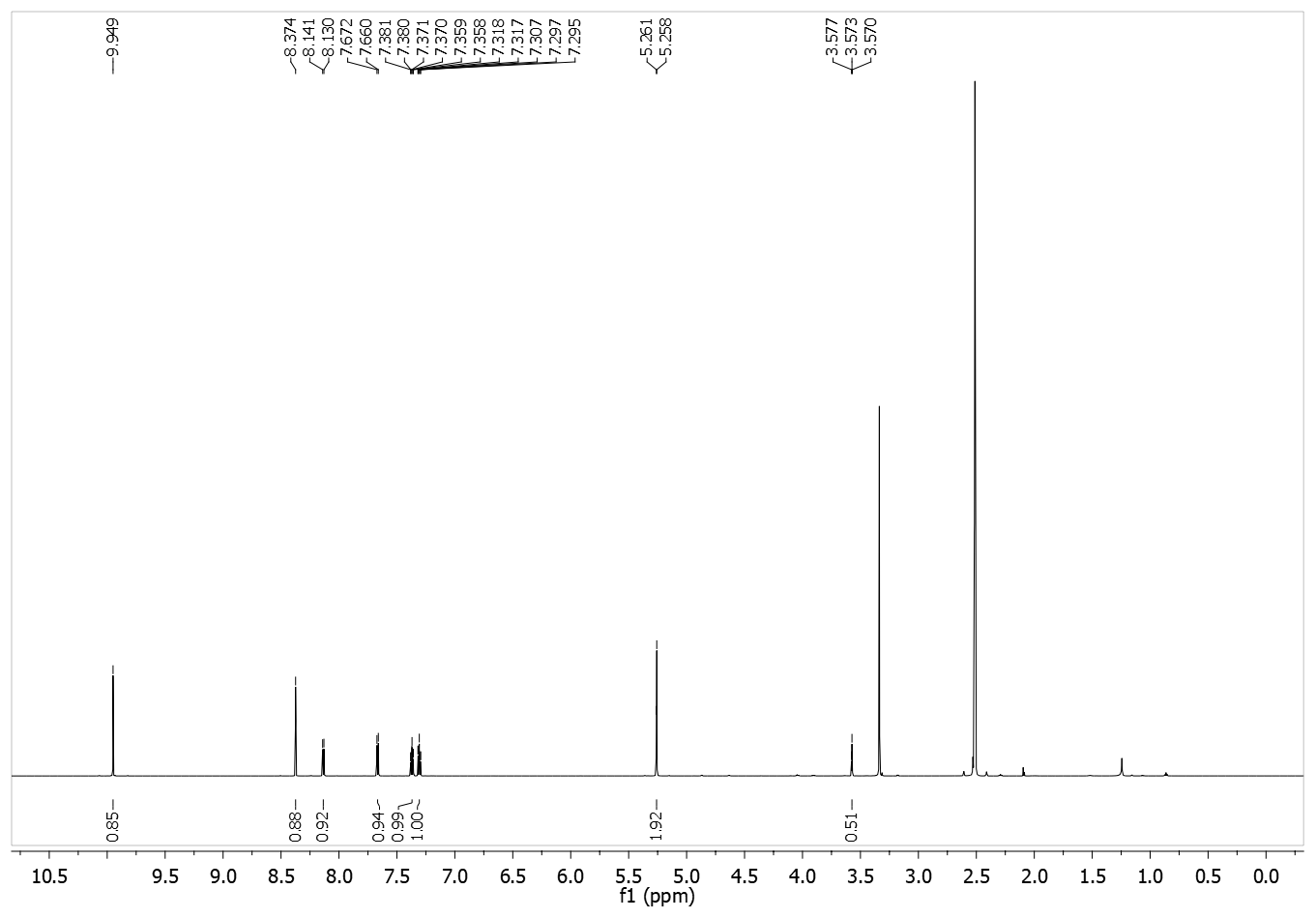

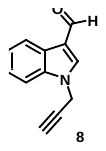

**Figure S.29:** ^1^H NMR spectrum of **8.**

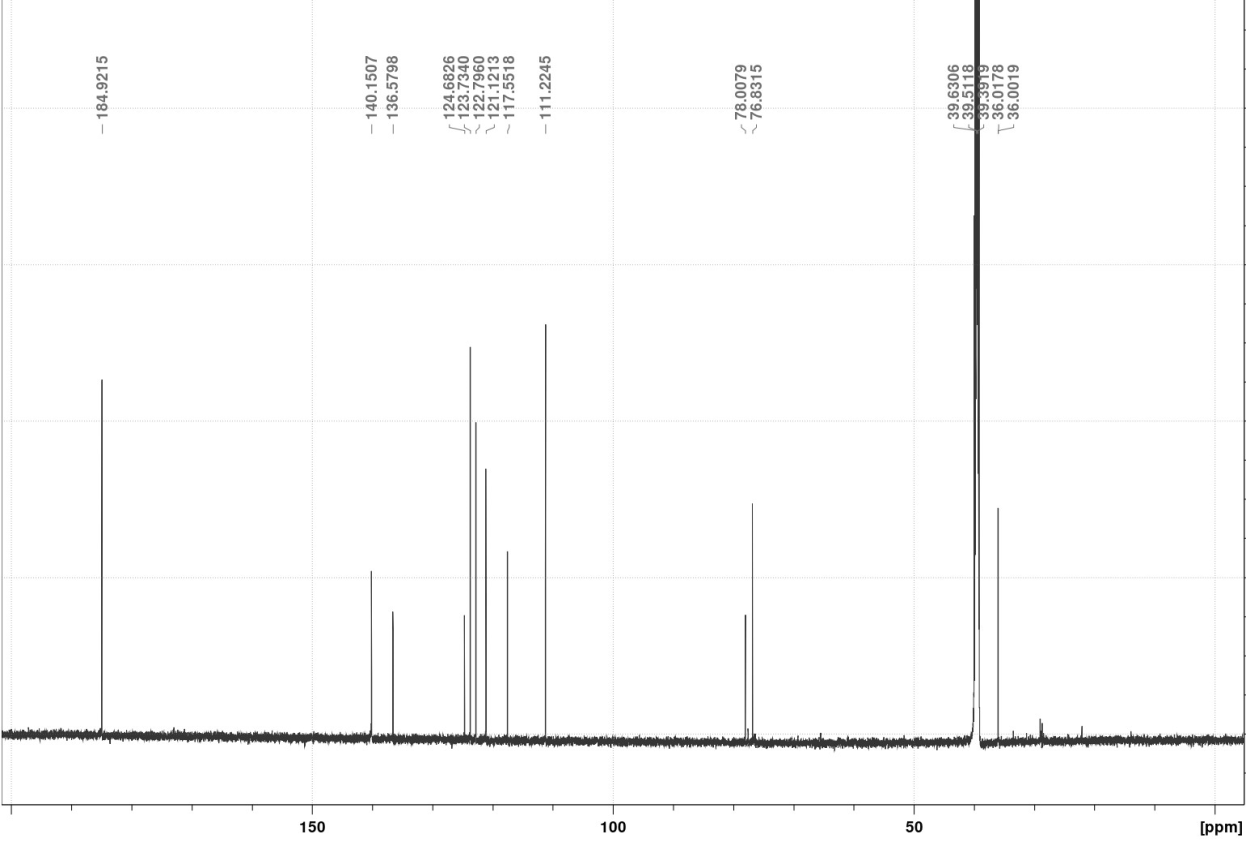

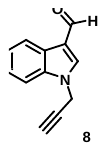

**Figure S.30:** ^13^C NMR spectrum of **8.**

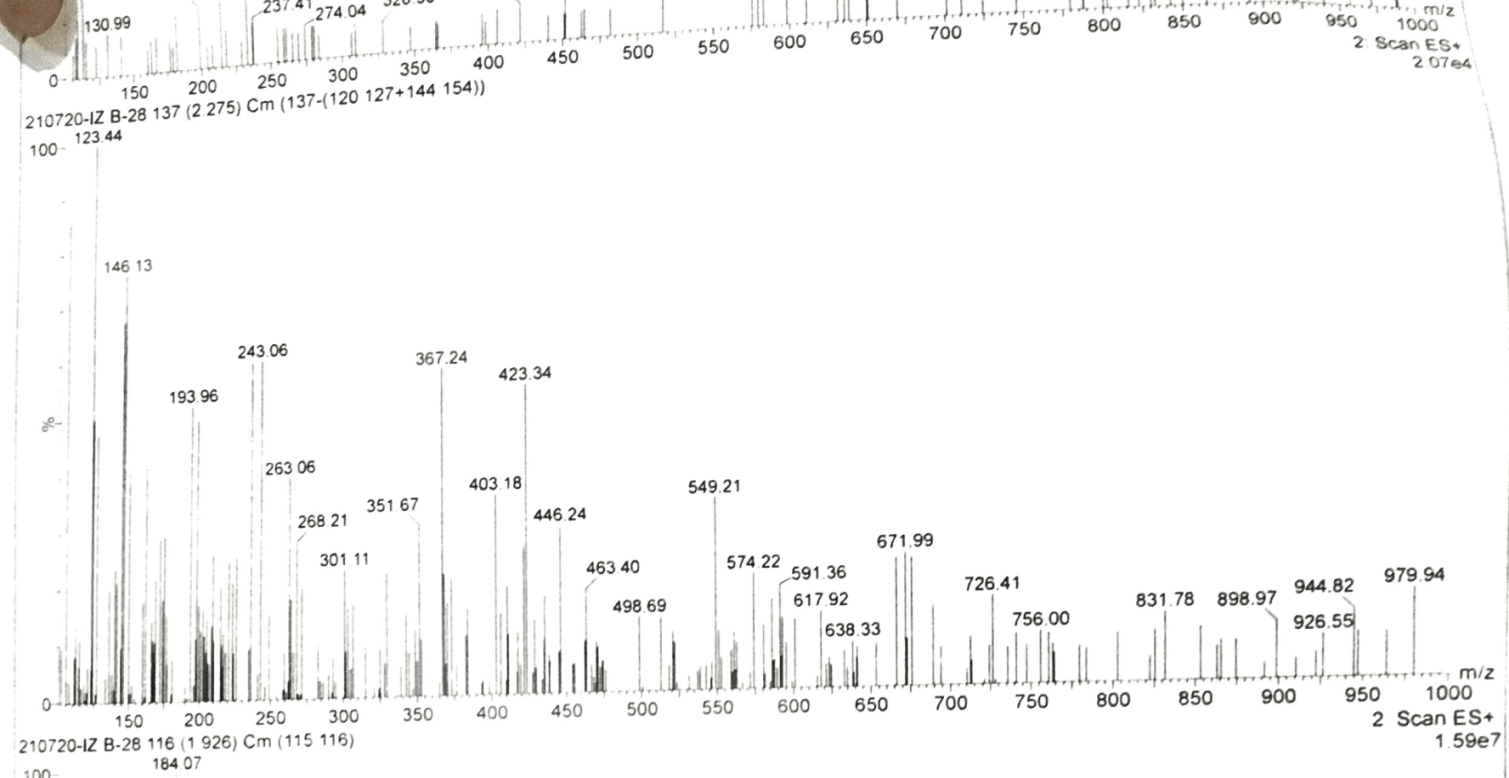

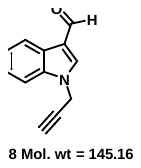

**Figure S.31:** Mass spectrum of **8.**

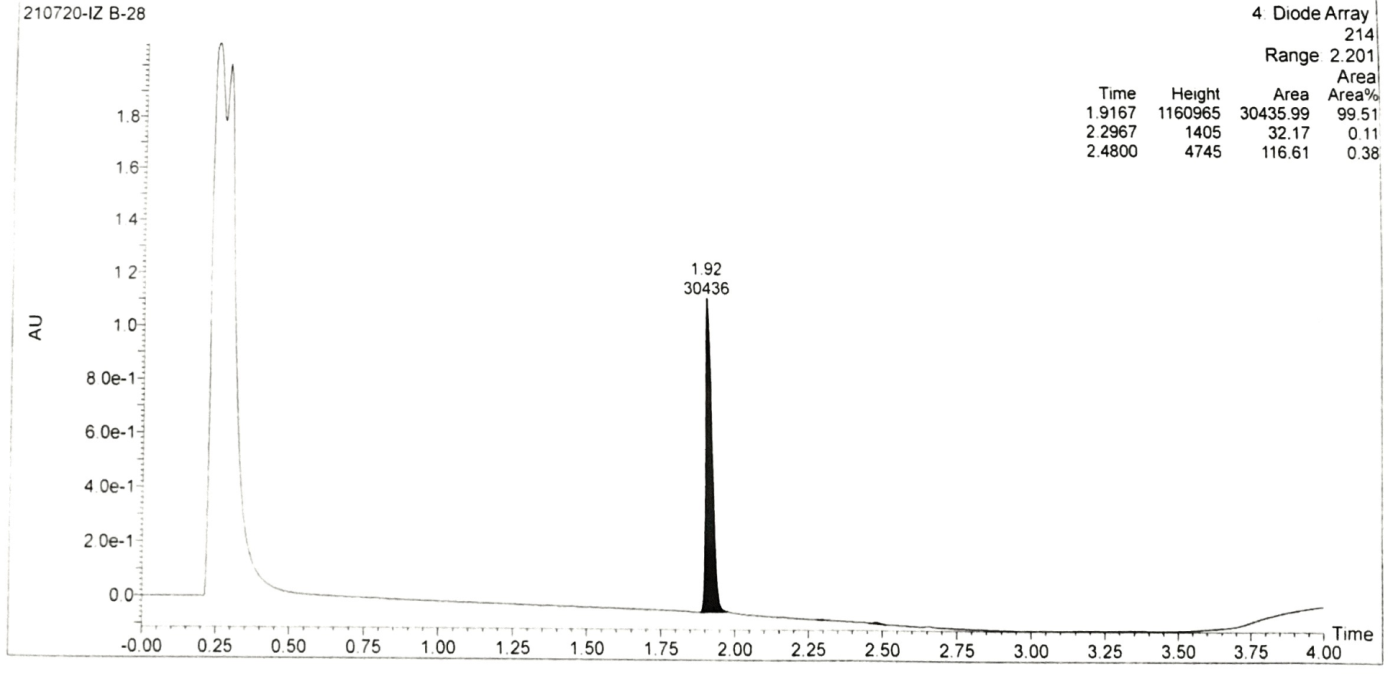

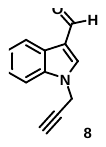

**Figure S.32:** Mass spectrum of **8.**

**9.** (*E*)-3-(1-((1-(7-chloroquinolin-4-yl)-1H-1,2,3-triazol-4-yl)methyl)-1H-indol-3-yl)-1-phenylprop-2-en-1-one **(7a)**:

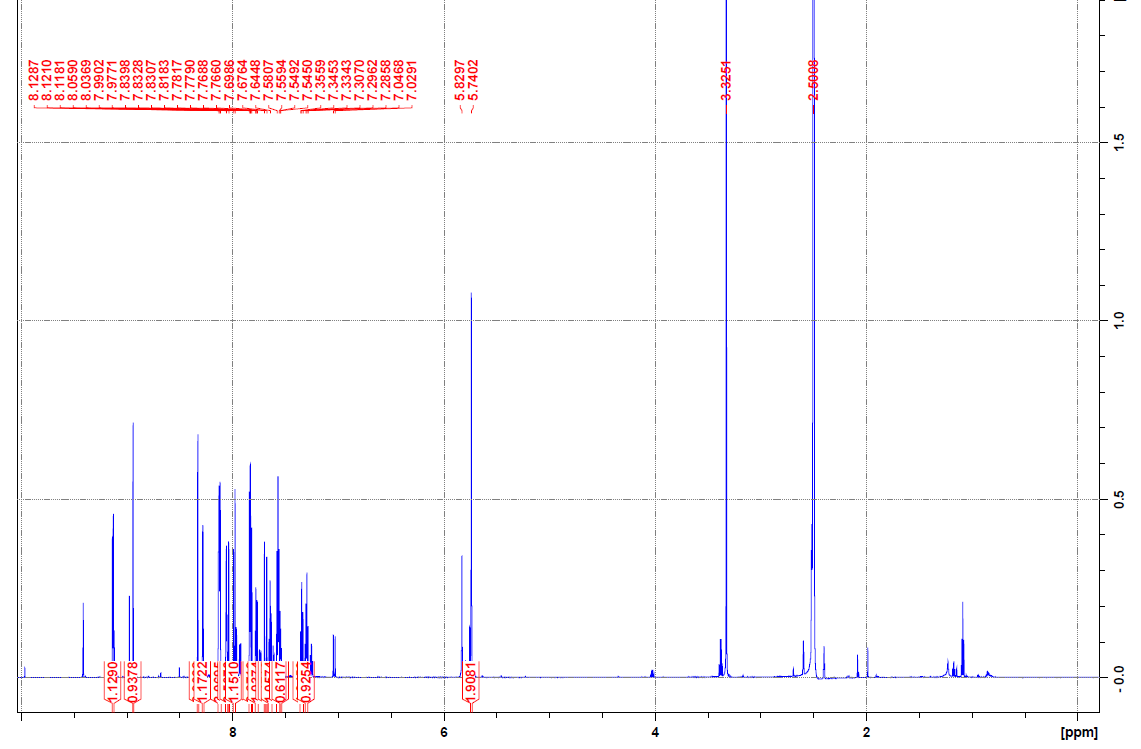

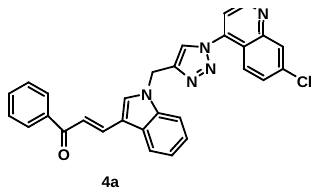

**Figure S.33A:** ^1^H NMR spectrum of **7a.**

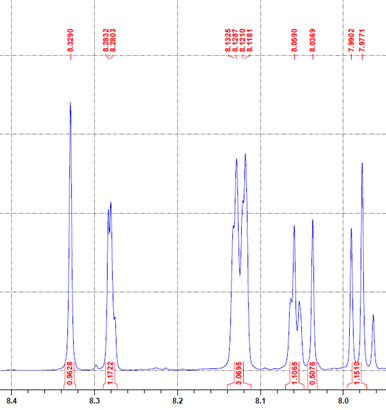

**Figure S.33B:** ^1^H NMR spectrum of **7a.**

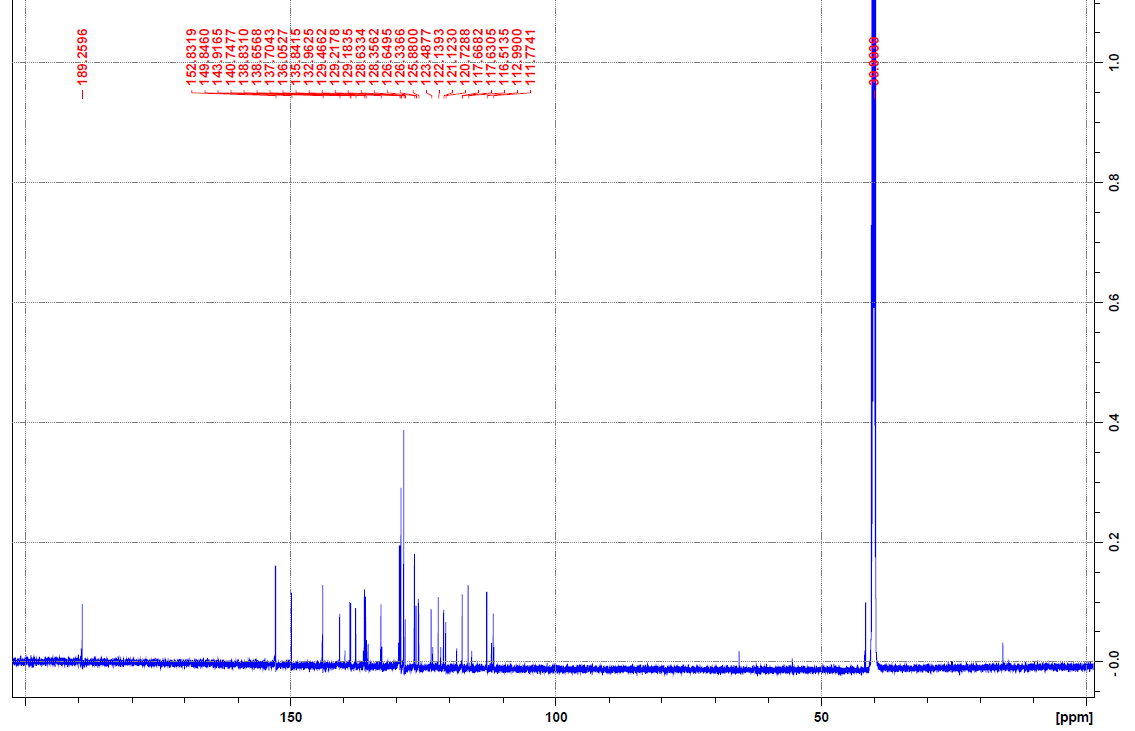

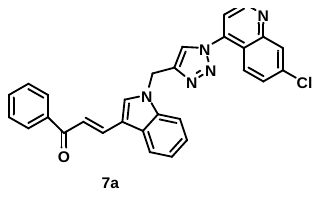

**Figure S.34:** ^13^C NMR spectrum of **7a.**

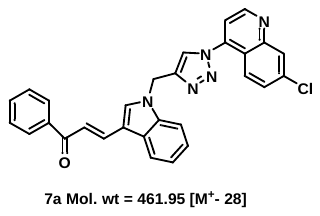

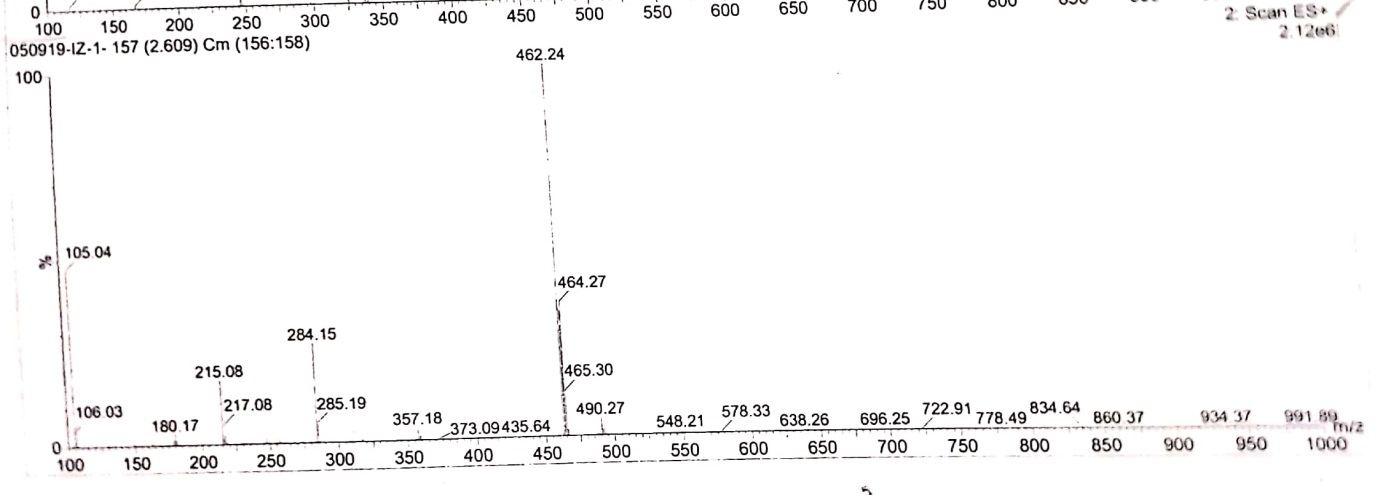

**Figure S.35:** Mass spectrum of **7a.**

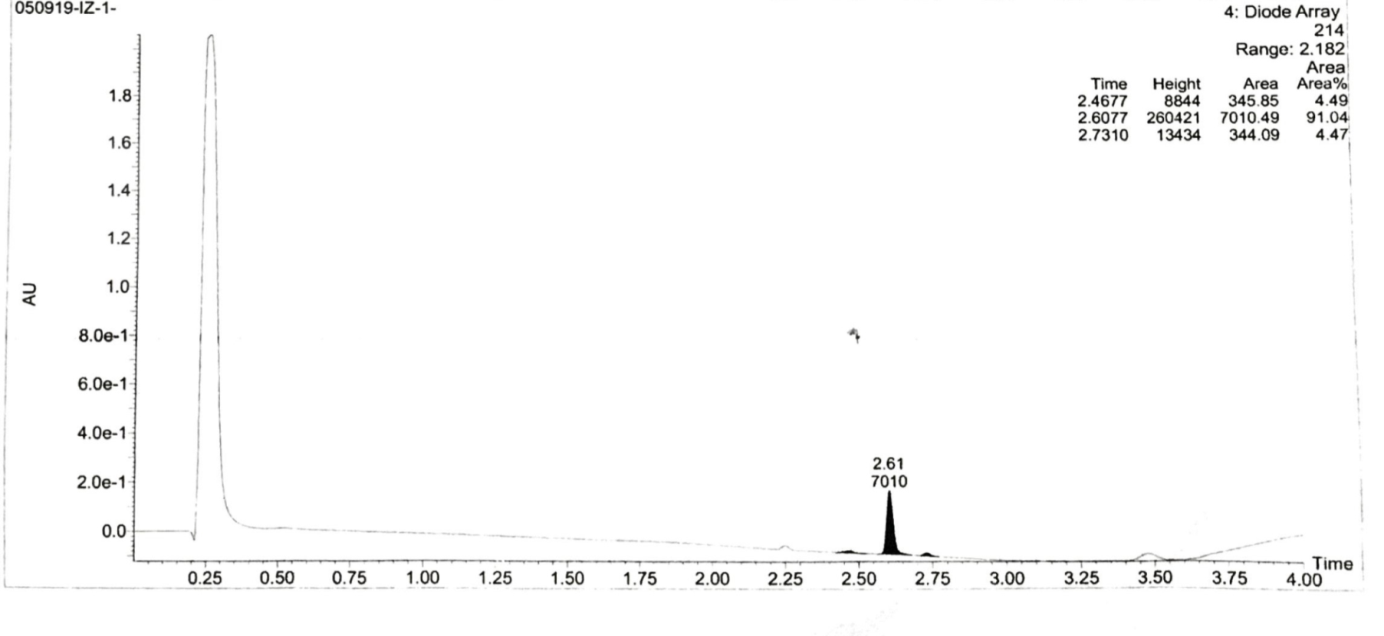

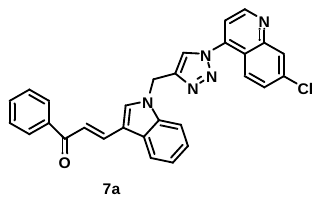

**Figure S.36:** Purity of **7a.**

**10.** (*E*)-3-(1-((1-(7-chloroquinolin-4-yl)-1H-1,2,3-triazol-4-yl)methyl)-1H-indol-3-yl)-1-(2-nitrophenyl)prop-2-en-1-one **(7b):**

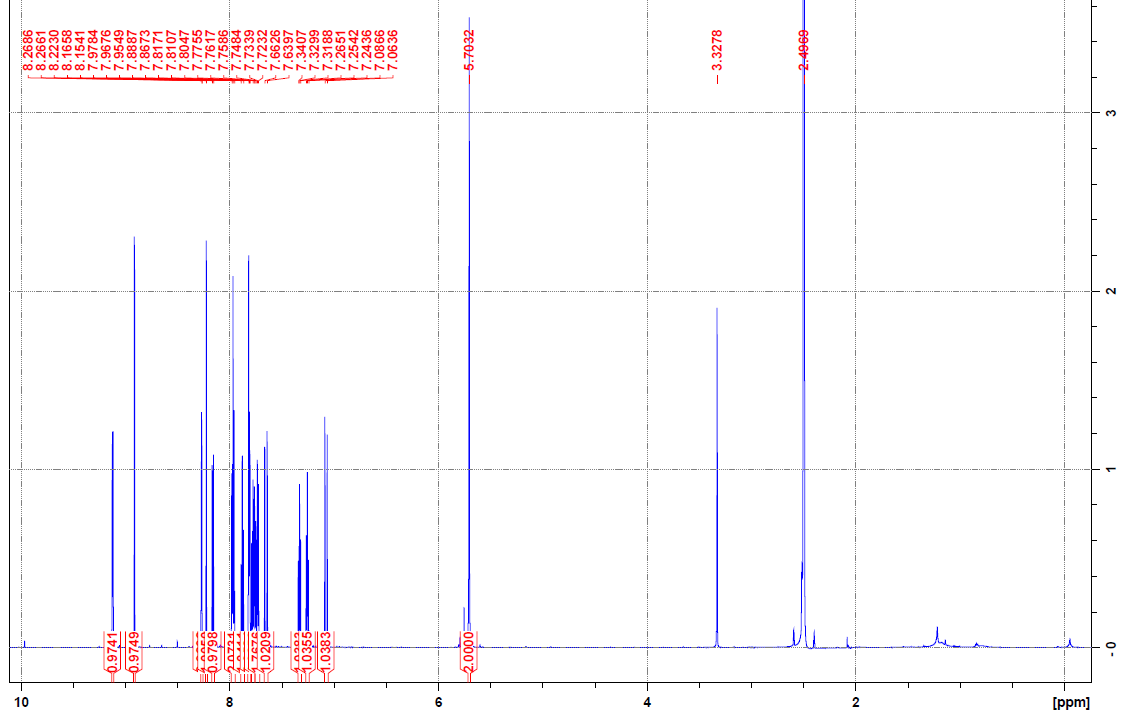

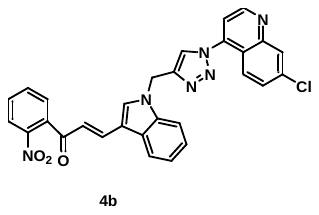

**Figure S.37A:** ^1^H NMR spectrum of **7b.**

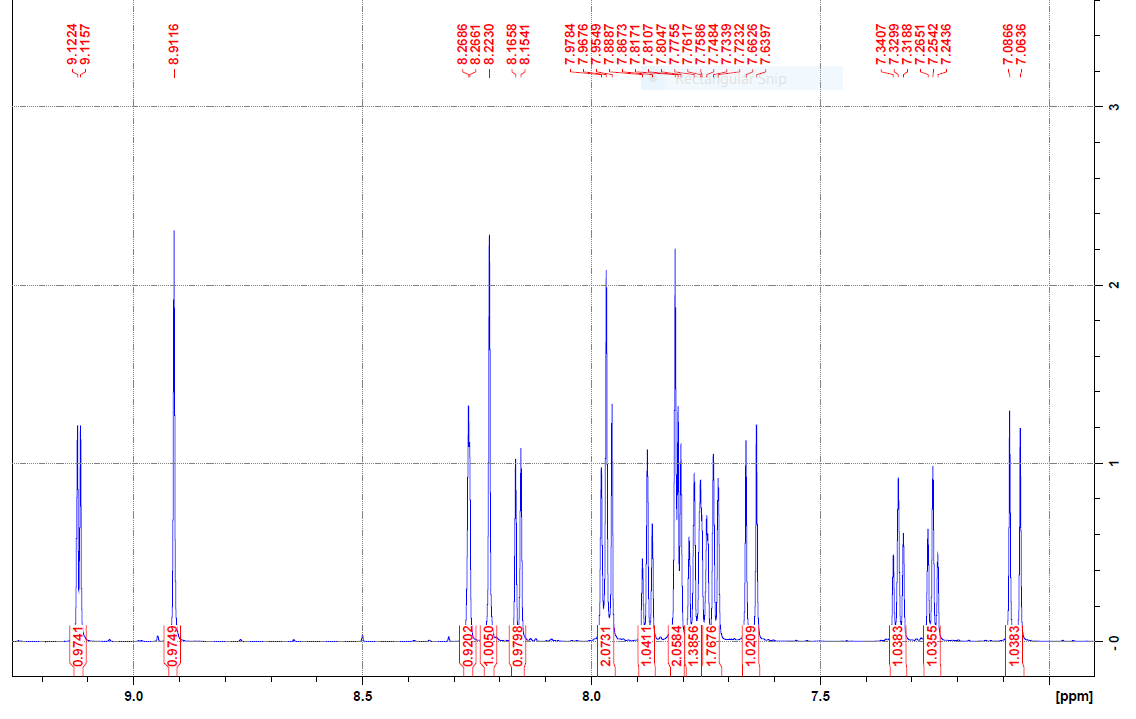

**Figure S.37B:** ^1^H NMR spectrum of **7b.**

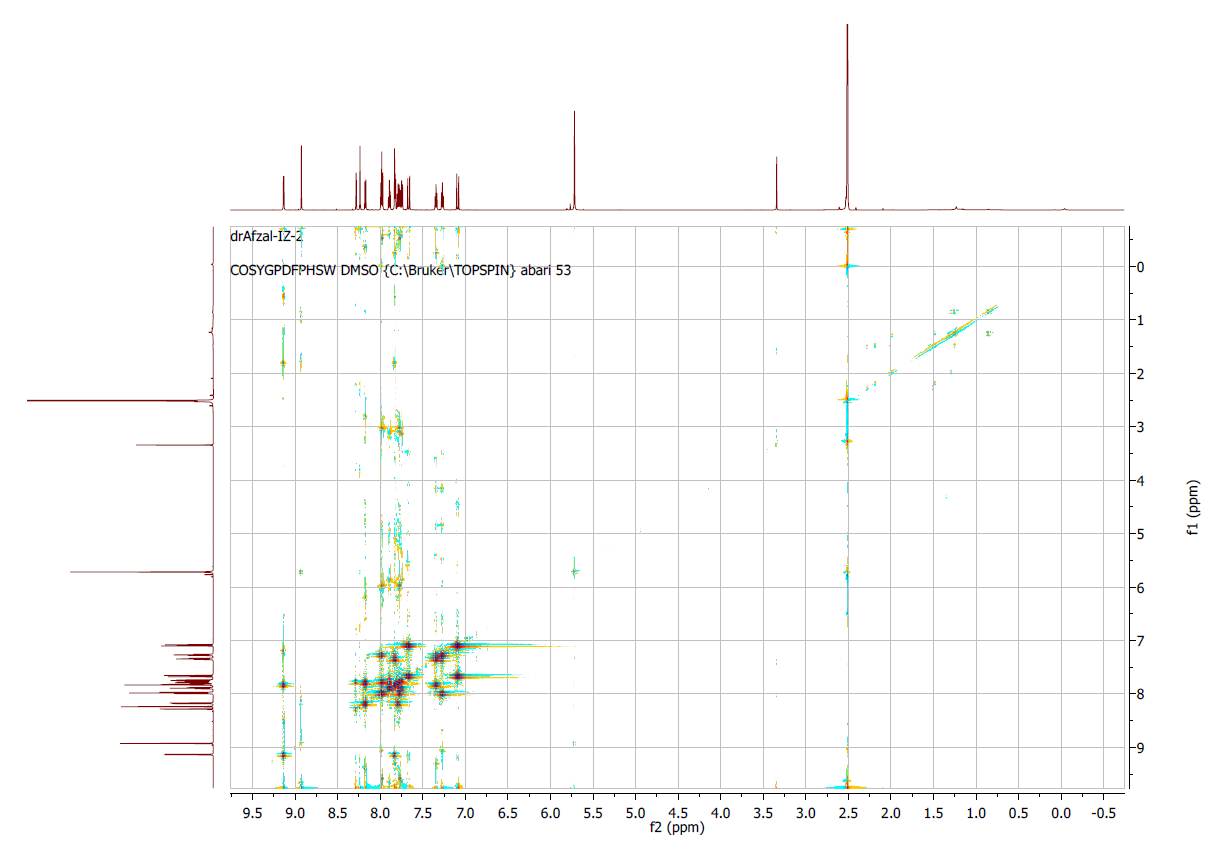

**Figure S.38:** 2D-COSY analyses of **7b.**

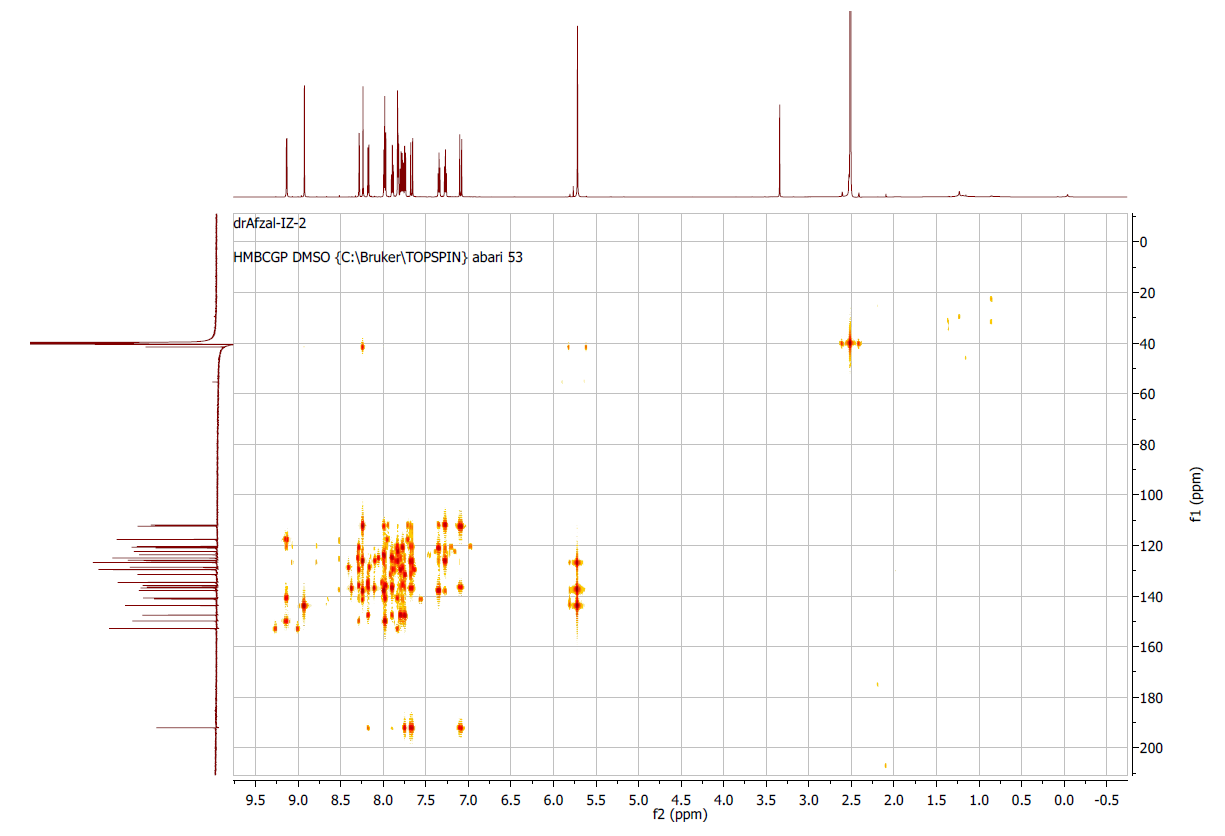

**Figure S.39:** 2D-HMBC analyses of **7b.**

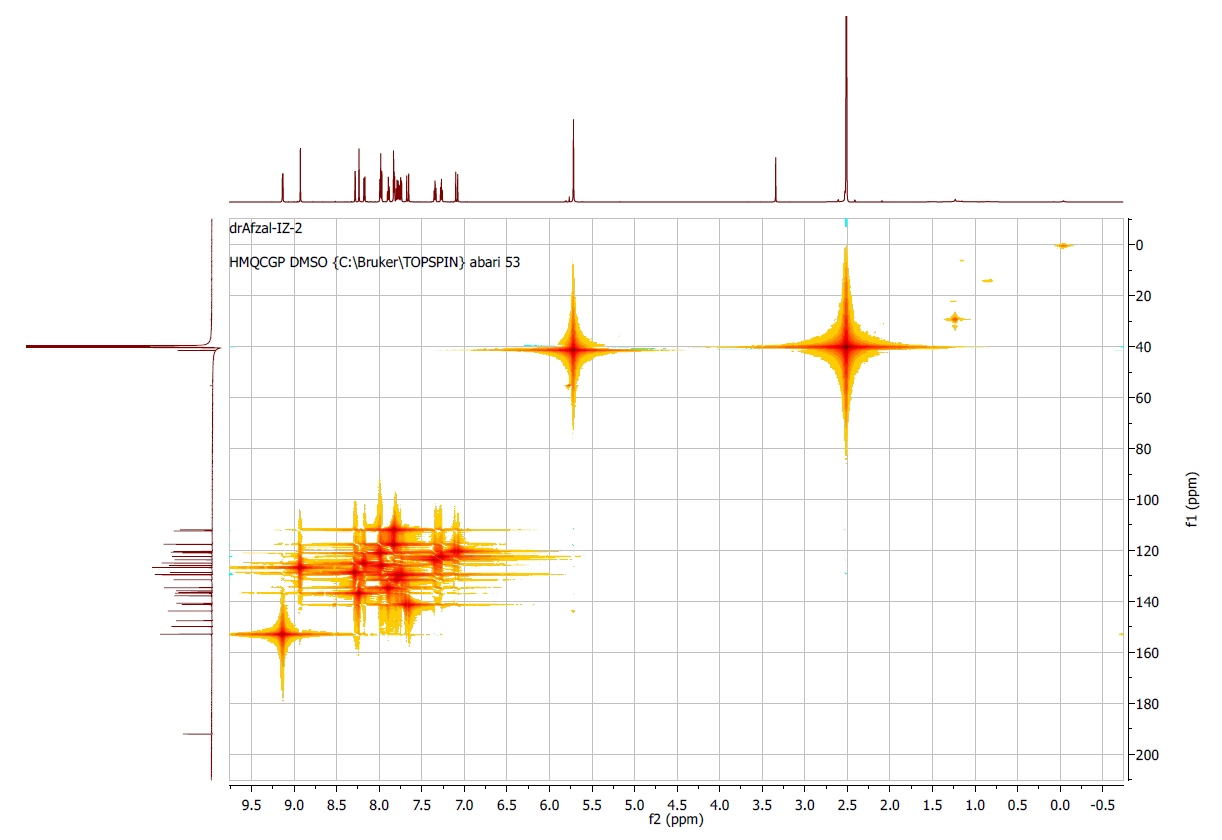

**Figure S.40:** 2D-HMQC analyses of **7b.**

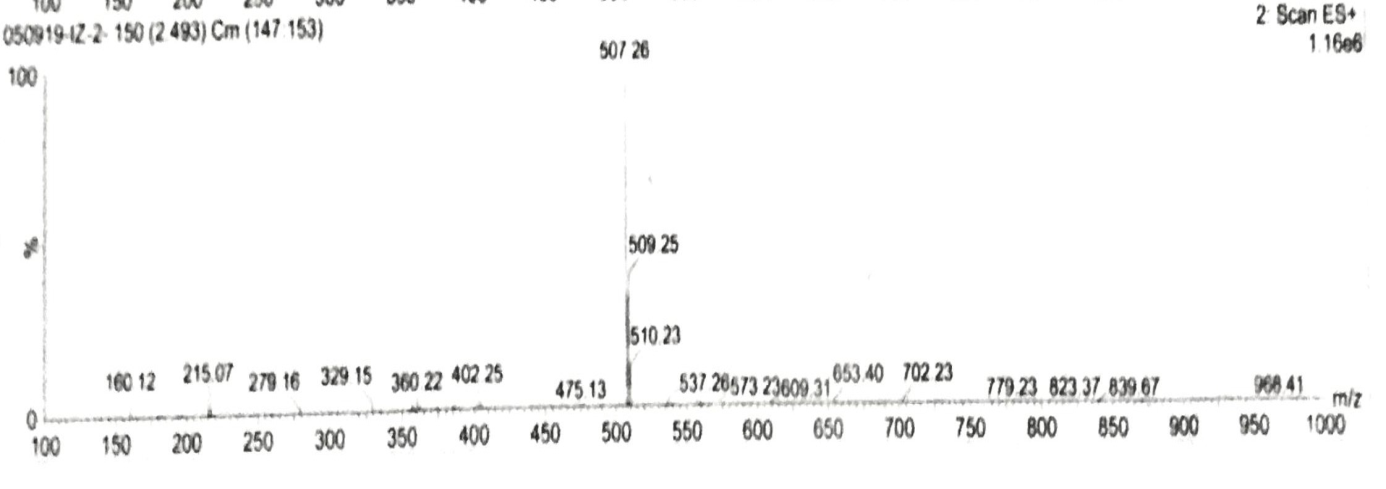

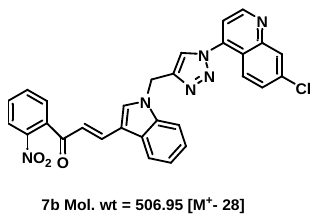

**Figure S.41:** Mass spectrum of **7b.**

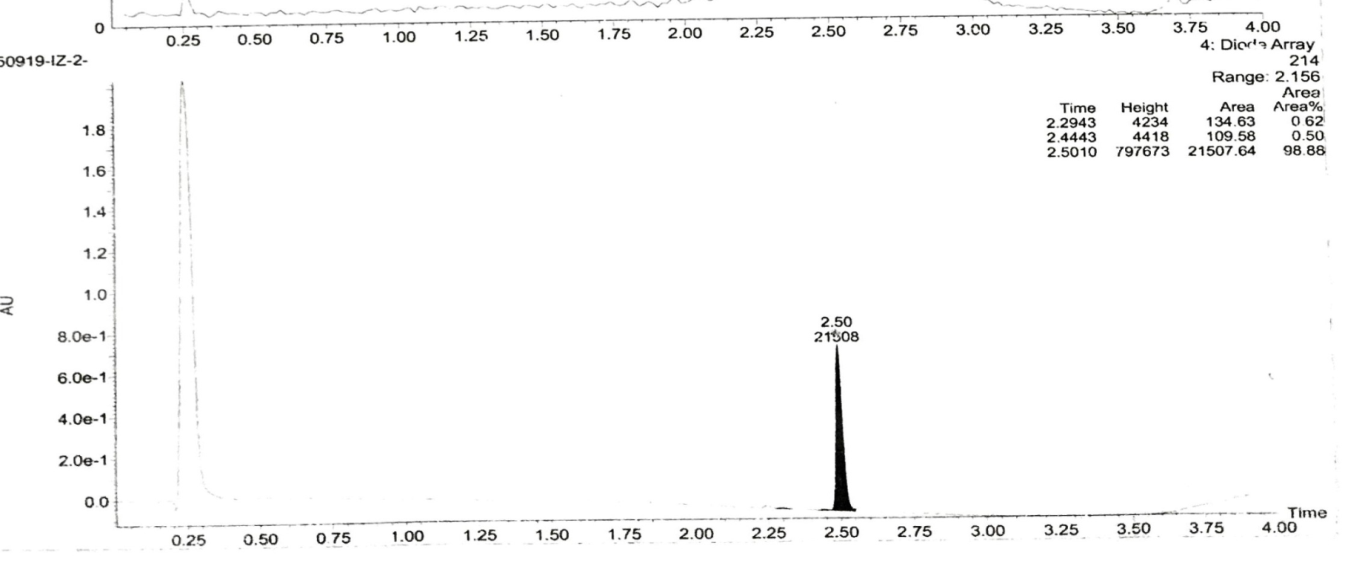

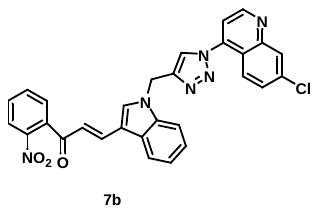

**Figure S.42:** Purity of **7b.**

**11.** (*E*)-3-(1-((1-(7-chloroquinolin-4-yl)-1H-1,2,3-triazol-4-yl)methyl)-1H-indol-3-yl)-1-(3-nitrophenyl)prop-2-en-1-one **(7c):**

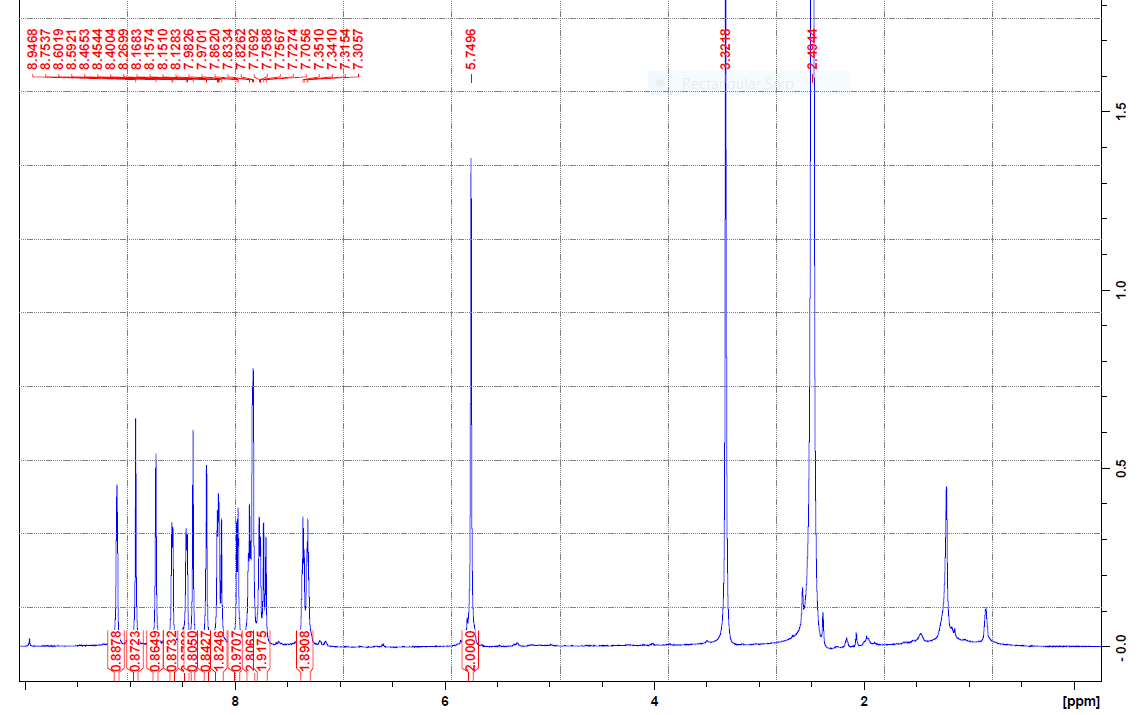

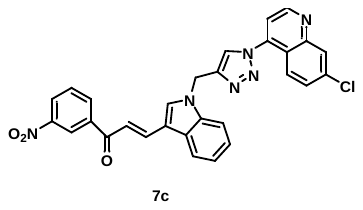

**Figure S. 43A:** ^1^H NMR spectrum of **7c.**

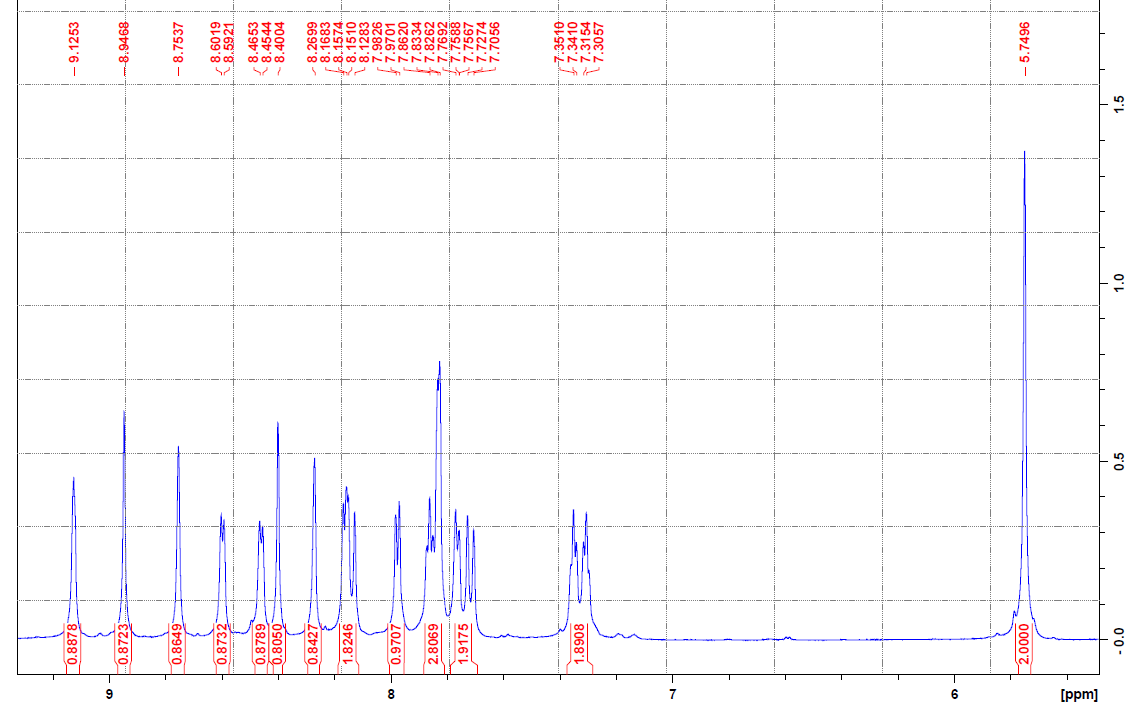

**Figure S. 43B:** ^1^H NMR spectrum of **7c.**

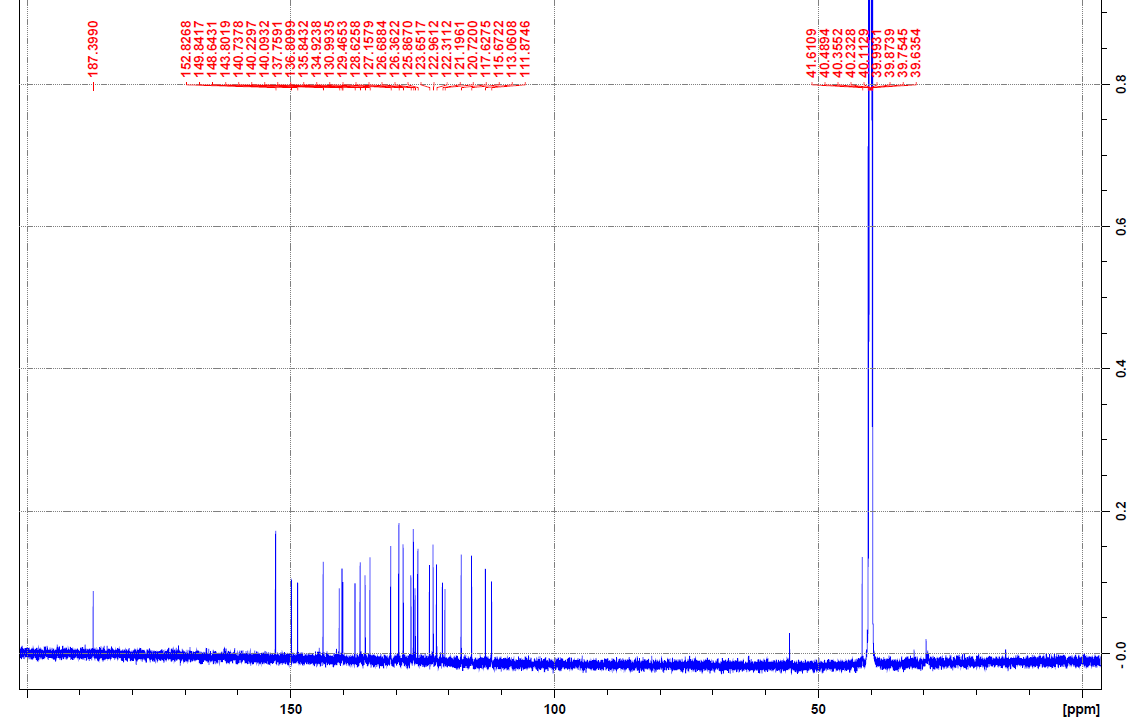

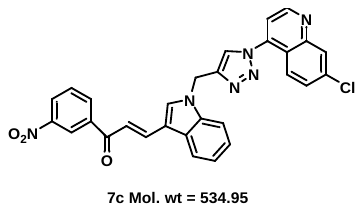

**Figure S.44:** ^13^C NMR spectrum of **7c.**

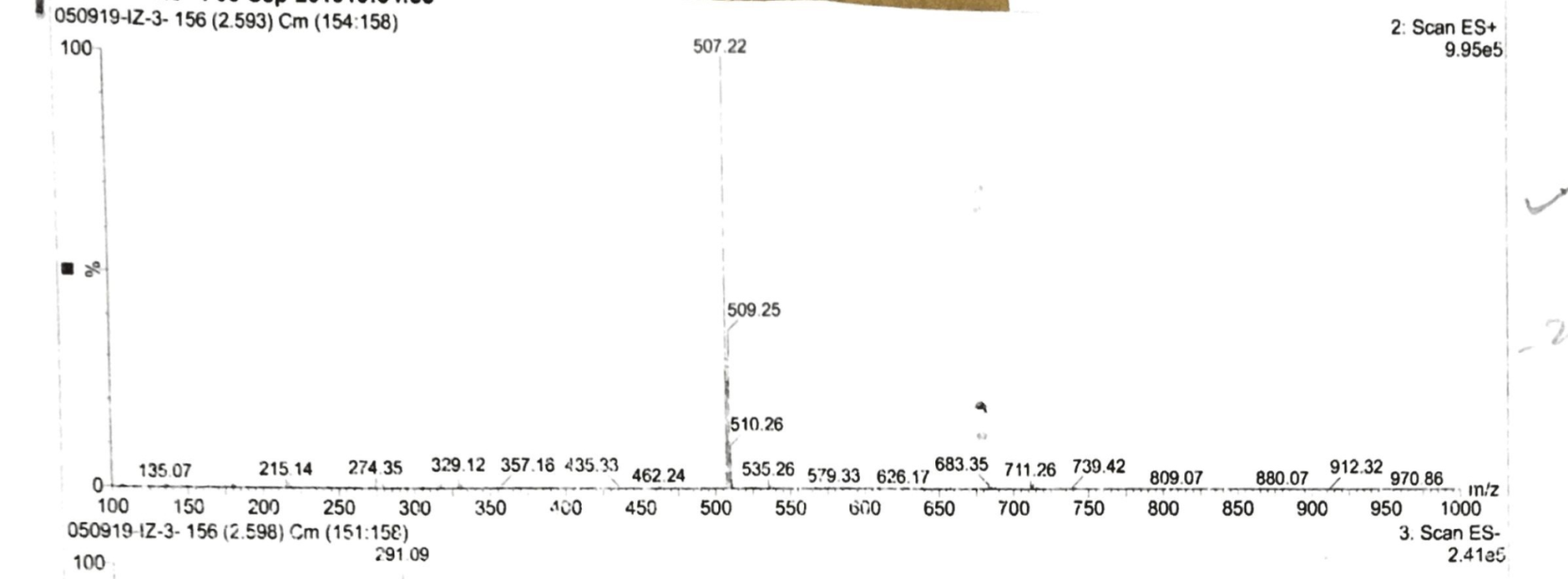

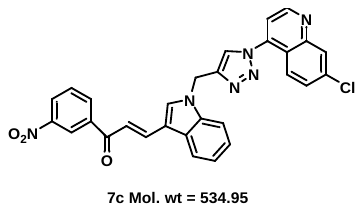

**Figure S.45:** Mass spectrum of **7c.**

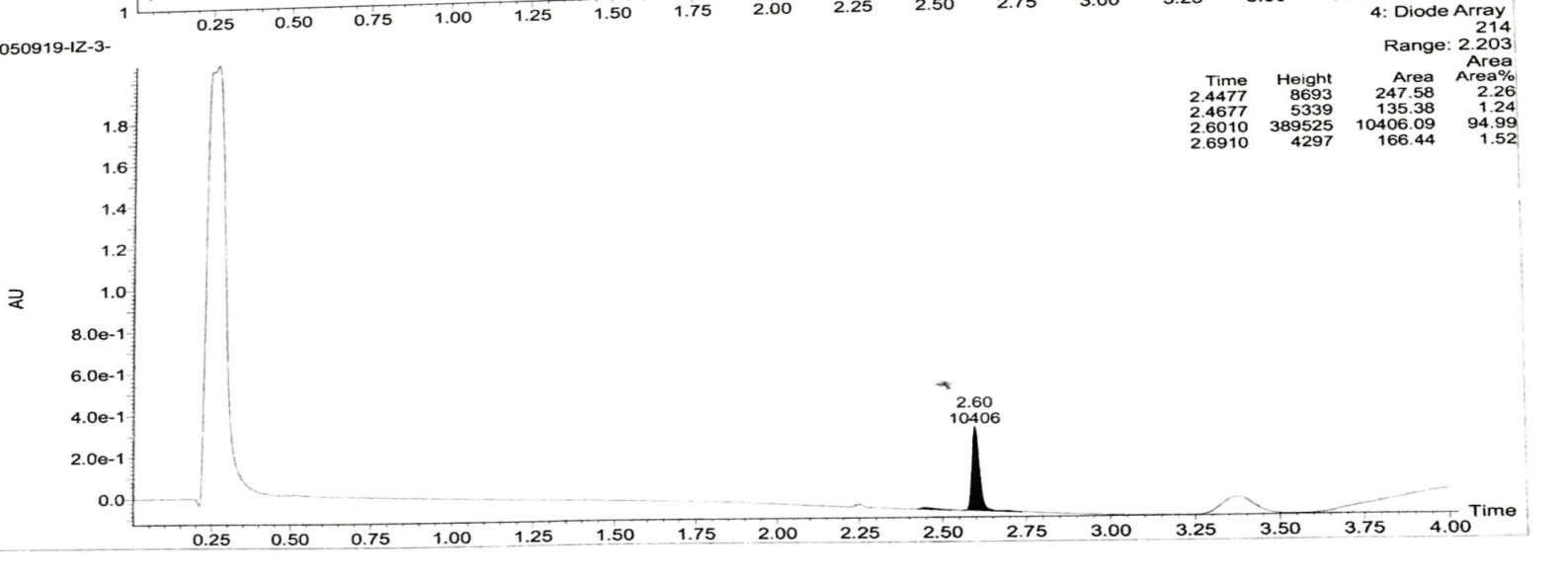

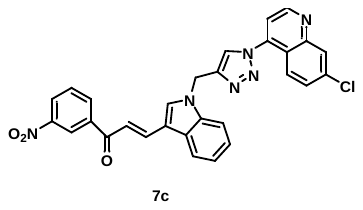

**Figure S.46:** purity of **7c.**

**12.** (*E*)-3-(1-((1-(7-chloroquinolin-4-yl)-1H-1,2,3-triazol-4-yl)methyl)-1H-indol-3-yl)-1-(4-nitrophenyl)prop-2-en-1-one **(7d):**

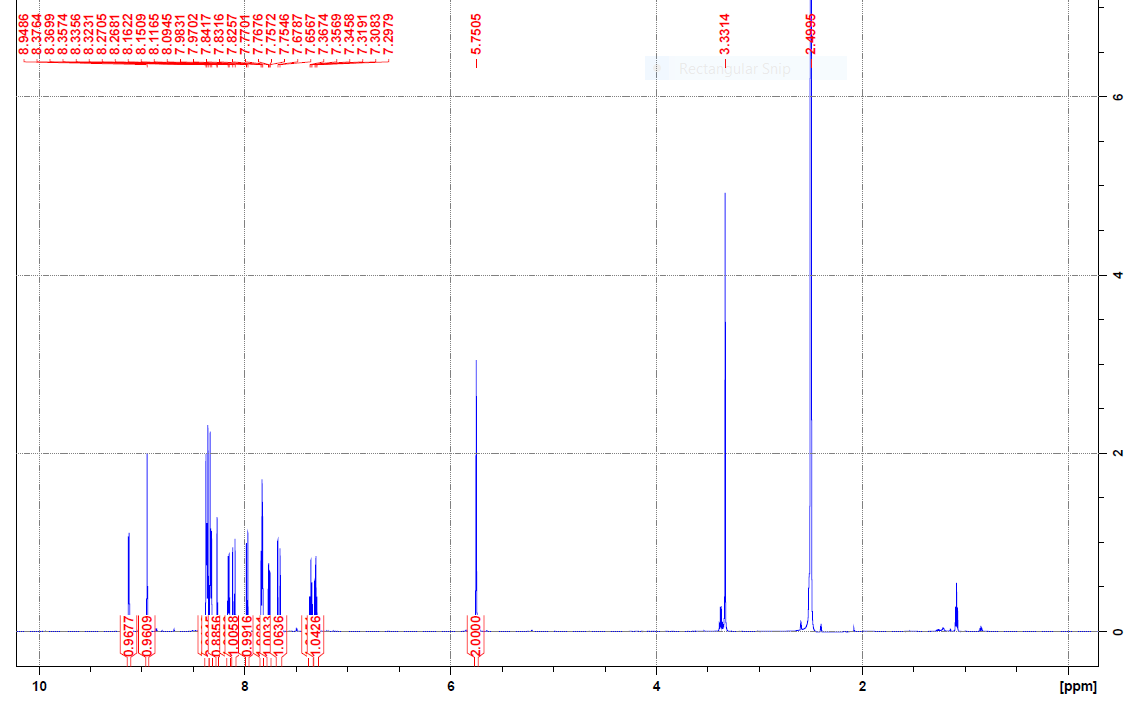

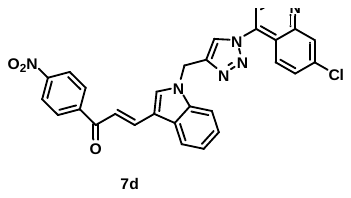

**Figure S.47A:** ^1^H NMR spectrum of **7d.**

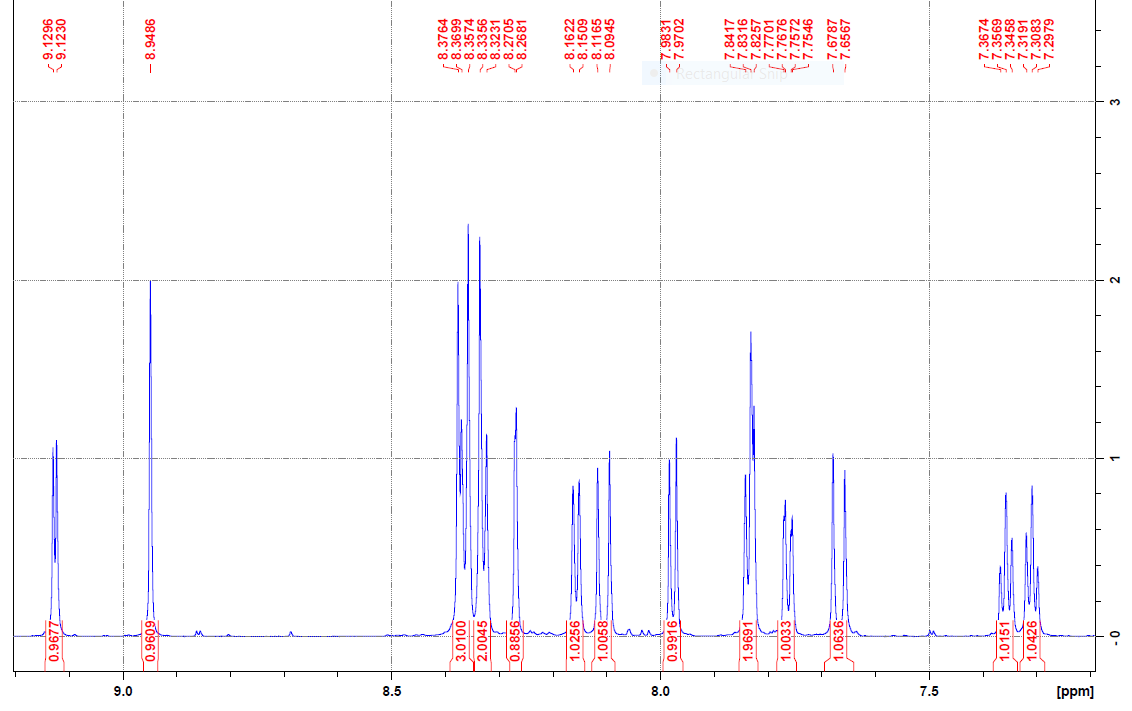

**Figure S.47B:** ^1^H NMR spectrum of **7d.**

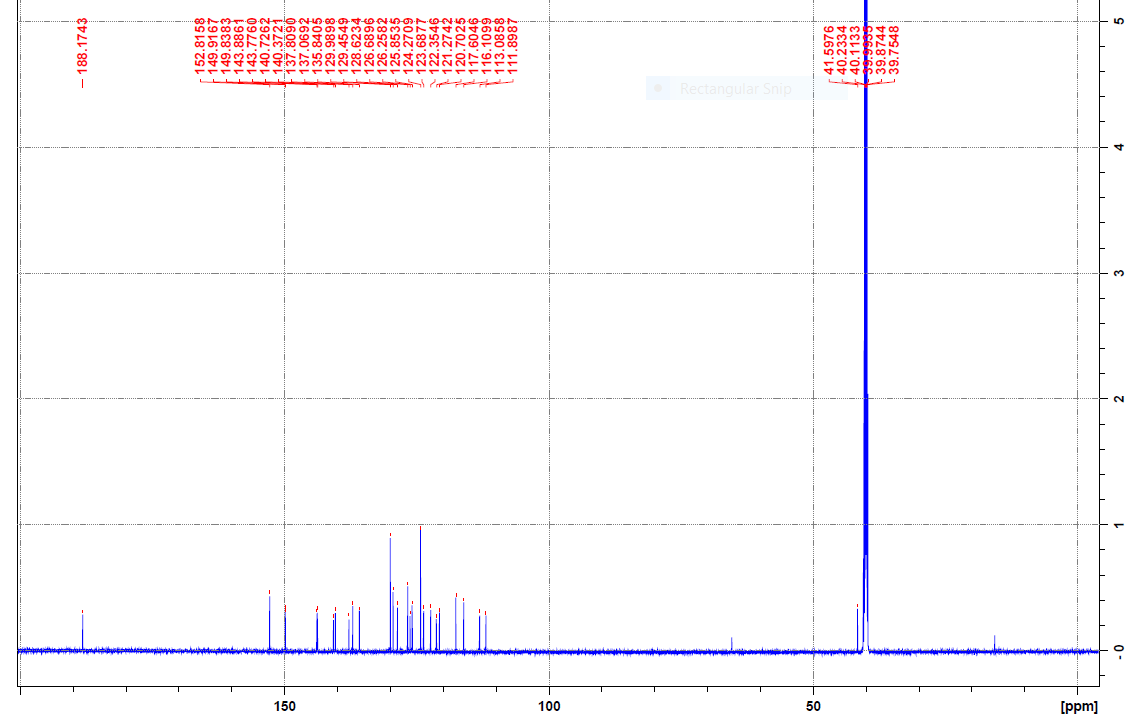

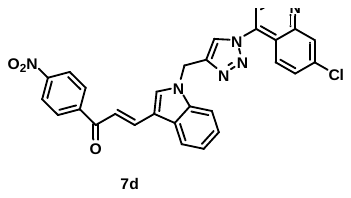

**Figure S.48:** ^13^C NMR spectrum of **7d.**

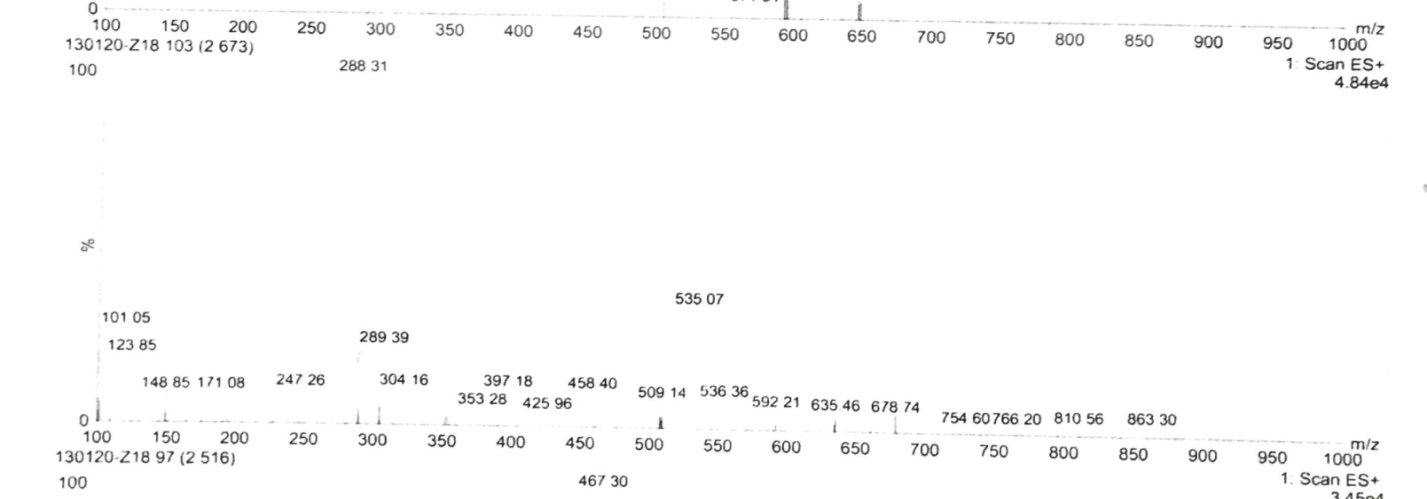

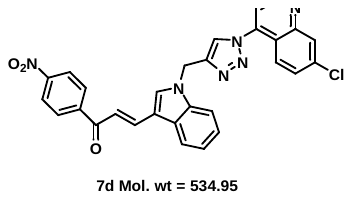

**Figure S.49:** Mass spectrum of **7d.**

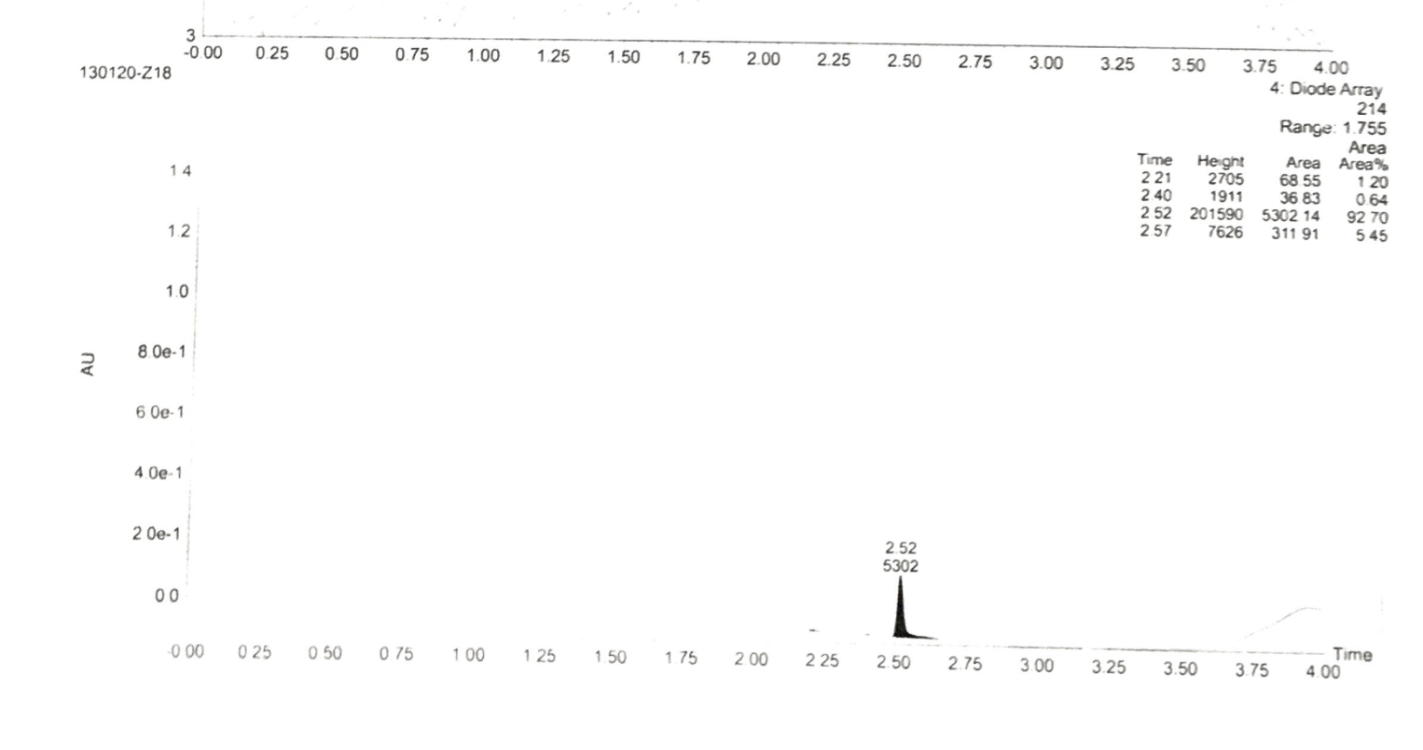

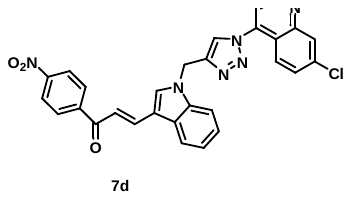

**Figure S.50:** Purity of **7d.**

**13.** (*E*)-1-(2-bromophenyl)-3-(1-((1-(7-chloroquinolin-4-yl)-1H-1,2,3-triazol-4-yl)methyl)-1H-indol-3-yl)prop-2-en-1-one **(7e):**

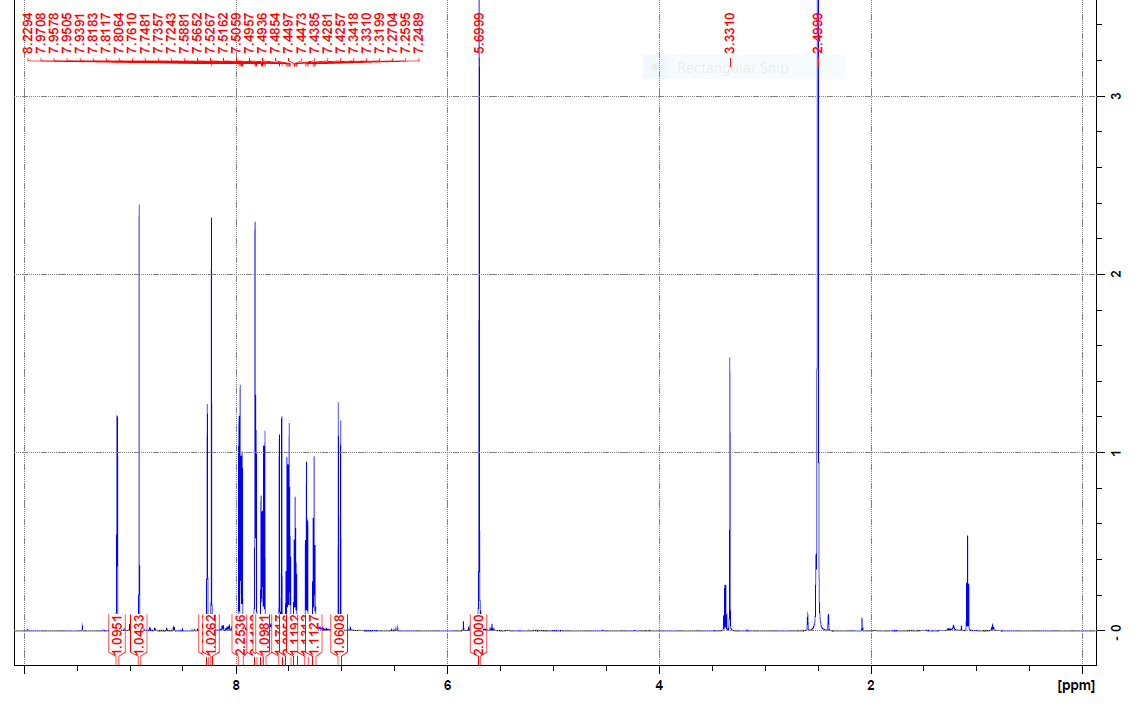

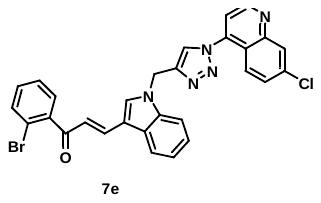

**Figure S.51A:** ^1^H NMR spectrum of **7e.**

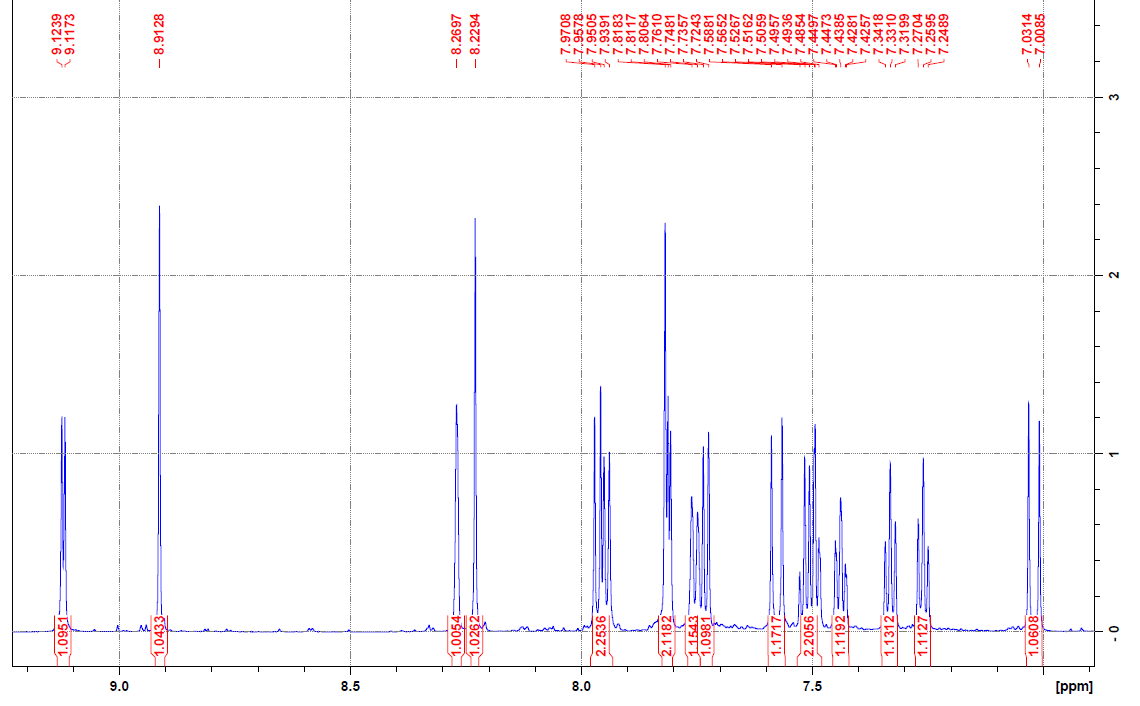

**Figure S.51B:** ^1^H NMR spectrum of **7e.**

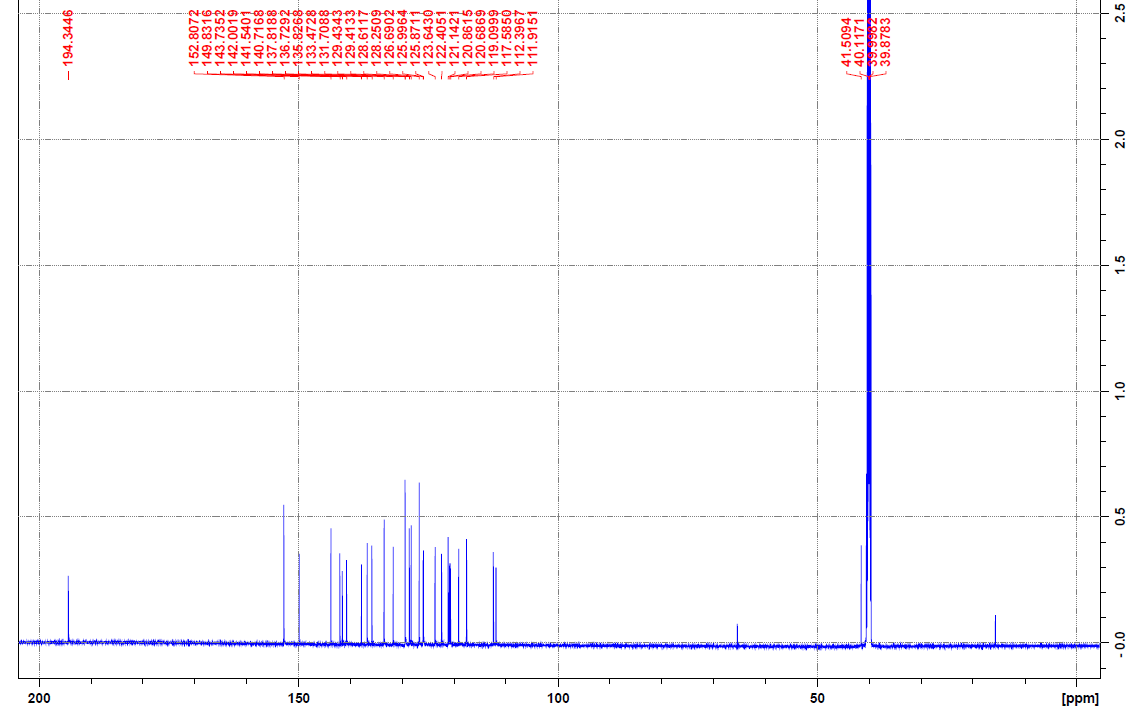

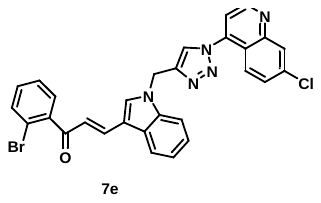

**Figure S.52:** ^13^C NMR spectrum of **7e.**

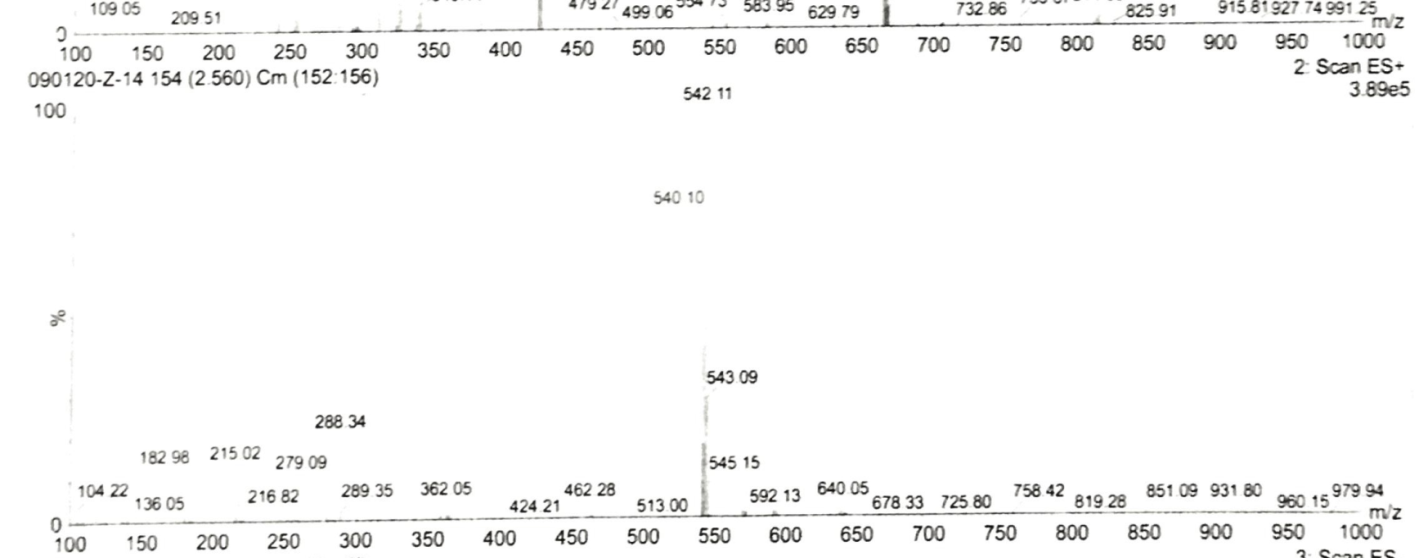

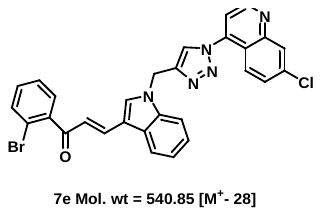

**Figure S.53:** Mass spectrum of **7e.**

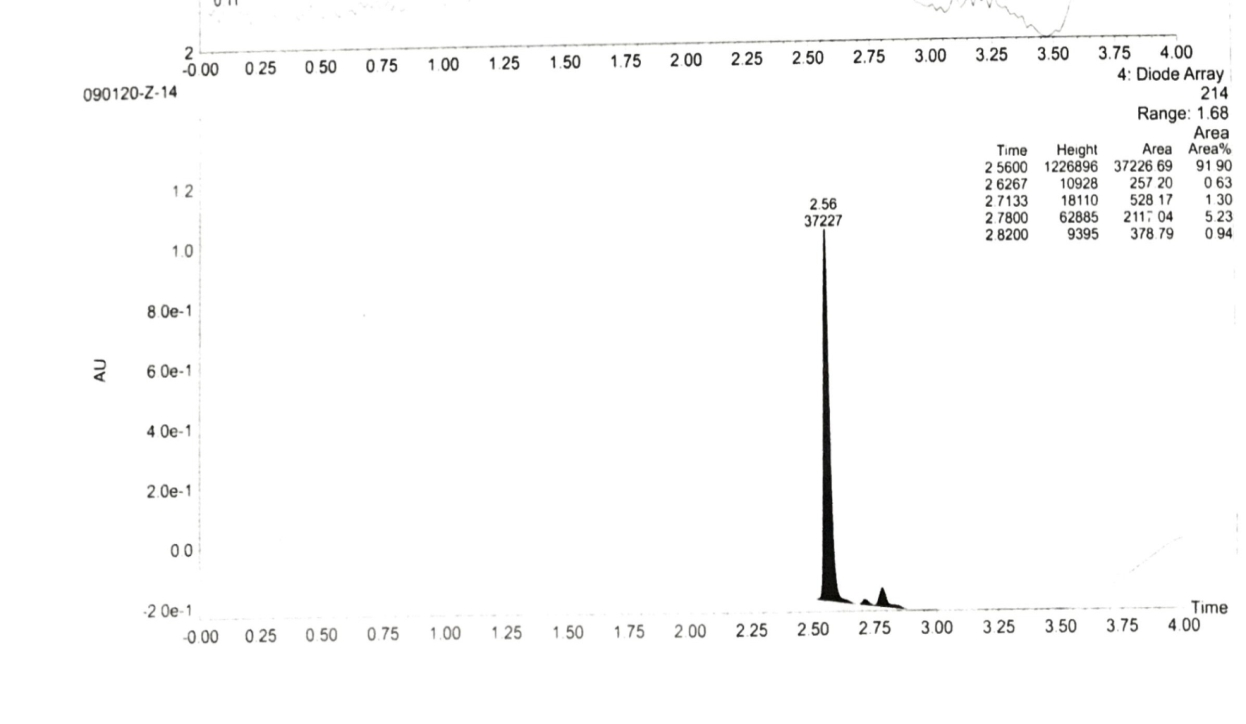

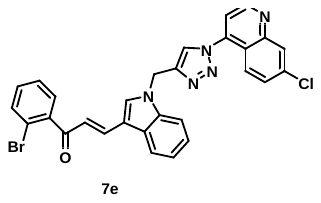

**Figure S.54:** Purity spectrum of **7e.**

**14.** (*E*)-1-(3-bromophenyl)-3-(1-((1-(7-chloroquinolin-4-yl)-1H-1,2,3-triazol-4-yl)methyl)-1H-indol-3-yl)prop-2-en-1-one **(7f):**

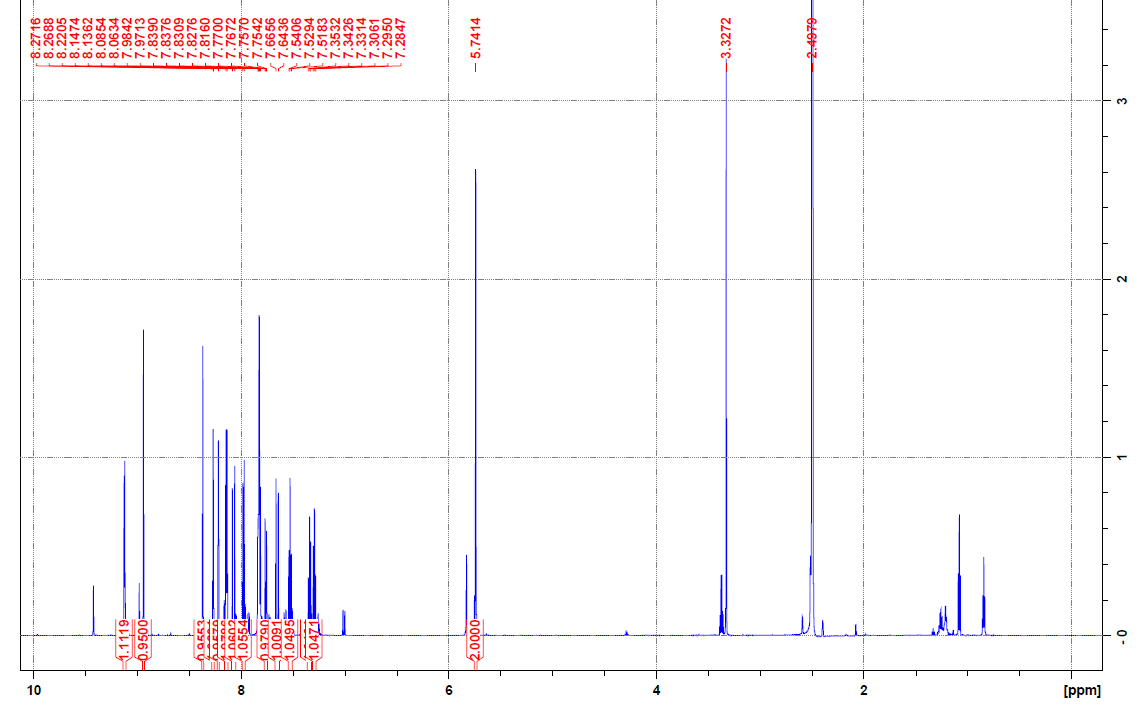

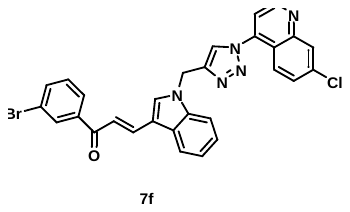

**Figure S.55A:** ^1^H NMR spectrum of **7f.**

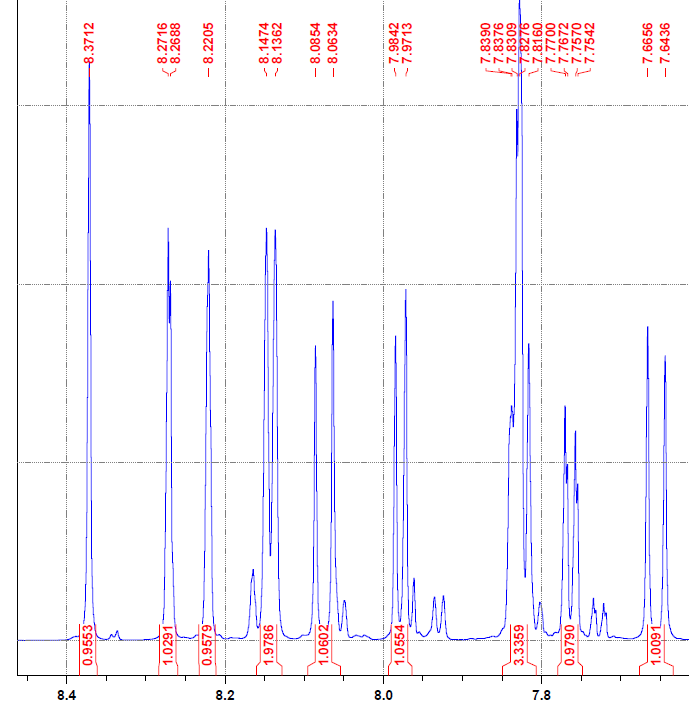

**Figure S.55B:** ^1^H NMR spectrum of **7f.**

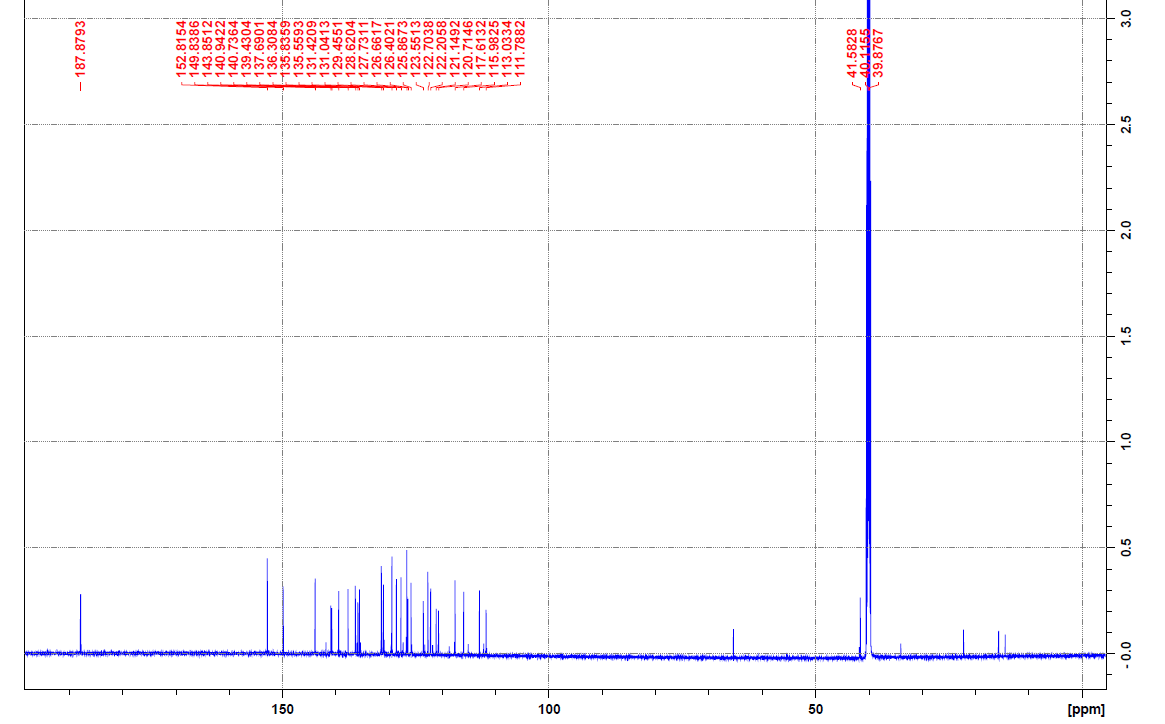

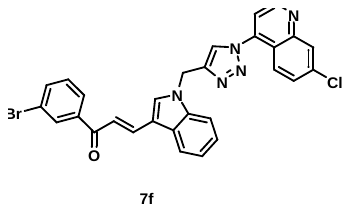

**Figure S.56:** ^13^C NMR spectrum of **7f.**

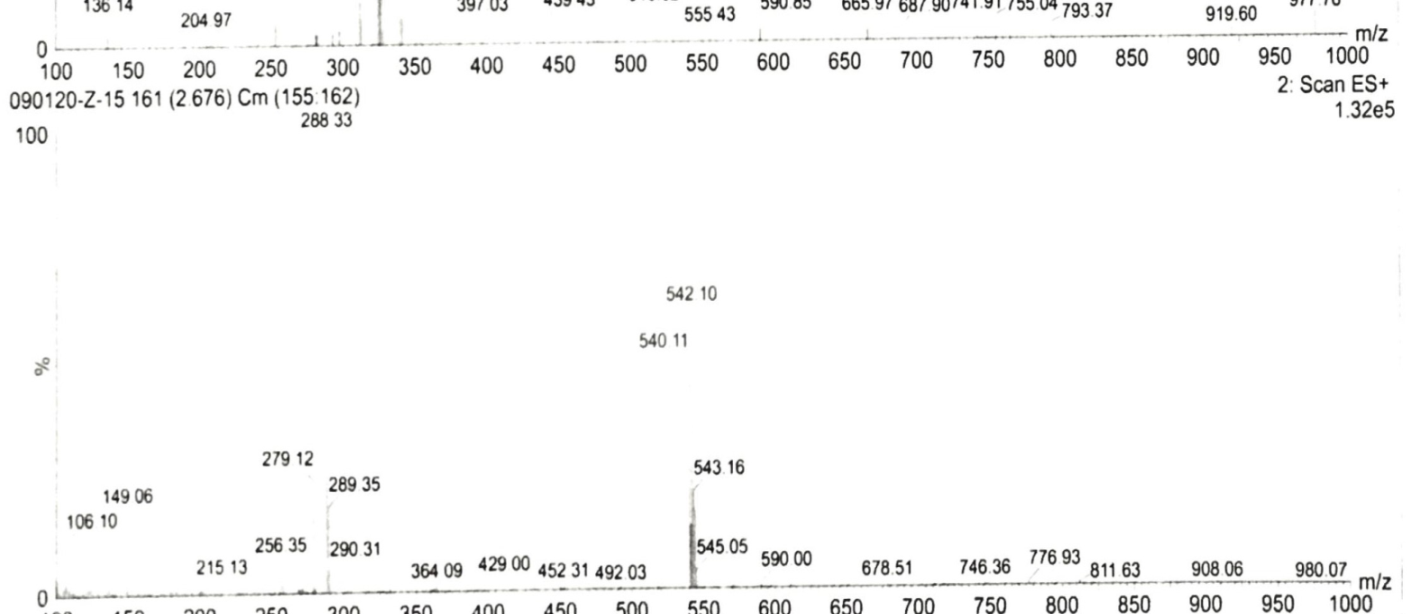

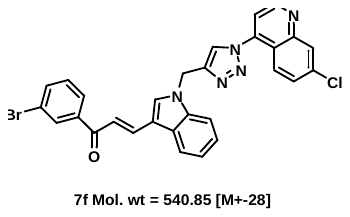

**Figure S.57:** Mass spectrum of **7f.**

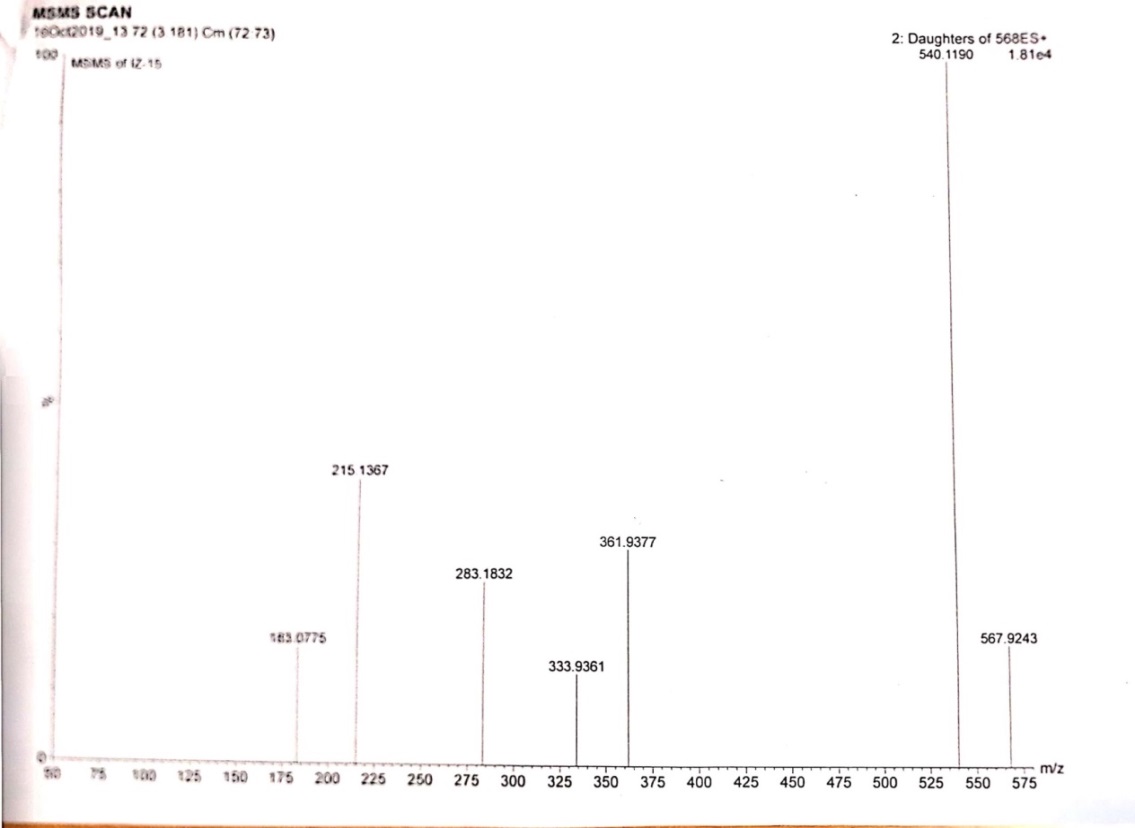

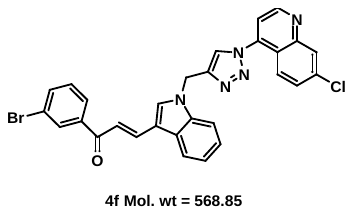

**Figure S.58:** MSMS spectrum of **7f.**

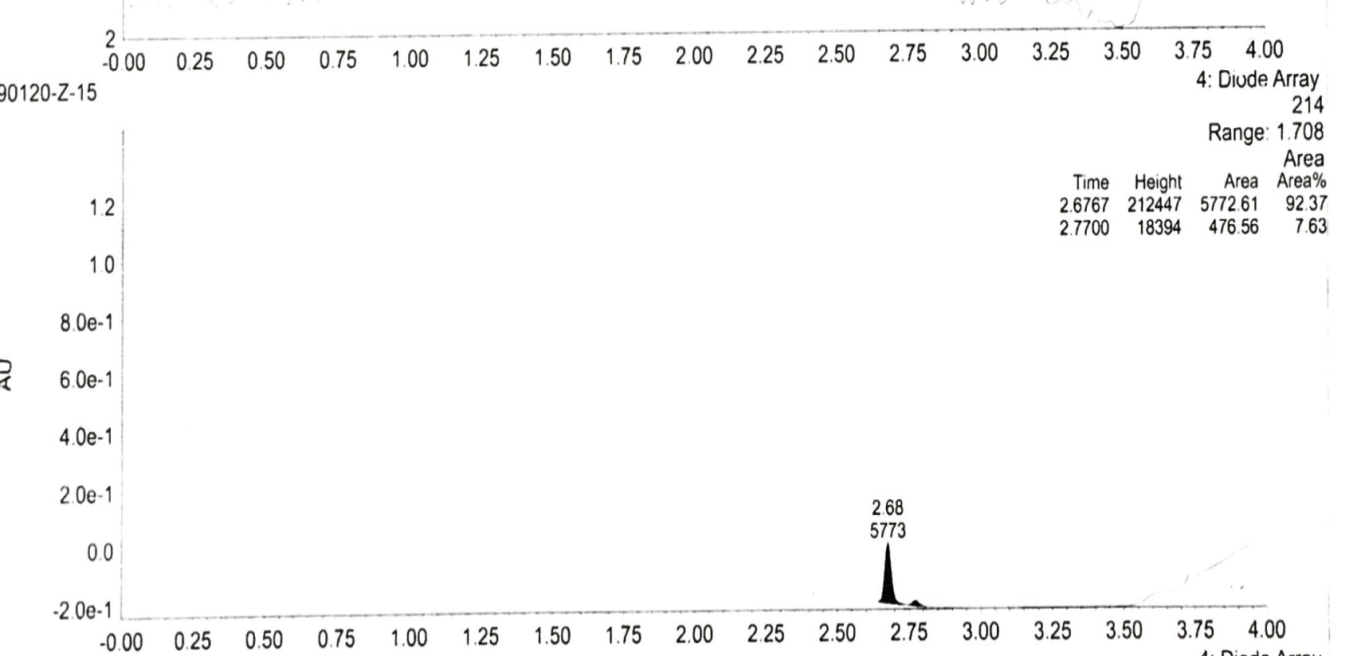

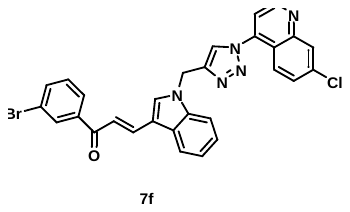

**Figure S.59:** Purity of **7f.**

**15.** (*E*)-1-(2-chlorophenyl)-3-(1-((1-(7-chloroquinolin-4-yl)-1H-1,2,3-triazol-4-yl)methyl)-1H-indol-3-yl)prop-2-en-1-one **(7h):**

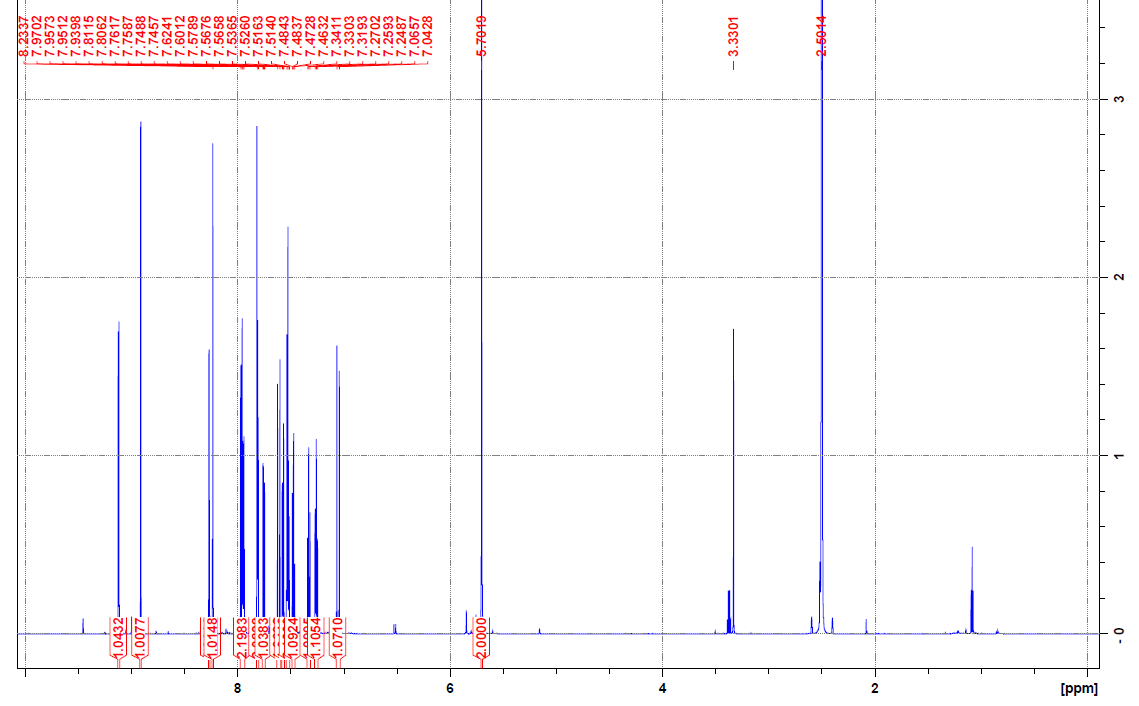

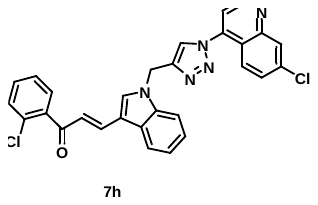

**Figure S.60A:** ^1^H NMR spectrum of **7h.**

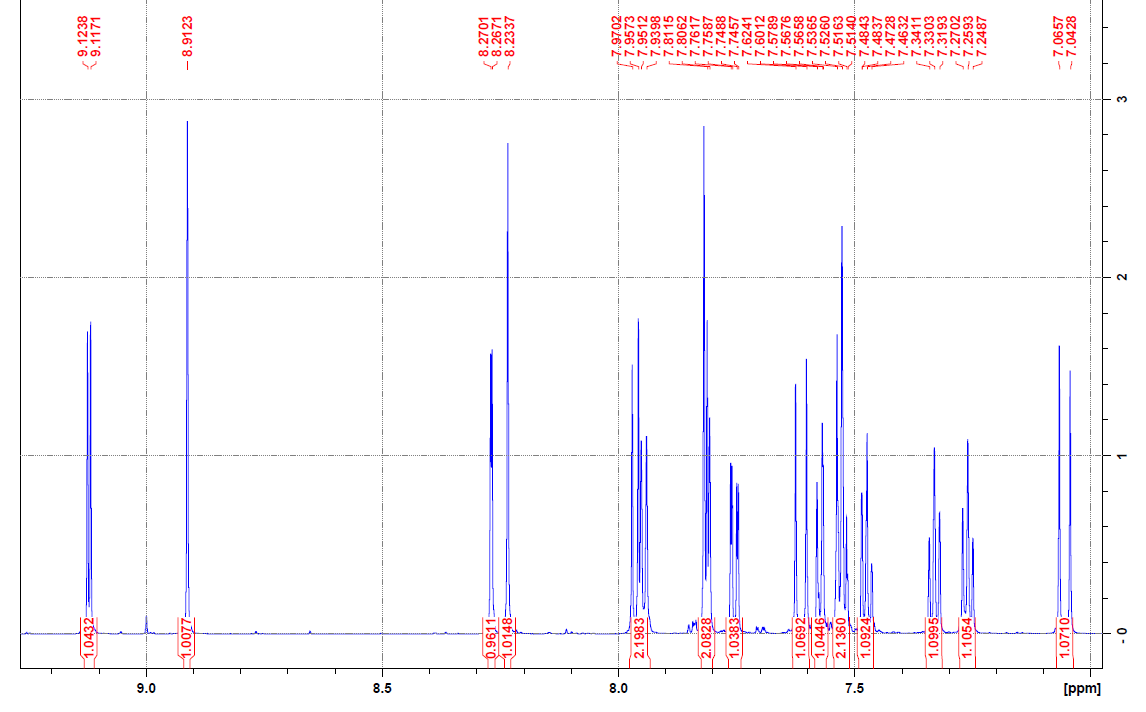

**Figure S.60B:** ^1^H NMR spectrum of **7h.**

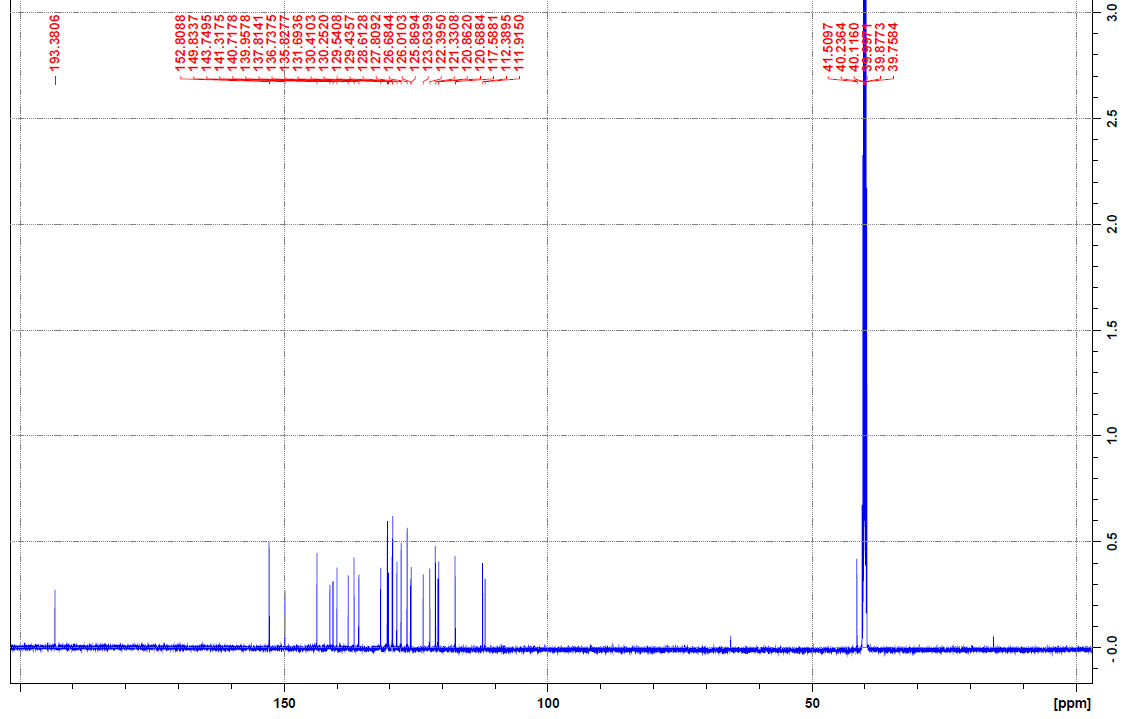

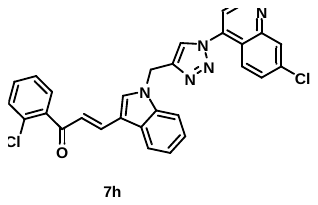

**Figure S.61:** ^13^ C NMR spectrum of **7h.**

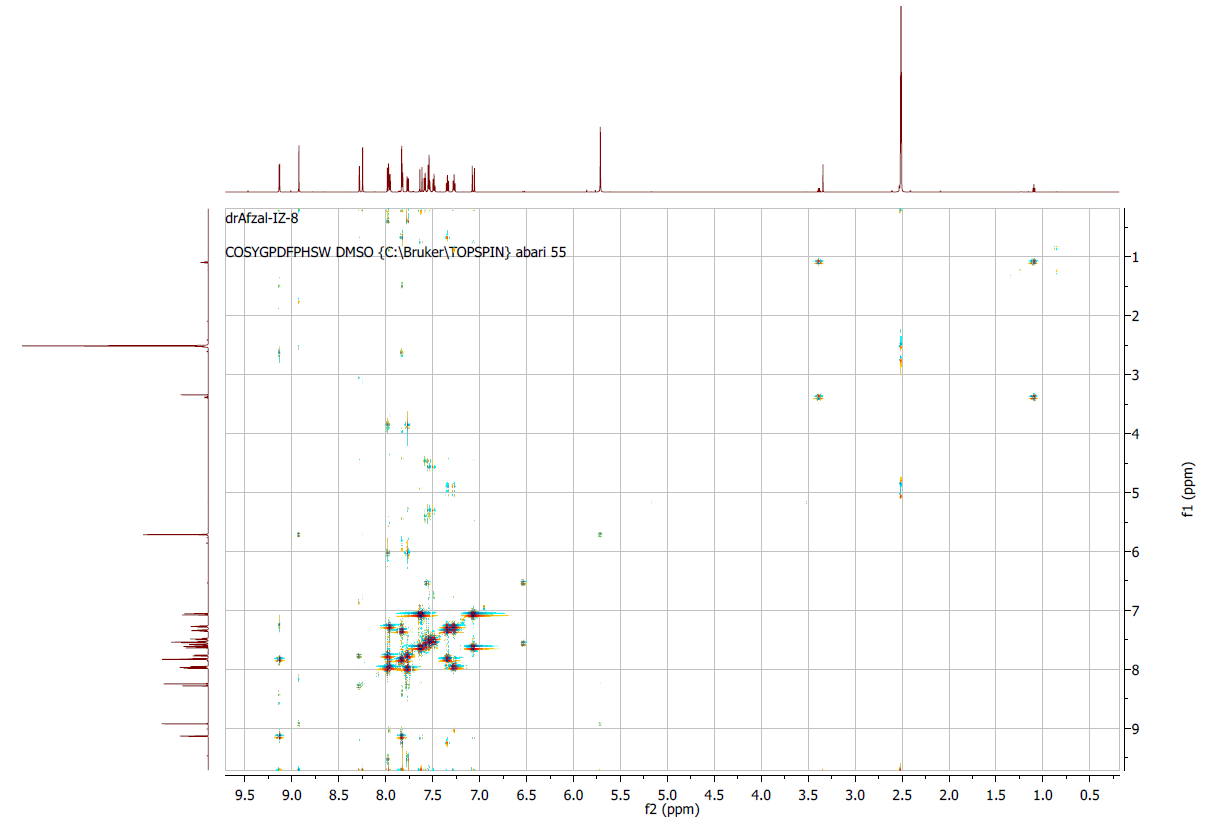

**Figure S.62:** 2D-COSY analyses of **7h.**

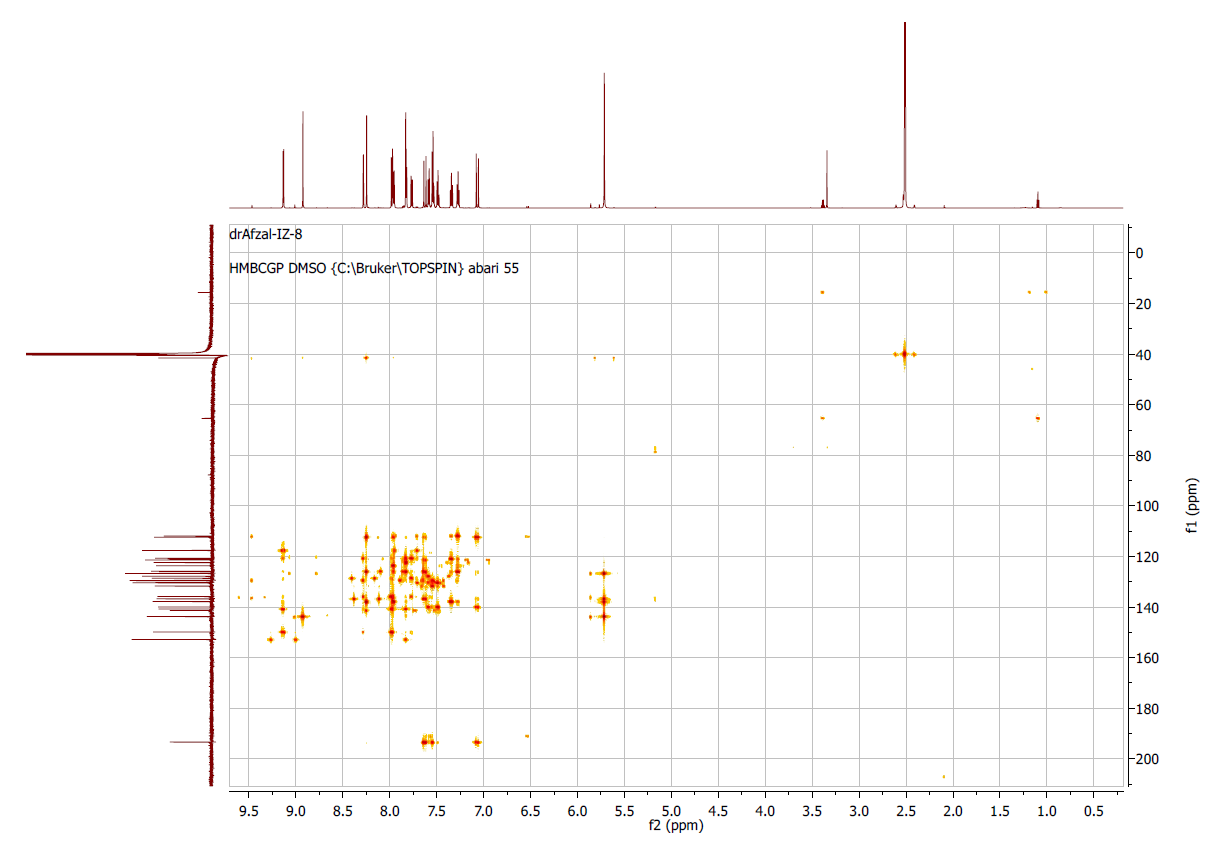

**Figure S.63:** 2D-HMBC analyses of **7h.**

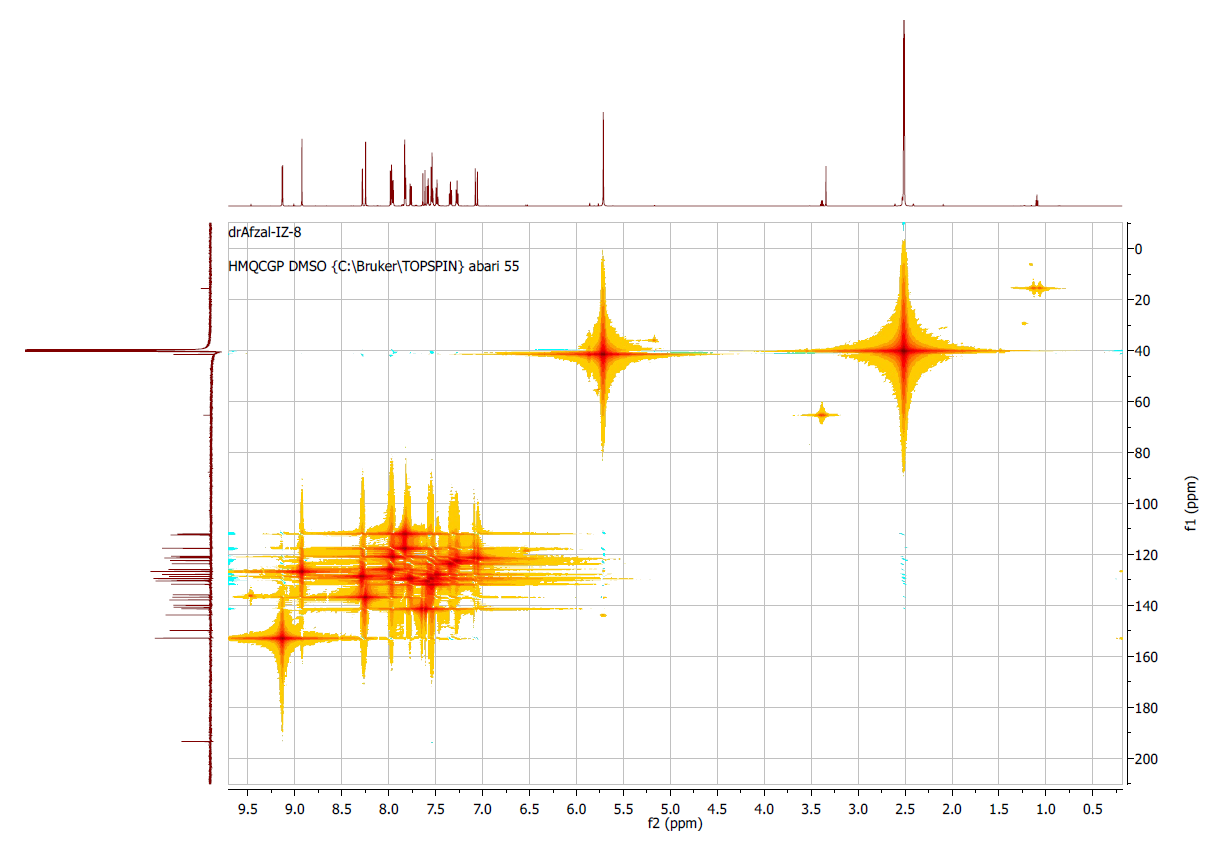

**Figure S.64:** 2D-HMQC analyses of **7h.**

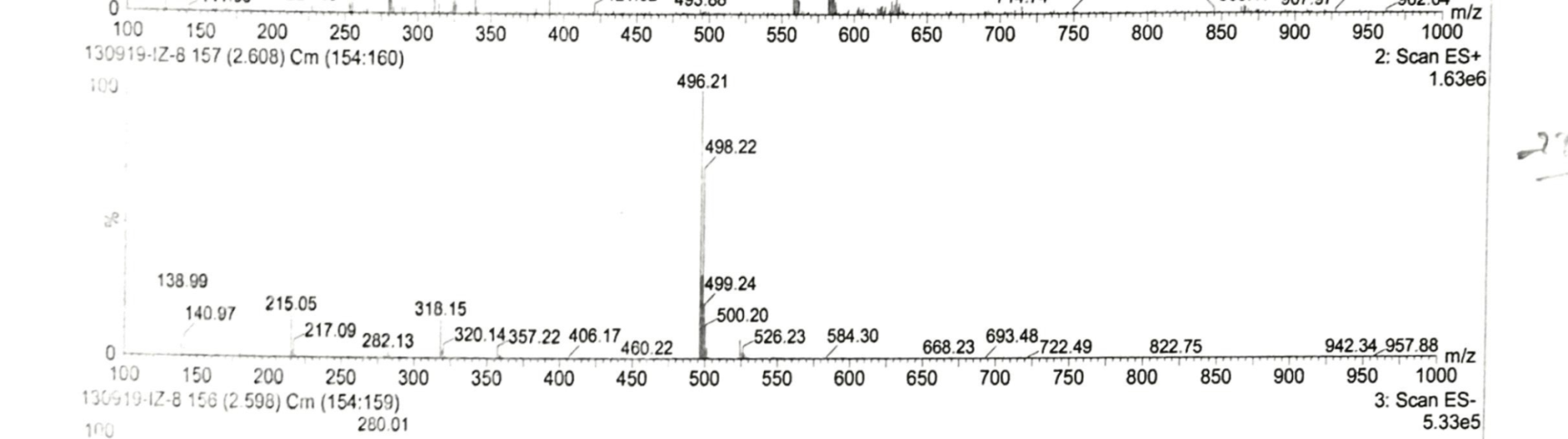

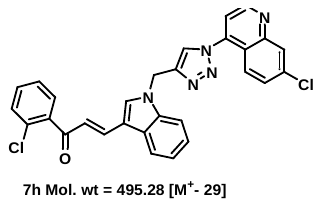

**Figure S.65:** Mass spectrum of **7h.**

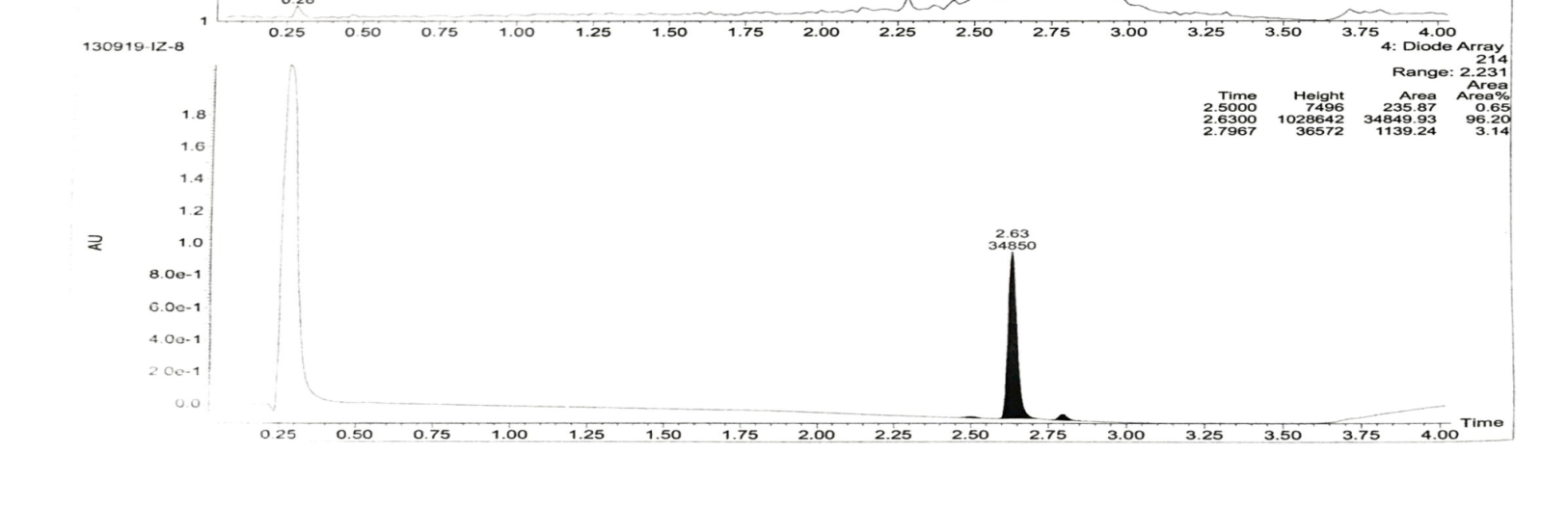

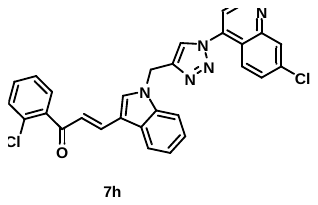

**Figure S.66:** Purity of **7h.**

**16.** (*E*)-1-(3-chlorophenyl)-3-(1-((1-(7-chloroquinolin-4-yl)-1H-1,2,3-triazol-4-yl)methyl)-1H-indol-3-yl)prop-2-en-1-one **(7i):**

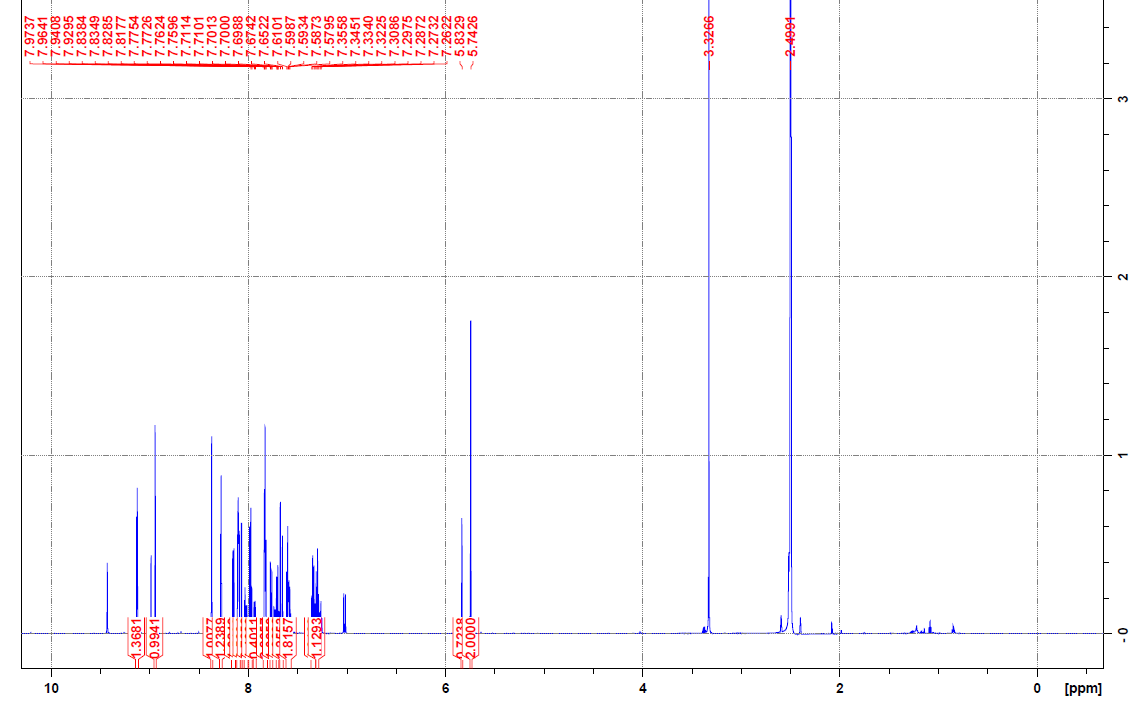

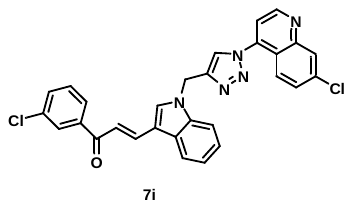

**Figure S.67A:** ^1^H NMR spectrum of **7i.**

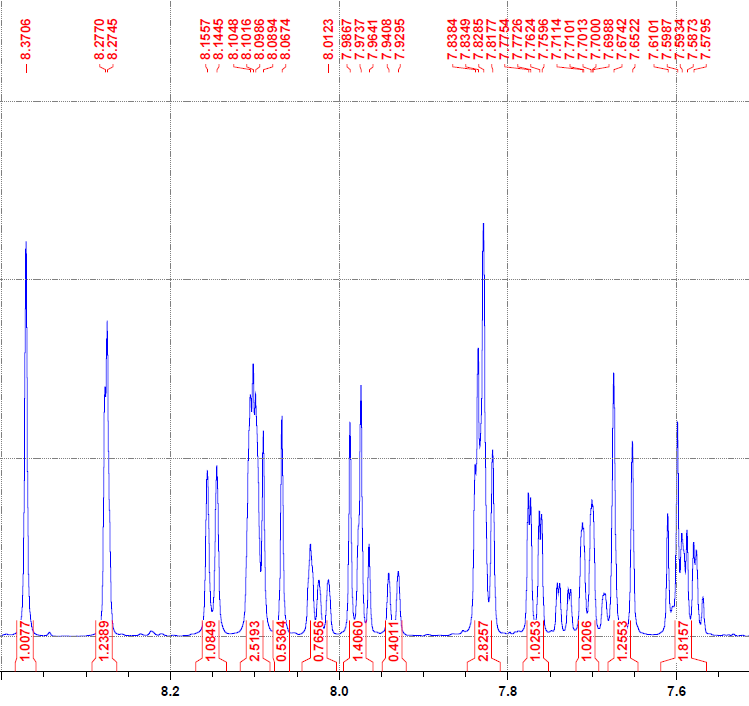

**Figure S.67B:** ^1^H NMR spectrum of **7i.**

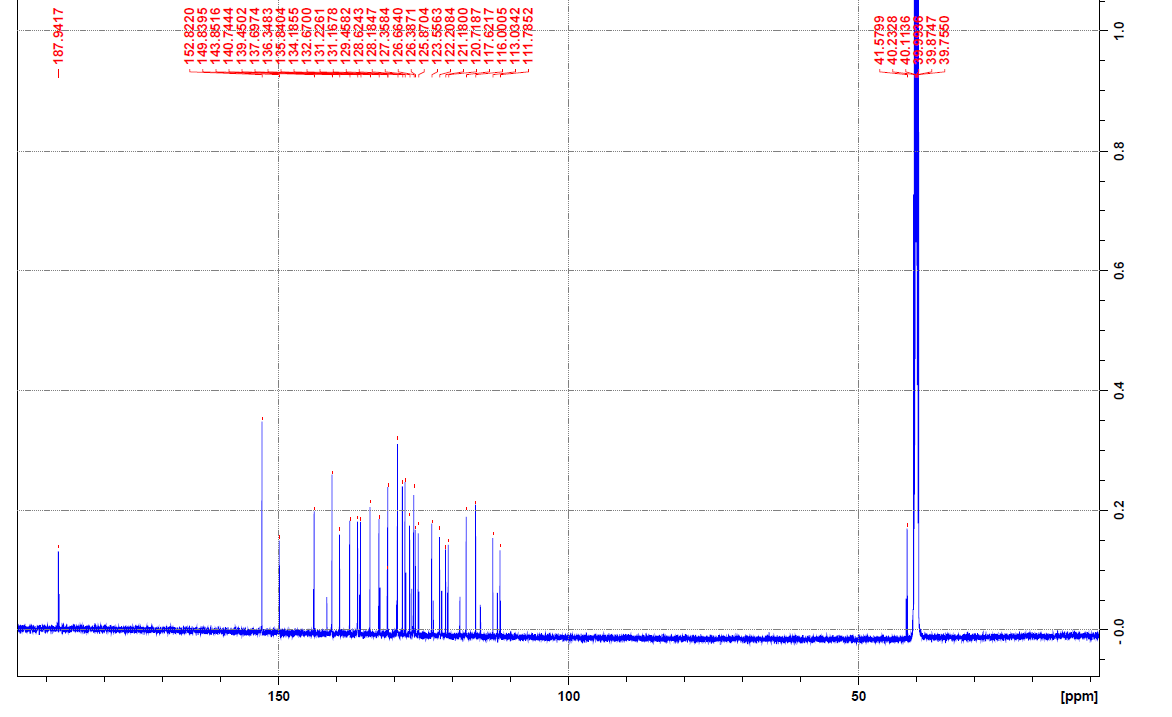

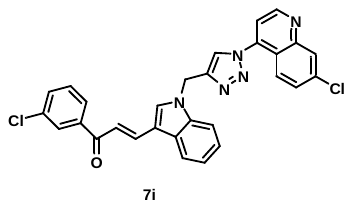

**Figure S.68:** ^13^ C NMR spectrum of **7i.**

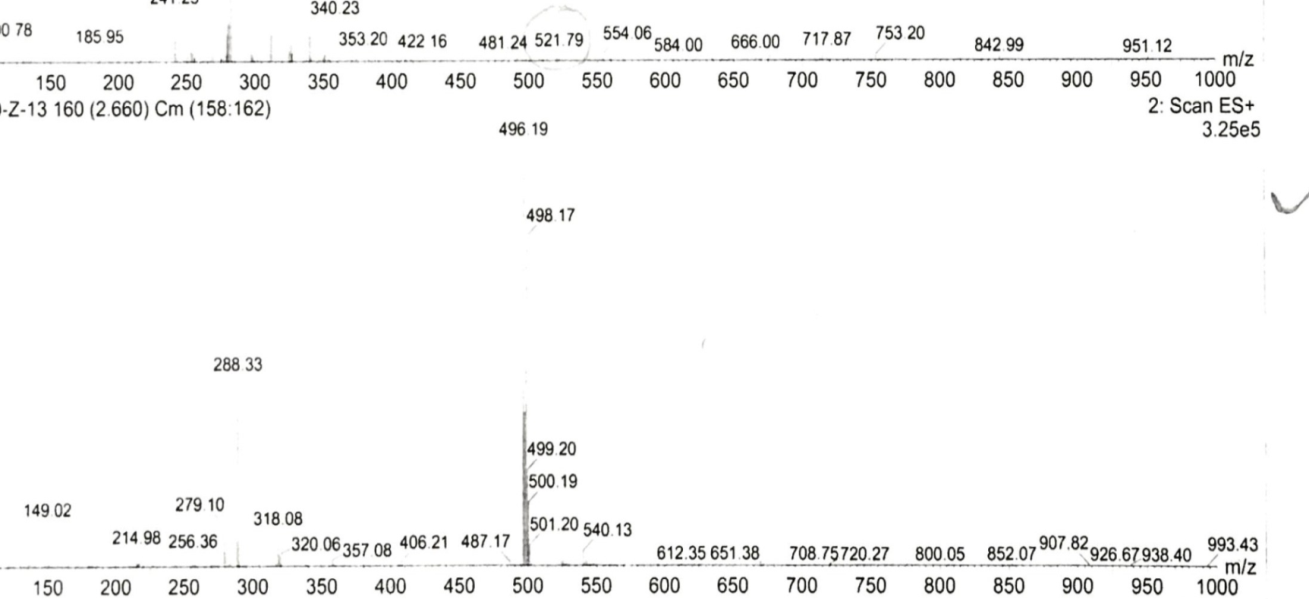

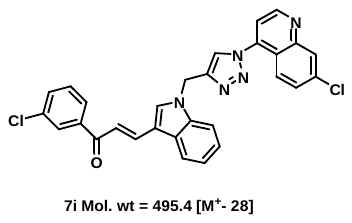

**.**

**Figure S.69:** Mass spectrum of **7i.**

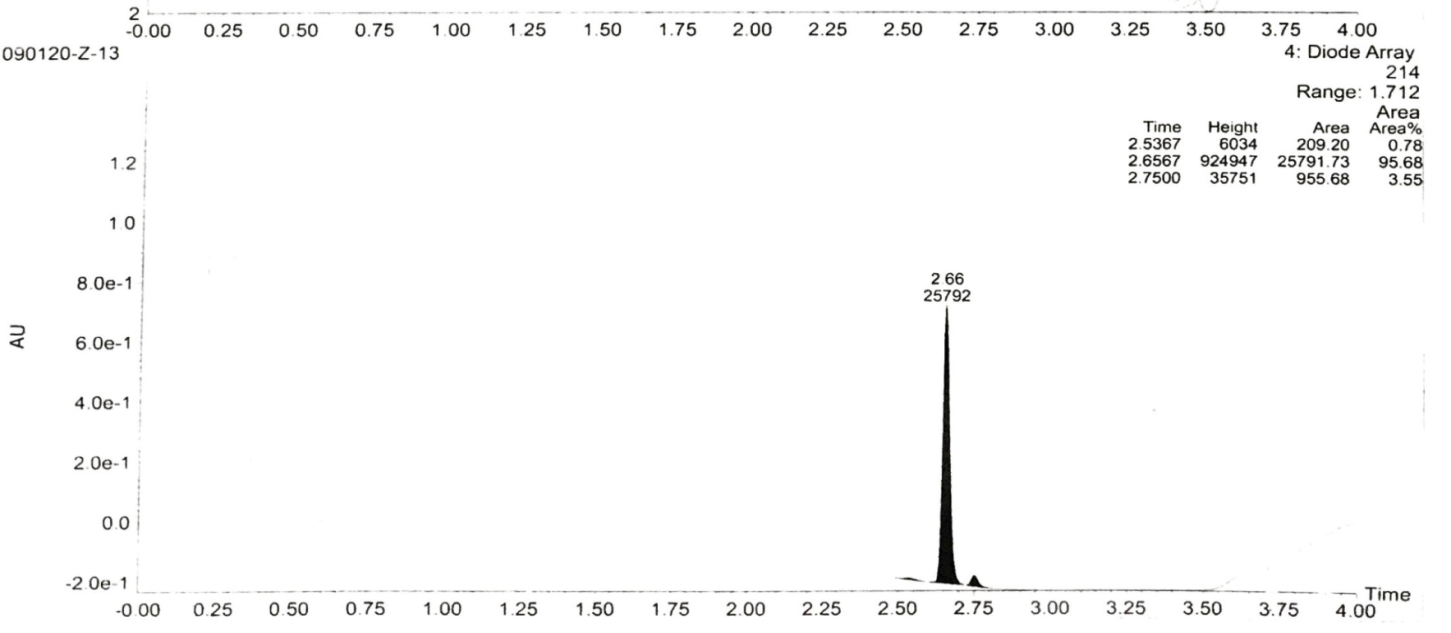

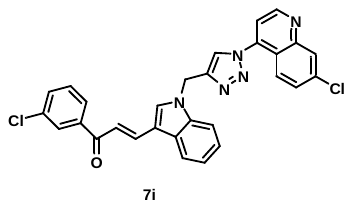

**Figure S.70:** Purity of **7i.**

**17.** (E)-3-(1-((1-(7-chloroquinolin-4-yl)-1H-1,2,3-triazol-4-yl)methyl)-1H-indol-3-yl)-1-(2-methoxyphenyl)prop-2-en-1-one **(7l**):

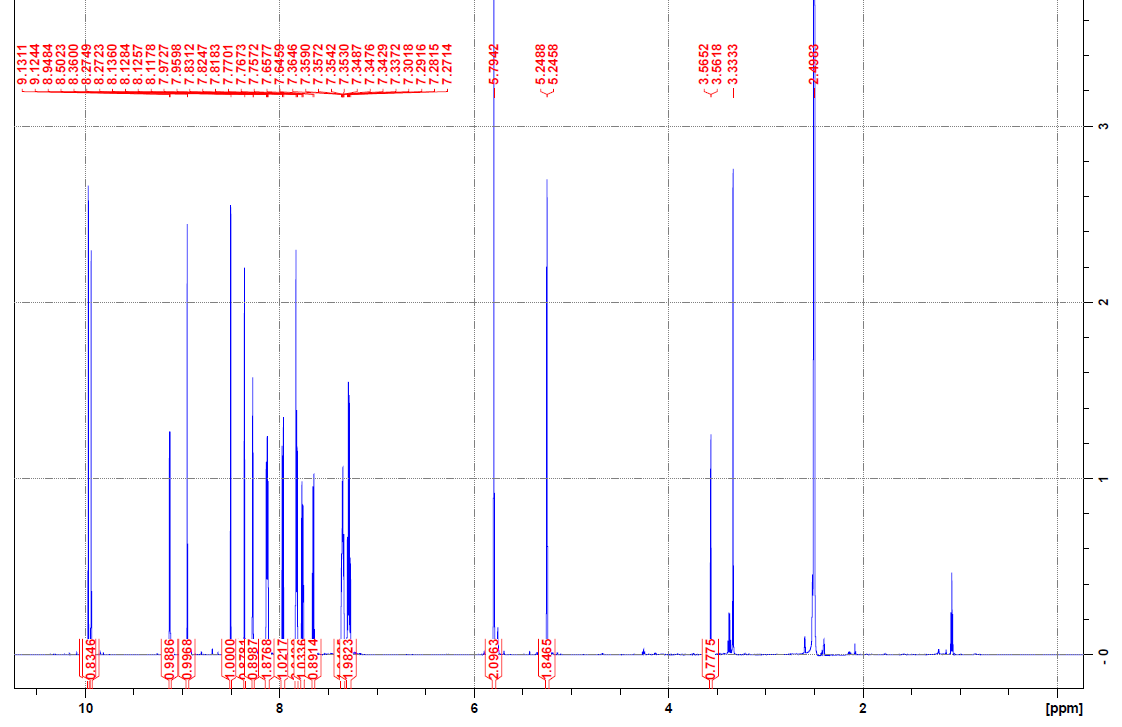

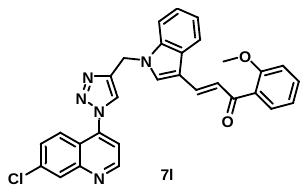

**Figure S.71A:** ^1^H NMR spectrum of **7l.**

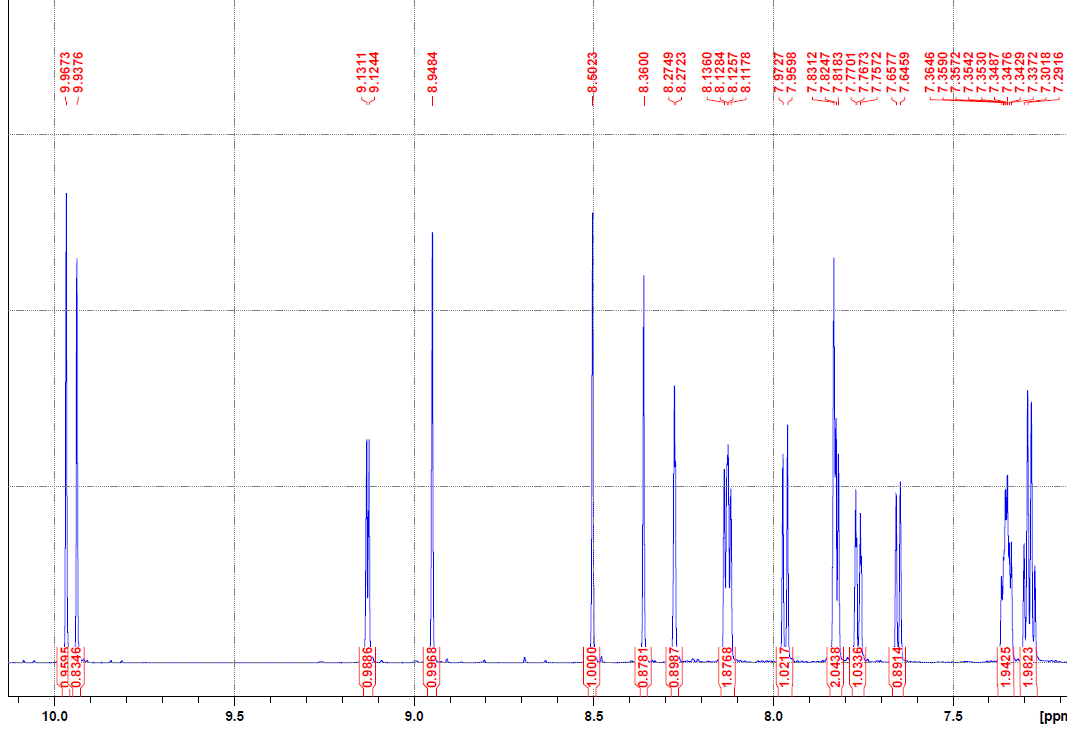

**Figure S.71B:** ^1^H NMR spectrum of **7l.**

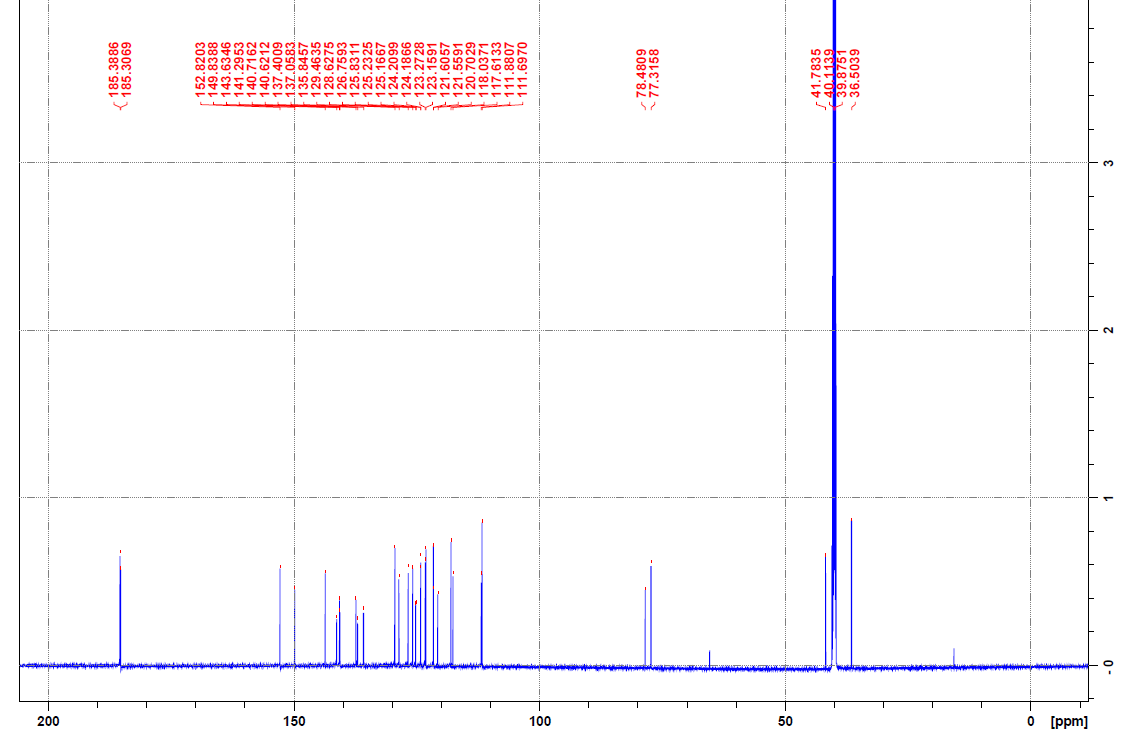

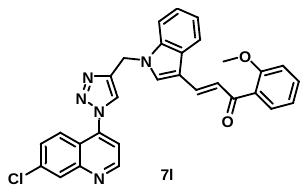

**Figure S.72:** ^13^C NMR spectrum of **7l.**

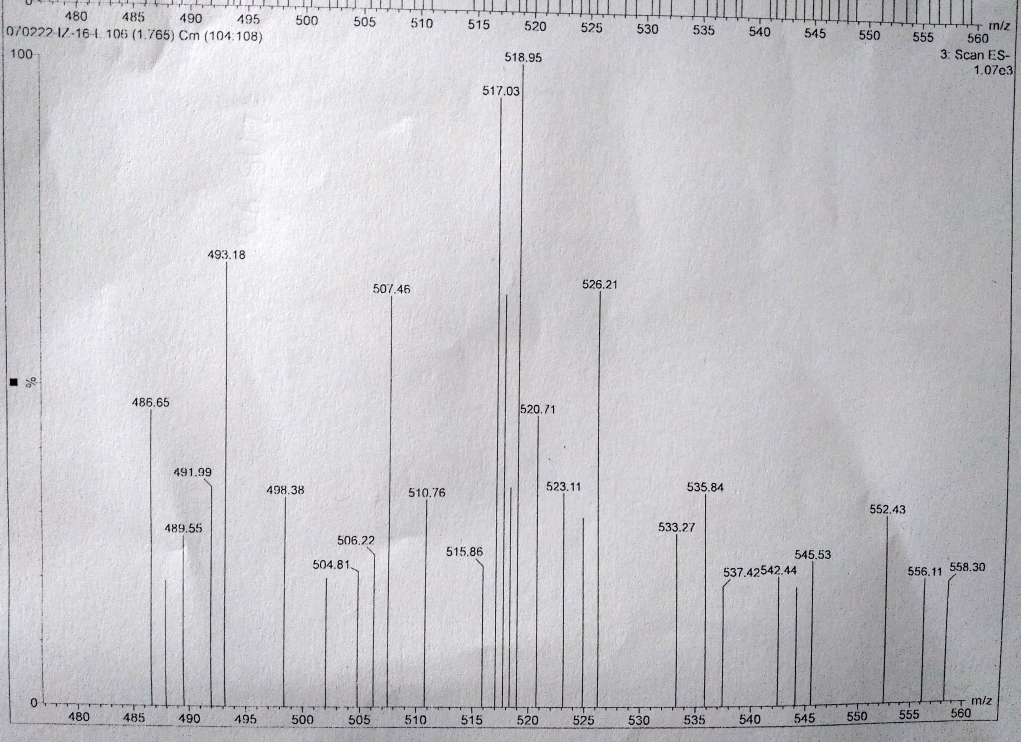

**Figure S.73:** Mass spectrum of **7l.**

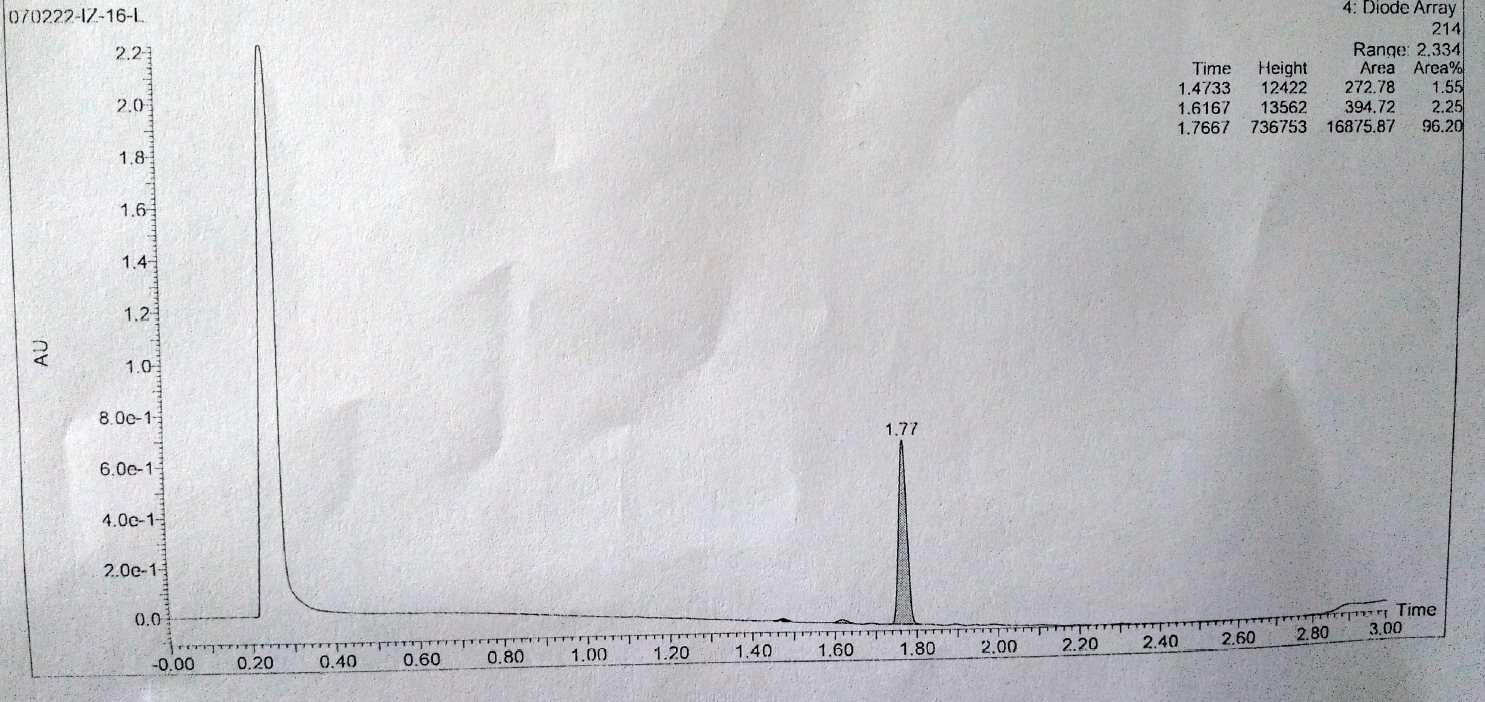

**Figure S.74:** Purity of **7l.**

**1**8**.** (*E*)-3-(1-((1-(7-chloroquinolin-4-yl)-1H-1,2,3-triazol-4-yl)methyl)-1H-indol-3-yl)-1-(3,4-dimethoxyphenyl)prop-2-en-1-one **(7o):**

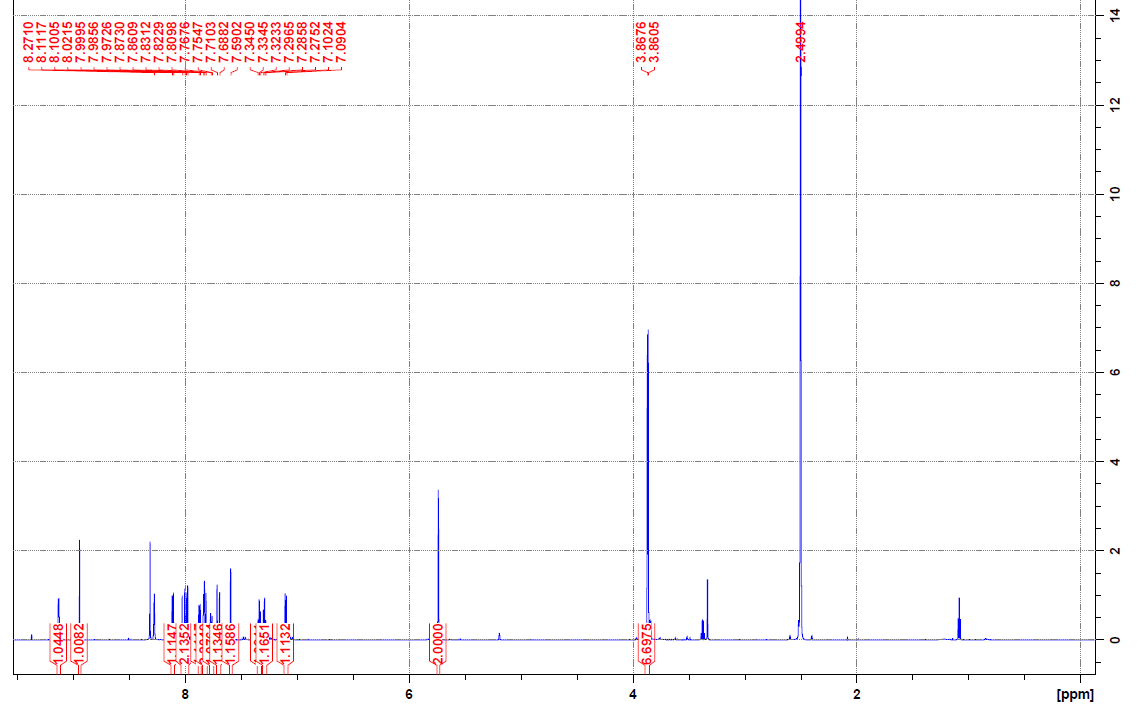

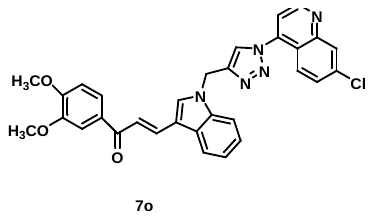

**Figure S.75A:** ^1^H NMR spectrum of **7o.**

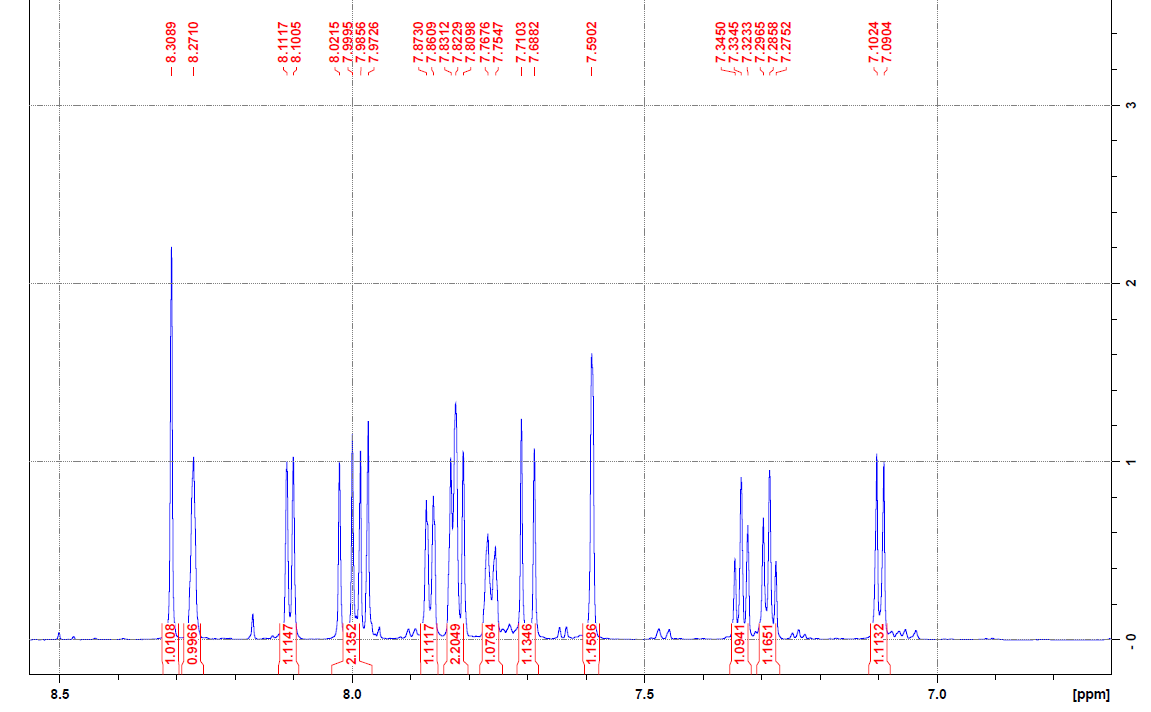

**Figure S.75B:** ^1^H NMR spectrum of **7o.**

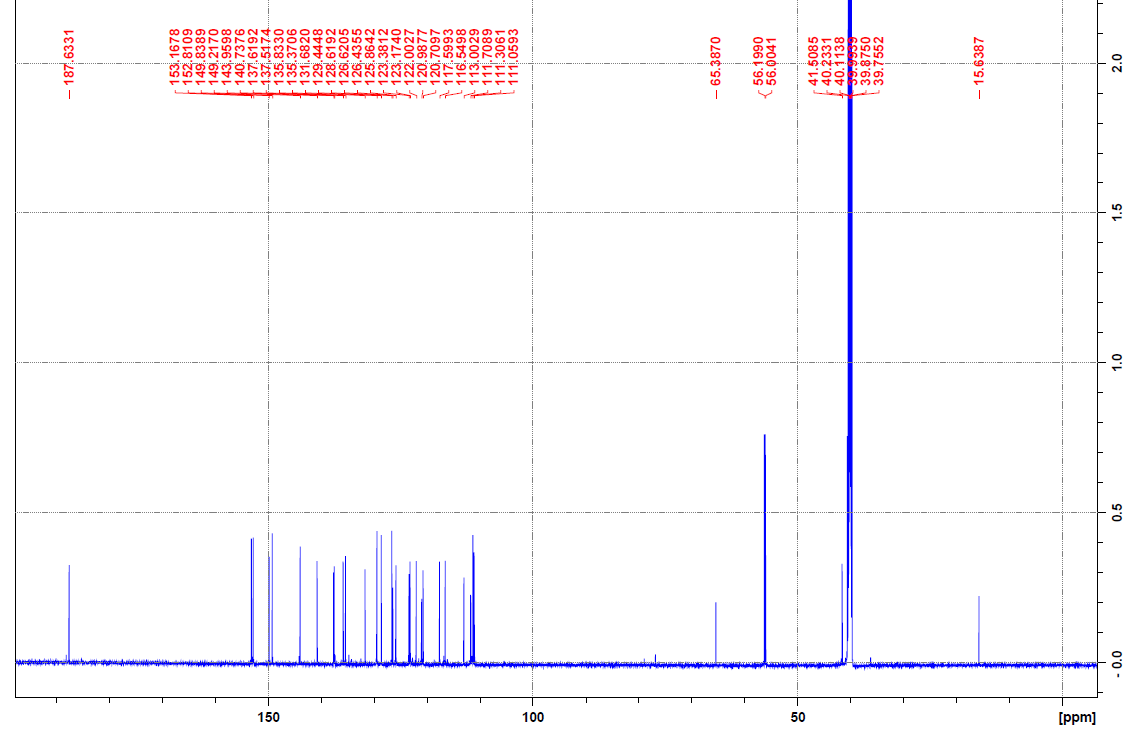

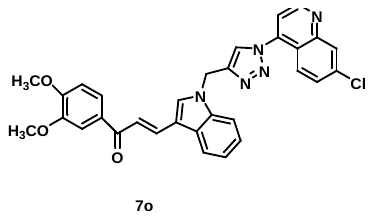

**Figure S.76:** ^13^ C NMR spectrum of **7o.**

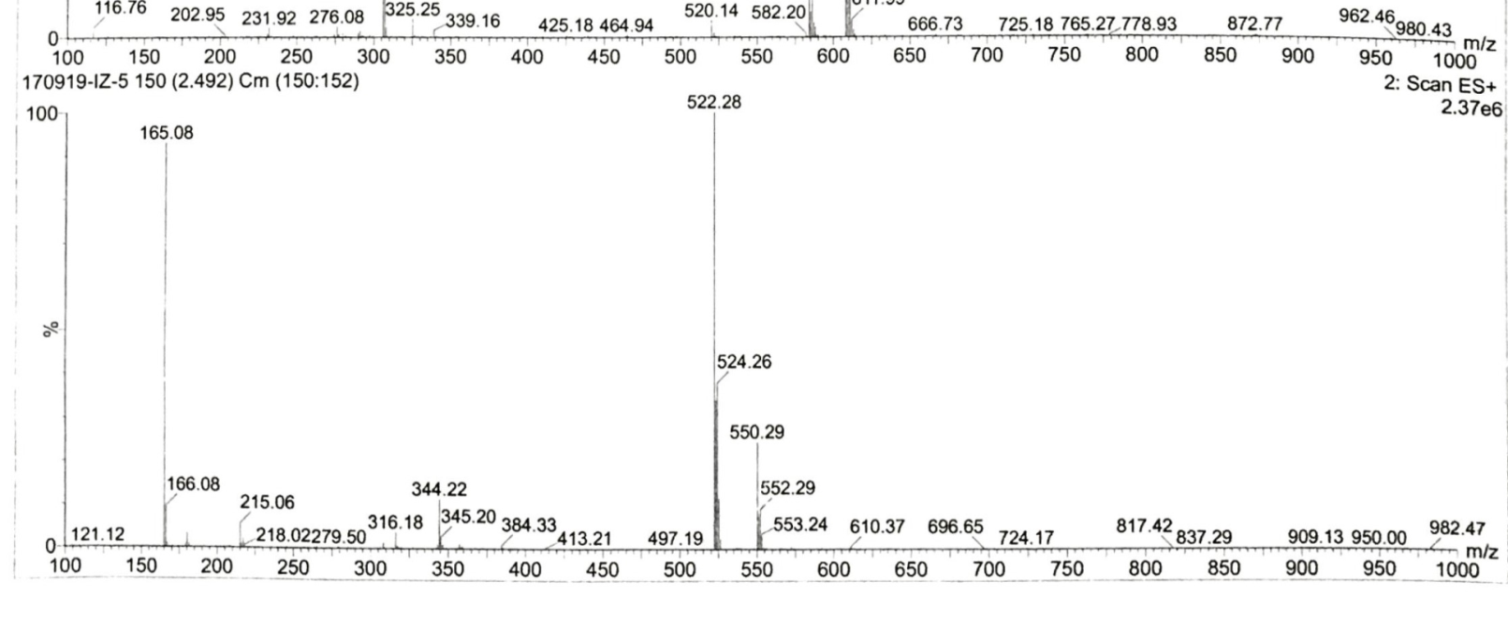

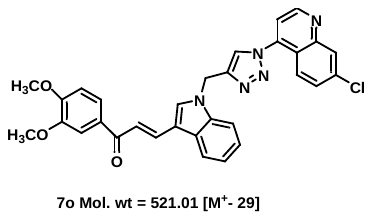

**Figure S.77:** Mass spectrum of **7o.**

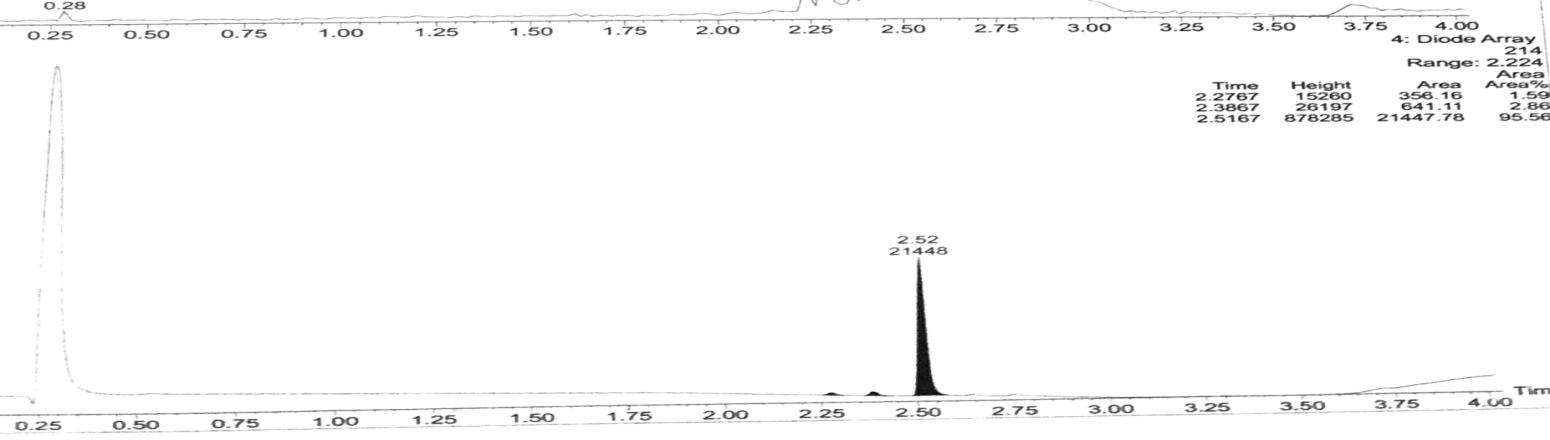

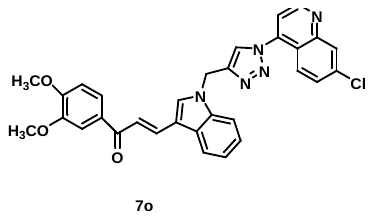

**Figure S.78:** Purity of **7o.**

**18.** (*E*)-3-(1-((1-(7-chloroquinolin-4-yl)-1H-1,2,3-triazol-4-yl)methyl)-1H-indol-3-yl)-1-(thiophen-2-yl)prop-2-en-1-one **(7p):**

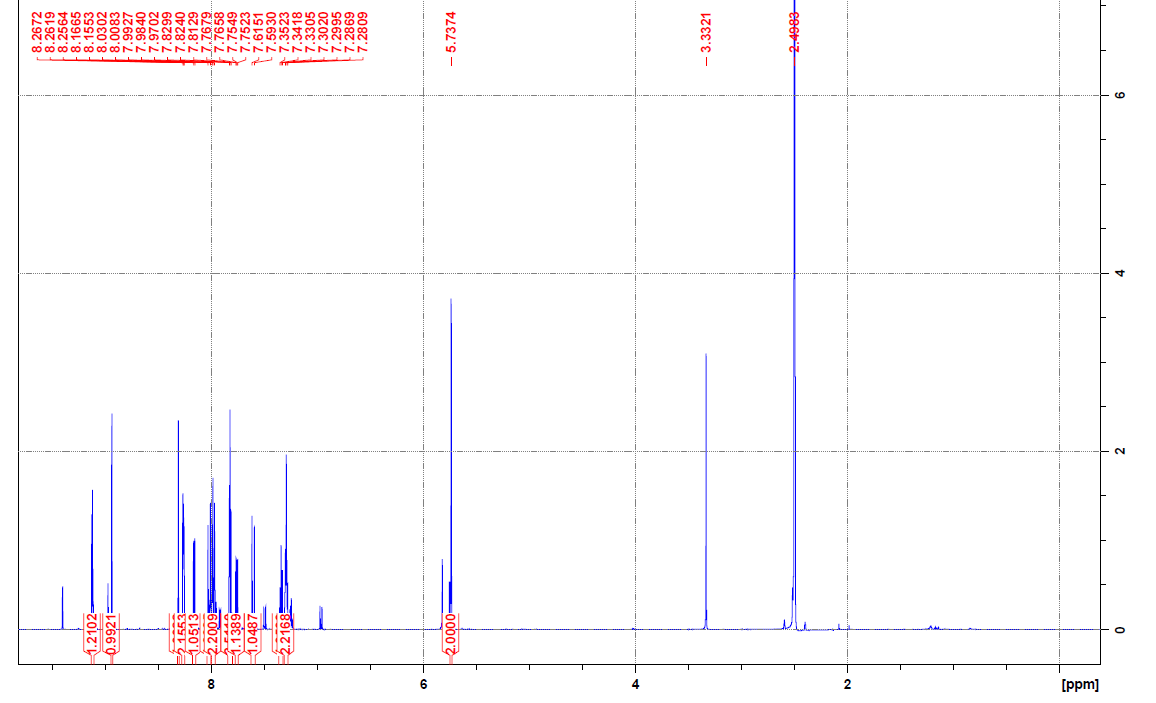

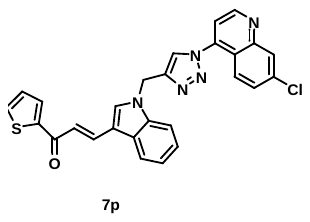

**Figure S.79A:** ^1^H NMR spectrum of **7p.**

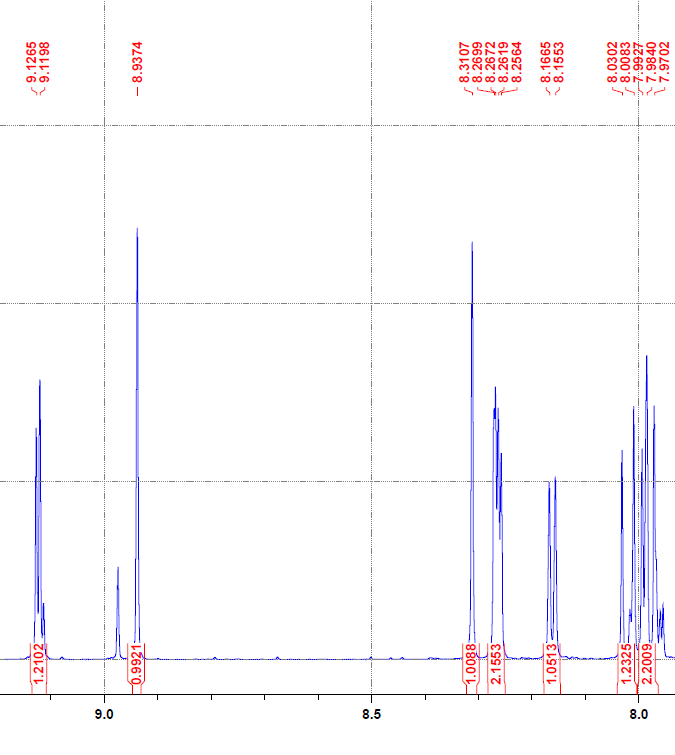

**Figure S.79B:** ^1^H NMR spectrum of **7p.**

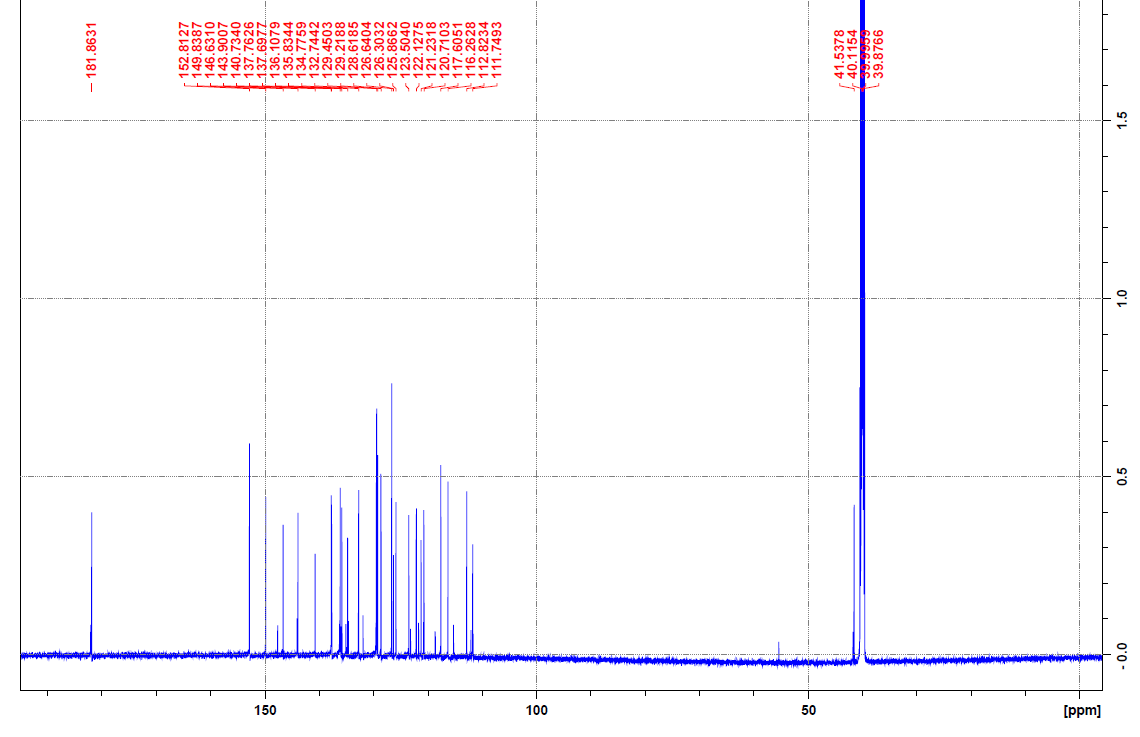

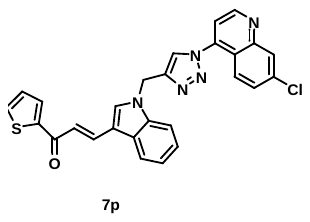

**Figure S.80:** ^13^ C NMR spectrum of **7p.**

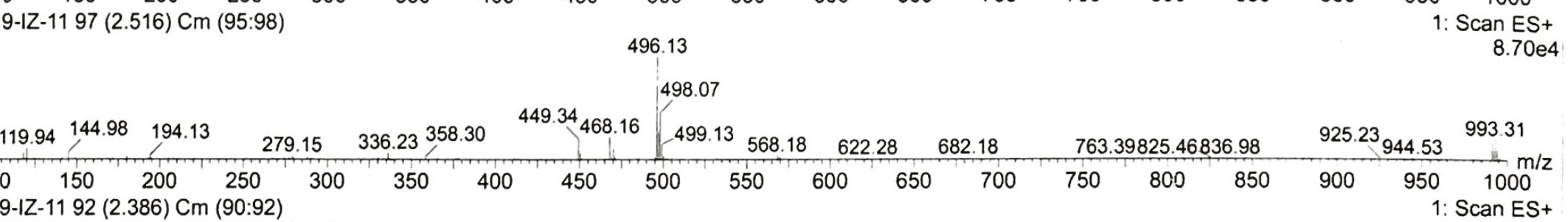

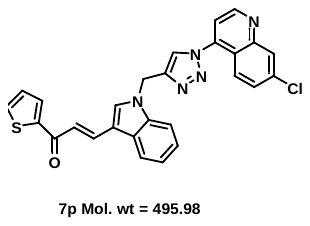

**Figure S.81:** Mass spectrum of **7p.**

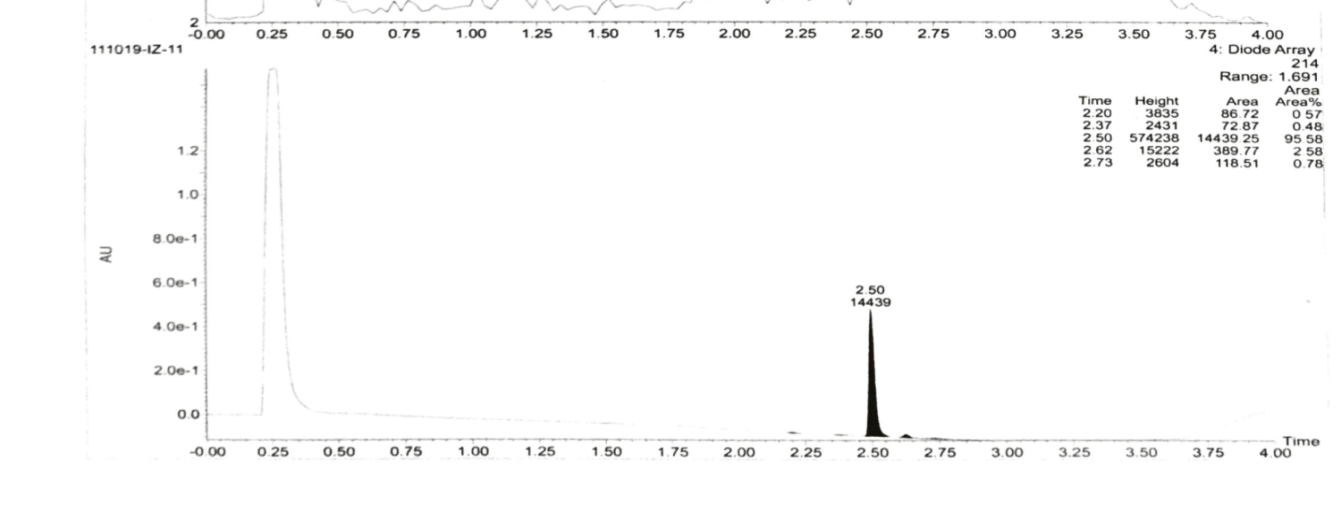

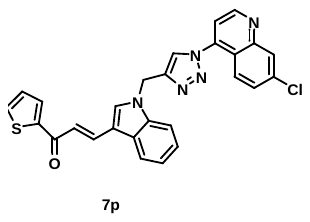

**Figure S82:** Purity of **7p.**

**19.** (*E*)-1-(3,5-bis(benzyloxy)phenyl)-3-(1-((1-(7-chloroquinolin-4-yl)-1H-1,2,3-triazol-4-yl)methyl)-1H-indol-3-yl)prop-2-en-1-one **(7q):**

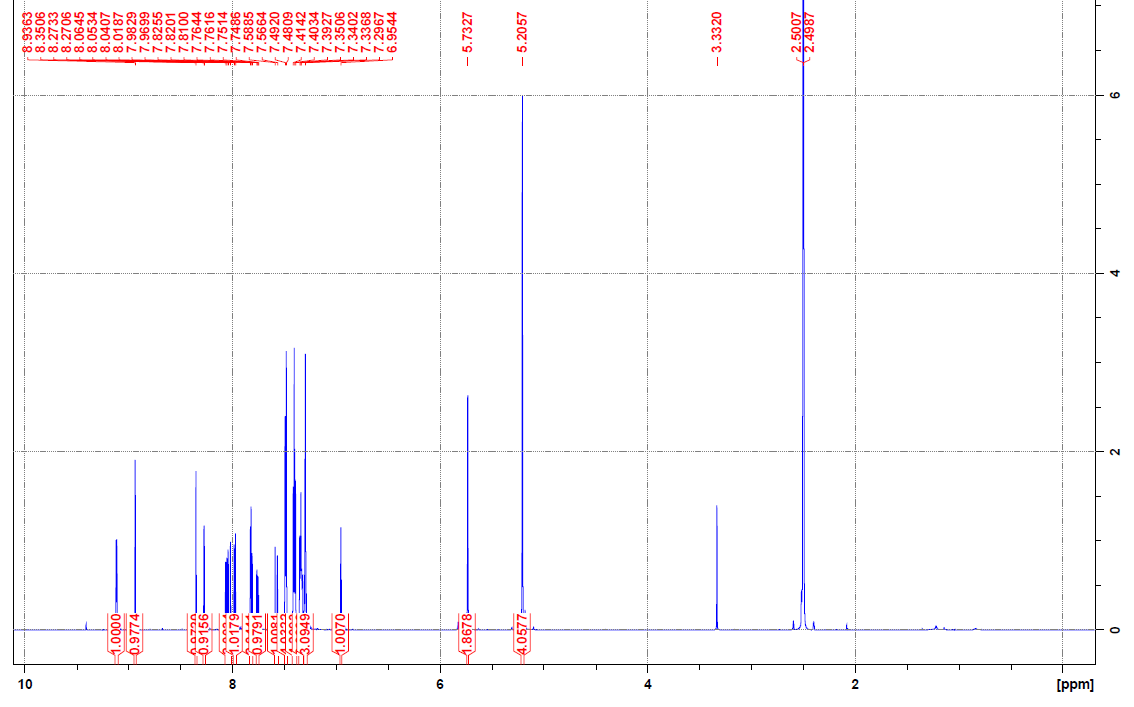

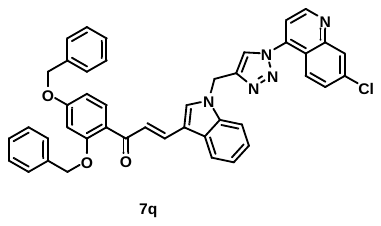

**Figure S.83A:** ^1^H NMR spectrum of **7q.**

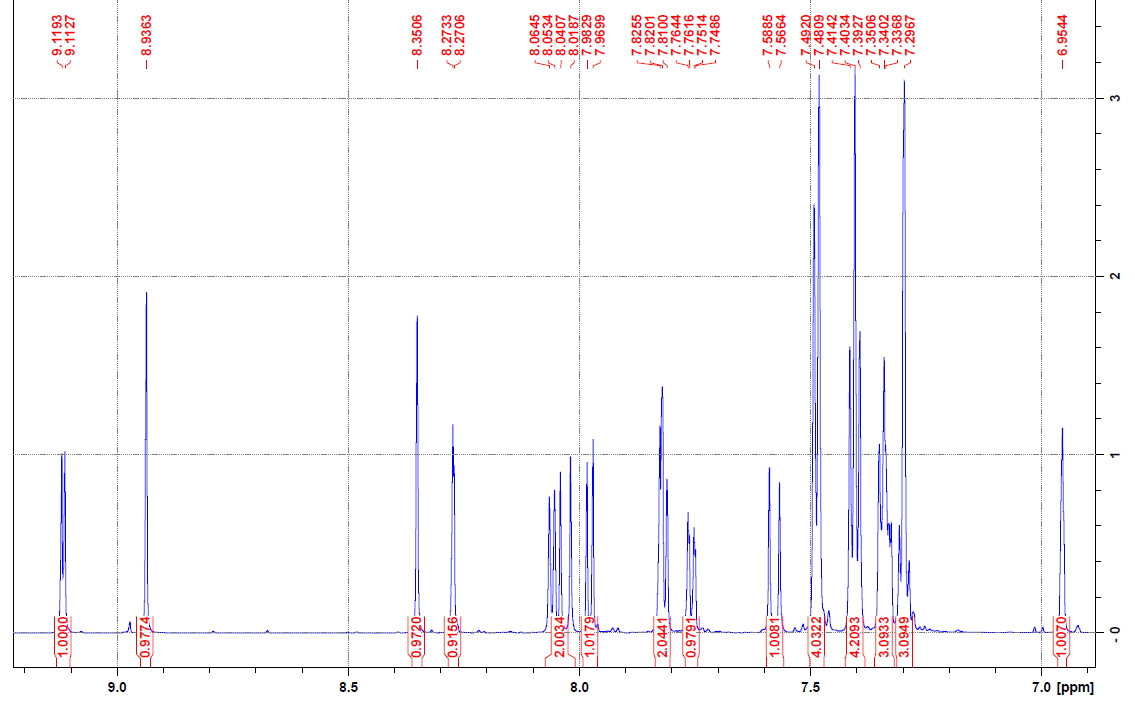

**Figure S.83B:** ^1^H NMR spectrum of **7q.**

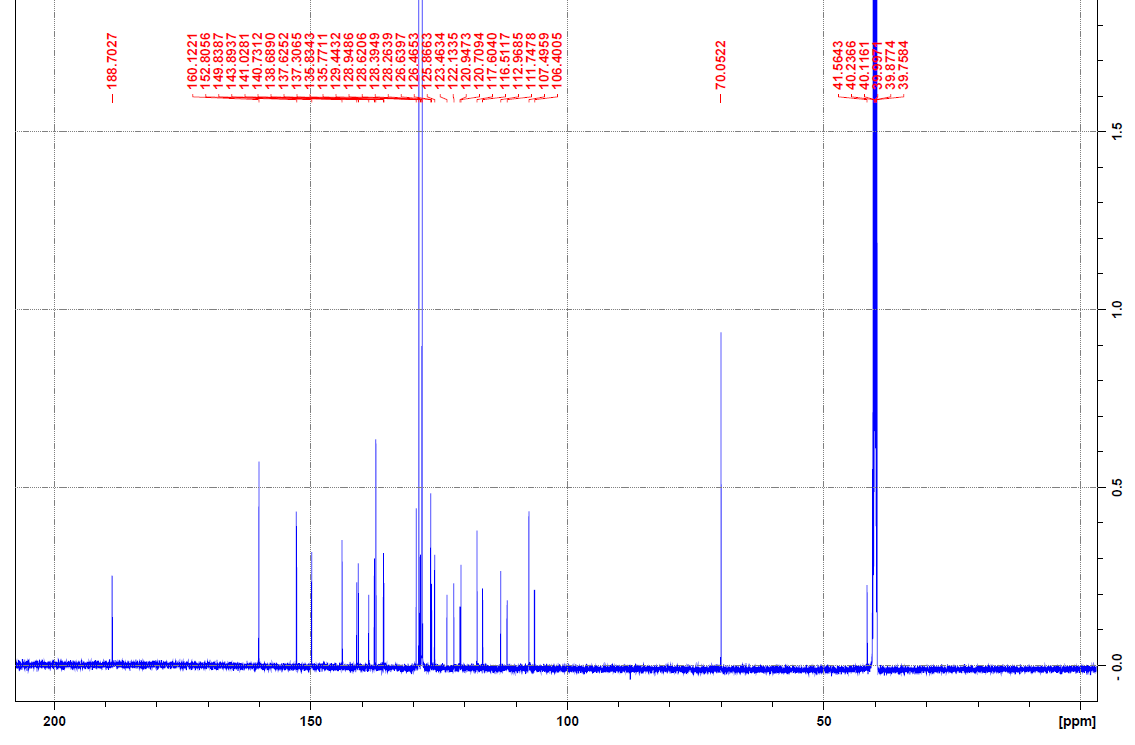

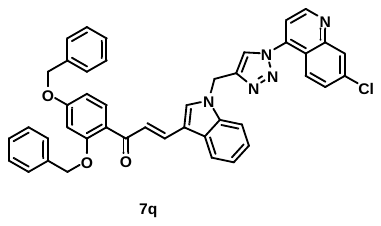

**Figure S.84:** ^13^C NMR spectrum of **7q.**

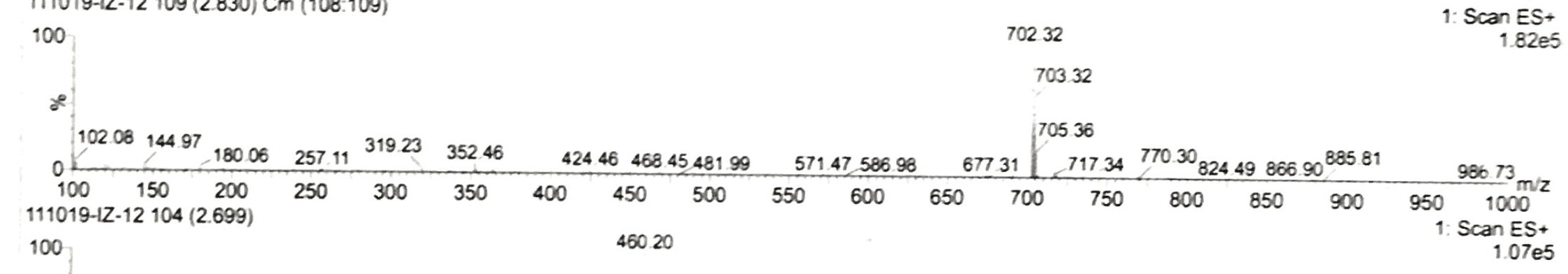

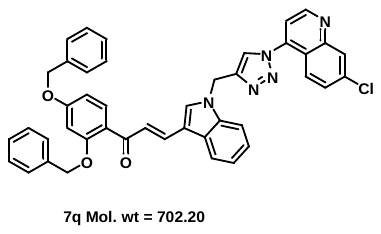

**Figure S.85:** Mass spectrum of **7q.**

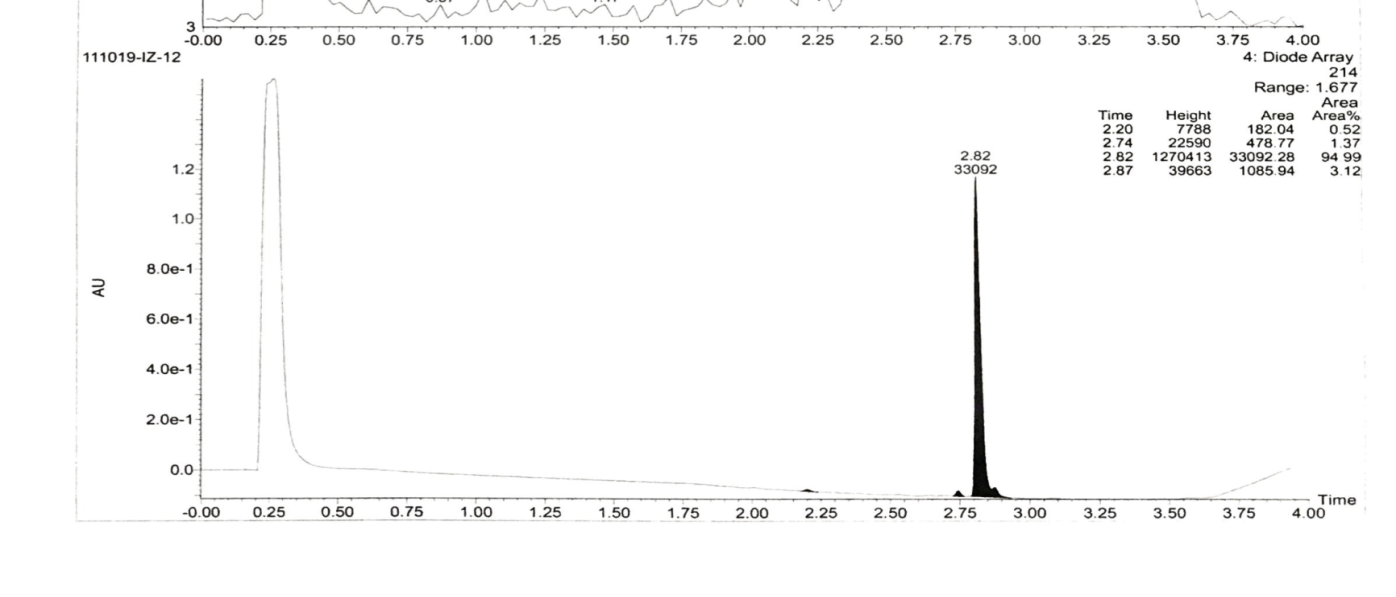

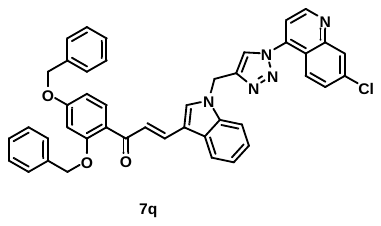

**Figure S.86:** Purity of **4q.**

**20.** (E)-3-(1-((1-(7-chloroquinolin-4-yl)-1H-1,2,3-triazol-4-yl)methyl)-1H-indol-3-yl)-1-o-tolylprop-2-en-1-one **(7r):**

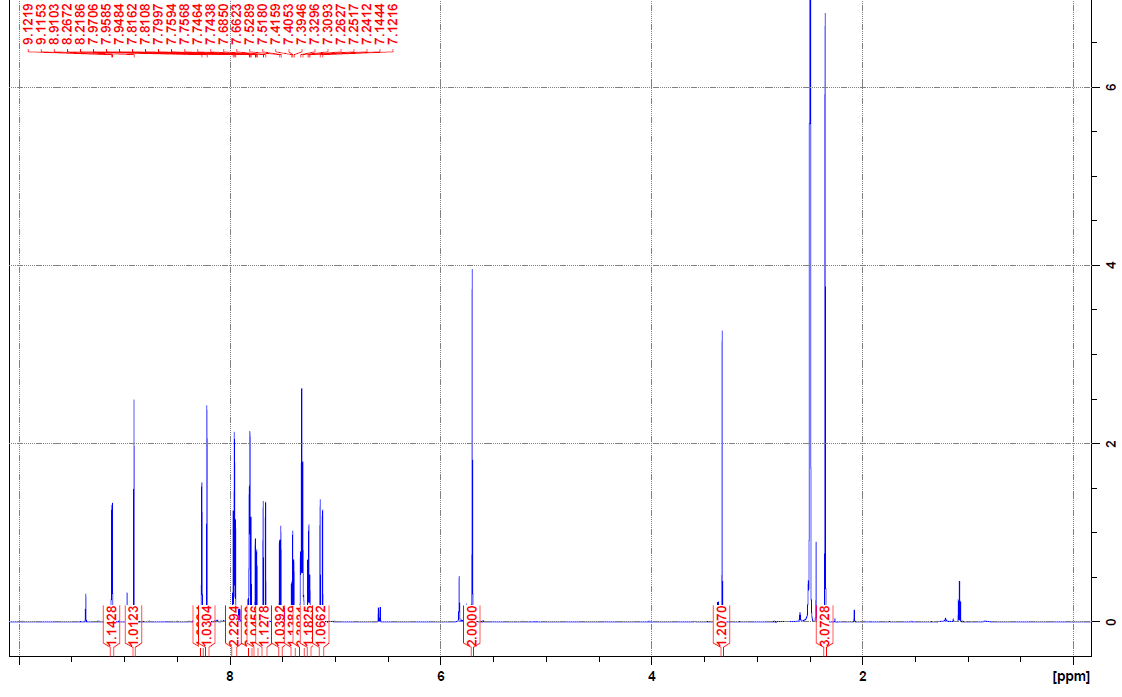

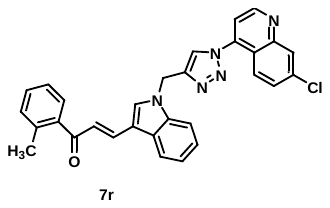

**Figure S.87A:** ^1^H NMR spectrum of **7r.**

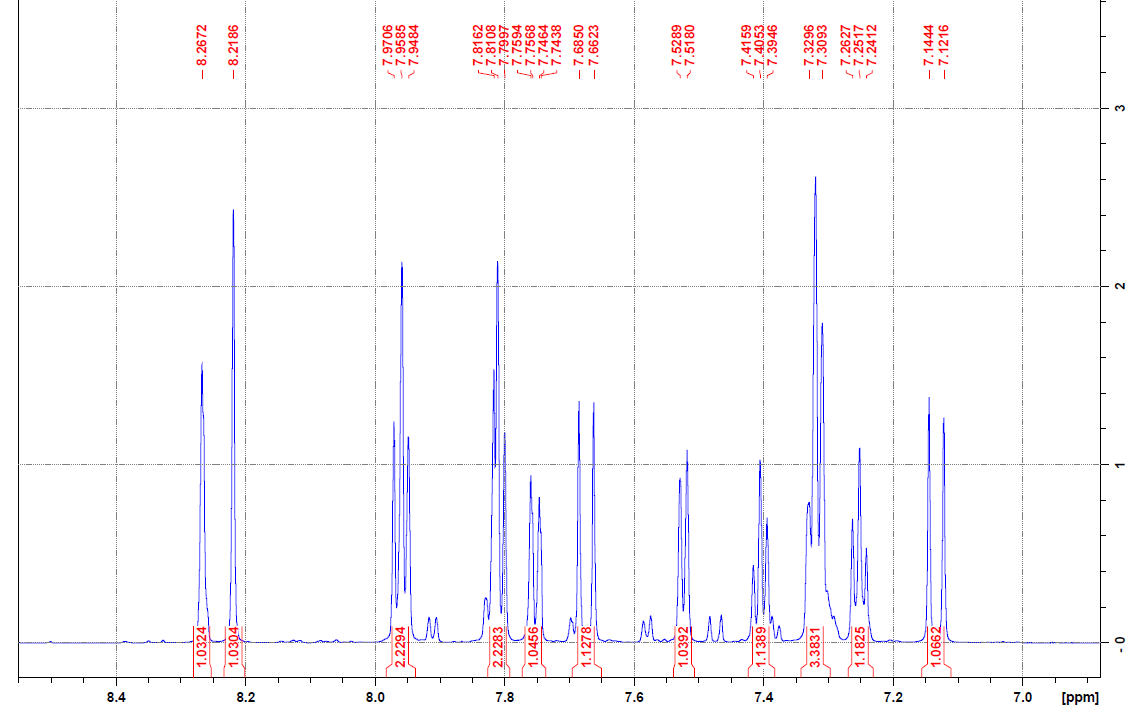

**Figure S.87B:** ^1^H NMR spectrum of **7r.**

**Figure S.88:** ^13^C NMR spectrum of **7r.**

**Figure S.89:** Mass spectrum of **7r.**

**Figure S.90:** Purity of **7r.**

**21.** 1-((1-(7-chloroquinolin-4-yl)-1H-1,2,3-triazol-4-yl)methyl)-1H-indole-3-carbaldehyde **(9):**

**Figure S.91A:** ^1^H NMR spectrum of **9.**

**Figure S.91B:** ^1^H NMR spectrum of **9.**

**Figure S.92:** ^13^C NMR spectrum of **9.**

**Figure S.93:** Mass spectrum of **9.**

**Figure S.94:** Purity of **9.**

**Fig. S.95: Possible fragmentation pattern in the mass spectrometry:**

The characteristic peaks observed within the mass spectra of triazole derivative **7f** are summarized in **Fig. S95**. This mass spectra exhibit molecular ion peaks and its fragmentation, that confirm the structure of all compounds. Similar trend was found for all compounds. Following is the mass fragmentation of compound **7f**.

**Fig. S.95:** Proposed fragmentation pathway of the protonated ion of the compound **7f**.

**Growth Inhibition assay of 7a-s and 9**

**Fig. S.96:** *In vitro* growth inhibition of **7a-s** and **9** at two concentrations (5 and 10 *µ*M) against 3D7 strains of *P. falciparum*.

**Table S1:** Physicochemical properties of **7a-s** and **9** compounds

| **Compound Id.** | **Mol. Weight** | **Heavy atoms** | **RB** | **HBA** | **HBD** | **TPSA** | **Log P** | **GI absorption** | **Lipinski violations** | **PAINS alerts** | **Carcinogen** |
| --- | --- | --- | --- | --- | --- | --- | --- | --- | --- | --- | --- |
| 7a | 489.96 | 36 | 6 | 4 | 0 | 65.6 | 3.91 | High | 0 | 0 | NC |
| 7b | 534.95 | 39 | 7 | 6 | 0 | 111.42 | 3.58 | Low | 1 | 0 | NC |
| 7c | 534.95 | 39 | 7 | 6 | 0 | 111.42 | 3.52 | Low | 1 | 0 | NC |
| 7d | 534.95 | 39 | 7 | 6 | 0 | 111.42 | 3.57 | Low | 1 | 0 | NC |
| 7e | 568.85 | 37 | 6 | 4 | 0 | 65.6 | 4.29 | Low | 2 | 0 | NC |
| 7f | 568.85 | 37 | 6 | 4 | 0 | 65.6 | 4.37 | Low | 2 | 0 | NC |
| 7g | 568.85 | 37 | 6 | 4 | 0 | 65.6 | 4.5 | Low | 2 | 0 | NC |
| 7h | 524.4 | 37 | 6 | 4 | 0 | 65.6 | 4.2 | Low | 2 | 0 | NC |
| 7i | 524.4 | 37 | 6 | 4 | 0 | 65.6 | 4.36 | Low | 2 | 0 | NC |
| 7j | 524.4 | 37 | 6 | 4 | 0 | 65.6 | 4.28 | Low | 2 | 0 | NC |
| 7k | 558.85 | 38 | 6 | 4 | 0 | 65.6 | 4.45 | Low | 2 | 0 | NC |
| 7l | 519.98 | 38 | 7 | 5 | 0 | 74.83 | 4.29 | High | 1 | 0 | NC |
| 7m | 519.98 | 38 | 7 | 5 | 0 | 74.83 | 4.42 | High | 1 | 0 | NC |
| 7n | 519.98 | 38 | 7 | 5 | 0 | 74.83 | 4.45 | High | 1 | 0 | NC |
| 7o | 550.01 | 40 | 8 | 6 | 0 | 84.06 | 4.25 | High | 1 | 0 | NC |
| 7p | 495.98 | 35 | 6 | 4 | 0 | 93.84 | 4.07 | Low | 0 | 0 | NC |
| 7q | 702.2 | 52 | 12 | 6 | 0 | 84.06 | 5.8 | Low | 2 | 0 | NC |
| 7r | 503.98 | 37 | 6 | 4 | 0 | 65.6 | 4.15 | High | 2 | 0 | NC |
| 7s | 507.95 | 37 | 6 | 5 | 0 | 65.6 | 4.14 | Low | 2 | 0 | NC |
| 9 | 387.82 | 28 | 4 | 4 | 0 | 65.6 | 3.01 | High | 0 | 0 | NC |

RB = Rotable bond, HBA = H acceptor, HBD = H doner, TPSA = Total polar surface area. PAIN = Pan assay interference assay.
